# A two-point IC_50_ method for evaluating the biochemical potency of irreversible enzyme inhibitors

**DOI:** 10.1101/2020.06.25.171207

**Authors:** Petr Kuzmič

## Abstract

Irreversible (covalent) enzyme inhibitors cannot be unambiguously ranked for biochemical potency by using IC_50_ values determined at a single point in time, because the same IC_50_ value could originate either from relatively low initial binding affinity accompanied by high chemical reactivity, or the other way around. To disambiguate the potency ranking of covalent inhibitors, here we describe a data-analytic procedure relying on two separate IC_50_ values, determined at two different reaction times. In the case of covalent inhibitors following the two-step kinetic mechanism *E* + *I* ⇌ *E*·*I* → *EI*, the two IC_50_ values alone can be used to estimate both the inhibition constant (*K*_i_) as a measure of binding affinity and the inactivation rate constant (*k*_inact_) as a measure of chemical reactivity. In the case of covalent inhibitors following the one-step kinetic mechanism *E* + *I* → *EI*, a simple algebraic formula can be used to estimate the covalent efficiency constant (*k*_inact_/*K*_i_) from a single experimental value of IC_50_. The two simplifying assumptions underlying the method are (1) zero inhibitor depletion, which implies that the inhibitor concentrations are always significantly higher than the enzyme concentration; and (2) constant reaction rate in the uninhibited control assay. The newly proposed method is validated by using a simulation study involving 64 irreversible inhibitors with covalent efficiency constants spanning seven orders of magnitude.

## 1. Introduction

Many medicines currently in use to treat various human diseases and symptoms are enzyme inhibitors. Furthermore, many important drugs and drug candidates are irreversible covalent inhibitors [1–4], which express their pharmacological effect by forming a permanent chemical bond with the protein target. Probably the most well known representative of this class is acetylsalicylate, or Aspirin, an irreversible covalent inhibitor of cyclooxygenase. Evaluating the biochemical potency of irreversible inhibitors in the process of pre-clinical drug discovery is exceedingly challenging. Even the task of simply arranging a list of potential drug candidates in order of their biochemical potency presents a serious obstacle. The main challenge is that the overall biochemical potency of irreversible enzyme inhibitors has two distinct components, namely, binding affinity measured by the inhibition constant (*K*_i_) and chemical reactivity measured by the inactivation rate constant (*k*_inact_).

Medicinal chemists and pharmacologists involved in drug discovery are accustomed to expressing the biochemical potency of enzyme inhibitors primarily in terms of the IC_50_ [5]^1^. However, in the case of irreversible inhibitors, the IC_50_ by definition decreases over time, until at asymptotically infinite time it approaches one half of the enzyme concentration. Therefore, reporting the IC_50_ for an irreversible inhibitor without also specifying the corresponding reaction time is essentially meaningless. An even more serious conceptual problem is that the same value of IC_50_ could originate either from relatively high initial binding affinity (low *K*_i_) and relatively low chemical reactivity (low *k*_inact_), or the other way around. Therefore, two or more inhibitors with disparate molecular properties could easily manifest as having “the same” biochemical potency if the IC_50_ assay is conducted at a single point in time.

This report presents a data-analytic procedure that relies on two separate IC_50_ determinations, conducted at two different reaction times. If a given inhibitor follows a stepwise mechanism of inhibition, involving a kinetically detectable noncovalent intermediate, we show that it is possible to estimate both *K*_i_ and *k*_inact_ from the two time-dependent IC_50_ measurements alone. Many highly potent covalent inhibitors display a one-step kinetic mechanism, apparently without the involvement of a clearly detectable noncovalent intermediate [6]. In those cases it is in principle impossible to measure the *K*_i_ and *k*_inact_ separately, but the data-analytic method described here allows for the determination of the covalent efficiency constant *k*_eff_, also known as *k*_inact_/*K*_i_, from any single measurement of IC_50_. The proposed method is validated by using a simulation study involving 64 computer-generated inhibitors with molecular properties (*K*_i_, *k*_inact_, and *k*_inact_/*K*_i_) spanning at least six orders of magnitude.

## 2. Methods

This section describes the theoretical and mathematical methods that were used in heuristic simulations described in this report. All computations were performed by using the software package DynaFit [7, 8]. Explanation of all mathematical symbols is given in the Appendix, see *Table A.1* and *Table A.2*.

### 2.1. Kinetic mechanisms of irreversible inhibition

In this report we will consider in various contexts the kinetics mechanisms of substrate catalysis and irreversible inhibition depicted in *Figure 1*. For details see ref. [9].

**Figure 1:**
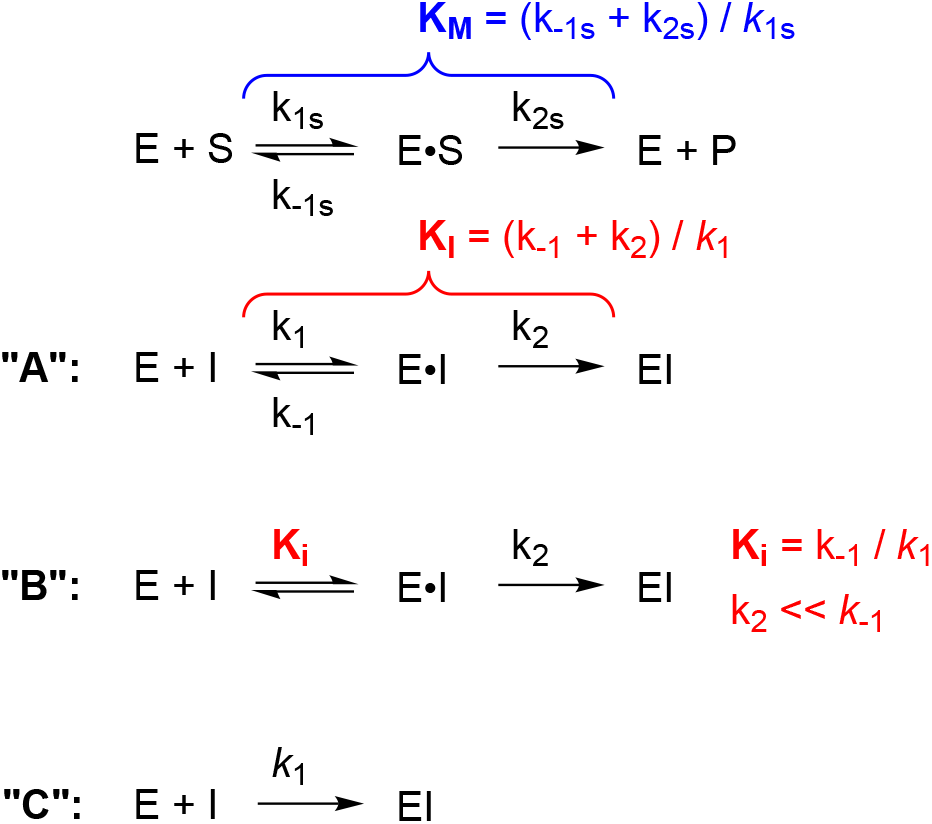
Kinetic mechanisms of substrate catalysis (top) and covalent inhibition (mechanisms **A** – **C**).

### 2.2. Mathematical models

#### 2.2.1. *Determination of* 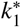 *from a single measurement of* IC_50_

On the assumption that the enzyme assay proceeds kinetically via the one-step inhibition mechanism **C**, the apparent second-order rate constant 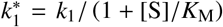 can be computed directly from a single measurement of IC_50_ by using Eqn (1), where *t*_50_ is the reaction time used to determine the experimental value of IC_50_. The requisite algebraic derivation is shown in the Appendix. Eqn (2) defines the second-order covalent efficiency constant *k*_eff_, also known as “*k*_inact_/*K*_i_” in the context of the one-step mechanism **C**.

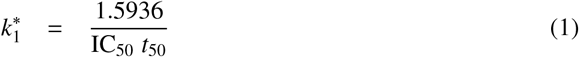

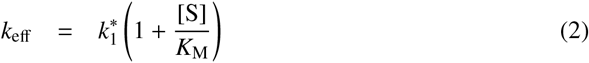

#### 2.2.2. Model discrimination analysis

On the assumption that the enzyme assay proceeds kinetically via the one-step inhibition mechanism **C**, the apparent second-order rate constant 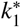 is by definition invariant with respect to time, according to Eqn (1). Thus, in the idealized case of zero experimental error, two values 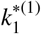 and 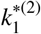 determined at two different stopping times *t*^(1)^ and *t*^(2)^ should be exactly identical. However, in the realistic case of non-zero experimental error, the two 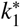 values will always be ever so slightly different even if the one-step kinetic mechanism **C** is actually operating. In order to decide whether or not 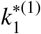 and 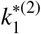 are sufficiently similar to warrant the acceptance of the one-step kinetic model, we use the geometric standard deviation defined by Eqn (4), where *μ_g_* is the geometric mean defined by Eqn (3).

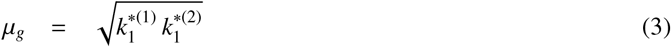

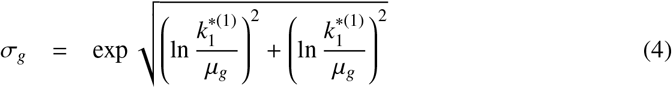

The maximum acceptable value of *σ_g_* will depend on the experimental situation. In the simulation study reported below, where the pseudo-random noise was equal to one percent of the maximum signal (e.g., fluorescence value), we found that *σ_g_* < 1.25 was producing satisfactory results. Note that the geometric standard deviation *σ_g_* is a dimensionless quantity measuring the “X-fold variation” associated with a group of numerical values. In our situation, *σ_g_* < 1.25 means that the two values 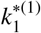 and 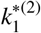 in a statistical sense differ by less than a factor of 1.25, or roughly by 25 percent in either direction (lower or higher).

#### 2.2.3. *Determination of k*_inact_ *and* 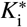 *from two values of* IC_50_

On the assumption that the enzyme assay proceeds kinetically via the two-step inhibition mechanism **B**, the apparent inhibition constant 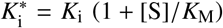 can be computed directly from any two measurements of IC_50_ by using Eqn (5), where 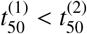 are the two reaction times used to determine 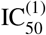 and 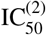, respectively, and *a* is an empirical constant (see below).

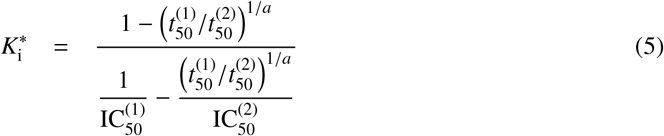

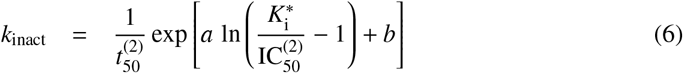

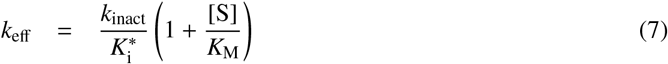

Once 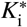 is determined from Eqn (5), *k*_inact_ can be computed from it and from 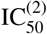 by using Eqn (6), where *a* and *b* are empirical constants. The value of *b* depends on the units used to express time and concentration. When time is expressed in seconds and concentrations in micromoles per liter, *b* = 0.558. The value of *a* = 0.9779 irrespective of units. Eqns (5)–(6) are derived in the Appendix. Eqn (7) defines the second-order covalent efficiency constant in the context of the two-step mechanism **B**.

For routine calculations with real-world experimental data, inevitably affected by finite random noise, it is convenient to utilize a simplified version of Eqns (5)–(6) shown in Eqns (8)–(9). Here utilize the fact that the empirical constant *a* = 0.9779 is very nearly equal to unity, which means that 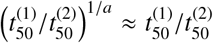. Thus, after setting *a* to unity in Eqn (5) and multiplying both the numerator and denominator by 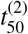, we obtain Eqn (8).

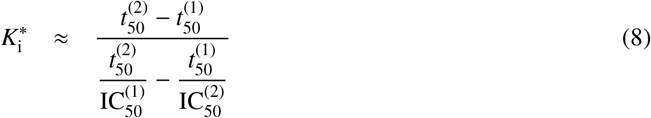

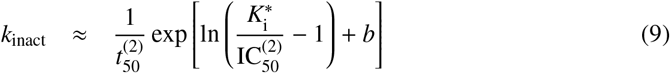

#### 2.2.4. *Implicit equation for* IC_50_ *vs. time in mechanism* **B**

For validation purposes, the dependence of IC_50_ on the reaction time was modeled by using the implicit algebraic Eqn (10), which is a minor variation of an equivalent implicit equation previously derived by Krippendorff *et al*. [10]. Given the values of *k*_inact_, 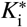 and *t*_50_, the iterative numerical solution to obtain the corresponding value of IC_50_ was accomplished by using the Newton-Raphson method [11].

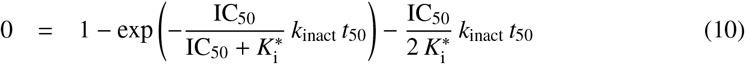

#### 2.2.5. ODE model for covalent enzyme inhibition

In the context of first-order ordinary differential-equation (ODE) modeling, the two-step inhibition mechanism **A** in *Figure 1* is mathematically represented by the ODE system defined by Eqns (11)–(17).

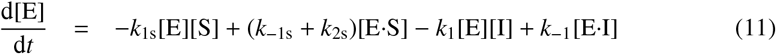

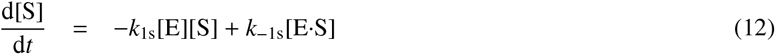

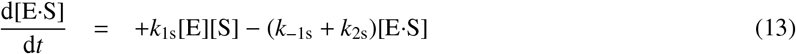

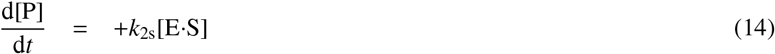

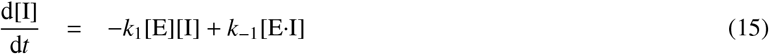

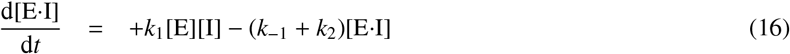

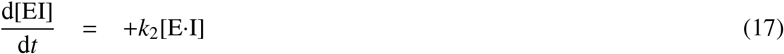

The ODE system defined by Eqns (11)–(17) was automatically generated by the software package DynaFit [8] from symbolic input. See the *Supporting Information* for details. The experimental signal was simulated according to Eqn (18), where *F* is the signal value, for example fluorescence intensity at time *t*, *F*_0_ is the baseline offset (a property of the instrument), *r*_P_ is the molar response coefficient of the product P, and (P) is the product concentration at time *t* computed by solving the initial value problem defined by Eqns (11)–(17).

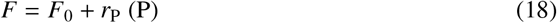

The microscopic rate constants *k*_1_, *k*_−1_ and *k*_2_ that we used in the simulation study described below can be related to the macroscopic kinetic constants as is shown in Eqns (19)–(21).

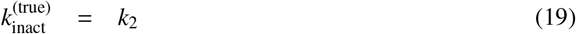

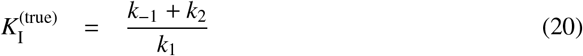

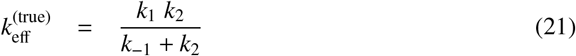

#### 2.2.6. *Determination of* IC_50_ *from simulated signal*

The IC_50_ values were determined by a fit of simulated florescence values to Eqn (22). The three adjustable model parameters were the control fluorescence intensity *F*_c_, corresponding to zero inhibitor concentration; the IC_50_; and the Hill constant *n*. It was assumed that at asymptotically infinite inhibitor concentration the fluorescence signal is by definition equal to zero.

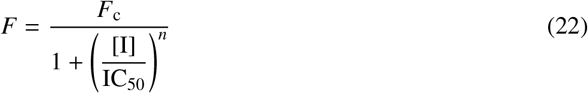

## 3. Results

This section describes the results of a simulation study that was designed to validate the determination of the kinetic constants *k*_inact_, *K*_i_, and *k*_cat_/*K*_i_ from two measurements of IC_50_. First we present an illustrative example of an irreversible inhibitor following the one-step mechanism **C**. Next we demonstrate the newly proposed method on an inhibitor following the two-step rapid-equilibrium kinetic mechanism **B**. Finally, a summary of results is given for all simulated compounds.

### 3.1. Simulation study design

The simulation study was designed such that each of the three microscopic rate constants appearing in the kinetic mechanism **A** varied by three orders of magnitude, stepping by a factor of 10. The association rate constant *k*_1_ varied from 10^4^ to 10^7^ M^−1^s^−1^; the dissociation rate constant *k*_−1_ varied from 0.001 to 1 s^−1^; and the inactivation rate constant varied from 0.0001 to 0.1 s^−1^. The corresponding covalent inhibition constant *K*_I_ ≡ (*k*_−1_+*k*_2_)/*k*_1_ [12] varied by six orders of magnitude from 110 pM to 110 *μ*M; the second-order covalent efficiency constants *k*_eff_ ≡ *k*_1_ *k*_2_/(*k*_2_ + *k*_−1_) varied by seven orders of magnitude from 0.99 M^−1^s^−1^ to 9.9 × 10^6^M^−1^s^−1^; the partition ratio *k*_2_/*k*_−1_ varied by six orders of magnitude from *k*_2_/*k*_−1_=0.0001 to *k*_2_/*k*_−1_=100. Note that the partition ratio determines the extent to which the conventionally invoked rapid equilibrium approximation (*k*_2_/*k*_−1_ << 1) is satisfied for any given compound. The corresponding “compound numbers” for the 4 × 4 × 4 = 64 simulated inhibitors are summarized in *Table 1*.

**Table 1:**
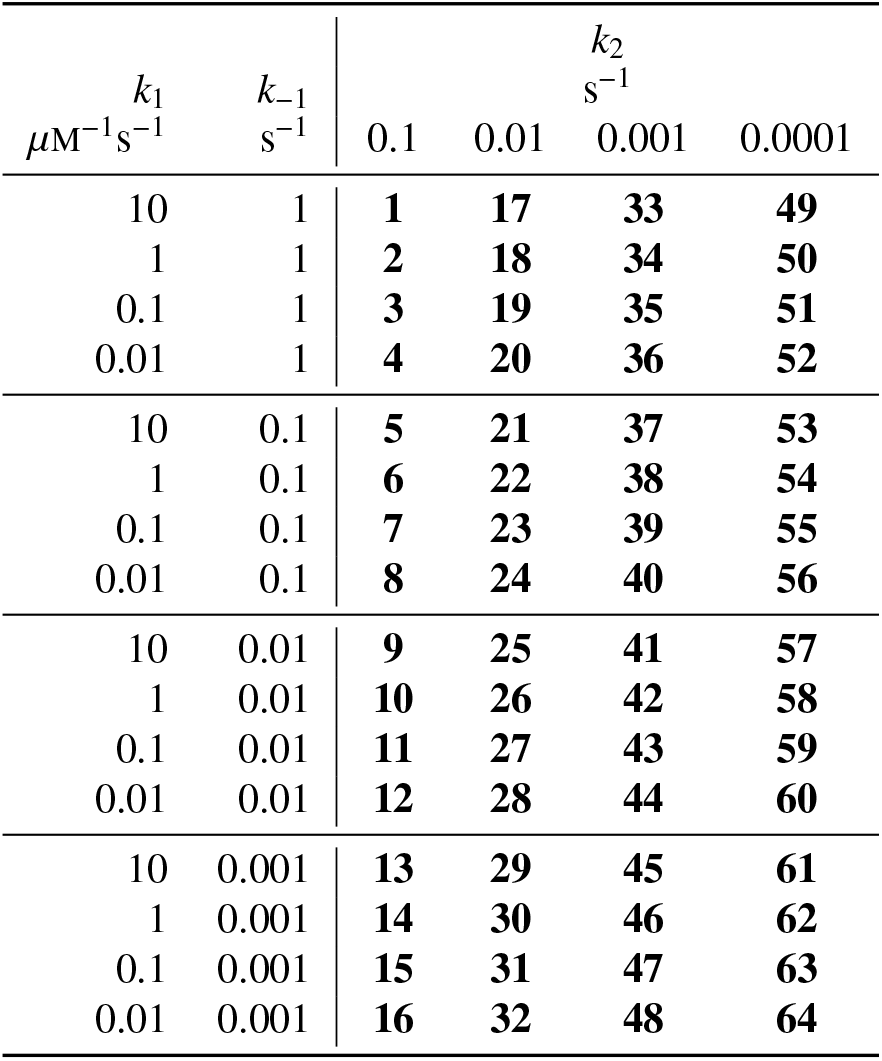
“Compound numbers” attached to each of the 64 simulated inhibitors.

| $k_1$<br>$\mu M^{-1}s^{-1}$ | $k_{-1}$<br>$s^{-1}$ | $k_2$<br>$s^{-1}$ | | | |
| --- | --- | --- | --- | --- | --- |
|  |  | 0.1 | 0.01 | 0.001 | 0.0001 |
| 10 | 1 | **1** | **17** | **33** | **49** |
| 1 | 1 | **2** | **18** | **34** | **50** |
| 0.1 | 1 | **3** | **19** | **35** | **51** |
| 0.01 | 1 | **4** | **20** | **36** | **52** |
| 10 | 0.1 | **5** | **21** | **37** | **53** |
| 1 | 0.1 | **6** | **22** | **38** | **54** |
| 0.1 | 0.1 | **7** | **23** | **39** | **55** |
| 0.01 | 0.1 | **8** | **24** | **40** | **56** |
| 10 | 0.01 | **9** | **25** | **41** | **57** |
| 1 | 0.01 | **10** | **26** | **42** | **58** |
| 0.1 | 0.01 | **11** | **27** | **43** | **59** |
| 0.01 | 0.01 | **12** | **28** | **44** | **60** |
| 10 | 0.001 | **13** | **29** | **45** | **61** |
| 1 | 0.001 | **14** | **30** | **46** | **62** |
| 0.1 | 0.001 | **15** | **31** | **47** | **63** |
| 0.01 | 0.001 | **16** | **32** | **48** | **64** |

The assumed values of substrate rate constants appearing in *Figure 1* were *k*_1s_ = 1 *μ*M^−1^s^−1^, *k*_−1s_ = 1 s^−1^, and *k*_2s_ = 1 s^−1^. Thus, the corresponding Michaelis constant had the value *K*_M_ ≡ (*k*_−1s_ + *k*_2s_)/*k*_1s_ = 2 *μ*M. The simulated substrate concentration was [S] = 2 *μ*M, such that the adjustment factor for competitive inhibition was 1 + [S]/*K*_M_ = 2. Each dose-response data set consisted of 12 progress curves corresponding to 11 nonzero inhibitor concentrations, plus the positive control progress curve at [I] = 0. The maximum inhibitor concentration was set to one fifth of the “true” covalent inhibition constant *K*_I_. For example the maximum concentration for compound **5** (*k*_1_ = 10 *μ*M^−1^s^−1^, *k*_−1_ = 0.1 s^−1^, and *k*_2_ = 0.1 s^−1^) was set to [I] = 0.2 × (0.1 +0.1)/10 = 0.004 *μ*M. The remaining 10 inhibitor concentrations represented a 1:2 dilution series; the resulting inhibitor concentration range spanned three orders of magnitude. The maximum inhibitor concentrations ranged from 22 *μ*M to 2 nM. The simulated enzyme concentration was [E] = 1 pM, which is lower by at least a factor of 2 than the minimum nonzero inhibitor concentration generated for any compound.

The simulated experimental signal was assumed to be directly proportional to the concentration of the reaction product, P. The assumed molar response coefficient of the enzymatic product was *r*_P_ = 10000 instrument units (for example, relative fluorescence units) per *μ*M. Each simulated fluorescence value was perturbed by normally distributed pseudo-random error equal to 1% of the maximum simulated signal value. Experimental signal values were simulated at five different stopping point, *t* = 15, 30, 60, 120, and 240 min. Importantly, *only two of the simulated signal values (generated t = 30 min and t = 2 hr) were used for the kinetic analysis*. The remaining time points were used merely to verify qualitative systematic trends in the simulated data, but were otherwise ignored for the purpose of determining the kinetic constants *k*_inact_, *K*_i_, and/or *k*_inact_/*K*_i_.

### 3.2. *Example 1: One-step kinetic mechanism* C

Compound **17** (*k*_1_ = 10 *μ*M^−1^s^−1^; *k*_−1_ = 1 s^−1^; *k*_2_ = 0.01 s^−1^) represents a typical example of a simulated inhibitor conforming to the one-step kinetic mechanism **C**, according to the present data-analytic procedure. The simulated reaction progress curves are shown in *Figure 2*, left panel, where the smooth theoretical model curves correspond to the numerical solution of the differential equation system Eqns (11)–(17), as described in section 2.2.5. The labels “A01” through “A11” in the left panel of *Figure 2* correspond to inhibitor concentrations ranging from 505 nM to 0.49 nM (1:2 serial dilution); progress curve “A12” represents the positive control.

**Figure 2:**
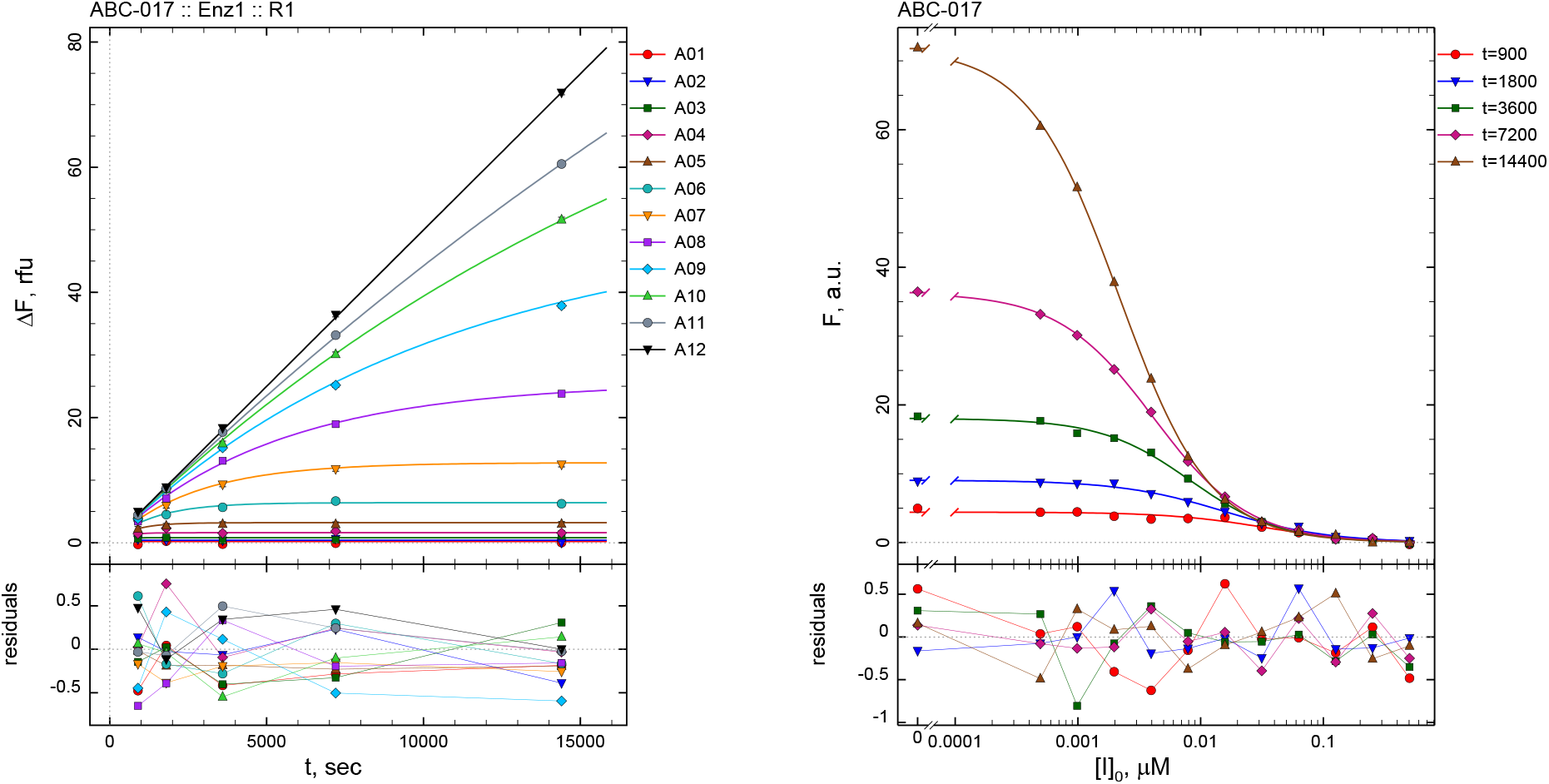
**Left:** Raw experimental signal simulated for compound **17**. **Right:** Results of fit of simulated experimental data for compound **17** to Eqn (22) to determine IC_50_ values.

**Figure 3:**
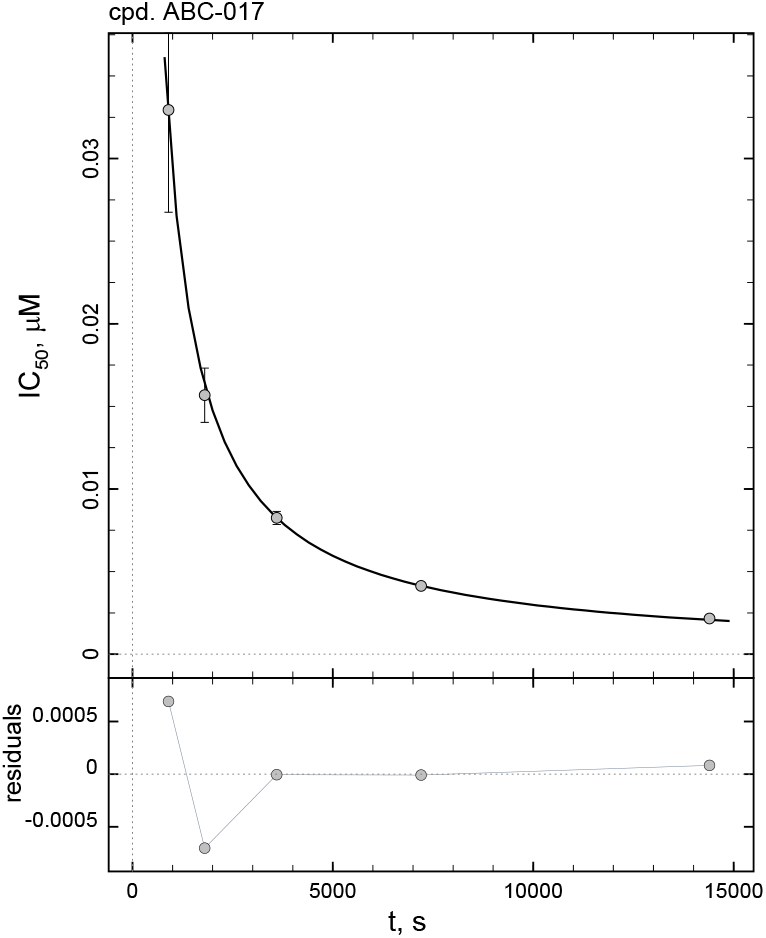
Results of fit of IC_50_ results for compound **17** (see *Figure 2*) to the implicit Eqn (10).

Each individual progress curve shown in *Figure 2* was fit separately to the three-parameter logistic Eqn (22). The best-fit values of the Hill constant *n* ranged from 1.01 to 1.17 for all 11 analyzed dose–response curves. Importantly, the best-fit values of IC_50_ were *inversely proportional to the reaction time*, which immediate alerts us to the involvement of the one-step kinetic mechanism, as predicted by Eqn (1). The details are summarized in the *Supporting Information*. Briefly, at stopping times equal to 15, 30, 60, 120 and 240 minutes (increasing systematically by a factor of 2) the best-fit values of IC_50_ were 33, 17, 8.2, 4.1, and 2.1 nM, again stepping approximately by a factor of 2, but in the opposite direction. Accordingly, as is predicted by the theoretical model specified by Eqn (1) for the one-step kinetic mechanism **C**, the calculated 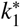 value is largely invariant with respect to time. In particular, the five calculated 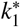 values were 53.8, 56.4, 53.7, 53.5, and 51.3 mM^−1^s^−1^, respectively (see *Supporting Information* for details).

For the purpose of the kinetic analysis, in this report we are purposely considering only two of the five stopping points, namely 30 min and 120 min. The geometric standard deviation for the two relevant values of 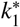 (in this case 56.4 and 53.5 mM^−1^s^−1^, respectively) was 1.04, which is lower than our acceptance criterion *σ_g_* < 1.25. Thus, we accept as the final result the apparent 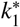 value corresponding to 120 min, in this case 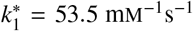. This corresponds to 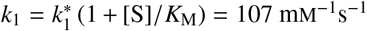. Similarly from the IC_50_ value determined at *t* = 30 min, *k*_1_ = 56.3×(1+2/2) = 113mM^−1^s^−1^. The “true” i.e. simulated value of the second-order covalent efficiency constant for compound **17** is *k*_inact_/*K*_I_ = *k*_eff_ = *k*_1_ *k*_2_/(*k*_−1_ + *k*_2_) = 10 × 0.01/(1 +0.01) = 0.099 *μ*M^−1^s^−1^ = 99 mM^−1^s^−1^. Thus, the “true” value (99 mM^−1^s^−1^) and the two calculated values each based on a single determination of IC_50_ (107 and 113 mM^−1^s^−1^, respectively) agree to within approximately ten to fifteen percent.

In conclusion, the covalent efficiency constant *k*_inact_/*K*_i_ for compound **17** could be determined from either of two IC_50_ determinations, either at 30 minutes or two hours, using the simple formula represented by Eqn (1). Importantly, the fact that the two efficiency constant values are in good agreement (*σ_g_* < 1.25) provides an internal check on the underlying kinetic mechanism, in this case the one-step kinetic mechanism **C**.

### 3.3. *Example 2: Two-step kinetic mechanism* **B**

Compound **33** (*k*_1_ = 10 *μ*M^−1^s^−1^; *k*_−1_ = 1 s^−1^; *k*_2_ = 0.001 s^−1^) represents a typical example of a simulated inhibitor conforming to the two-step kinetic mechanism **B**. Note that the only difference between compound **33** and compound **17** analyzed as Example 1 above is a ten-fold difference in the inactivation rate constant *k*_2_. In the case of compound **17**, the inactivation rate constant was ten times higher (*k*_2_ = 0.01 s^−1^) compared to compound **33** (*k*_2_ = 0.001 s^−1^). The association rate constant *k*_1_ and the dissociation rate constant *k*_−1_ have identical values for both compounds.

The simulated experimental signal (*Figure 4*, left panel) was fit to the three-parameter logistic Eqn (22). The best-fit values of the Hill constant *n* ranged from 0.97 to 1.13 for all 11 analyzed progress curves. The best-fit values of IC_50_ corresponding to each of the five different stopping times (15, 30,…, 240 min) are displayed in *Figure 4*, right panel. Full details of the kinetic analysis are shown in the *Supporting Information*. Briefly, the best-fit values of IC_50_ at stopping times 30 and 120 min were 92.7 and 36.4 nM respectively, which corresponds to 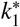 values 9.6 and 6.1 mM^−1^s^−1^ according to Eqn (1). The geometric standard deviation associated with these two numerical values is 1.38, which is higher than the cut-off acceptance criterion *σ_g_* < 1.25. Therefore compound **33** is assigned the two-step kinetic mechanism **B**. The computation of the efficiency constant then proceeds in three consecutive steps. In the first step, we compute an estimate of the apparent inhibition constant using the simplified empirical Eqn (8), as follows:

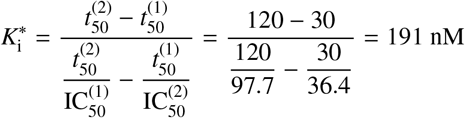

**Figure 4:**
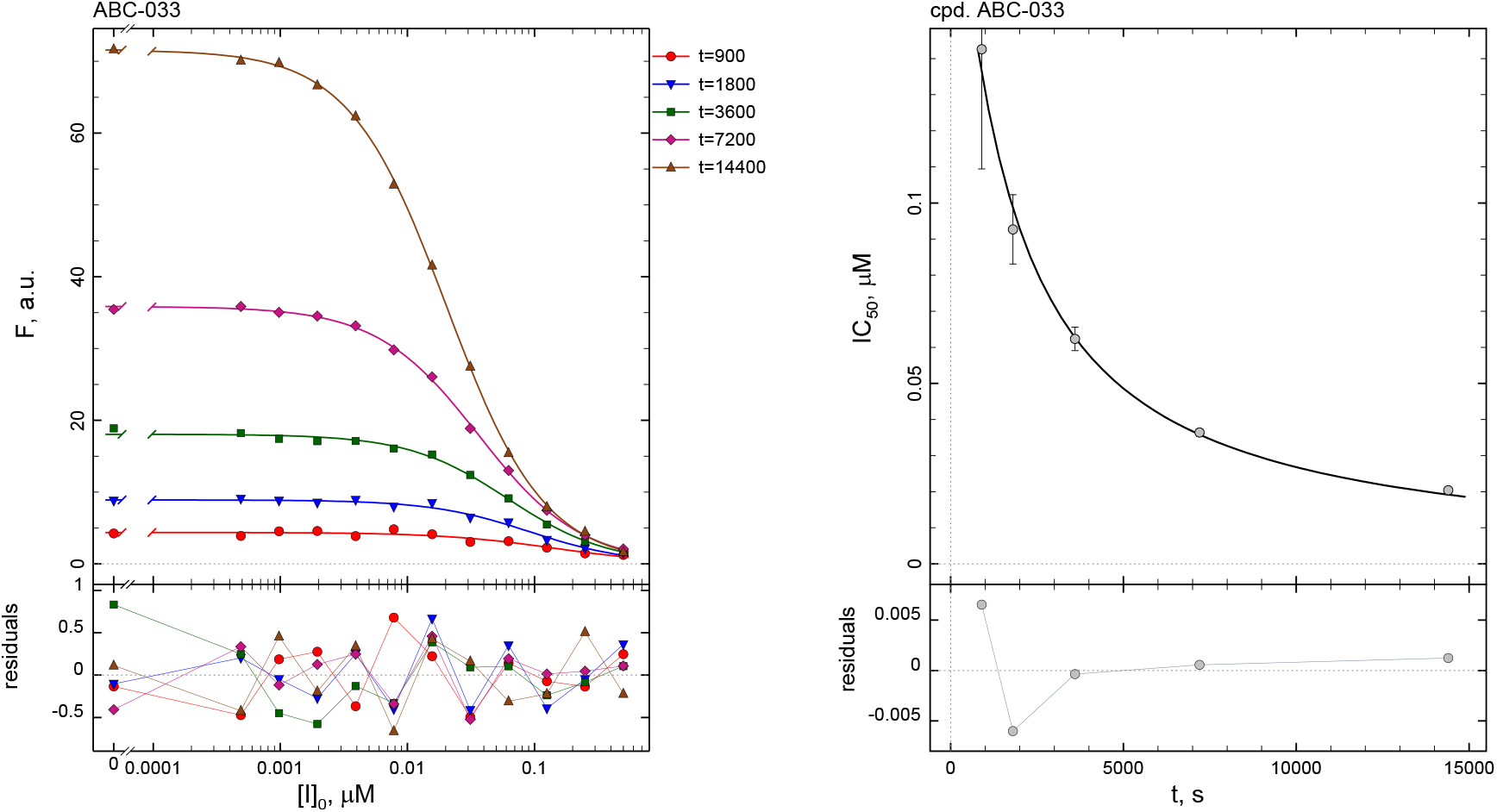
**Left:** Results of fit of simulated experimental data for compound **33** to Eqn (22) to determine IC_50_ values. **Right:** Results of fit of IC_50_ results for compound **33** to the implicit Eqn (10).

After converting to micromoles per liter, 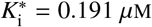. In the second step, we use the Eqn (6) to compute *k*_inact_ from 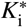 and one of the IC_50_ values obtained at the later stopping time, in this case 7200 sec (IC_50_ = 0.0364 *μ*M):^2^

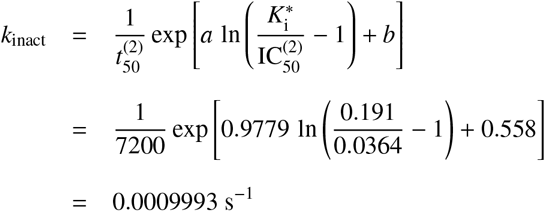

In the third and final step, we compute the covalent efficiency constant *k*_eff_ as a ratio of *k*_inact_ over *K*_i_ and simultaneously adjust both *k*_eff_ and *K*_i_ for the assumed kinetically *competitive* initial binding. Thus, 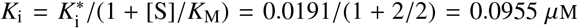 and *k*_eff_ = *k*_inact_/*K*_i_ = 0.0105 *μ*M^−1^ s^−1^. The “true” i.e. simulated value of the second-order covalent efficiency constant for compound **33** is *k*_eff_ = *k*_1_ *k*_2_/(*k*_−1_ + *k*_2_) = 10 × 0.001/(1 +0.001) = 0.0999 *μ*M^−1^s^−1^, which agrees within less than 5% with the calculated value *k*_eff_ = 0.0105 *μ*M^−1^s^−1^. The “true” i.e. simulated value of the covalent inhibition constant is *K*_I_ = (*k*_−1_ + *k*_2_)/*k*_1_ = (1+0.001)/10 = 0.1001 *μ*M, which agrees within less than 5% with the calculated value *K*_i_ = 0.0955 *μ*M.

In conclusion, the covalent efficiency constant *k*_inact_/*K*_i_ for compound **33**, as well as the covalent inhibition constant *K*_i_ and the inactivation rate constant *k*_inact_, could be determined from only two IC_50_ determinations conducted at 30 minutes and two hours, using simple algebraic formulas that can be implemented in a common spreadsheet or a calculator. The theoretically expected and calculated values for all three kinetic constants differ by less than 10%.

### 3.4. Summary of results for all 64 inhibitors

#### 3.4.1. Assignment of kinetic mechanism

Preliminary investigations revealed that the assignment of the optimal kinetic mechanism (either one-step or two-step) to each inhibitor depends on the choice of stopping times 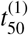 and 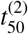, as well as on the choice of the empirical model-acceptance criterion *σ_g_*. Using 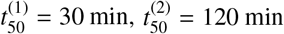, and *σ_g_* < 1.25, the results are summarized in *Table 2*.

**Table 2:** Kinetic mechanisms assigned to the simulated compounds using 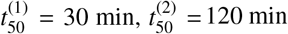, and *σ_g_* < 1.25.

| $k_1$<br>$\mu\text{M}^{-1}\text{s}^{-1}$ | $k_{-1}$<br>$\text{s}^{-1}$ | $k_2$<br>$\text{s}^{-1}$ | | | |
| --- | --- | --- | --- | --- | --- |
|  |  | 0.1 | 0.01 | 0.001 | 0.0001 |
| 10 | 1 | **C** | **C** | **B** | **B** |
| 1 | 1 | **C** | **C** | **B** | **B** |
| 0.1 | 1 | **C** | **C** | **B** | **B** |
| 0.01 | 1 | **C** | **C** | **B** | **B** |
| 10 | 0.1 | **C** | **C** | **C** | **B** |
| 1 | 0.1 | **C** | **C** | **B** | **B** |
| 0.1 | 0.1 | **C** | **C** | **B** | **B** |
| 0.01 | 0.1 | **C** | **C** | **B** | **B** |
| 10 | 0.01 | **C** | **C** | **B** | **B** |
| 1 | 0.01 | **C** | **C** | **B** | **B** |
| 0.1 | 0.01 | **C** | **C** | **B** | **B** |
| 0.01 | 0.01 | **C** | **C** | **B** | **B** |
| 10 | 0.001 | **C** | **C** | **C** | **B** |
| 1 | 0.001 | **C** | **C** | **C** | **B** |
| 0.1 | 0.001 | **C** | **C** | **C** | **B** |
| 0.01 | 0.001 | **C** | **C** | **C** | **B** |

The results shown in *Table 2* can be summarized as follows. (i) Compounds **1** – **32**, characterized by a relatively *fast* chemical step with *k*_2_ ≡ *k*_inact_ ≥ 0.01 s^−1^, were judged by the model selection algorithm to be following the *one-step* kinetic mechanism **C**, irrespective of the particular value of the partition ratio *k*_2_/*k*_−1_. (ii) Compounds **49** – **64**, characterized by a relatively *slow* chemical step with *k*_2_ ≡ *k*_inact_ ≤ 0.0001 s^−1^, were judged by the model selection algorithm to be following the *two-step* kinetic mechanism **B**, again irrespective of the particular value of the partition ratio *k*_2_/*k*_−1_. (iii) Compounds **33** – **48** associated with an intermediate value of *k*_2_ = 0.001 s^−1^ followed either of the two kinetic mechanisms, depending on the partition ratio *k*_2_/*k*_−1_. In particular, compounds **33** – **44** with the exception of **37** displayed the two-step kinetic mechanism **B**. In all those cases, the chemical step is slower than the dissociation step (*k*_2_/*k*_−1_ < 1). In contrast, compounds **45** – **48**, for which *k*_2_/*k*_−1_ = 1, conformed to the one-step kinetic mechanism **C**.

#### 3.4.2. Calculated values of macroscopic kinetic constants

The calculated values of the second-order covalent efficiency constant *k*_eff_ for all 64 simulated inhibitors, as determined by the two-point method, are summarized in *Figure 5*, left panel. See *Supporting Information* for details. Note that the semantics of *k*_eff_ differs depending on the kinetic mechanism of inhibition (either **B** or **C**) assigned to each individual compound, as shown in *Table 2*. Thus, *k*_eff_ = *k*_1_ for compounds that follow the one-step kinetic mechanism **C**, whereas *k*_eff_ = *k*_inact_/*K*_i_ for compounds that follow the two-step kinetic mechanism **B**.

**Figure 5:**
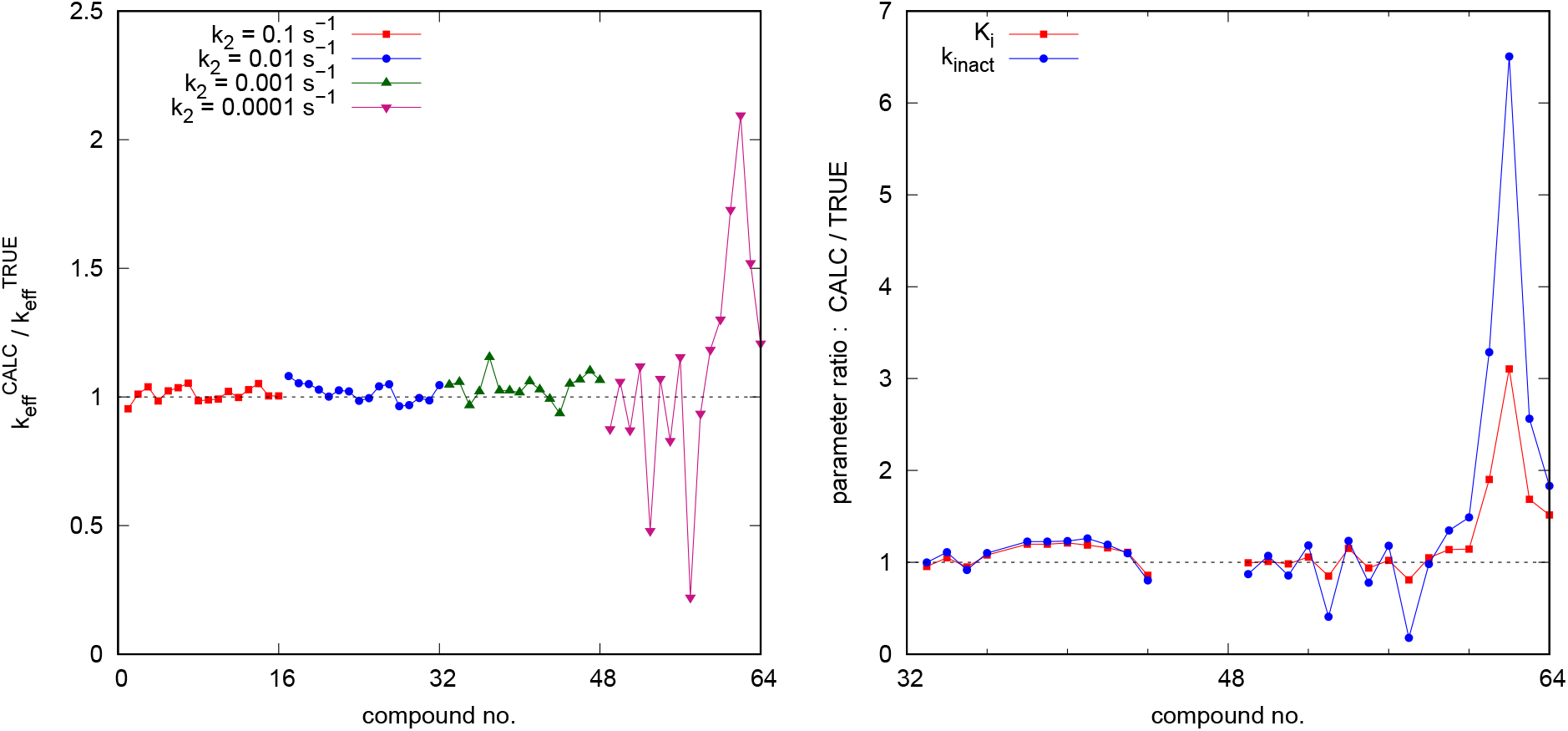
The ratios of calculated vs. “true” (i.e., simulated) values of kinetic constants. **Left:** *k*_eff_ a.k.a. *k*_inact_/*K*_i_, compounds **1**–**64**. **Right:** *k*_inact_ and *K*_i_, compounds **33**–**64** only.

The results displayed in the left-hand panel of *Figure 5* indicate that *k*_eff_ is determined with better than approximately 25% accuracy for all inhibitors except compounds **49**– **64**, which are associated with very slow chemical inactivation step (*k*_inact_ = *k*_2_ = 0.0001 s^−1^). In the latter group of compounds, the ratio of the calculated over “true” i.e. simulated *k*_eff_ varies approximately from 0.2 to 2.0. Note that for all inhibitors following the two-step kinetic mechanism, the efficiency constant *k*_eff_ is computed after the fact, as a *ratio* of independently determined *k*_inact_ and *K*_i_ values. Thus, the question remains which of these two contributing factor, if any, is principally responsible for the lack of accuracy.

An explanation for the relatively large uncertainty of *k*_eff_ seen in most “slow” inactivators is presented in *Figure 5*, which shows that in most cases (in particular, compounds **49** – **60**) the inhibition constant *K*_i_ is determined quite accurately, whereas the inactivation constant *k*_inact_ shows a large degree of uncertainty. Only compounds **61** – **64** show a relatively large discrepancy between the “true” and calculated values for both *k*_inact_ and *K*_i_. Note that compounds **61** – **64** are genuinely exceptional in two different respects. Not only their chemical reactivity is exceptionally low, as measured by *k*_inact_ = *k*_2_ = 0.0001 s^−1^, but also their dissociation rate constant *k*_−1_ = 0.0001 s^−1^ is the lowest in the entire compound collection. An examination of instantaneous rate plots for these four compounds (see *Supporting Information*) shows that the there is a kinetic transient (a “slow binding” phenomenon [13, 14]) that dramatically distorts the IC_50_ values determined at *t* = 30 min.

## 4. Discussion

### Assumptions and limitations of the present method

The theoretical model represented by Eqns (1)–(7) is based on two simplifying assumptions. Note that the two assumptions are those that also underlie the standard “*k*_obs_” method [5] and the Krippendorff method [10] of analyzing the time-dependence of IC_50_ from multiple measurements. First, it is assumed that there is no inhibitor depletion, in the sense that during the assay only a negligibly small mole fraction of the inhibitor is bound to the enzyme, either covalently or non-covalently. This in turn implies that the total or analytic concentration of the inhibitor is always significantly higher than the initial concentration of the enzyme. In practical terms, we found that an approximately three fold excess of the lowest inhibitor concentration in a dilution series over the enzyme concentration is satisfactory. A situation that should be strenuously avoided is allowing any of the inhibitor concentrations become lower than the enzyme concentration, if and when those inhibitor concentrations are associated with any observable inhibitory effect.

Of course, depending on the nature of the assay, it may not be practically possible to lower the enzyme concentration sufficiently and still maintain assay sensitivity. For example, it may not be practically possible to use enzyme concentrations as low as [E] = 1 pM, as was done in the simulation study presented here. In fact, in many assays it becomes necessary to use enzyme concentrations as high as [E] ≈ 10 nM or even higher, because of sensitivity concerns. However, note that the binding affinity of many therapeutically important enzyme inhibitors also lies in the nanomolar region, which means that these molecules express their inhibitory potency already at [I] ≈ 1 nM or lower. Neither the “*k*_obs_” method [5], the Krippendorff method [10], nor the method presented here, can be used under such experimental circumstances, where the zero-inhibitor depletion assumption is violated. The only remedy is to deploy a data-analytic procedure that does not rely on any simplifying assumptions, meaning a mathematical model based on the numerical solution of differential equation. For an illustrative example involving the inhibition of drug-resistant EGFR mutants, see ref. [15].

The second simplifying assumption underlying the data-ana-lytic procedure presented here is that the reaction rate in the positive control experiment ([I] = 0) remains strictly constant over the entire duration of the assay. In other words, it is assumed that the positive control progress curve (time vs. experimental signal) is strictly linear. This can only be achieved if the mole fraction of the substrate ultimately consumed in the control assay remains negligibly low; if the initial concentration of the substrate is very much higher than the corresponding Michaelis constant ([S] >> *K*_M_); or if both of the above conditions are satisfied simultaneously. It should be noted that simple visual inspection can often be extremely misleading when it comes to judging linearity vs. nonlinearity of positive control assays. Instead of relying on a subjective assessment, it is preferable to deploy an objective *cross-validation procedure* described in ref. [16].

### Experimental design

Krippendorff’s [10] implicit algebraic Eqn (10) for time-dependence of IC_50_, as well as the data-analytic formulas derived in this report, are both based on the important assumption that there is *no preincubation of the enzyme with inhibitor prior to adding the substrate* to trigger the enzymatic assay. Instead, the enzyme’s interactions with the substrate and with the inhibitor need to be initiated at the same time, by adding the enzyme catalyst as the last component into the assay.

At least some practitioners in enzyme kinetics apparently misunderstand this very important aspect of covalent IC_50_ assays analyzed specifically by Krippendorff’s method [10] (and also by the two-point method presented here). For example, Fassunke *et al*. [17] reported that “for kinetic characterization (*k*_inact_/*K*_i_), the inhibitors were incubated with EGFR-mutants over different periods of time (2–90 min), whereas durations of enzymatic reactions [25 min after adding the substrate at the end of enzyme–inhibitor preincubation, note added by P.K.] were kept constant. […] Calculated IC_50_-values were […] fit as described in the literature to determine *k*_inact_ and *K*_i_”. The “literature” method mentioned immediately above is a reference to the Krippendorff method [10]. However, to repeat for emphasis, Krippendorff’s equation Eqn (10) was derived under the assumption that the onset of product formation occurs simultaneously with the onset of enzyme inhibition, otherwise Eqn (10) cannot be used. On that basis, the *k*_inact_ and *K*_i_ values reported for EGFR inhibitors in ref. [17] are almost certainly invalid.

For the purposes of utilizing the newly proposed two-point IC_50_ method, it is important to arrange the experiment such that the two stopping times are spaced sufficiently widely. Based on preliminary investigations, it appears sufficient to maintain at least a four-fold difference between 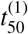 and 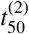, for example 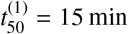 and 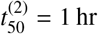, or alternately 30 min and 2 hr. The objective is to assure that the two IC_50_ values are sufficiently different from each other, such that it becomes possible to discern whether or not the two resulting IC_50_ values are inversely proportional to the stopping time according to Eqn (1). In this respect, it is advantageous to choose 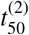 as high as is practically possible, while also keeping in mind that substrate depletion, enzyme deactivation, and other “nuisance” factors might cause the positive control progress curve to become nonlinear, which should be avoided as much as possible.

The optimal duration of the covalent inhibition assay is also closely related to the expected inactivation rate constant *k*_inact_. In the hypothetical scenario where the enzyme is instantaneously saturated with the inhibitor because the inhibitor concentration is very much higher than the covalent inhibition constant *K*_i_, the covalent conjugate EI is formed with the first-order rate constant *k*_inact_. Under these hypothetical circumstances, the half-time for inactivation is equal to ln(2)/*k*_inact_. For example, in the specific case of *k*_inact_ = 0.0001 s^−1^, the expected half-time for inactivation is *t*_1/2_ = ln(2)/0.0001 = 0.693/0.0001 = 6930 s = 115 min, or approximately two hours. Assuming that nearly full inactivation is achieved at about *t*_max_ ≈ 3 × *t*_1/2_, the assay would have to last almost six hours in order to see the enzyme fully inactivated. An enzyme assay that long of course might not be possible for numerous practical reasons, which also means that covalent inhibitors with *k*_inact_ ≤ 0.0001 s^−1^ are exceedingly difficult to characterize accurately; see also the results reported here for simulated compounds **49** – **64**.

### Choice of the model selection criterion σ_g_

A successful application of the two-point data analytic procedure described in this report depends on a suitable choice of the model selection criterion *σ_g_*. Recall that *σ_g_* is the geometric standard deviation between two numerical values of IC_50_, defined by Eqn (4), and is used to decide between the one-step kinetic mechanism **C** and the two-step kinetic mechanism **B**. The optimal choice *σ_g_* depends on the nature of assay and also on the choice of the two stopping times, 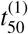 and 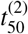. We found that for more closely spaced stopping time values, *σ_g_* ≈ 1.25 performs satisfactorily, whereas for stopping times separated by a factor of five or higher, *σ_g_* ≈ 1.5 works better. The optimal value of *σ_g_* may need to be adjusted in the course of an inhibitor screening campaign, as practical experiences are being accumulated.

### Similarities and differences with the Krippendorff method

The present method is similar to the method of Krippendorff *et al*. [10] in that both methods use the same theoretical foundation represented by Eqn (10). There are also three major differences. First, our method requires only two measurements of IC_50_ whereas the Krippendorff method requires several times as many data points. Second, our method can be computationally implemented very simply by using a common spreadsheet program, whereas the Krippendorff method requires a highly specialized software package that allow nonlinear least-squares fit to an *implicitly* defined algebraic model, of the general form *f*(*X*, *Y*) = 0, as opposed to the much more common *explicit* algebraic equation, of the general form *Y* = *f*(*X*). Third, and most important, the Krippendorff method is based on an *assumption* that all inhibitors follow the two-step kinetic mechanism **B**, whereas our method allows the actual kinetic mechanism to be detected from the experimental data, without making prior assumptions.

In fact, we have previously documented [6, 9] that covalent inhibitors characterized by high initial binding affinity, high chemical reactivity, or both, will outwardly display one-step kinetics. In the specific case of highly “tight binding” inhibitors [14, 18], which are characterized by relatively low dissociation rate constant *k*_−1_ in the initial noncovalent step, the noncovalent complex dissociates only very slowly on the time-scale of the experiment, which renders even the first (strictly speaking, noncovalent) binding step effectively irreversible. In the case of highly reactive covalent inhibitors, which are characterized by relatively high inactivation rate constant *k*_2_, the initial noncovalent complex may be pulled forward through the reaction pathway so rapidly that the mole fraction of E·I remains kinetically undetectable. That is why covalent inhibitors characterized by relatively high inactivation constant (*k*_2_ ≥ 0.01 s^−1^) are not expected to yield a meaningful value of the dissociation constant *K*_i_, even though the initial noncovalent complex must be physically present – however fleetingly.

### *Steady-state K*_I_ *vs. rapid-equilibrium K*_i_

Throughout this report, depending on context, we have been purposely alternating between two fundamentally distinct conceptions of the inhibition constant. The classical definition of the covalent inhibition constant, found in nearly all medicinal chemistry and biochemistry literature on irreversible enzyme inhibition, is implicitly based on the *rapid-equilibrium* approximation. Accordingly, the rapid-equilibrium inhibition constant, denoted as *K*_i_ in this report, is treated as a true dissociation constant of the initial noncovalent complex *E*·*I* and is thus defined as a simple ratio of two microscopic rate constants, *k*_i_ = *k*_−1_/*k*_1_. In the history of enzyme kinetics, this definition of the inhibition constant is equivalent to the original conception of the Michaelis constant as a simple dissociation equilibrium constant [19, 20]. The rapid-equilibrium approximation was first invoked in the context of irreversible enzyme inhibition by Kitz & Wilson [21]. This is the definition of the inhibition constant invoked in this report when discussing the value of kinetic parameters determined from simulated pseudo-experimental data.

An alternate understanding of the inhibition constant in the context of irreversible enzyme inhibition, first introduced by Malcolm & Radda [12], is based on the *steady-state* approximation in enzyme kinetics. Accordingly, the steady-state inhibition constant, denoted as *K*_I_ in this report, is defined in terms of all three microscopic rate constants appearing in mechanism **A**, *K*_I_ = (*k*_−1_ + *k*_2_)/*k*_1_. Historically, this definition of the inhibition constant is equivalent to a more refined understanding of the Michaelis constant according to Briggs & Haldane [22]. This is the definition of the inhibition constant invoked in this report when discussing the “true” or simulated values of kinetic parameters.

Under most experimental situations arising in the evaluation covalent inhibitory potency, the distinction between *K*_i_ and *K*_I_ is blurred, in the sense that the individual microscopic rate constants *k*_1_, *k*_−1_ and *k*_2_ remain inaccessible to routine enzyme kinetic measurements. In fact all three microscopic constants are essentially accessible only through meticulous rapid-kinetic (e.g. stopped-flow) experimental setup. See for example a recent report on the kinetics of Bruton tyrosine kinase inhibition by the irreversible inhibitor osimertinib [23]. However, the *conceptual* distinction between *K*_i_ and *K*_I_ should always be kept firmly in mind, because it can help explain potentially puzzling experimental observations.

For example, the rapid-equilibrium dissociation constant *K*_i_ for an irreversible inhibitor characterized by *k*_1_ = 10^6^ M^−1^s^−1^ and *k*_−1_ = 0.0001 s^−1^ is *K*_i_ = *k*_−1_/*k*_1_ = 0.0001/1.0 = 0.0001 *μ*M = 0.1 nm. In contrast, the steady-state inhibition constant for the same inhibitor is a thousand times higher, *K*_I_ = (*k*_−1_ + *k*_2_)/*k*_1_ = (0.0001 + 0.1)/1 = 0.1001 *μ*M = 100.1 nm. This massive difference between *K*_i_ and *K*_I_ for the same covalent compound could potentially explain major expected differences in the dose-response (“saturation”) behavior of (a) the covalent inhibitor and (b) a corresponding non-covalent analogue, even under the assumption that both inhibitors possess approximately identical noncovalent binding affinity. In particular, the *K*_i_ for the noncovalent compound will be easily measurable at or below [I]^(max)^ ≈ 10 × *K*_i_ = 1 nM. In contrast, the covalent analogue even at a ten-fold higher inhibitor concentration, [I] = 10 nM, will be nowhere near the saturation point because 10 nm is only 10% of the *K*_I_ value. Thus, according to the rule of thumb formulated by Kitz & Wilson [21], at [I]^(max)^ = 10 nm (a value 100 times higher than the equilibrium dissociation constant) the covalent analogue will appear kinetically as a “one-step” irreversible inhibitor with immeasurably weak initial binding affinity.

### *Significance and utility of the two-point* IC_50_ *method*

The cost of successfully developing new medications and bringing them to market past unavoidable regulatory hurdles is enormous, amounting to approximately 2.6 billion US dollars per compound in 2016 [24]. Even assuming that the largest fraction of the overall expenditure is taken up by clinical trials, the cost of pre-clinical discovery processes such as the evaluation of enzyme inhibitors for biochemical potency is very significant, both in terms of human energy and in terms of material supplies. In this context, irreversible enzyme inhibitors present an exceptional challenge, because even the “simple” task of meaningfully *ranking* a series of drug candidates by biochemical potency is complicated by the fact that the overall potency of covalent inhibitors consists of two entirely separate components, namely, their initial binding affinity (*K*_i_) and their chemical reactivity (*k*_inact_). This conceptual difficulty often leads to low-information experiments that are potentially wasteful.

For example, a number of drug discovery projects begin by testing each irreversible inhibitor in a “one-hour IC_50_” assay, or in a similar single time-point IC_50_ assay. However, any value of covalent IC_50_ observed at a single time-point is by definition non-unique, because it could be produced either by an inhibitor that has high affinity (low *K*_i_) and low reactivity (low *k*_inact_) or alternately by another inhibitor that has low affinity (high *K*_i_) and high reactivity (high *k*_inact_). Because of this inherent redundancy and ambiguity, a covalent IC_50_ value determined at a single time-point cannot be used to rank irreversible inhibitors by potency in a meaningful way. In contrast, the two-point IC_50_ method presented here is guaranteed to produce a *unique* value of the covalent efficiency constant *k*_eff_ for all inhibitors, irrespective of the kinetic mechanism, and additionally also a *unique* value of *k*_inact_ and *K*_i_ for those inhibitors that formally follow the two-step kinetic mechanism **B**.

## Supporting information

Simulated data

## Acknowledgements

I thank Dr. Claire McWhirter (Artios Pharma, Cambridge, UK) for helpful discussions. Dr. Ben Allsop (GSK Medicines Research Centre, Stevenage, UK) is gratefully acknowledged for a correction to the initial release of this paper.

## Supporting information

1. File BioKinPub-2020-04-SI.pdf: Algebraic derivations; detailed kinetic analysis of all simulated inhibitors.
2. File BioKinPub-2020-04.zip: Simulated pseudo-experimental data files.

## Appendix

## A. Explanation of symbols

**Table A.1:** Explanation of symbols: Microscopic rate constants and derived kinetic constants.

| Symbol | Unit | Explanation |
| --- | --- | --- |
| $k_{1s}$ | $M^{-1}s^{-1}$ | association rate constant for $E+S \rightarrow E \cdot S$ |
| $k_{-1s}$ | $s^{-1}$ | dissociation rate constant for $E \cdot S \rightarrow E + S$ |
| $k_{2s}$ | $s^{-1}$ | turnover number; $k_{2s} \equiv k_{cat}$ |
| $K_M$ | $M$ | Michaelis constant; $K_M = (k_{-1s} + k_{2s})/k_1$ |
| $k_S$ | $M^{-1}s^{-1}$ | catalytic efficiency constant; specificity constant; $k_S \equiv k_{cat}/K_M$ |
| $k_1$ | $M^{-1}s^{-1}$ | association rate constant for $E+I \rightarrow \dots$ |
| $k_{-1}$ | $s^{-1}$ | dissociation rate constant for $E \cdot I \rightarrow E + I$ |
| $k_2$ | $s^{-1}$ | inactivation rate constant; $k_2 \equiv k_{inact}$ |
| $k_1^*$ | $M^{-1}s^{-1}$ | apparent association rate constant:<br>competitive: $k_1^* = k_1/(1 + [S]/K_M)$<br>uncompetitive: $k_1^* = k_1 (1 + [S]/K_M)$<br>noncompetitive $k_1^* = k_1$ |
| $k_{eff}$ | $M^{-1}s^{-1}$ | second-order inhibition efficiency constant; $k_{eff} \equiv "k_{inact}/K_i"$:<br>steady-state two-step mechanism **A**: $k_{eff} = k_1 k_2/(k_{-1} + k_2)$<br>rapid-equilibrium two-step mechanism **B**: $k_{eff} = k_1 k_2/k_{-1}$<br>one-step mechanism **C**: $k_{eff} = k_1$ |
| $k_{eff}^*$ | $M^{-1}s^{-1}$ | apparent inhibition efficiency constant:<br>competitive: $k_{eff}^* = k_{eff}/(1 + [S]/K_M)$<br>uncompetitive: $k_{eff}^* = k_{eff} (1 + [S]/K_M)$<br>noncompetitive $k_{eff}^* = k_{eff}$ |
| $K_i$ | $M$ | equilibrium dissociation constant of the $E \cdot I$ complex; $K_i = k_{-1}/k_1$ |
| $K_i^*$ | $M$ | apparent equilibrium dissociation constant:<br>competitive: $K_i^* = K_i (1 + [S]/K_M)$<br>uncompetitive: $K_i^* = K_i/(1 + [S]/K_M)$<br>noncompetitive $K_i^* = K_i$ |
| $K_I$ | $M$ | inhibition constant; $K_I = (k_{-1} + k_2)/k_1$ |
| $K_I^*$ | $M$ | apparent inhibition constant:<br>competitive: $K_I^* = K_I (1 + [S]/K_M)$<br>uncompetitive: $K_I^* = K_I/(1 + [S]/K_M)$<br>noncompetitive $K_I^* = K_I$ |
| $k_{obs}$ | $s^{-1}$ | apparent first-order rate constant for enzyme inactivation |

## B. Algebraic derivations

### B.1. Derivation of the “one-step” algebraic equation

Under the simplifying assumption that the uninhibited positive-control rate is constant over time and in the absence of inhibitor depletion, Kitz & Wilson [21] derived Eqn (B.2) as the mathematical model for the progress of a covalent inactivation reaction. In Eqn (B.1), (P)_c_ is the product concentration formed at time *t* in the positive control assay proceeding with the constant rate *v*_0_. In Eqn (B.2), (P)_i_ is the product concentration formed at time *t* in the inhibited assay; *v*_i_ is the initial reaction rate; and *k*_obs_ is the apparent first-order rate constant.

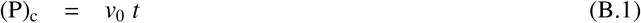

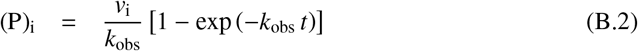

**Table A.2:** Explanation of symbols: Concentrations, reaction rates, and auxiliary symbols.

| Symbol | Unit | Explanation |
| --- | --- | --- |
| (X) | M | concentration of reactant X, where X = S, P, I, or E |
| [X] | M | initial (total, analytic) concentration of reactant X |
| $F$ | AIU | observed experimental signal in arbitrary instrument units (AIU) |
| $F_0$ | AIU | baseline signal; baseline offset |
| $r_P$ | AIU/M | molar response coefficient of the reaction product P |
| $v_0$ | $\text{Ms}^{-1}$ | initial rate of the uninhibited enzyme reaction, at $[I] = 0$ |
| $V_0$ | $\text{AIU s}^{-1}$ | observed uninhibited initial rate in arbitrary instrument units |
| $v_i$ | $\text{Ms}^{-1}$ | initial rate of the inhibited enzyme reaction, at $[I] > 0$ |
| $V_i$ | $\text{AIU s}^{-1}$ | observed inhibited initial rate in arbitrary instrument units |
| $\alpha, \beta, \gamma$ | $\text{s}^{-1}$ | auxiliary variables (groupings of rate constants) |
| $r_1, r_2$ | $\text{s}^{-1}$ | apparent bi-exponential rate constants |
| $a_1, a_2$ | — | bi-exponential amplitudes |

Kitz & Wilson [21] demonstrated that if the irreversible inhibition assay formally follows the one-step kinetic mechanism *E*+*I* → *EI*, perhaps because the inhibitor concentration is very much lower than the apparent inhibition constant 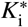, the initial reaction rate *v*_i_ is invariant with respect to the inhibitor concentration, according to Eqn (B.3), and the apparent first order rate constant *k*_obs_ is a linear function of [I], according to Eqn (B.4). Therefore the product concentration changes over time according to Eqn (B.5).

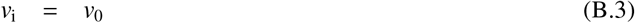

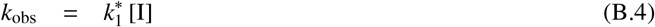

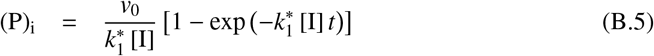

Following the line of reasoning first introduced by Krippendorff *at al*. [10], we can focus on a particular reaction time (*t*_50_) in the inhibited assay conducted at a certain nonzero inhibitor concentration (IC_50_) when the concentration of product P become exactly identical to one half of product concentration formed in the uninhibited assay. This condition is formally expressed in Eqn (B.6).

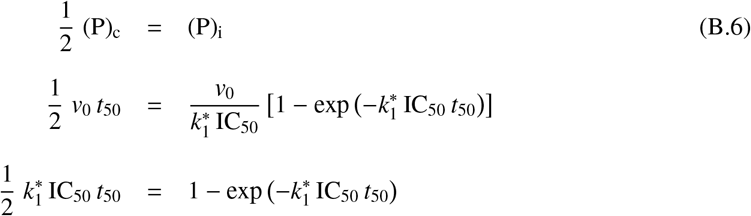

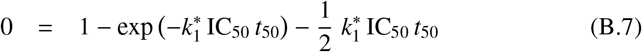

Substituting for (P)_i_ in Eqn (B.6) from Eqn (B.5), substituting for (P)_c_ from Eqn (B.1), and rearranging the resulting expression, we obtain the *implicit* algebraic Eqn (B.7). Note that the parameters 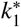, IC_50_, and *t*_50_ always appear as a product 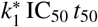. This means that infinitely many combinations of 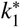, IC_50_, and *t*_50_ will satisfy the implicit Eqn (B.7) as long as the product 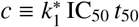 has a certain unique numerical value. To find the value of *c* that satisfies Eqn (B.9), we can conveniently use the *fixed-point iteration* [25] formula defined by Eqn (B.10), where (m+1) and (m) represent the current and the immediately preceding iteration. Starting from the initial estimate *c* = 1, the fixed-point iteration formula converged to within 14 significant digits precision at m = 25, yielding *c* = 1.5936 as the solution.

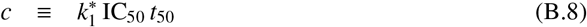

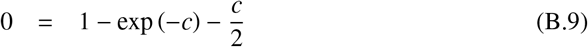

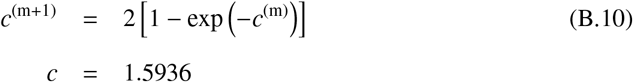

In conclusion, assuming that the one-step kinetic mechanism is operating, the apparent second-order covalent efficiency constant 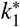 (also known as “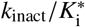”) can be determined from any single measurement of IC_50_ conducted at the reaction time time *t*_50_, according to Eqn (B.11) where *c* = 1.5936. Equivalently, given any particular value of 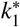, the IC_50_ at reaction time *t*_50_ can be predicted by using Eqn (B.12).

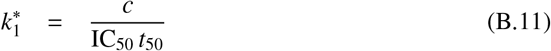

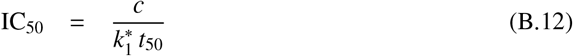

### B.2. Derivation of the “two-step” algebraic equation

Kitz & Wilson [21] demonstrated that if the irreversible inhibition assay formally follows the two-step kinetic mechanism *E* + *I* ⇌ *E*·*I* → *EI*, the initial reaction rate *v*_i_ depends on the inhibitor concentration [I] according to Eqn (B.13), and the apparent first order rate constant *k*_obs_ depends on the inhibitor concentration [I] according to Eqn (B.14). Therefore the product concentration changes over time according to Eqn (B.15).

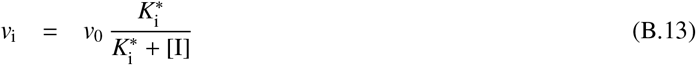

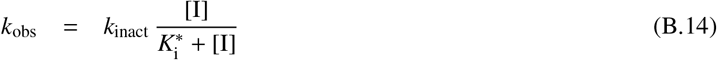

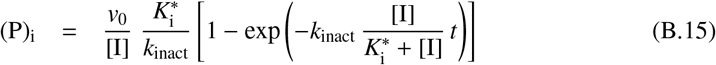

Again, following the line of reasoning first introduced by Krippendorff *at al*. [10], at a particular reaction time (*t*_50_) in the inhibited assay conducted the concentration of product P become exactly identical to one half of product concentration formed in the uninhibited assay, as shown in Eqn (B.6). Substituting for (P)_i_ in Eqn (B.6) from Eqn (B.15), substituting for (P)_c_ from Eqn (B.1), and rearranging the resulting expression, we obtain the implicit algebraic Eqn (B.16).

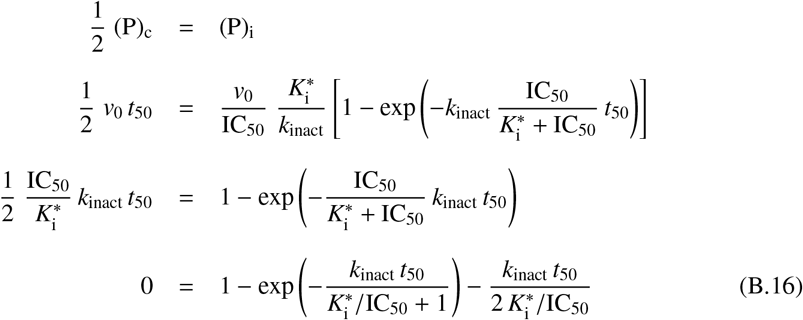

Eqn (B.16) contains four variables: *k*_inact_, 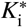, IC_50_ and *t*_50_. Howewer, note that *k*_inact_ *t*_50_ always appear as a product whereas 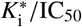 always appear as a ratio. This means that infinitely many combinations of *k*_inact_, 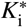, IC_50_ and *t*_50_ will satisfy Eqn (B.16) as long as the product *k*_inact_ × *t*_50_ has a particular value *and* the ratio 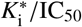 has a particular value.

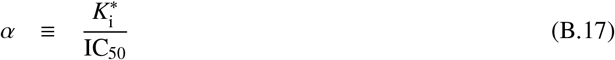

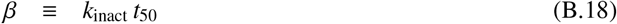

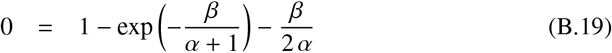

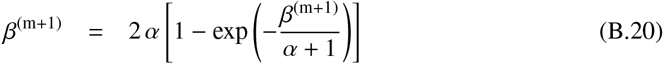

In order to discover which pairs of *k*_inact_×*t*_50_ and 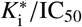 will satisfy Eqn (B.16), the equation was converted into a dimensionless variant form represented by Eqn (B.19). Given a suitably chosen value of *α* the corresponding value of *β* was computed by using the fixed-point iteration Eqn (B.20). The algorithm was implemented in the Perl language code listed immediately below. The results are summarized in the first two columns of *Table B.1*. Note that *α* > 1 by definition, because 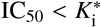.

~~~
use strict;
package main;
$main::itMax = 10000;
$main::relDiffStop = 1e-14;
calculate ();
write_report ();
#-------------------------------------------------------
sub calculate
{
  $main::report = “alpha,alpha-1,beta,zero,ln alpha-1, ln beta\n”;
  my $alphaMinus1 = 0.001;
  my $n = 21;
  my $i = 0;
  for (; $i < $n; ++$i) {
   my $alpha = 1 + $alphaMinus1;
   my $beta = solve_beta ($alpha);
   my $zero = test_zero_alpha_beta ($alpha, $beta);
   my $loga = log($alpha-1);
   my $logb = log($beta);
   $main::report .= “$alpha,$alphaMinus1,$beta,$zero,$loga,$logb\n”;
   $alphaMinus1 _*_= 2;
 }
}
#-------------------------------------------------------
sub solve_beta
{
  my ($alpha) = @_;
  my $beta = 1;
  my $it = 0;
  my $absDiff;
  my $relDiff;
  for (; $it < $main::itMax; ++$it) {
   my $betanew = $alpha*2* (1 - exp(-$beta/($alpha + 1)));
   $absDiff = abs($betanew - $beta);
   $relDiff = $absDiff/$beta;
   if ($relDiff < $main::relDiffStop) {
      last;
   }
   $beta = $betanew;
  }
  return $beta;
}
#-------------------------------------------------------
sub test_zero_alpha_beta
{
  my ($alpha, $beta) = @_;
  my $zero = 1.0;
  $zero -= exp(-$beta/($alpha + 1));
  $zero -= $beta/(2_*_$alpha);
  return $zero;
}
#-------------------------------------------------------
sub write_report
{
  my $path = “alpha-beta-report.csv”;
  open (my $out, “>$path”) or die $!;
  print $out $main::report;
  close ($out);
  print “-> $path\n”;
}
~~~

For purposes of kinetic modeling, the deterministic relationship between *α* and *β* can be empirically described as a straight line in the *X* = ln *α* − 1, *Y* = ln*β* coordinates. The slope and intercept of the empirical linear model was determined by using the software package DynaFit [8], according to the input script file listed immediately below. The best-fit values of the slope and intercept, respectively, were *a* = 0.9779 and *b* = 0.5850. The results of fit are summarized graphically in *Figure B.1*.

~~~
[task]
  task = fit
  data = generic
[parameters]
~~~

**Table B.1:** Pairs of *α* and *β* values that satisfy Eqn (B.19), generated by the fixed-point iteration formula Eqn (B.20). The “scientific notation” *E*±*NN* represents × 10±^*NN*^.

| $\alpha$ | $\beta$ | $f \approx 0$ | $\ln(\alpha - 1)$ | $\ln\beta$ |
| --- | --- | --- | --- | --- |
| 1.001 | 2.01323E-03 | -3.41E-09 | -6.90776 | -6.20801 |
| 1.002 | 3.99885E-03 | -9.10E-11 | -6.21461 | -5.52175 |
| 1.004 | 7.99468E-03 | -1.70E-14 | -5.52146 | -4.82898 |
| 1.008 | 1.59788E-02 | 0.00E+00 | -4.82831 | -4.13649 |
| 1.016 | 3.19158E-02 | -1.11E-16 | -4.13517 | -3.44465 |
| 1.032 | 6.36675E-02 | -2.26E-16 | -3.44202 | -2.75408 |
| 1.064 | 1.26704E-01 | -5.55E-16 | -2.74887 | -2.06590 |
| 1.128 | 2.51064E-01 | -1.12E-15 | -2.05573 | -1.38205 |
| 1.256 | 4.93987E-01 | -1.89E-15 | -1.36258 | -0.70525 |
| 1.512 | 9.62664E-01 | -3.00E-15 | -0.66943 | -0.03805 |
| 2.024 | 1.85877E+00 | 4.00E-15 | 0.02372 | 0.61991 |
| 3.048 | 3.57617E+00 | 4.44E-15 | 0.71686 | 1.27429 |
| 5.096 | 6.91270E+00 | 4.33E-15 | 1.41001 | 1.93336 |
| 9.192 | 1.34911E+01 | 4.66E-15 | 2.10316 | 2.60203 |
| 17.384 | 2.65770E+01 | 4.22E-15 | 2.79631 | 3.28004 |
| 33.768 | 5.27041E+01 | 3.89E-15 | 3.48945 | 3.96469 |
| 66.536 | 1.04933E+02 | 3.55E-15 | 4.18260 | 4.65332 |
| 132.072 | 2.09377E+02 | 5.33E-15 | 4.87575 | 5.34414 |
| 263.144 | 4.18259E+02 | 4.11E-15 | 5.56889 | 6.03610 |
| 525.288 | 8.36020E+02 | 3.55E-15 | 6.26204 | 6.72865 |
| 1049.576 | 1.67154E+03 | 3.33E-15 | 6.95519 | 7.42150 |

~~~
  x, a, b
[model]
  a = 1 ??
  b = 0.1 ??
  y = a*x + b
[data]
  variable x
graph ln {/Symbol b} = 0.9779 ln ({/Symbol a} - 1) +  0.5850
  set data
[output]
  directory ./TN/2020/03/output/fit-001
[settings]
{ConfidenceIntervals}
  LevelPercent = 99
{Output}
  XAxisLabel = ln ({/Symbol a} - 1)
  YAxisLabel = ln {/Symbol b}
[set:data]
ln(alpha-1) ln beta
-6.90776 -6.20801
-6.21461 -5.52175
-5.52146 -4.82898
-4.82831 -4.13649
-4.13517 -3.44465
-3.44202 -2.75408
-2.74887 -2.0659
-2.05573 -1.38205
-1.36258 -0.70525
-0.66943 -0.03805
0.02372 0.61991
0.71686 1.27429
1.41001 1.93336
2.10316 2.60203
2.79631 3.28004
3.48945 3.96469
4.1826 4.65332
4.87575 5.34414
5.56889 6.0361
6.26204 6.72865
6.95519 7.4215
[end]
~~~

**Figure B.1:**
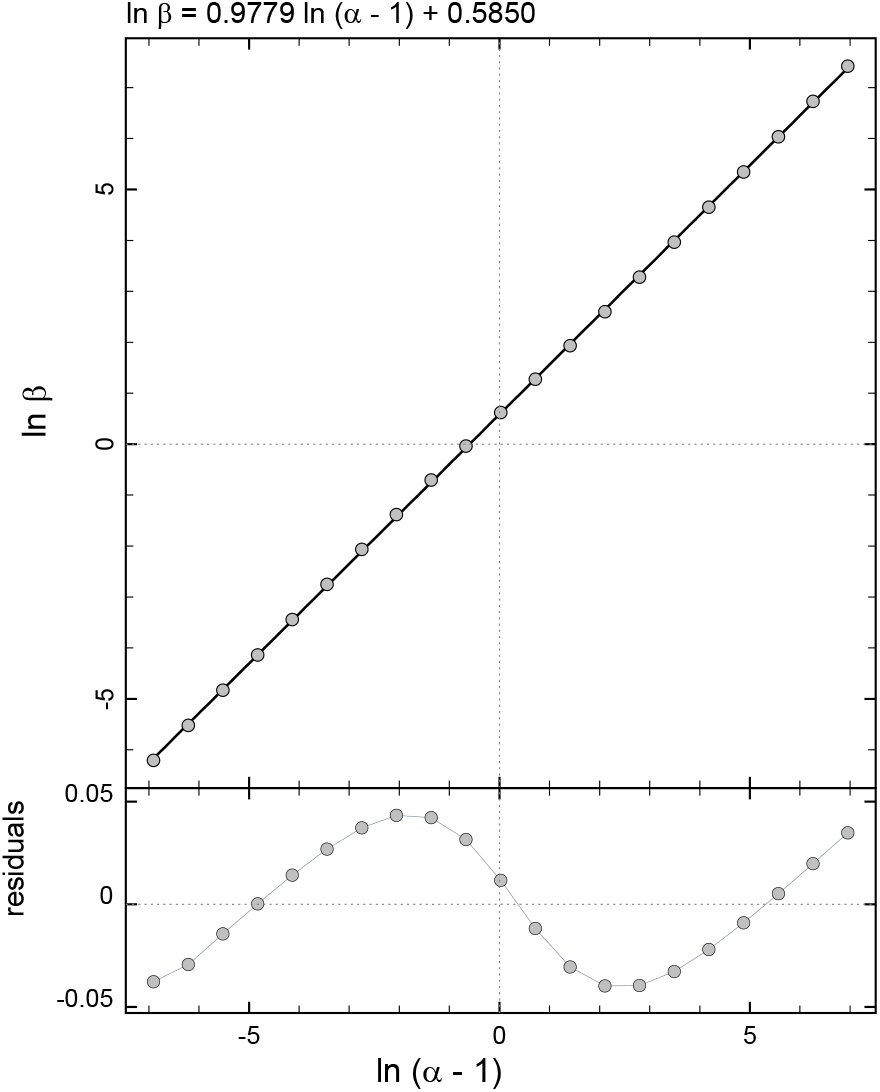
Results of linear least-squares fit of ln *β* vs. ln(*α* − 1) to determine the empirical coefficients *a* and *b*.

Thus, given any arbitrary numerical values of 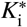, IC_50_ and *t*_50_, the corresponding *k*_inact_ value that satisfies the implicit algebraic Eqn (B.16) can be computed by using Eqn (B.23). Similarly, given any arbitrary numerical values of *k*_inact_, IC_50_ and *t*_50_, the corresponding 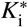 value can be computed by using Eqn (B.24).

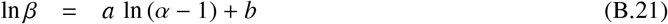

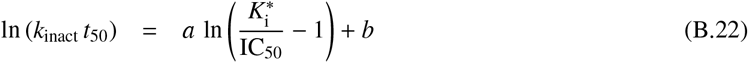

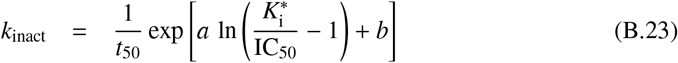

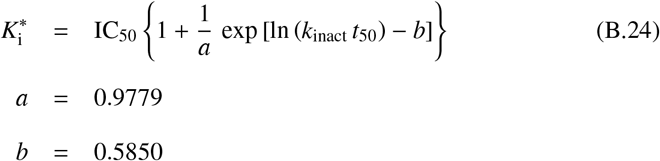

Let us now consider an experimental scenario involving two independent determinations of IC_50_ (referred to below as 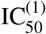 and 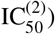) obtained at two different reaction times 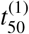 and 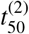, respectively. Treating these four experimentally determined values as fixed constants, we can now rewrite Eqn (B.23) as a system of two simultaneous nonlinear algebraic equations for two unknowns *k*_inact_ and 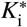, as shown in Eqns (B.25)–(B.26).

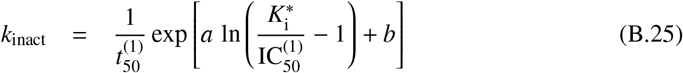

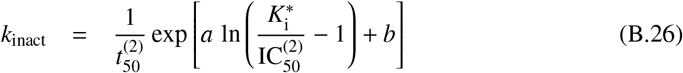

The nonlinear algebraic system of Eqns (B.25)–(B.26) can be solved as follows. First, we can very simply eliminate *k*_inact_ by setting up an equality of the right-hand sides of Eqns (B.25)–(B.26). Next, we can solve for 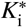 as is shown below. The final result shown as Eqn (5) represents the fact that given any two measurements of IC_50_ (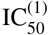 and 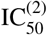) performed at two different reaction times (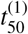 and 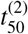) we can immediately estimate the inhibition constant *K*_i_ from those two experimental results alone.

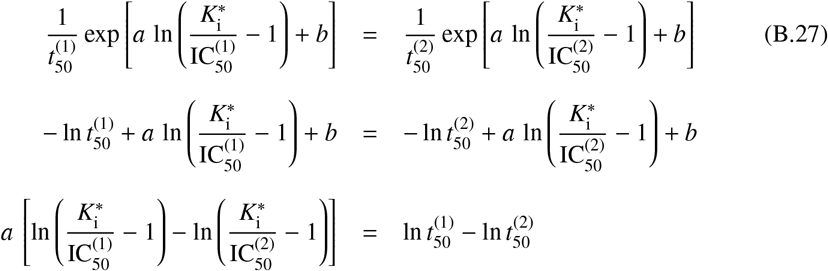

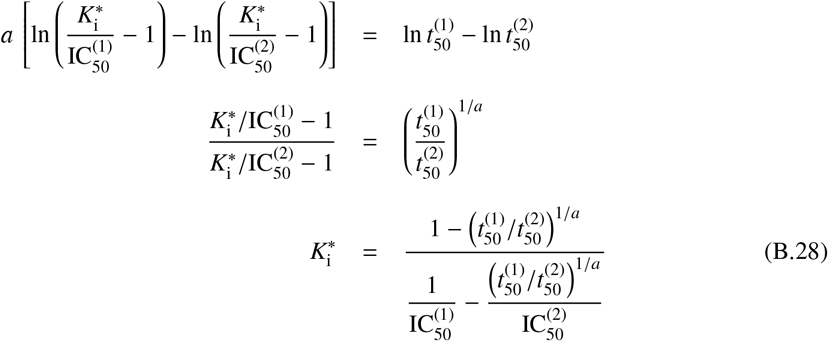

## Footnotes

1 Throughout this manuscript, the conventionally used notation IC_50_ is abbreviated as IC_50_.

2 Recall that the empirical constants *a* and *b* in Eqn (6) strictly require that all concentrations be expressed in micromoles per liter and the reaction time is in seconds.

