## Supporting Information for "A two-point IC_50_ method for evaluating the biochemical potency of irreversible enzyme inhibitors"

Petr Kuzmič

*BioKin Ltd., Watertown, Massachusetts, USA*
*http://www.biokin.com*

**References** **133**

#### 1. Input script files for the DynaFit software package

The listing below shows the complete text of input script files for the DynaFit [1] software package.

##### *1.1. Simulation of enzymatic progress curves*

```
;________________________________________________________________________________
[task]
   task = simulate
   data = progress
   model = C2Sc
[mechanism]
   E + S <==> E.S       :    ka.S     kd.S
   E.S ---> E + P       :    kd.P
   E + I <==> E.I       :    ka.I      kd.I
   E.I ---> EI          :    k.for
[constants]
   ka.S = 1
   kd.S = 1
   kd.P = 1
   ka.I = 10
   kd.I = 1
   k.for = 0.1
[concentrations]
   E = 1e-006
   S = 2
[responses]
   P = 10000
[data]
   mesh from 900 to 14400 step 2 logarithmic
   error constant 0.5 percent
   delay 0
   directory ./project-directory/data
   sheet simul-I01-R1.csv
   monitor E, E.S, E.I, EI
graph ABC-001 :: Enz1 :: R1
   column 2  | conc I =  0.022        | label  A01
   column 3  | conc I =  0.011        | label  A02
   column 4  | conc I =  0.0055        | label  A03
   column 5  | conc I =  0.00275        | label  A04
   column 6  | conc I =  0.001375        | label  A05
   column 7  | conc I =  0.0006875        | label  A06
   column 8  | conc I =  0.00034375        | label  A07
   column 9  | conc I =  0.000171875        | label  A08
   column 10  | conc I =  8.59375e-005        | label  A09
   column 11  | conc I =  4.29688e-005        | label  A10
   column 12  | conc I =  2.14844e-005        | label  A11
   column 13  | conc I =  0        | label  A12
[output]
   directory ./project-directory/output/Sub-MM-Inh-C2Sc-I01-R1
[settings]
```

```
{Output}
   XAxisLabel = t, sec
   YAxisLabel = {/Symbol D}F, rfu
{DynaFit}
   RandomizationSeed = 0
[end]
;_______________________________________________________________________
;_______________________________________________________________________
```

##### 1.2. *Determination of $I_{50}$*

```
[task]
   task = fit
   data = generic
[parameters]
   Io, Fo, Ic50, n
[model]
   F = Fo/(1 + (Io/Ic50)^n)
[data]
   variable Io
   directory ./project-directory/data
graph ABC-001
   sheet simtr-I01-R1.csv
   plot logarithmic
   column 2 | param Ic50 = 0.01 ?, n = 1 ?, Fo = 3 ? | label t=900
   column 3 | param Ic50 = 0.01 ?, n = 1 ?, Fo = 6 ? | label t=1800
   column 4 | param Ic50 = 0.01 ?, n = 1 ?, Fo = 12 ? | label t=3600
   column 5 | param Ic50 = 0.001 ?, n = 1 ?, Fo = 24 ? | label t=7200
   column 6 | param Ic50 = 0.001 ?, n = 1 ?, Fo = 48 ? | label t=14400
[output]
   directory ./project-directory/output/FitIc50-I01-R1
[settings]
{Output}
   XAxisLabel = [I]_0, {/Symbol m}M
   YAxisLabel = F, a.u.
[end]
```

#### 2. Detailed results for individual simulated compounds

##### *2.1. Compound No. 1*

*Cpd.* **1***: Simulated data*

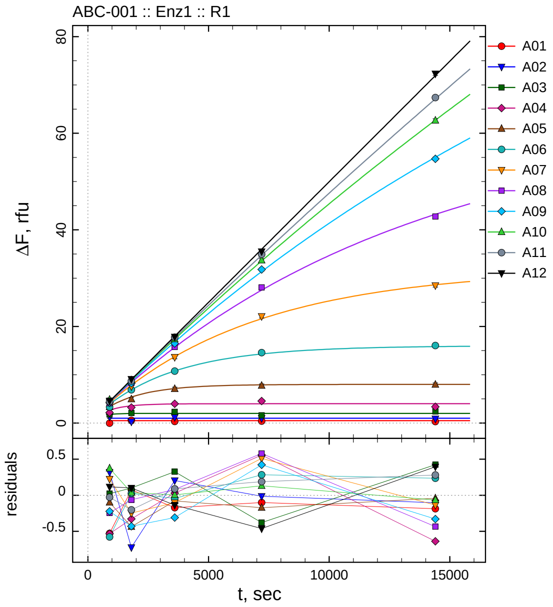

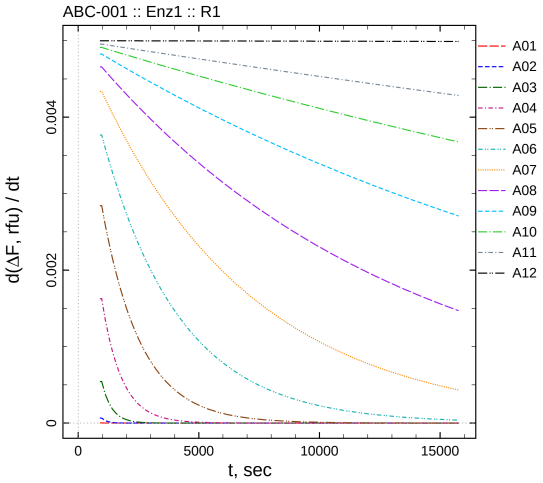

| $[I]_0$, nM | t = 15 | 30 | 60 | 120 | 240 min |
|---:|---:|---:|---:|---:|---:|
| 22 | -0.029 | 0.576 | 0.328 | 0.399 | 0.314 |
| 11 | 1.307 | 0.278 | 1.204 | 0.988 | 0.895 |
| 5.5 | 1.800 | 2.077 | 2.328 | 1.621 | 2.425 |
| 2.75 | 2.150 | 3.240 | 3.982 | 4.557 | 3.362 |
| 1.375 | 3.327 | 4.950 | 7.067 | 7.738 | 7.958 |
| 0.6875 | 3.333 | 6.893 | 10.744 | 14.589 | 16.062 |
| 0.34375 | 4.417 | 7.586 | 13.648 | 22.105 | 28.488 |
| 0.171875 | 4.098 | 8.326 | 15.758 | 28.093 | 42.770 |
| 0.0859375 | 4.196 | 8.256 | 16.473 | 31.783 | 54.703 |
| 0.0429688 | 4.833 | 8.876 | 17.380 | 33.695 | 62.647 |
| 0.0214844 | 4.450 | 8.716 | 17.769 | 34.934 | 67.391 |
| 0 | 4.615 | 9.099 | 17.858 | 35.520 | 72.330 |

*Cpd.* **1***: Determination of* $IC_{50}$ *and* $k_1^*$

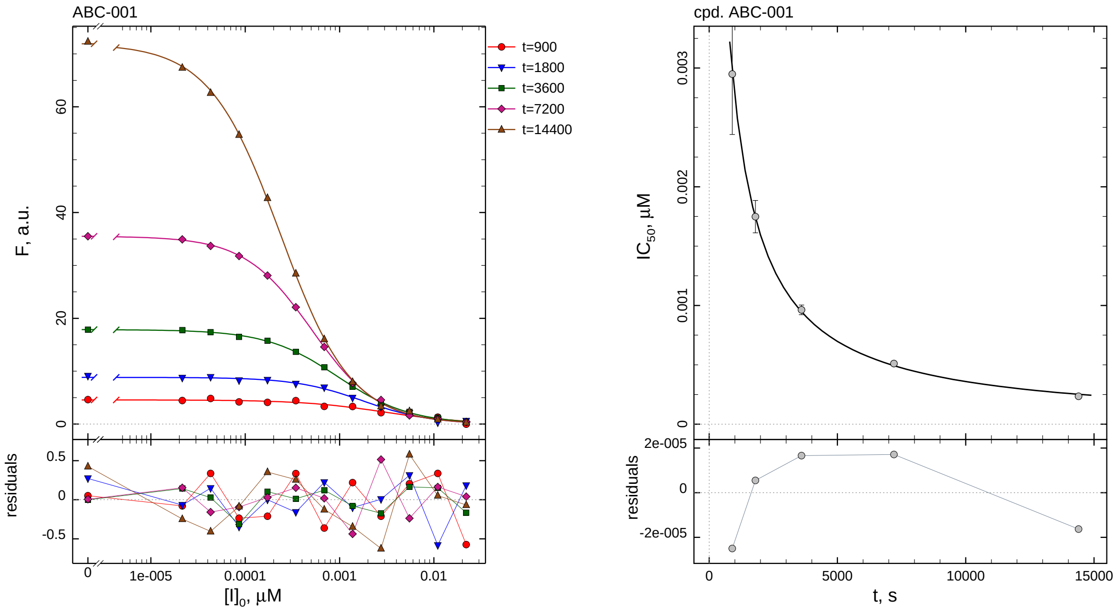

| $i$ | $t$, min | $I_{50}$, nM | ± | std.err. | CV, % | $k_1^*$, $mM^{-1}s^{-1}$ |
|---|---|---|---|---|---|---|
| 1 | 15 | 2.949 | ± | 0.5086 | 17.2 | 600.4 |
| 2 | 30 | 1.748 | ± | 0.1362 | 7.8 | 506.4 |
| 3 | 60 | 0.9632 | ± | 0.04070 | 4.2 | 459.6 |
| 4 | 120 | 0.5104 | ± | 0.01074 | 2.1 | 433.6 |
| 5 | 240 | 0.2355 | ± | 0.002748 | 1.2 | 469.9 |

*Cpd.* **1***: Mechanistic analysis*

| parameter | unit | "true" (T) | calculated (C) | C/T ratio | note |
|---|---|---|---|---|---|
| mechanism | | **C2S** | **C1** | | |
| $k_{\mathrm{eff}}$ | $mM^{-1}s^{-1}$ | 909.1 | 867.2 | 0.95 | $= k_1^* \,(1 + [S]_0/K_M)$ |

| | |
|---|---|
| Reaction times used for analysis: | 30 and 120 min |
| Maximum GSD for accepting one-step model **C1**: | 1.25 |
| Observed GSD: | 1.12 |
| Assumed $[S]_0/K_M$ ratio: | 1.0 |

*2.2. Compound No. 2*

*Cpd.* **2***: Simulated data*

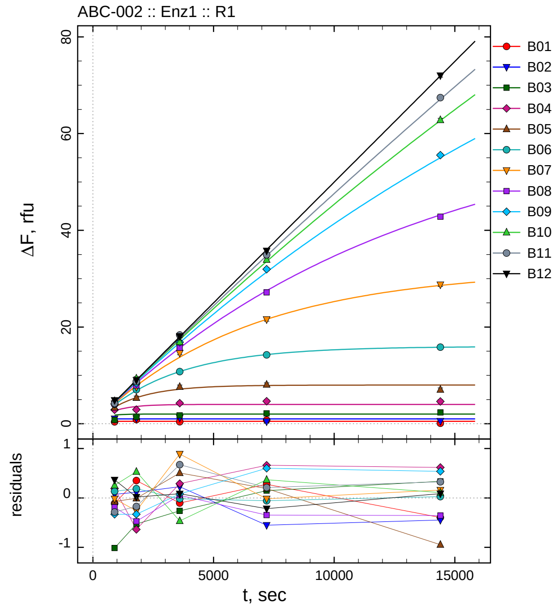

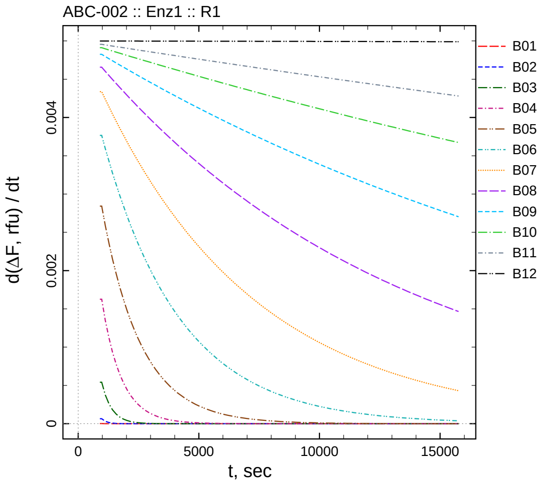

| $[I]_0$, nM | t = 15 | 30 | 60 | 120 | 240 min |
|---:|---:|---:|---:|---:|---:|
| 220 | 0.390 | 0.852 | 0.396 | 0.775 | 0.104 |
| 110 | 1.060 | 1.114 | 1.227 | 0.449 | 0.554 |
| 55 | 0.765 | 1.446 | 1.739 | 2.147 | 2.337 |
| 27.5 | 2.873 | 2.928 | 4.243 | 4.660 | 4.616 |
| 13.75 | 3.354 | 5.376 | 7.650 | 8.098 | 7.051 |
| 6.875 | 4.045 | 7.054 | 10.768 | 14.241 | 15.843 |
| 3.4375 | 4.174 | 7.572 | 14.637 | 21.569 | 28.761 |
| 1.71875 | 4.155 | 7.915 | 15.728 | 27.161 | 42.819 |
| 0.859375 | 4.088 | 8.355 | 16.863 | 31.958 | 55.547 |
| 0.429688 | 4.713 | 9.372 | 16.907 | 33.929 | 62.806 |
| 0.214844 | 4.194 | 8.746 | 18.353 | 34.955 | 67.429 |
| 0 | 4.865 | 9.016 | 18.079 | 35.766 | 72.021 |

*Cpd.* **2**: *Determination of* $IC_{50}$ *and* $k_1^*$

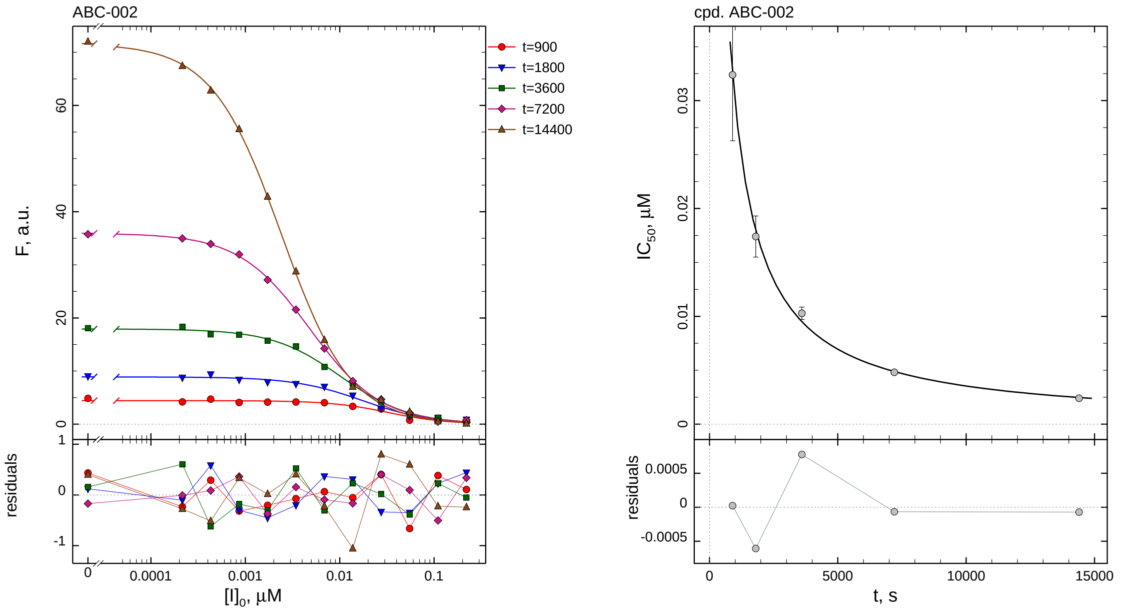

| $i$ | $t$, min | $I_{50}$, nM | ± | std.err. | CV, % | $k_1^*$, mM$^{-1}$s$^{-1}$ |
|---|---|---|---|---|---|---|
| 1 | 15 | 32.39 | ± | 6.118 | 18.9 | 54.66 |
| 2 | 30 | 17.41 | ± | 1.903 | 10.9 | 50.87 |
| 3 | 60 | 10.28 | ± | 0.5767 | 5.6 | 43.05 |
| 4 | 120 | 4.816 | ± | 0.1482 | 3.1 | 45.96 |
| 5 | 240 | 2.403 | ± | 0.03852 | 1.6 | 46.06 |

*Cpd.* **2**: *Mechanistic analysis*

| parameter | unit | "true" (T) | calculated (C) | C/T ratio | note |
|---|---|---|---|---|---|
| mechanism | | **C2S** | **C1** | | |
| $k_{\text{eff}}$ | mM$^{-1}$s$^{-1}$ | 90.91 | 91.92 | 1.01 | $= k_1^* \, (1 + [S]_0/K_M)$ |

| | |
|---|---|
| Reaction times used for analysis: | 30 and 120 min |
| Maximum GSD for accepting one-step model **C1**: | 1.25 |
| Observed GSD: | 1.07 |
| Assumed $[S]_0/K_M$ ratio: | 1.0 |

#### *2.3. Compound No. 3*

*Cpd.* **3***: Simulated data*

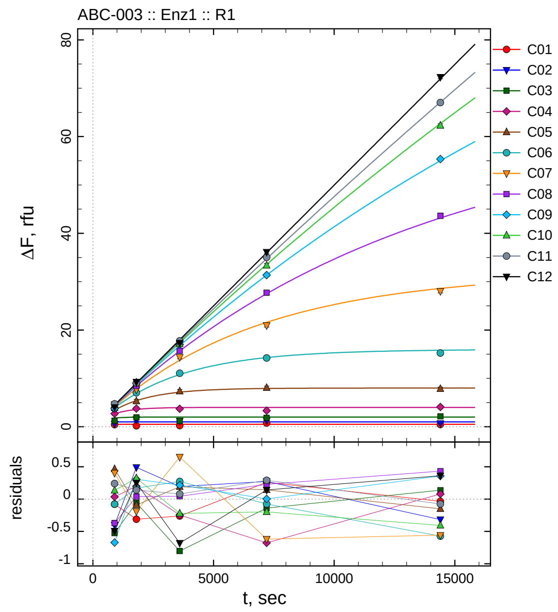

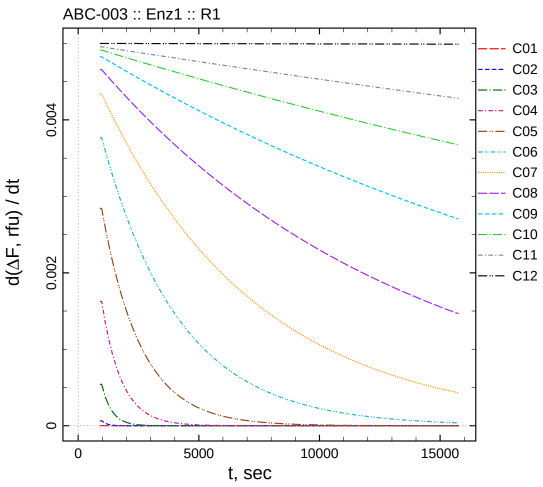

| $[I]_0$, nM | t = 15 | 30 | 60 | 120 | 240 min |
|---:|---:|---:|---:|---:|---:|
| 2200 | 0.421 | 0.189 | 0.239 | 0.761 | 0.466 |
| 1100 | 0.586 | 1.492 | 1.196 | 1.273 | 0.683 |
| 550 | 1.251 | 1.917 | 1.197 | 1.861 | 2.137 |
| 275 | 2.719 | 3.765 | 3.707 | 3.322 | 4.079 |
| 137.5 | 3.893 | 5.295 | 7.336 | 8.050 | 7.847 |
| 68.75 | 3.833 | 7.037 | 11.060 | 14.224 | 15.242 |
| 34.375 | 4.595 | 7.644 | 14.399 | 20.967 | 28.041 |
| 17.1875 | 3.969 | 8.427 | 15.720 | 27.736 | 43.606 |
| 8.59375 | 3.746 | 8.992 | 16.998 | 31.360 | 55.365 |
| 4.29688 | 4.592 | 9.165 | 17.152 | 33.365 | 62.289 |
| 2.14844 | 4.719 | 9.059 | 17.759 | 35.034 | 67.025 |
| 0 | 4.005 | 9.249 | 17.314 | 36.124 | 72.296 |

*Cpd.* **3***: Determination of* $IC_{50}$ *and* $k_1^*$

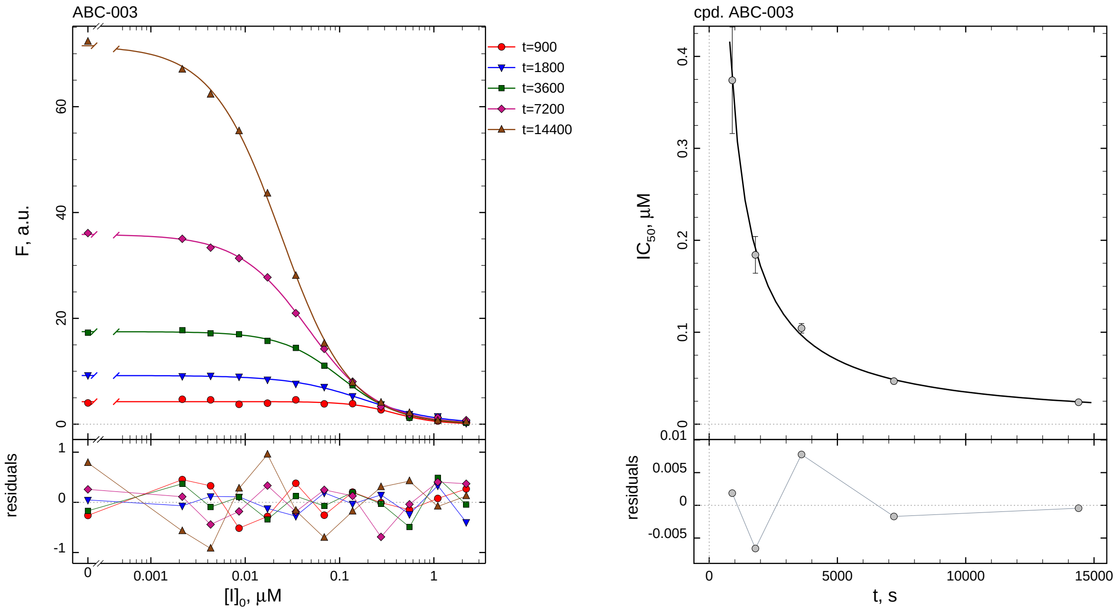

| $i$ | $t$, min | $I_{50}$, nM ± std.err. | CV, % | $k_1^*$, mM$^{-1}$s$^{-1}$ |
|---|---|---|---|---|
| 1 | 15 | 374.1 ± 57.95 | 15.5 | 4.734 |
| 2 | 30 | 184.2 ± 20.01 | 10.9 | 4.807 |
| 3 | 60 | 104.3 ± 5.165 | 5.0 | 4.243 |
| 4 | 120 | 46.86 ± 1.356 | 2.9 | 4.723 |
| 5 | 240 | 23.92 ± 0.3649 | 1.5 | 4.627 |

*Cpd.* **3***: Mechanistic analysis*

| parameter | unit | "true" (T) | calculated (C) | C/T ratio | note |
|---|---|---|---|---|---|
| mechanism | | **C2S** | **C1** | | |
| $k_{\text{eff}}$ | mM$^{-1}$s$^{-1}$ | 9.091 | 9.446 | 1.04 | $= k_1^* \, (1 + [\text{S}]_0/K_\text{M})$ |

| | |
|---|---|
| Reaction times used for analysis: | 30 and 120 min |
| Maximum GSD for accepting one-step model **C1**: | 1.25 |
| Observed GSD: | 1.01 |
| Assumed $[\text{S}]_0/K_\text{M}$ ratio: | 1.0 |

*2.4. Compound No. 4*

*Cpd.* **4***: Simulated data*

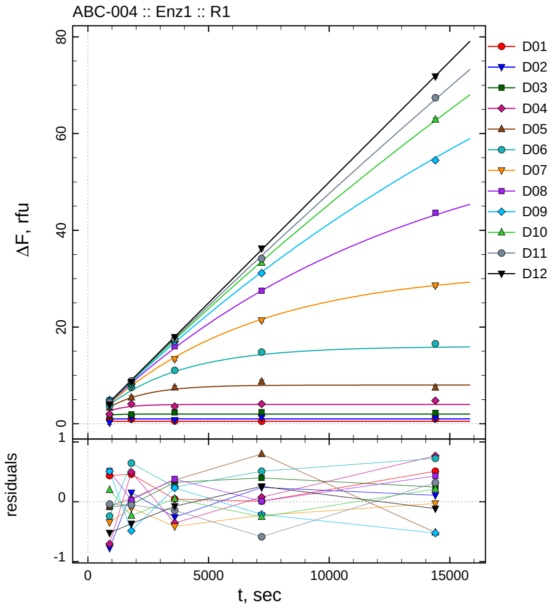

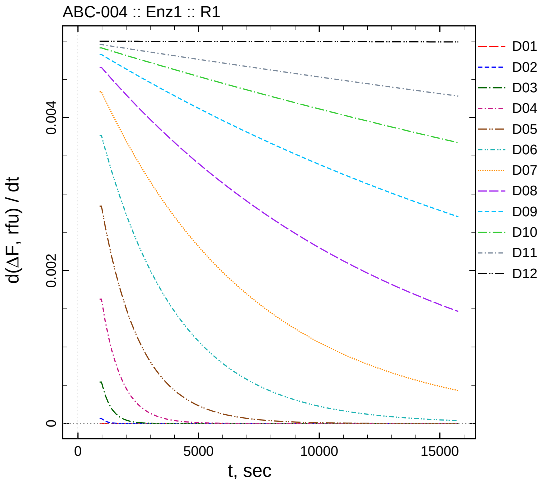

| $[I]_0$, nM | t = 15 | 30 | 60 | 120 | 240 min |
|---|---|---|---|---|---|
| 22000 | 0.939 | 0.962 | 0.545 | 0.511 | 1.012 |
| 11000 | 0.214 | 1.151 | 0.744 | 1.241 | 1.109 |
| 5500 | 1.691 | 1.929 | 2.324 | 2.400 | 2.247 |
| 2750 | 1.981 | 4.065 | 3.597 | 4.076 | 4.768 |
| 1375 | 3.353 | 5.440 | 7.508 | 8.706 | 7.486 |
| 687.5 | 3.672 | 7.514 | 11.032 | 14.810 | 16.540 |
| 343.75 | 3.848 | 7.731 | 13.336 | 21.361 | 28.572 |
| 171.875 | 4.856 | 8.419 | 16.056 | 27.517 | 43.606 |
| 85.9375 | 4.929 | 8.200 | 17.012 | 31.146 | 54.484 |
| 42.9688 | 4.656 | 8.607 | 17.426 | 33.311 | 62.910 |
| 21.4844 | 4.440 | 8.863 | 17.538 | 34.159 | 67.419 |
| 0 | 3.978 | 8.631 | 17.921 | 36.241 | 71.820 |

*Cpd.* **4***: Determination of* $IC_{50}$ *and* $k_1^*$

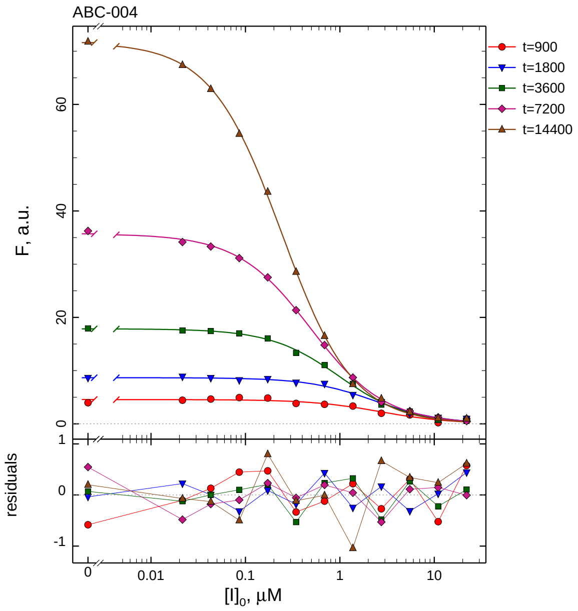

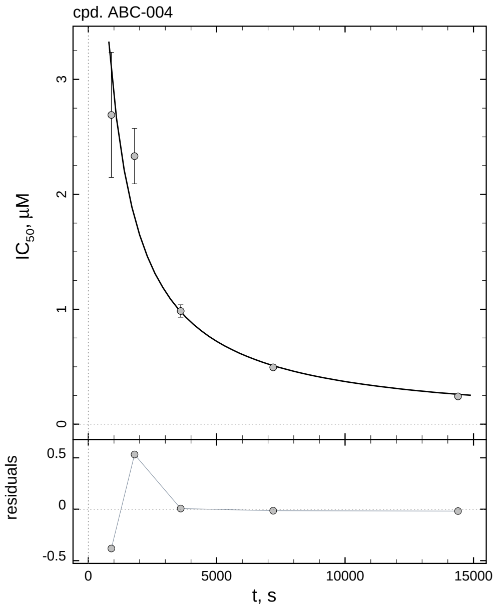

| $i$ | $t$, min | $I_{50}$, nM | ± | std.err. | CV, % | $k_1^*$, $mM^{-1}s^{-1}$ |
|---|---|---|---|---|---|---|
| 1 | 15 | 2690.8 | ± | 544.3 | 20.2 | 0.6581 |
| 2 | 30 | 2332.5 | ± | 240.7 | 10.3 | 0.3796 |
| 3 | 60 | 984.9 | ± | 53.48 | 5.4 | 0.4495 |
| 4 | 120 | 494.8 | ± | 15.05 | 3.0 | 0.4474 |
| 5 | 240 | 242.3 | ± | 3.781 | 1.6 | 0.4567 |

*Cpd.* **4***: Mechanistic analysis*

| parameter | unit | "true" (T) | calculated (C) | C/T ratio | note |
|---|---|---|---|---|---|
| mechanism | | **C2S** | **C1** | | |
| $k_{\mathrm{eff}}$ | $mM^{-1}s^{-1}$ | 0.9091 | 0.8947 | 0.98 | $= k_1^* \, (1 + [S]_0/K_M)$ |

| | |
|---|---|
| Reaction times used for analysis: | 30 and 120 min |
| Maximum GSD for accepting one-step model **C1**: | 1.25 |
| Observed GSD: | 1.12 |
| Assumed $[S]_0/K_M$ ratio: | 1.0 |

*2.5. Compound No. 5*

*Cpd.* ***5****: Simulated data*

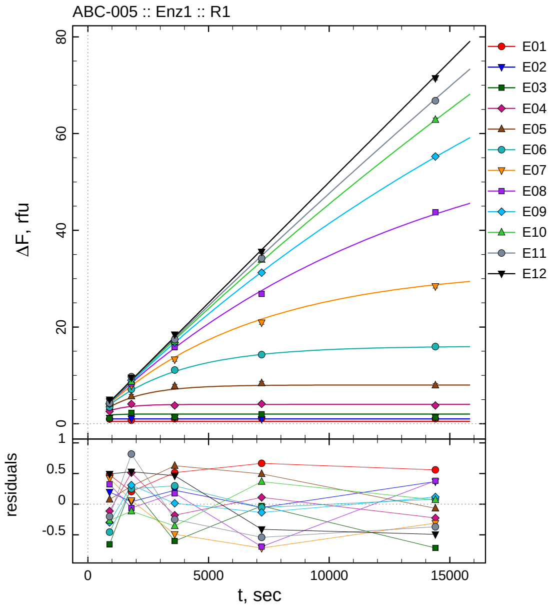

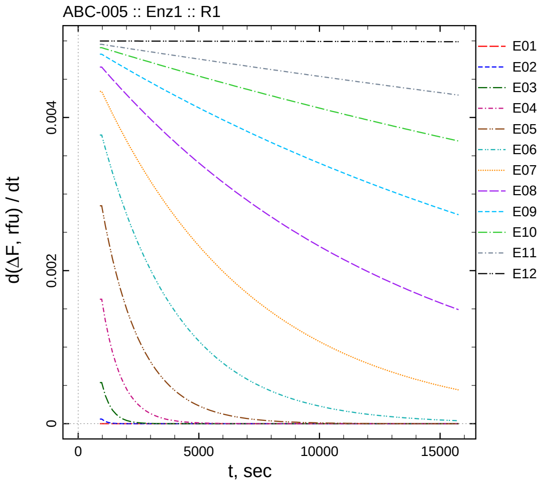

| $[I]_0$, nM | t = 15 | 30 | 60 | 120 | 240 min |
|---:|---:|---:|---:|---:|---:|
| 4 | 0.979 | 0.706 | 1.015 | 1.167 | 1.059 |
| 2 | 1.186 | 1.040 | 1.227 | 0.945 | 1.369 |
| 1 | 1.131 | 2.257 | 1.400 | 1.974 | 1.287 |
| 0.5 | 2.579 | 4.087 | 3.779 | 4.114 | 3.780 |
| 0.25 | 3.510 | 5.690 | 7.785 | 8.416 | 7.946 |
| 0.125 | 3.461 | 7.129 | 11.104 | 14.282 | 15.955 |
| 0.0625 | 4.615 | 7.901 | 13.273 | 20.921 | 28.428 |
| 0.03125 | 4.668 | 8.330 | 15.864 | 26.857 | 43.730 |
| 0.015625 | 4.121 | 8.999 | 16.803 | 31.249 | 55.273 |
| 0.0078125 | 4.192 | 8.727 | 17.022 | 33.944 | 62.864 |
| 0.00390625 | 4.277 | 9.736 | 17.434 | 34.212 | 66.788 |
| 0 | 4.991 | 9.527 | 18.458 | 35.572 | 71.442 |

*Cpd.* **5***: Determination of* $IC_{50}$ *and* $k_1^*$

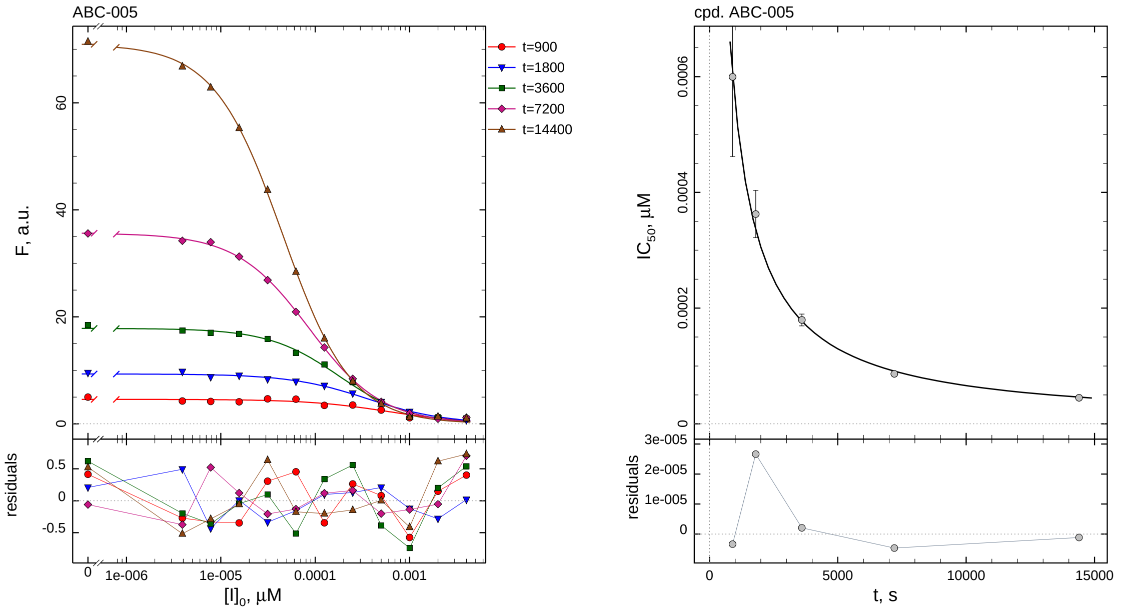

| $i$ | $t$, min | $I_{50}$, nM | ± | std.err. | CV, % | $k_1^*$, $\mathrm{mM}^{-1}\mathrm{s}^{-1}$ |
|---|---|---|---|---|---|---|
| 1 | 15 | 0.5996 | ± | 0.1377 | 23.0 | 2953.3 |
| 2 | 30 | 0.3626 | ± | 0.04093 | 11.3 | 2441.9 |
| 3 | 60 | 0.1795 | ± | 0.01022 | 5.7 | 2466.2 |
| 4 | 120 | 0.08650 | ± | 0.002685 | 3.1 | 2558.8 |
| 5 | 240 | 0.04505 | ± | 0.0007057 | 1.6 | 2456.7 |

*Cpd.* **5***: Mechanistic analysis*

| parameter | unit | "true" (T) | calculated (C) | C/T ratio | note |
|---|---|---|---|---|---|
| mechanism | | **C2S** | **C1** | | |
| $k_{\mathrm{eff}}$ | $\mathrm{mM}^{-1}\mathrm{s}^{-1}$ | 5000.0 | 5117.5 | 1.02 | $= k_1^* \, (1 + [\mathrm{S}]_0/K_\mathrm{M})$ |

| | |
|---|---|
| Reaction times used for analysis: | 30 and 120 min |
| Maximum GSD for accepting one-step model **C1**: | 1.25 |
| Observed GSD: | 1.03 |
| Assumed $[\mathrm{S}]_0/K_\mathrm{M}$ ratio: | 1.0 |

2.6. *Compound No. 6*

*Cpd.* **6**: *Simulated data*

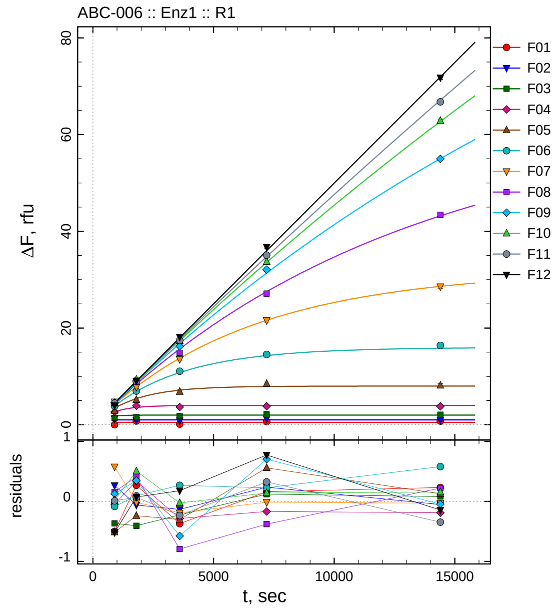

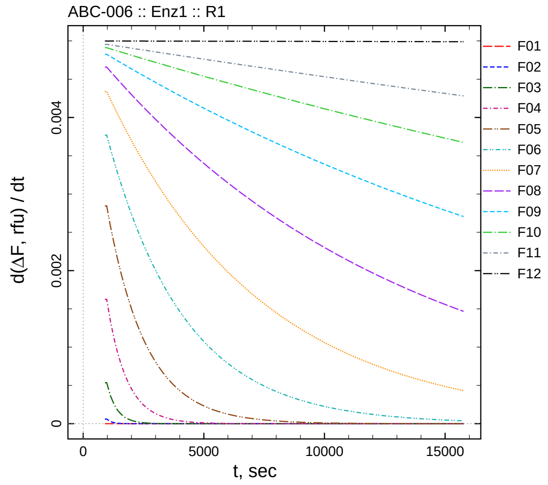

| $[I]_0$, nM | t = 15 | 30 | 60 | 120 | 240 min |
|---:|---:|---:|---:|---:|---:|
| 40 | 0.001 | 0.769 | 0.129 | 0.662 | 0.733 |
| 20 | 1.253 | 0.940 | 0.867 | 1.238 | 0.946 |
| 10 | 1.415 | 1.572 | 1.774 | 2.122 | 2.082 |
| 5 | 2.646 | 3.911 | 3.670 | 3.832 | 3.812 |
| 2.5 | 2.919 | 5.154 | 6.849 | 8.470 | 8.126 |
| 1.25 | 3.832 | 6.967 | 11.064 | 14.533 | 16.402 |
| 0.625 | 4.772 | 7.823 | 13.565 | 21.585 | 28.588 |
| 0.3125 | 4.499 | 8.806 | 14.888 | 27.139 | 43.420 |
| 0.15625 | 4.544 | 9.037 | 16.209 | 32.069 | 54.983 |
| 0.078125 | 4.479 | 9.345 | 17.350 | 33.706 | 62.865 |
| 0.0390625 | 4.488 | 8.987 | 17.447 | 35.071 | 66.763 |
| 0 | 3.982 | 9.072 | 18.173 | 36.756 | 71.790 |

*Cpd.* **6**: *Determination of* $IC_{50}$ *and* $k_1^*$

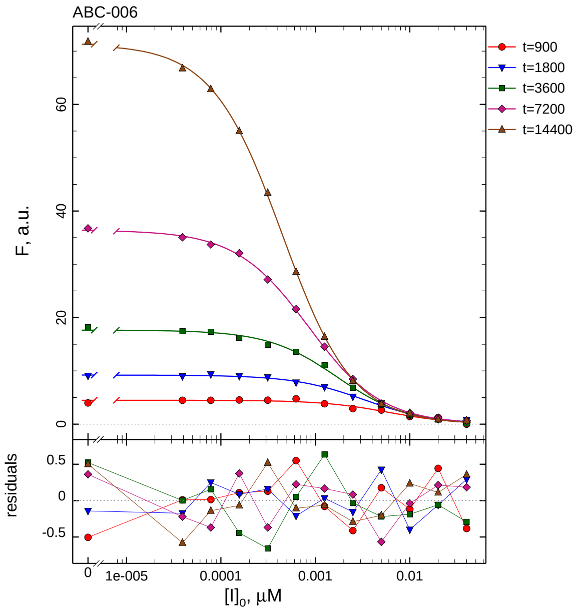

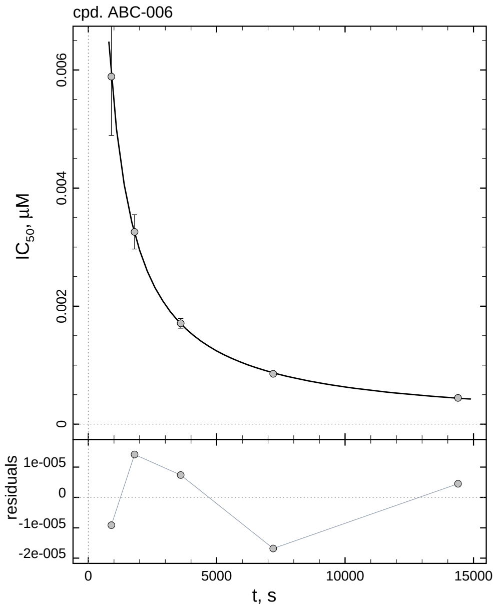

| $i$ | $t$, min | $I_{50}$, nM | ± | std.err. | CV, % | $k_1^*$, mM$^{-1}$s$^{-1}$ |
|---|---|---|---|---|---|---|
| 1 | 15 | 5.888 | ± | 0.9987 | 17.0 | 300.7 |
| 2 | 30 | 3.258 | ± | 0.2911 | 8.9 | 271.8 |
| 3 | 60 | 1.709 | ± | 0.08229 | 4.8 | 259.0 |
| 4 | 120 | 0.8544 | ± | 0.02206 | 2.6 | 259.0 |
| 5 | 240 | 0.4453 | ± | 0.006001 | 1.3 | 248.5 |

*Cpd.* **6**: *Mechanistic analysis*

| parameter | unit | "true" (T) | calculated (C) | C/T ratio | note |
|---|---|---|---|---|---|
| mechanism | | **C2S** | **C1** | | |
| $k_{\text{eff}}$ | mM$^{-1}$s$^{-1}$ | 500.0 | 518.1 | 1.04 | $= k_1^* \, (1 + [S]_0/K_M)$ |

| | |
|---|---|
| Reaction times used for analysis: | 30 and 120 min |
| Maximum GSD for accepting one-step model **C1**: | 1.25 |
| Observed GSD: | 1.03 |
| Assumed $[S]_0/K_M$ ratio: | 1.0 |

#### *2.7. Compound No. 7*

*Cpd.* **7***: Simulated data*

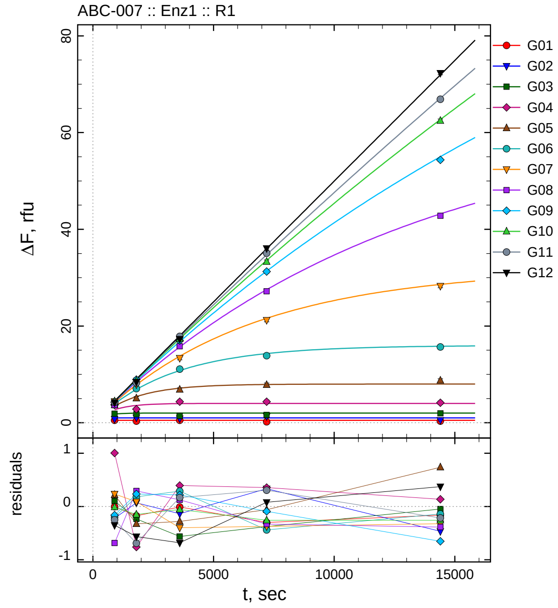

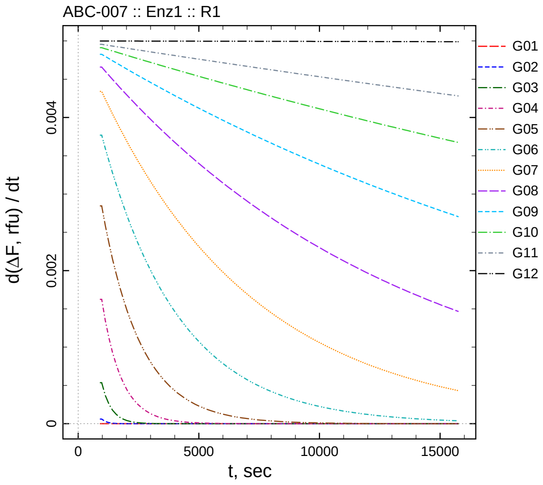

| $[I]_0$, nM | t = 15 | 30 | 60 | 120 | 240 min |
|---:|---:|---:|---:|---:|---:|
| 400 | 0.498 | 0.326 | 0.492 | 0.201 | 0.340 |
| 200 | 0.768 | 1.068 | 0.865 | 1.334 | 0.535 |
| 100 | 1.887 | 1.744 | 1.438 | 1.623 | 1.952 |
| 50 | 3.696 | 2.812 | 4.348 | 4.355 | 4.136 |
| 25 | 3.654 | 5.066 | 6.869 | 7.849 | 8.735 |
| 12.5 | 3.714 | 7.056 | 11.083 | 13.863 | 15.676 |
| 6.25 | 4.429 | 7.928 | 13.357 | 21.221 | 28.281 |
| 3.125 | 3.657 | 8.684 | 15.805 | 27.185 | 42.797 |
| 1.5625 | 4.259 | 8.922 | 17.003 | 31.270 | 54.362 |
| 0.78125 | 4.449 | 8.692 | 17.320 | 33.307 | 62.443 |
| 0.390625 | 4.229 | 8.229 | 17.850 | 35.053 | 66.888 |
| 0 | 4.147 | 8.433 | 17.314 | 36.062 | 72.307 |

*Cpd.* **7**: *Determination of* $IC_{50}$ *and* $k_1^*$

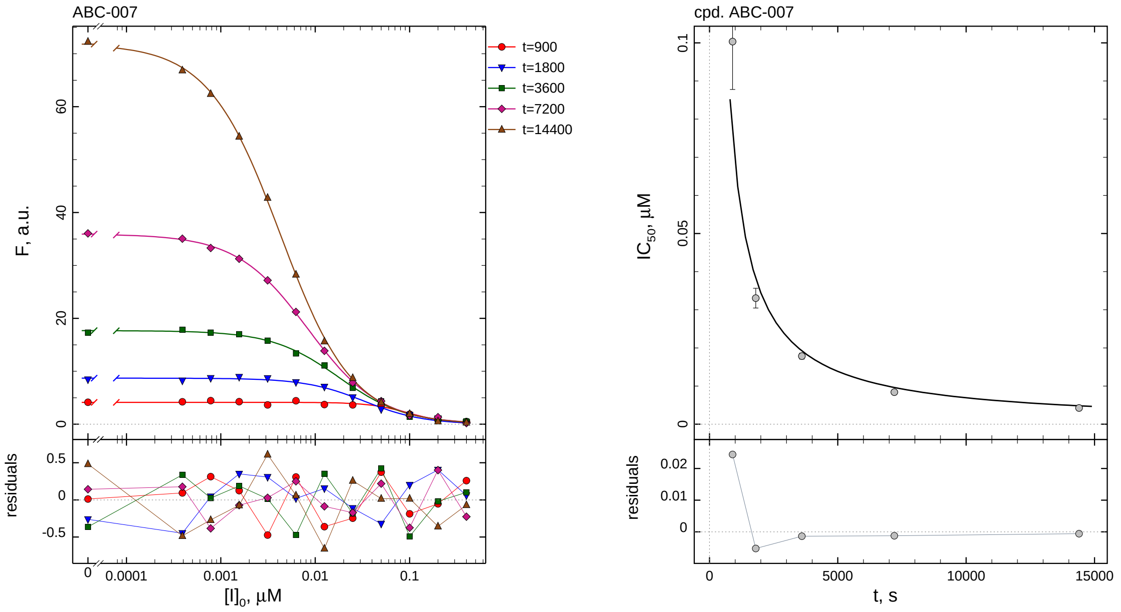

| $i$ | $t$, min | $I_{50}$, nM ± std.err. | CV, % | $k_1^*$, mM$^{-1}$s$^{-1}$ |
|---|---|---|---|---|
| 1 | 15 | 100.3 ± 12.57 | 12.5 | 17.65 |
| 2 | 30 | 33.05 ± 2.594 | 7.8 | 26.78 |
| 3 | 60 | 17.86 ± 0.7915 | 4.4 | 24.79 |
| 4 | 120 | 8.404 ± 0.2065 | 2.5 | 26.34 |
| 5 | 240 | 4.263 ± 0.05573 | 1.3 | 25.96 |

*Cpd.* **7**: *Mechanistic analysis*

| parameter | unit | "true" (T) | calculated (C) | C/T ratio | note |
|---|---|---|---|---|---|
| mechanism | | **C2S** | **C1** | | |
| $k_{\text{eff}}$ | mM$^{-1}$s$^{-1}$ | 50.00 | 52.67 | 1.05 | $= k_1^* \; (1 + [S]_0/K_M)$ |

| | |
|---|---|
| Reaction times used for analysis: | 30 and 120 min |
| Maximum GSD for accepting one-step model **C1**: | 1.25 |
| Observed GSD: | 1.01 |
| Assumed $[S]_0/K_M$ ratio: | 1.0 |

*2.8. Compound No. 8*

*Cpd.* **8**: *Simulated data*

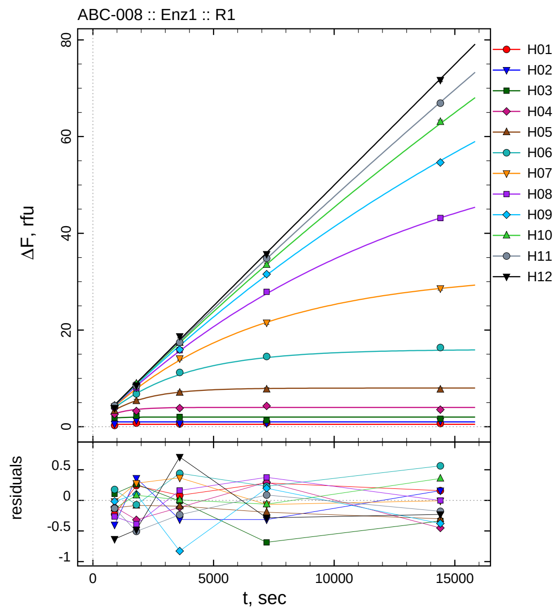

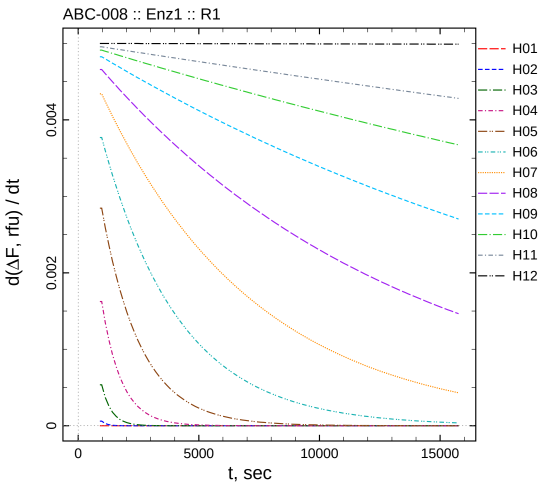

| $[I]_0$, nM | t = 15 | 30 | 60 | 120 | 240 min |
|---|---|---|---|---|---|
| 4000 | 0.273 | 0.741 | 0.581 | 0.784 | 0.660 |
| 2000 | 0.590 | 1.363 | 0.688 | 0.687 | 1.158 |
| 1000 | 1.890 | 2.240 | 1.989 | 1.315 | 1.663 |
| 500 | 2.584 | 3.255 | 3.838 | 4.296 | 3.548 |
| 250 | 3.310 | 5.309 | 7.055 | 7.714 | 7.697 |
| 125 | 4.094 | 6.802 | 11.236 | 14.533 | 16.378 |
| 62.5 | 4.176 | 8.119 | 14.124 | 21.525 | 28.597 |
| 31.25 | 4.078 | 8.002 | 15.843 | 27.890 | 43.179 |
| 15.625 | 4.404 | 8.787 | 15.957 | 31.559 | 54.637 |
| 7.8125 | 4.327 | 8.925 | 17.383 | 33.502 | 63.059 |
| 3.90625 | 4.349 | 8.411 | 17.446 | 34.832 | 66.925 |
| 0 | 3.866 | 8.517 | 18.702 | 35.696 | 71.704 |

*Cpd.* **8***: Determination of* $IC_{50}$ *and* $k_1^*$

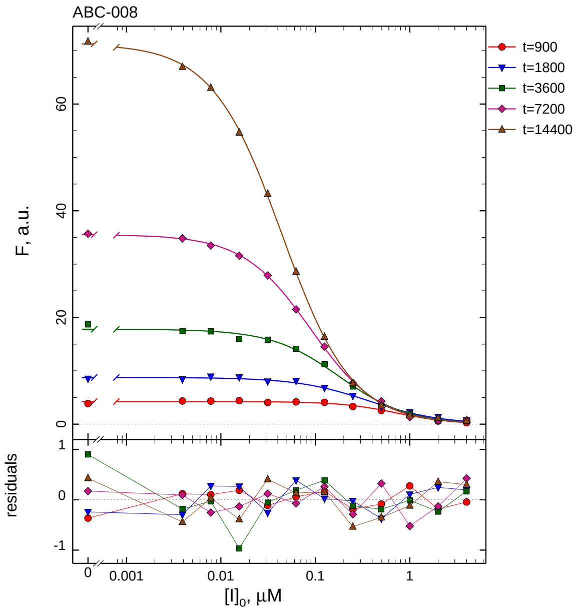

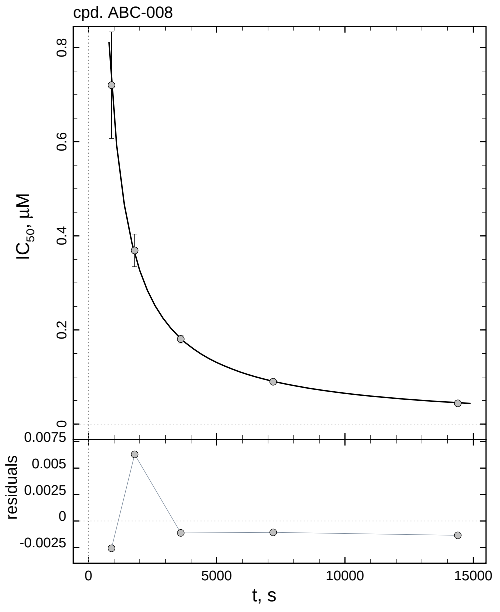

| $i$ | $t$, min | $I_{50}$, nM | ± | std.err. | CV, % | $k_1^*$, $\text{mM}^{-1}\text{s}^{-1}$ |
|---|---|---|---|---|---|---|
| 1 | 15 | 720.1 | ± | 113.0 | 15.7 | 2.459 |
| 2 | 30 | 369.1 | ± | 34.74 | 9.4 | 2.399 |
| 3 | 60 | 180.6 | ± | 8.322 | 4.6 | 2.451 |
| 4 | 120 | 89.89 | ± | 2.196 | 2.4 | 2.462 |
| 5 | 240 | 44.15 | ± | 0.5866 | 1.3 | 2.507 |

*Cpd.* **8***: Mechanistic analysis*

| parameter | unit | "true" (T) | calculated (C) | C/T ratio | note |
|---|---|---|---|---|---|
| mechanism | | **C2S** | **C1** | | |
| $k_{\text{eff}}$ | $\text{mM}^{-1}\text{s}^{-1}$ | 5.000 | 4.925 | 0.98 | $= k_1^* \, (1 + [\text{S}]_0/K_\text{M})$ |

| | |
|---|---|
| Reaction times used for analysis: | 30 and 120 min |
| Maximum GSD for accepting one-step model **C1**: | 1.25 |
| Observed GSD: | 1.02 |
| Assumed $[\text{S}]_0/K_\text{M}$ ratio: | 1.0 |

*2.9. Compound No. 9*

*Cpd.* **9***: Simulated data*

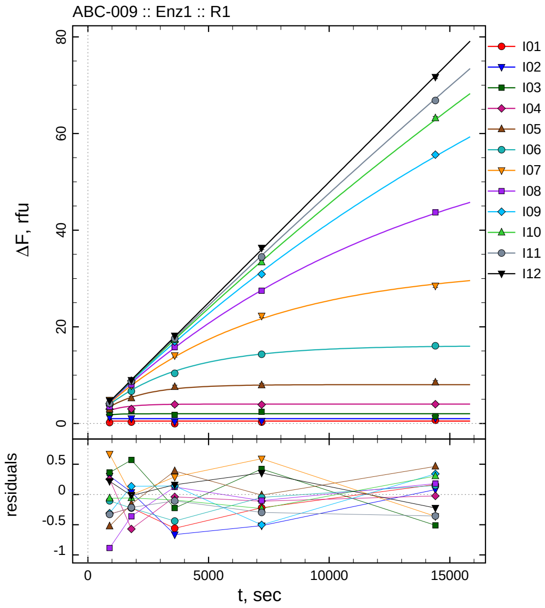

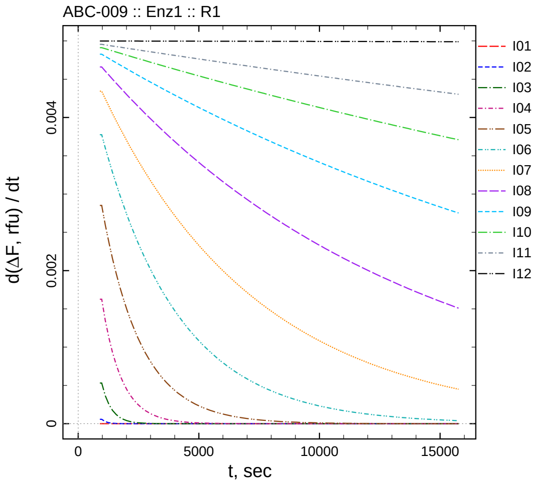

| $[I]_0$, nM | t = 15 | 30 | 60 | 120 | 240 min |
|---:|---:|---:|---:|---:|---:|
| 2.2 | 0.175 | 0.280 | -0.060 | 0.280 | 0.664 |
| 1.1 | 1.297 | 1.037 | 0.336 | 0.486 | 1.081 |
| 0.55 | 2.151 | 2.550 | 1.776 | 2.425 | 1.492 |
| 0.275 | 2.953 | 3.010 | 3.924 | 3.896 | 3.983 |
| 0.1375 | 2.915 | 5.247 | 7.558 | 7.925 | 8.492 |
| 0.06875 | 3.815 | 6.660 | 10.381 | 14.320 | 16.070 |
| 0.034375 | 4.865 | 7.837 | 14.069 | 22.268 | 28.488 |
| 0.0171875 | 3.459 | 8.036 | 15.820 | 27.483 | 43.672 |
| 0.00859375 | 4.111 | 8.825 | 16.933 | 30.905 | 55.616 |
| 0.00429688 | 4.398 | 8.784 | 17.288 | 33.359 | 63.176 |
| 0.00214844 | 4.150 | 8.703 | 17.578 | 34.462 | 66.845 |
| 0 | 4.718 | 8.980 | 18.156 | 36.339 | 71.711 |

*Cpd.* **9***: Determination of* $IC_{50}$ *and* $k_1^*$

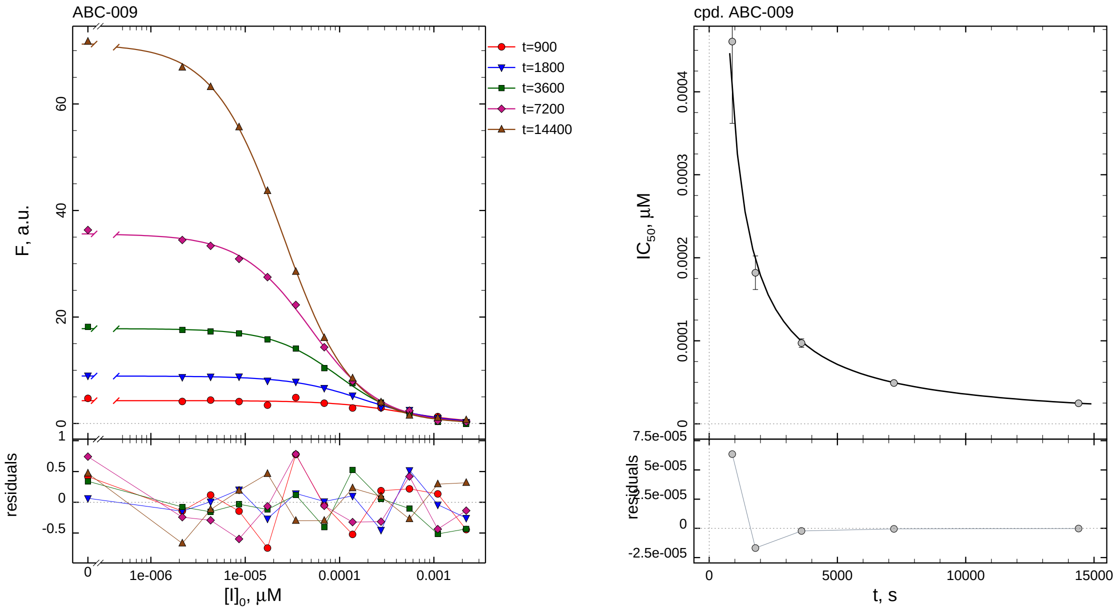

| $i$ | $t$, min | $I_{50}$, nM | ± | std.err. | CV, % | $k_1^*$, mM$^{-1}$s$^{-1}$ |
|---|---|---|---|---|---|---|
| 1 | 15 | 0.4605 | ± | 0.09849 | 21.4 | 3845.4 |
| 2 | 30 | 0.1820 | ± | 0.02020 | 11.1 | 4864.2 |
| 3 | 60 | 0.09727 | ± | 0.005121 | 5.3 | 4551.0 |
| 4 | 120 | 0.04927 | ± | 0.001440 | 2.9 | 4492.7 |
| 5 | 240 | 0.02477 | ± | 0.0003791 | 1.5 | 4468.5 |

*Cpd.* **9***: Mechanistic analysis*

| parameter | unit | "true" (T) | calculated (C) | C/T ratio | note |
|---|---|---|---|---|---|
| mechanism | | **C2S** | **C1** | | |
| $k_{\mathrm{eff}}$ | mM$^{-1}$s$^{-1}$ | 9090.9 | 8985.4 | 0.99 | $= k_1^*\ (1 + [\mathrm{S}]_0/K_\mathrm{M})$ |

| | |
|---|---|
| Reaction times used for analysis: | 30 and 120 min |
| Maximum GSD for accepting one-step model **C1**: | 1.25 |
| Observed GSD: | 1.06 |
| Assumed $[\mathrm{S}]_0/K_\mathrm{M}$ ratio: | 1.0 |

*2.10. Compound No. 10*

*Cpd.* **10***: Simulated data*

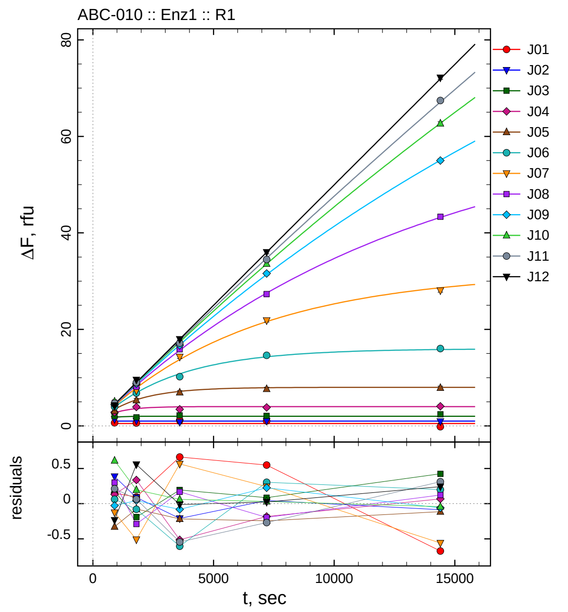

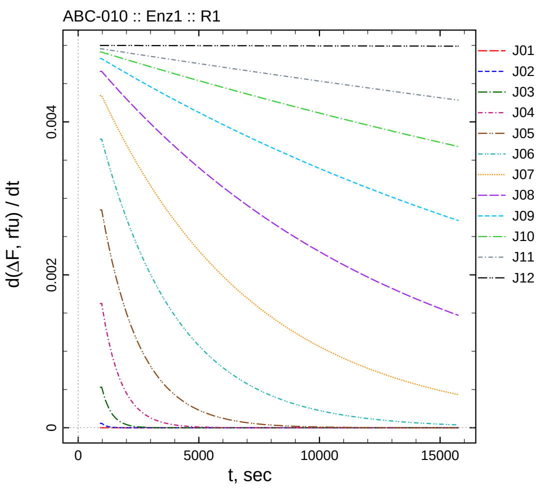

| $[I]_0$, nM | t = 15 | 30 | 60 | 120 | 240 min |
|---:|---:|---:|---:|---:|---:|
| 22 | 0.649 | 0.595 | 1.162 | 1.049 | -0.173 |
| 11 | 1.374 | 1.089 | 0.790 | 1.044 | 0.914 |
| 5.5 | 1.961 | 1.786 | 2.195 | 2.085 | 2.425 |
| 2.75 | 2.830 | 3.914 | 3.442 | 3.817 | 4.068 |
| 1.375 | 3.112 | 5.324 | 6.939 | 7.667 | 7.886 |
| 0.6875 | 3.982 | 6.801 | 10.199 | 14.618 | 16.026 |
| 0.34375 | 4.068 | 7.333 | 14.328 | 21.850 | 28.074 |
| 0.171875 | 4.642 | 8.104 | 15.854 | 27.331 | 43.342 |
| 0.0859375 | 4.394 | 8.731 | 16.706 | 31.595 | 54.986 |
| 0.0429688 | 5.072 | 9.037 | 17.437 | 33.601 | 62.679 |
| 0.0214844 | 4.691 | 8.983 | 17.138 | 34.479 | 67.427 |
| 0 | 4.264 | 9.555 | 17.982 | 36.007 | 72.176 |

*Cpd.* **10***: Determination of $IC_{50}$ and $k_1^*$*

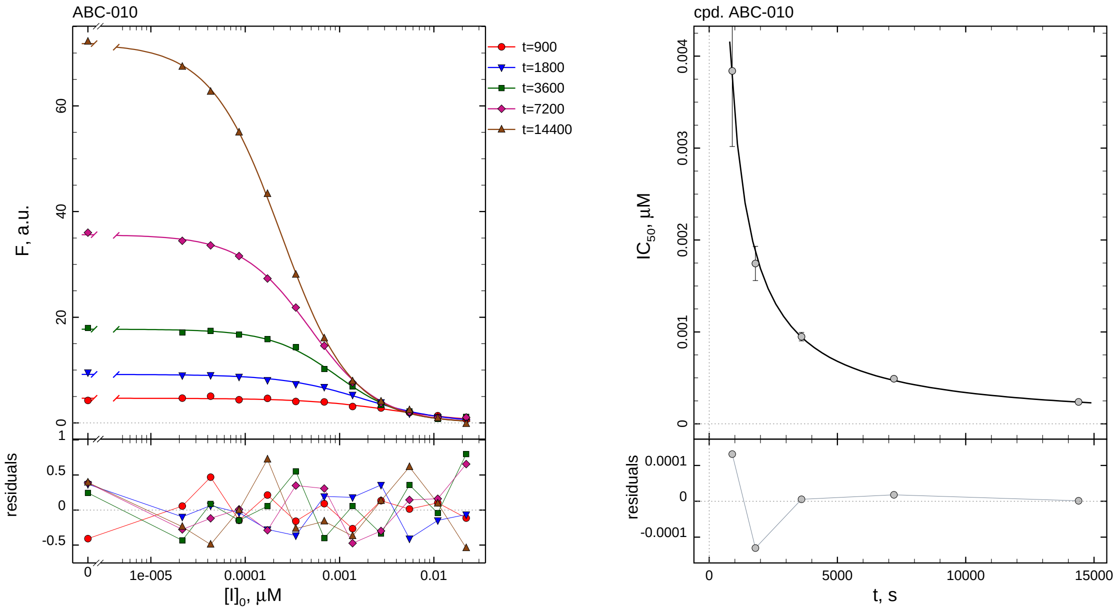

| $i$ | $t$, min | $I_{50}$, nM | ± | std.err. | CV, % | $k_1^*$, mM$^{-1}$s$^{-1}$ |
|---|---|---|---|---|---|---|
| 1 | 15 | 3.839 | ± | 0.8218 | 21.4 | 461.2 |
| 2 | 30 | 1.745 | ± | 0.1864 | 10.7 | 507.4 |
| 3 | 60 | 0.9485 | ± | 0.04613 | 4.9 | 466.7 |
| 4 | 120 | 0.4908 | ± | 0.01298 | 2.6 | 451.0 |
| 5 | 240 | 0.2381 | ± | 0.003370 | 1.4 | 464.7 |

*Cpd.* **10***: Mechanistic analysis*

| parameter | unit | "true" (T) | calculated (C) | C/T ratio | note |
|---|---|---|---|---|---|
| mechanism | | **C2S** | **C1** | | |
| $k_{\mathrm{eff}}$ | mM$^{-1}$s$^{-1}$ | 909.1 | 901.9 | 0.99 | $= k_1^* \, (1 + [S]_0/K_M)$ |

| | |
|---|---|
| Reaction times used for analysis: | 30 and 120 min |
| Maximum GSD for accepting one-step model **C1**: | 1.25 |
| Observed GSD: | 1.09 |
| Assumed $[S]_0/K_M$ ratio: | 1.0 |

*2.11. Compound No. 11*

*Cpd.* **11***: Simulated data*

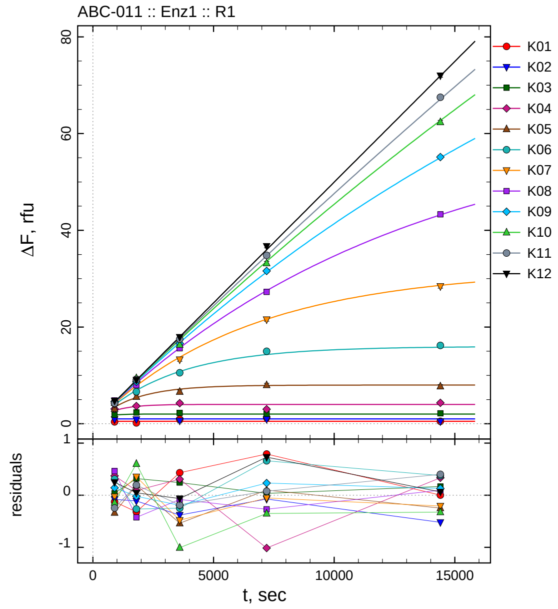

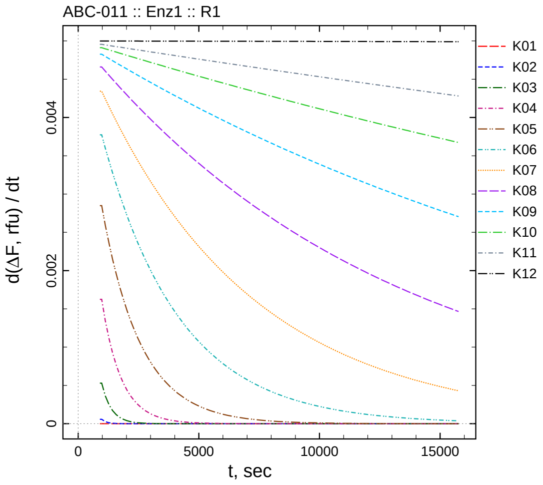

| $[I]_0$, nM | t = 15 | 30 | 60 | 120 | 240 min |
|---:|---:|---:|---:|---:|---:|
| 220 | 0.375 | 0.180 | 0.932 | 1.290 | 0.502 |
| 110 | 0.909 | 0.889 | 0.618 | 0.927 | 0.480 |
| 55 | 1.826 | 2.297 | 2.242 | 2.034 | 2.167 |
| 27.5 | 3.072 | 3.668 | 4.264 | 2.986 | 4.334 |
| 13.75 | 3.104 | 5.593 | 6.616 | 8.009 | 7.750 |
| 6.875 | 4.231 | 6.616 | 10.552 | 14.970 | 16.195 |
| 3.4375 | 4.169 | 8.198 | 13.281 | 21.548 | 28.404 |
| 1.71875 | 4.806 | 7.968 | 15.602 | 27.250 | 43.288 |
| 0.859375 | 4.562 | 8.668 | 16.591 | 31.598 | 55.154 |
| 0.429688 | 4.317 | 9.444 | 16.368 | 33.216 | 62.382 |
| 0.214844 | 4.234 | 9.115 | 17.492 | 34.824 | 67.509 |
| 0 | 4.748 | 9.043 | 17.930 | 36.715 | 71.998 |

*Cpd.* **11***: Determination of* $IC_{50}$ *and* $k_1^*$

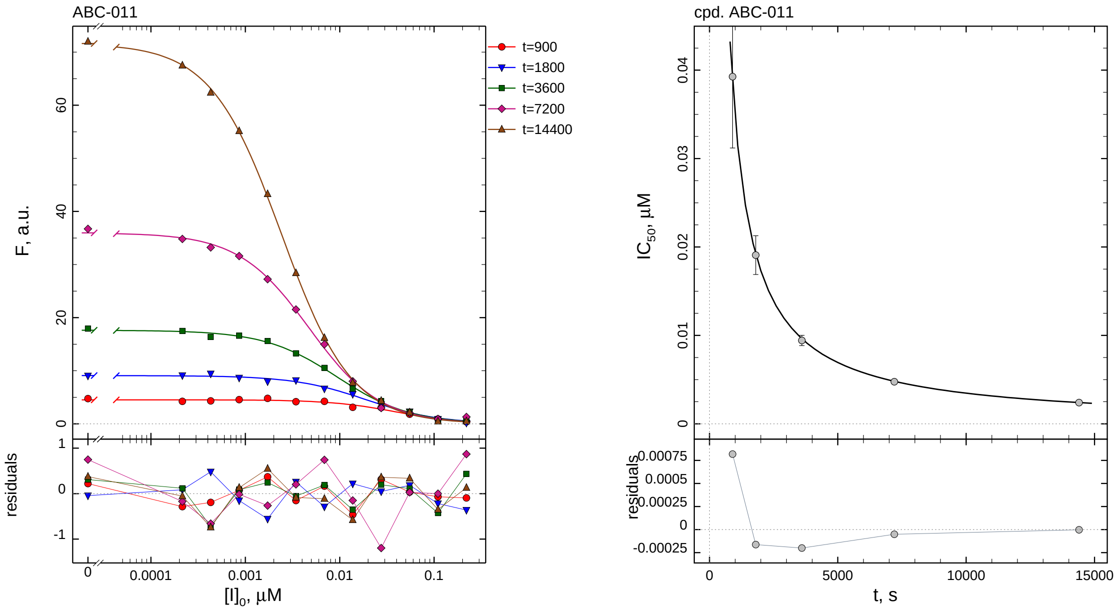

| $i$ | $t$, min | $I_{50}$, nM | ± | std.err. | CV, % | $k_1^*$, mM$^{-1}$s$^{-1}$ |
|---|---|---|---|---|---|---|
| 1 | 15 | 39.26 | ± | 8.065 | 20.5 | 45.10 |
| 2 | 30 | 19.09 | ± | 2.190 | 11.5 | 46.39 |
| 3 | 60 | 9.431 | ± | 0.5876 | 6.2 | 46.94 |
| 4 | 120 | 4.766 | ± | 0.1501 | 3.2 | 46.44 |
| 5 | 240 | 2.407 | ± | 0.04017 | 1.7 | 45.98 |

*Cpd.* **11***: Mechanistic analysis*

| parameter | unit | "true" (T) | calculated (C) | C/T ratio | note |
|---|---|---|---|---|---|
| mechanism | | **C2S** | **C1** | | |
| $k_{\mathrm{eff}}$ | mM$^{-1}$s$^{-1}$ | 90.91 | 92.88 | 1.02 | $= k_1^* \, (1 + [\mathrm{S}]_0/K_\mathrm{M})$ |

| | |
|---|---|
| Reaction times used for analysis: | 30 and 120 min |
| Maximum GSD for accepting one-step model **C1**: | 1.25 |
| Observed GSD: | 1.00 |
| Assumed $[\mathrm{S}]_0/K_\mathrm{M}$ ratio: | 1.0 |

*2.12. Compound No. 12*

*Cpd.* **12***: Simulated data*

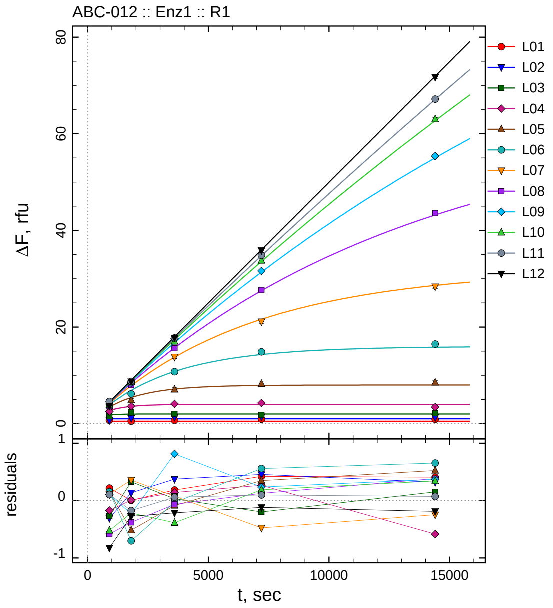

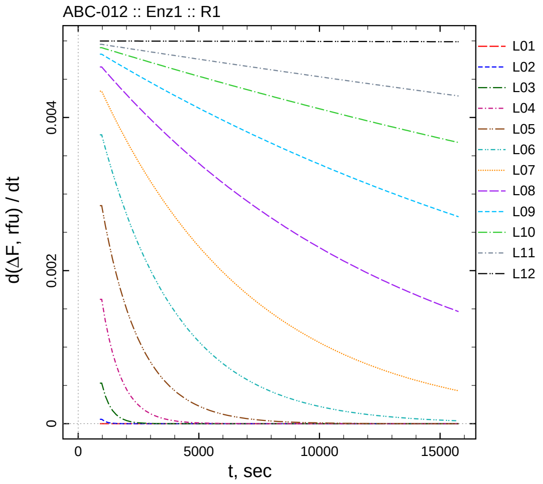

| $[I]_0$, nM | t = 15 | 30 | 60 | 120 | 240 min |
|---|---|---|---|---|---|
| 2200 | 0.717 | 0.503 | 0.683 | 0.922 | 0.908 |
| 1100 | 0.681 | 1.136 | 1.372 | 1.456 | 1.320 |
| 550 | 1.519 | 2.305 | 2.030 | 1.802 | 2.152 |
| 275 | 2.525 | 3.584 | 4.095 | 4.265 | 3.414 |
| 137.5 | 3.589 | 4.882 | 7.068 | 8.256 | 8.520 |
| 68.75 | 4.074 | 6.177 | 10.768 | 14.866 | 16.468 |
| 34.375 | 4.310 | 8.200 | 13.837 | 21.123 | 28.362 |
| 17.1875 | 3.760 | 8.012 | 15.617 | 27.645 | 43.572 |
| 8.59375 | 4.524 | 8.459 | 17.599 | 31.601 | 55.390 |
| 4.29688 | 3.937 | 8.624 | 16.990 | 33.754 | 63.047 |
| 2.14844 | 4.585 | 8.745 | 17.739 | 34.843 | 67.177 |
| 0 | 3.674 | 8.722 | 17.779 | 35.864 | 71.746 |

*Cpd.* **12***: Determination of $IC_{50}$ and $k_1^*$*

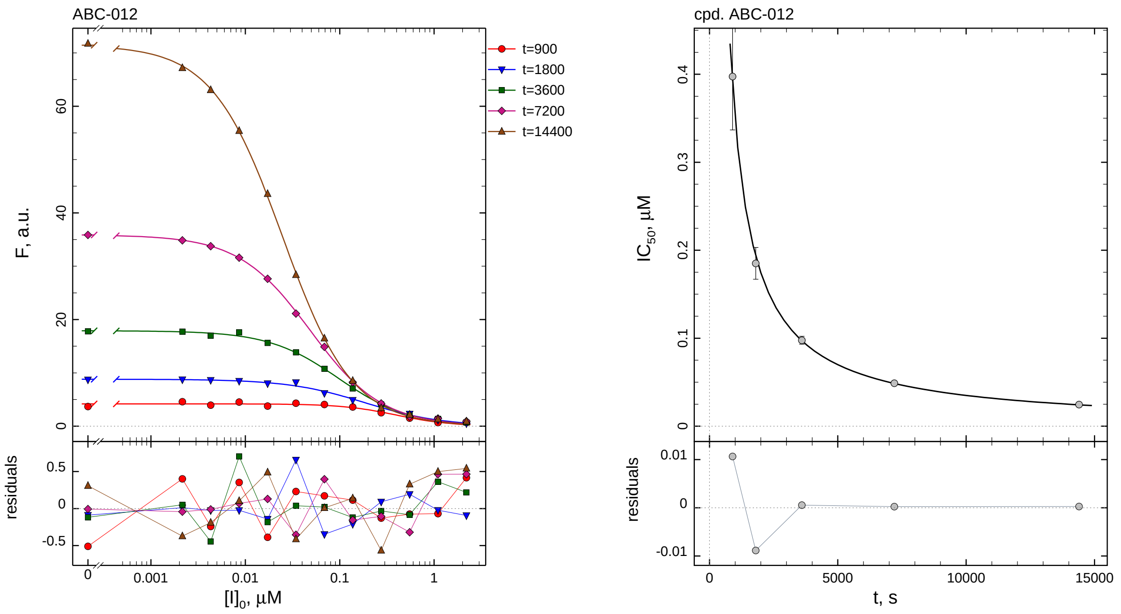

| $i$ | $t$, min | $I_{50}$, nM | ± | std.err. | CV, % | $k_1^*$, mM$^{-1}$s$^{-1}$ |
|---|---|---|---|---|---|---|
| 1 | 15 | 397.4 | ± | 60.65 | 15.3 | 4.455 |
| 2 | 30 | 185.1 | ± | 18.06 | 9.8 | 4.784 |
| 3 | 60 | 97.63 | ± | 4.519 | 4.6 | 4.534 |
| 4 | 120 | 48.80 | ± | 1.214 | 2.5 | 4.535 |
| 5 | 240 | 24.56 | ± | 0.3180 | 1.3 | 4.506 |

*Cpd.* **12***: Mechanistic analysis*

| parameter | unit | "true" (T) | calculated (C) | C/T ratio | note |
|---|---|---|---|---|---|
| mechanism | | **C2S** | **C1** | | |
| $k_{\mathrm{eff}}$ | mM$^{-1}$s$^{-1}$ | 9.091 | 9.070 | 1.00 | $= k_1^*\ (1 + [\mathrm{S}]_0/K_\mathrm{M})$ |

| | |
|---|---|
| Reaction times used for analysis: | 30 and 120 min |
| Maximum GSD for accepting one-step model **C1**: | 1.25 |
| Observed GSD: | 1.04 |
| Assumed $[\mathrm{S}]_0/K_\mathrm{M}$ ratio: | 1.0 |

#### *2.13. Compound No. 13*

*Cpd.* **13***: Simulated data*

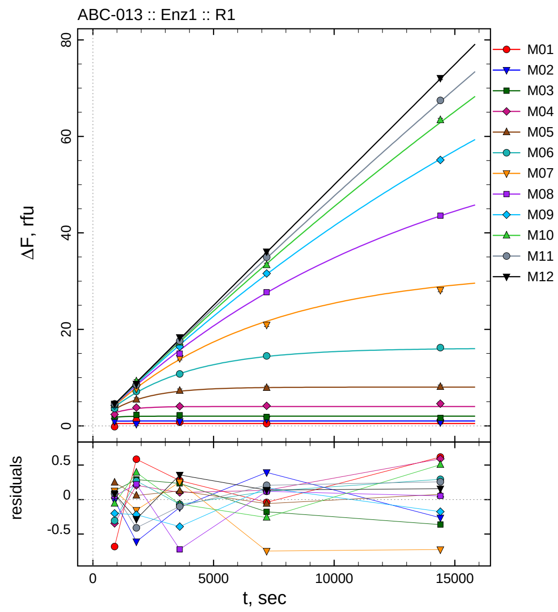

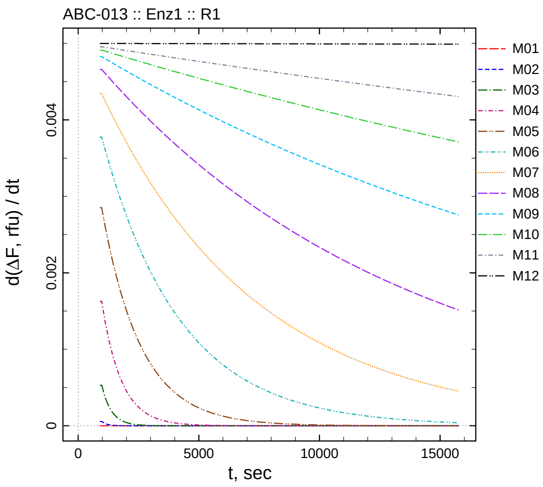

| $[I]_0$, nM | t = 15 | 30 | 60 | 120 | 240 min |
|---:|---:|---:|---:|---:|---:|
| 2.02 | -0.182 | 1.086 | 0.776 | 0.459 | 1.117 |
| 1.01 | 0.976 | 0.390 | 0.887 | 1.394 | 0.737 |
| 0.505 | 1.917 | 2.270 | 2.236 | 1.825 | 1.638 |
| 0.2525 | 2.357 | 3.789 | 4.065 | 4.141 | 4.598 |
| 0.12625 | 3.689 | 5.466 | 7.298 | 7.879 | 8.106 |
| 0.063125 | 3.616 | 7.158 | 10.755 | 14.493 | 16.226 |
| 0.0315625 | 4.325 | 7.692 | 14.034 | 20.940 | 28.148 |
| 0.0157813 | 4.398 | 8.617 | 14.975 | 27.710 | 43.574 |
| 0.00789063 | 4.221 | 8.470 | 16.402 | 31.580 | 55.124 |
| 0.00394531 | 4.403 | 9.234 | 17.316 | 33.334 | 63.392 |
| 0.00197266 | 4.566 | 8.510 | 17.586 | 34.969 | 67.468 |
| 0 | 4.581 | 8.714 | 18.352 | 36.121 | 72.092 |

*Cpd.* **13***: Determination of* $IC_{50}$ *and* $k_1^*$

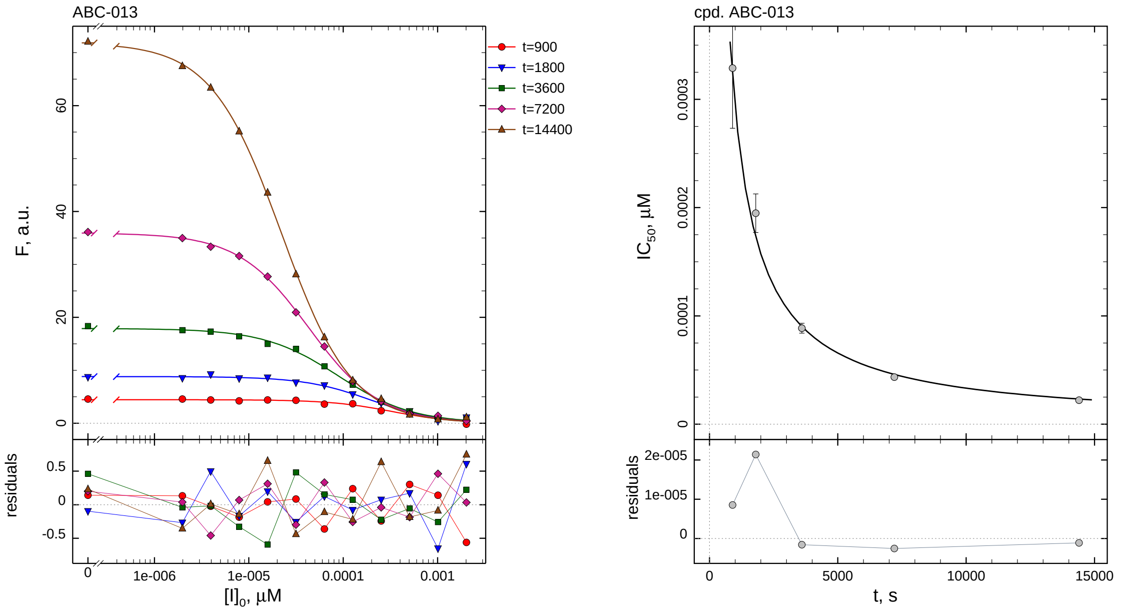

| $i$ | $t$, min | $I_{50}$, nM | ± | std.err. | CV, % | $k_1^*$, mM$^{-1}$s$^{-1}$ |
|---|---|---|---|---|---|---|
| 1 | 15 | 0.3289 | ± | 0.05555 | 16.9 | 5384.1 |
| 2 | 30 | 0.1948 | ± | 0.01783 | 9.1 | 4544.3 |
| 3 | 60 | 0.08866 | ± | 0.004586 | 5.2 | 4993.1 |
| 4 | 120 | 0.04348 | ± | 0.001143 | 2.6 | 5090.2 |
| 5 | 240 | 0.02212 | ± | 0.0003045 | 1.4 | 5002.6 |

*Cpd.* **13***: Mechanistic analysis*

| parameter | unit | "true" (T) | calculated (C) | C/T ratio | note |
|---|---|---|---|---|---|
| mechanism | | **C2S** | **C1** | | |
| $k_{\text{eff}}$ | mM$^{-1}$s$^{-1}$ | 9901.0 | 10180 | 1.03 | $= k_1^* \, (1 + [S]_0/K_M)$ |

| | |
|---|---|
| Reaction times used for analysis: | 30 and 120 min |
| Maximum GSD for accepting one-step model **C1**: | 1.25 |
| Observed GSD: | 1.08 |
| Assumed $[S]_0/K_M$ ratio: | 1.0 |

#### 2.14. *Compound No. 14*

*Cpd.* **14***: Simulated data*

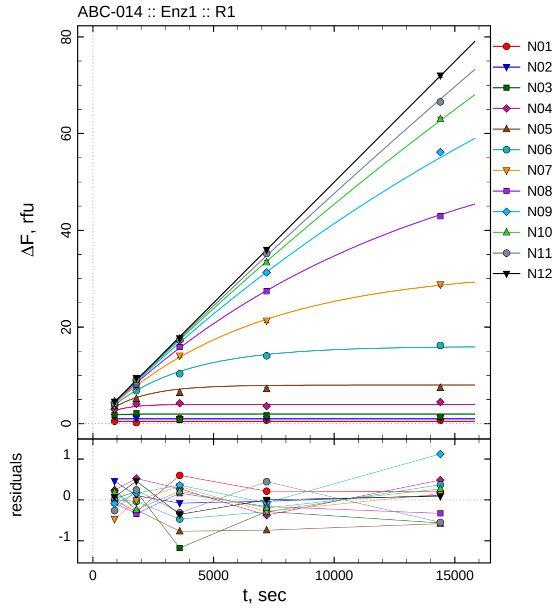

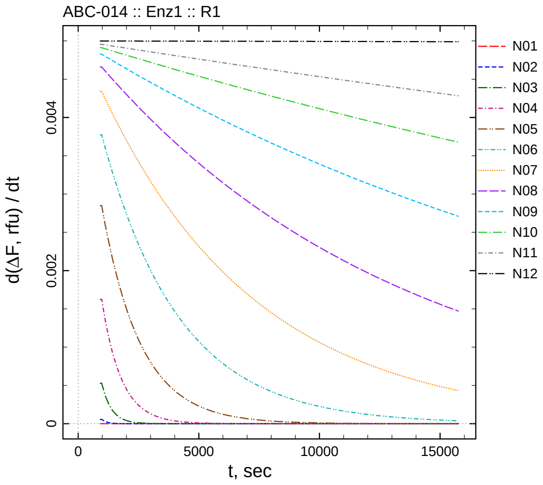

| $[I]_0$, nM | t = 15 | 30 | 60 | 120 | 240 min |
|---|---|---|---|---|---|
| 20.2 | 0.501 | 0.208 | 1.101 | 0.713 | 0.698 |
| 10.1 | 1.449 | 1.108 | 0.923 | 0.966 | 1.110 |
| 5.05 | 1.820 | 2.181 | 0.825 | 1.717 | 1.438 |
| 2.525 | 2.941 | 4.105 | 4.229 | 3.626 | 4.487 |
| 1.2625 | 3.694 | 5.159 | 6.392 | 7.172 | 7.420 |
| 0.63125 | 3.927 | 6.843 | 10.336 | 14.026 | 16.198 |
| 0.315625 | 3.729 | 7.830 | 14.091 | 21.327 | 28.787 |
| 0.157813 | 4.270 | 8.055 | 15.852 | 27.365 | 42.896 |
| 0.0789063 | 4.331 | 8.853 | 17.148 | 31.299 | 56.172 |
| 0.0394531 | 4.644 | 8.606 | 17.592 | 33.374 | 62.994 |
| 0.0197266 | 4.220 | 9.164 | 17.367 | 35.196 | 66.568 |
| 0 | 4.565 | 9.460 | 17.641 | 35.985 | 72.026 |

*Cpd.* **14***: Determination of* $IC_{50}$ *and* $k_1^*$

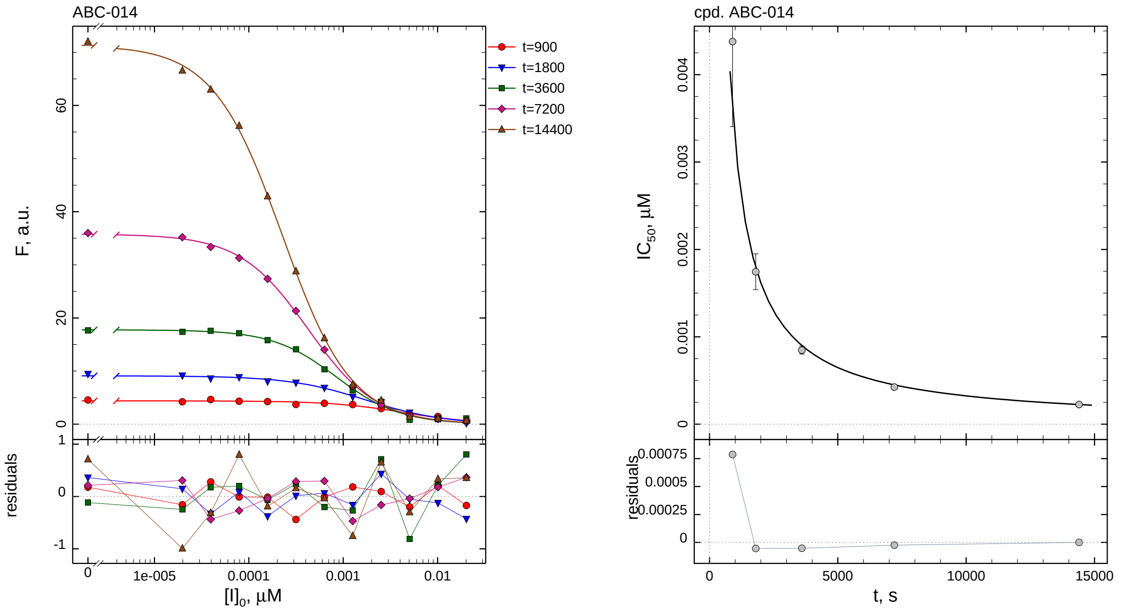

| $i$ | $t$, min | $I_{50}$, nM | ± | std.err. | CV, % | $k_1^*$, mM$^{-1}$s$^{-1}$ |
|---|---|---|---|---|---|---|
| 1 | 15 | 4.379 | ± | 0.9724 | 22.2 | 404.4 |
| 2 | 30 | 1.745 | ± | 0.2050 | 11.8 | 507.5 |
| 3 | 60 | 0.8479 | ± | 0.04539 | 5.4 | 522.1 |
| 4 | 120 | 0.4251 | ± | 0.01268 | 3.0 | 520.7 |
| 5 | 240 | 0.2254 | ± | 0.003592 | 1.6 | 491.0 |

*Cpd.* **14***: Mechanistic analysis*

| parameter | unit | "true" (T) | calculated (C) | C/T ratio | note |
|---|---|---|---|---|---|
| mechanism | | **C2S** | **C1** | | |
| $k_{\mathrm{eff}}$ | mM$^{-1}$s$^{-1}$ | 990.1 | 1041.4 | 1.05 | $= k_1^* \, (1 + [\mathrm{S}]_0/K_\mathrm{M})$ |

| | |
|---|---|
| Reaction times used for analysis: | 30 and 120 min |
| Maximum GSD for accepting one-step model **C1**: | 1.25 |
| Observed GSD: | 1.02 |
| Assumed $[\mathrm{S}]_0/K_\mathrm{M}$ ratio: | 1.0 |

#### 2.15. *Compound No. 15*

*Cpd.* **15***: Simulated data*

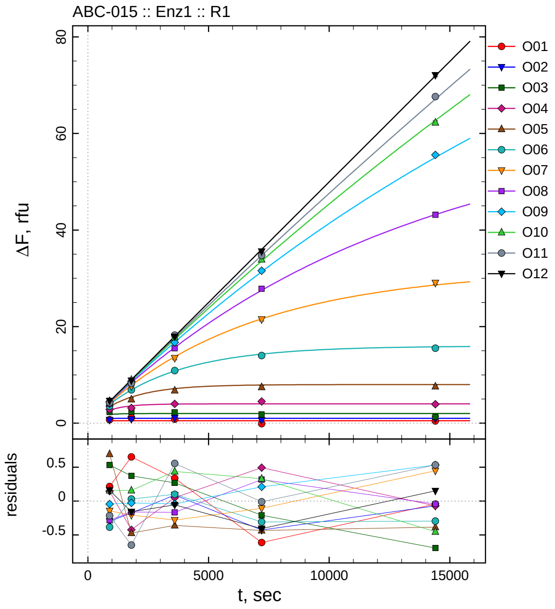

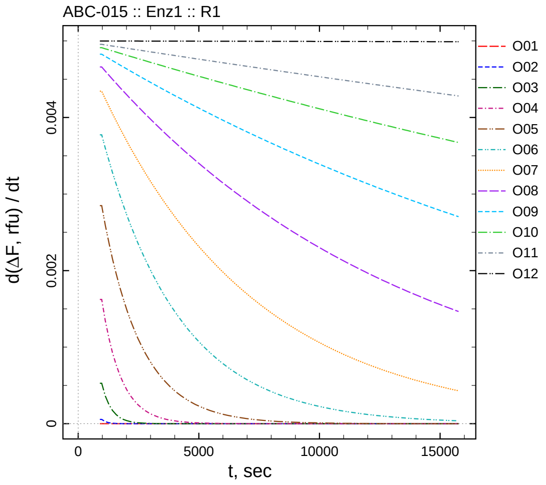

| $[I]_0$, nM | t = 15 | 30 | 60 | 120 | 240 min |
|---:|---:|---:|---:|---:|---:|
| 202 | 0.715 | 1.152 | 0.837 | -0.112 | 0.455 |
| 101 | 0.688 | 0.825 | 1.089 | 0.572 | 0.927 |
| 50.5 | 2.320 | 2.349 | 2.269 | 1.791 | 1.307 |
| 25.25 | 2.848 | 3.153 | 4.005 | 4.492 | 3.927 |
| 12.625 | 4.134 | 4.929 | 6.796 | 7.475 | 7.612 |
| 6.3125 | 3.536 | 6.912 | 10.902 | 14.001 | 15.521 |
| 3.15625 | 4.052 | 7.636 | 13.482 | 21.496 | 29.059 |
| 1.57813 | 4.061 | 8.232 | 15.519 | 27.838 | 43.149 |
| 0.789063 | 4.376 | 8.661 | 16.744 | 31.575 | 55.556 |
| 0.394531 | 4.615 | 8.999 | 17.813 | 33.897 | 62.254 |
| 0.197266 | 4.263 | 8.270 | 18.237 | 34.736 | 67.641 |
| 0 | 4.657 | 8.839 | 17.938 | 35.575 | 72.085 |

*Cpd.* **15***: Determination of* $IC_{50}$ *and* $k_1^*$

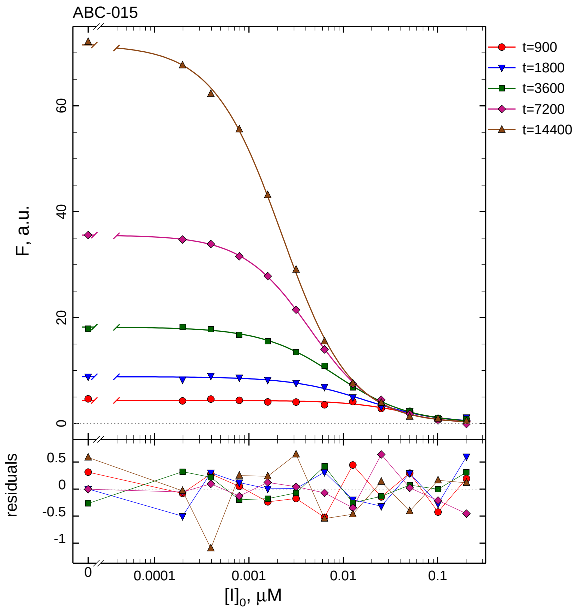

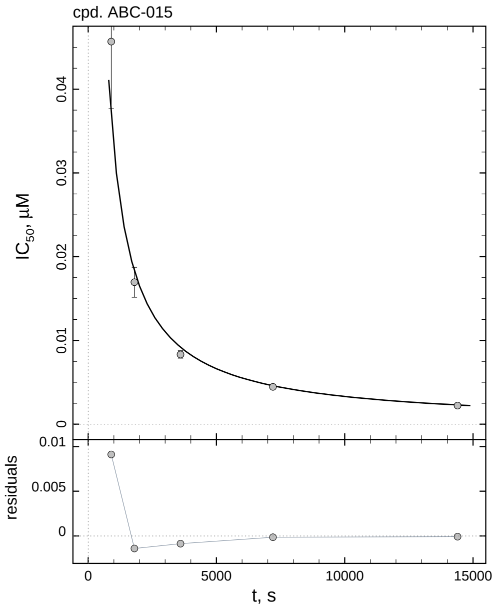

| $i$ | $t$, min | $I_{50}$, nM | ± | std.err. | CV, % | $k_1^*$, mM$^{-1}$s$^{-1}$ |
|---|---|---|---|---|---|---|
| 1 | 15 | 45.68 | ± | 8.017 | 17.5 | 38.76 |
| 2 | 30 | 16.95 | ± | 1.787 | 10.5 | 52.23 |
| 3 | 60 | 8.326 | ± | 0.4456 | 5.4 | 53.16 |
| 4 | 120 | 4.452 | ± | 0.1176 | 2.6 | 49.72 |
| 5 | 240 | 2.222 | ± | 0.03173 | 1.4 | 49.79 |

*Cpd.* **15***: Mechanistic analysis*

| parameter | unit | "true" (T) | calculated (C) | C/T ratio | note |
|---|---|---|---|---|---|
| mechanism | | **C2S** | **C1** | | |
| $k_{\text{eff}}$ | mM$^{-1}$s$^{-1}$ | 99.01 | 99.44 | 1.00 | $= k_1^* \, (1 + [\text{S}]_0/K_\text{M})$ |

| | |
|---|---|
| Reaction times used for analysis: | 30 and 120 min |
| Maximum GSD for accepting one-step model **C1**: | 1.25 |
| Observed GSD: | 1.04 |
| Assumed $[\text{S}]_0/K_\text{M}$ ratio: | 1.0 |

*2.16. Compound No. 16*

*Cpd.* **16***: Simulated data*

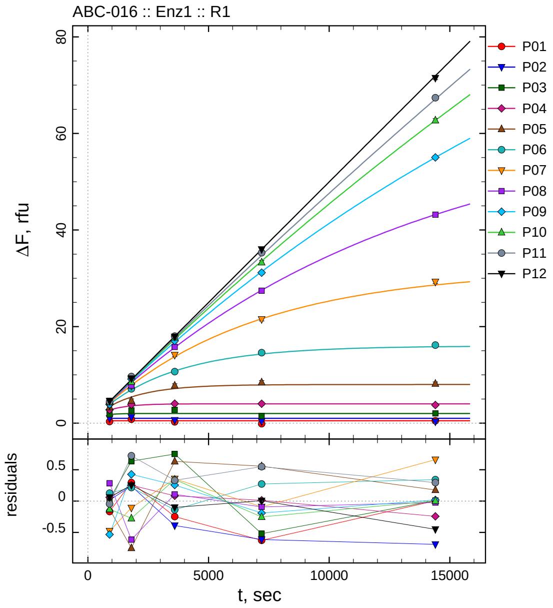

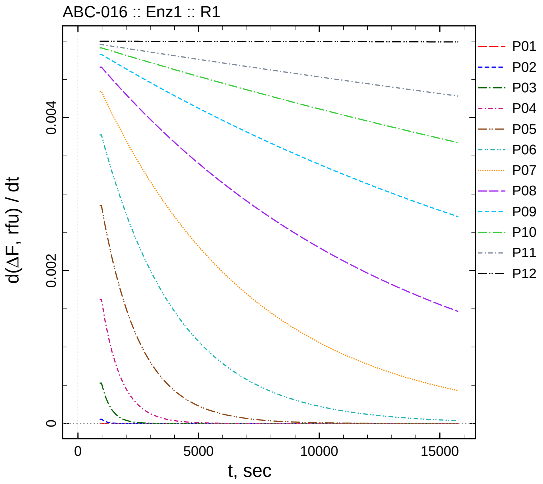

| $[I]_0$, nM | t = 15 | 30 | 60 | 120 | 240 min |
|---|---|---|---|---|---|
| 2020 | 0.333 | 0.797 | 0.252 | -0.126 | 0.505 |
| 1010 | 1.005 | 1.240 | 0.610 | 0.386 | 0.309 |
| 505 | 1.820 | 2.616 | 2.750 | 1.479 | 2.010 |
| 252.5 | 2.760 | 3.831 | 4.039 | 4.011 | 3.757 |
| 126.25 | 3.309 | 4.652 | 7.789 | 8.467 | 8.175 |
| 63.125 | 4.047 | 7.098 | 10.663 | 14.582 | 16.158 |
| 31.5625 | 3.717 | 7.734 | 14.118 | 21.510 | 29.271 |
| 15.7813 | 4.629 | 7.778 | 15.793 | 27.427 | 43.166 |
| 7.89063 | 3.887 | 9.115 | 17.043 | 31.176 | 55.039 |
| 3.94531 | 4.331 | 8.566 | 17.721 | 33.314 | 62.709 |
| 1.97266 | 4.434 | 9.643 | 18.013 | 35.296 | 67.401 |
| 0 | 4.560 | 9.250 | 17.894 | 35.989 | 71.482 |

*Cpd.* **16**: *Determination of $IC_{50}$ and $k_1^*$*

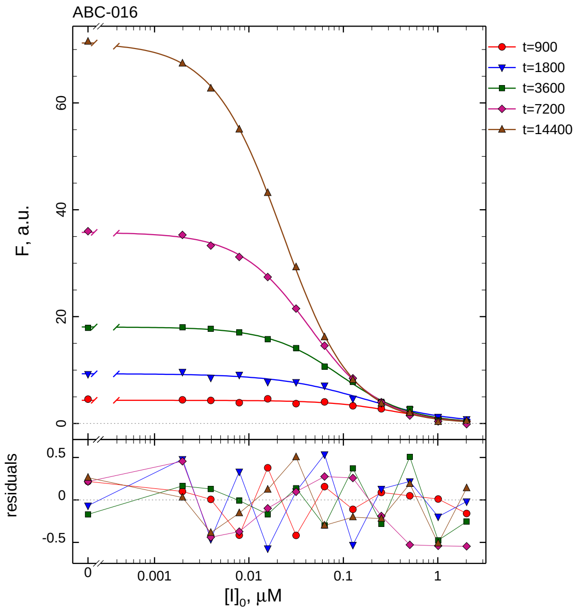

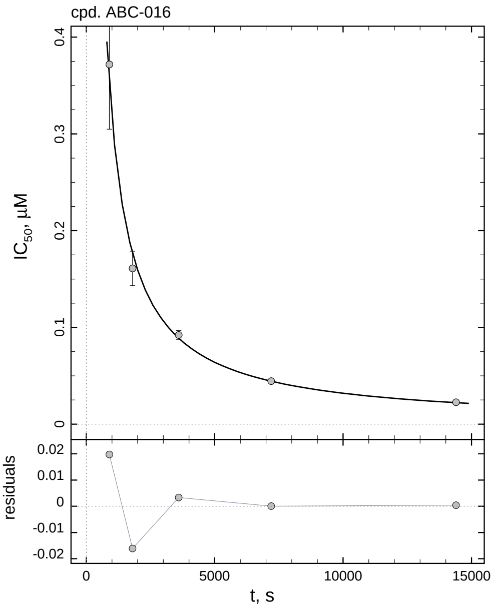

| $i$ | $t$, min | $I_{50}$, nM | ± | std.err. | CV, % | $k_1^*$, $mM^{-1}s^{-1}$ |
|---|---|---|---|---|---|---|
| 1 | 15 | 371.7 | ± | 66.81 | 18.0 | 4.763 |
| 2 | 30 | 161.0 | ± | 17.78 | 11.0 | 5.498 |
| 3 | 60 | 92.16 | ± | 4.499 | 4.9 | 4.803 |
| 4 | 120 | 44.52 | ± | 1.153 | 2.6 | 4.972 |
| 5 | 240 | 22.65 | ± | 0.3099 | 1.4 | 4.886 |

*Cpd.* **16**: *Mechanistic analysis*

| parameter | unit | “true” (T) | calculated (C) | C/T ratio | note |
|---|---|---|---|---|---|
| mechanism | | **C2S** | **C1** | | |
| $k_{eff}$ | $mM^{-1}s^{-1}$ | 9.901 | 9.943 | 1.00 | $= k_1^* \ (1 + [S]_0/K_M)$ |

| | |
|---|---|
| Reaction times used for analysis: | 30 and 120 min |
| Maximum GSD for accepting one-step model **C1**: | 1.25 |
| Observed GSD: | 1.07 |
| Assumed $[S]_0/K_M$ ratio: | 1.0 |

*2.17. Compound No. 17*

*Cpd.* **17***: Simulated data*

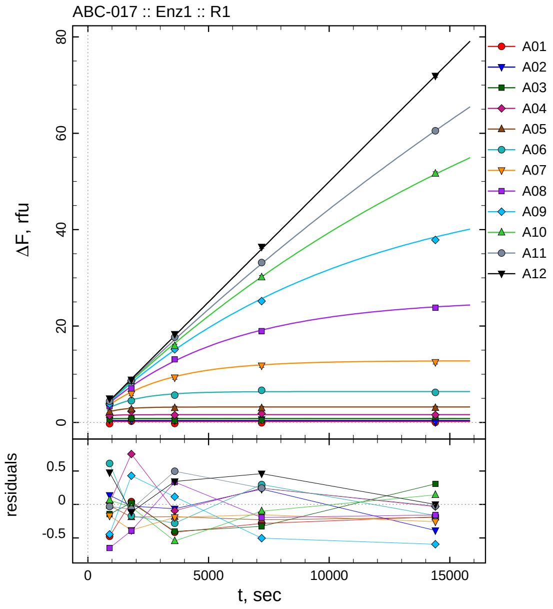

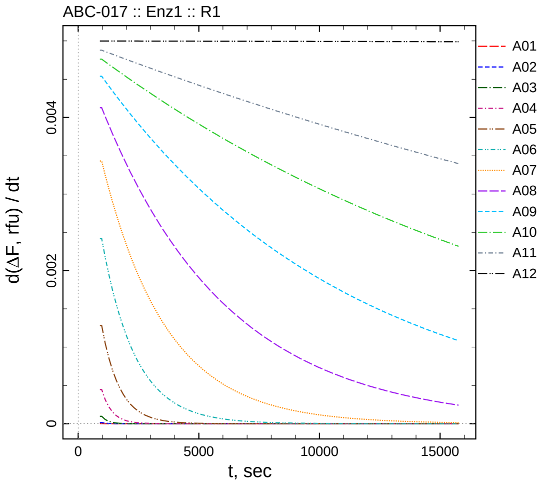

| $[I]_0$, nM | t = 15 | 30 | 60 | 120 | 240 min |
|---:|---:|---:|---:|---:|---:|
| 505 | -0.276 | 0.243 | -0.216 | -0.084 | 0.015 |
| 252.5 | 0.535 | 0.376 | 0.335 | 0.631 | 0.012 |
| 126.25 | 0.629 | 0.824 | 0.398 | 0.473 | 1.108 |
| 63.125 | 1.437 | 2.331 | 1.506 | 1.843 | 1.567 |
| 31.5625 | 2.236 | 2.730 | 2.991 | 2.969 | 3.007 |
| 15.7813 | 3.680 | 4.492 | 5.647 | 6.661 | 6.233 |
| 7.89063 | 3.505 | 5.908 | 9.300 | 11.790 | 12.483 |
| 3.94531 | 3.403 | 7.073 | 13.088 | 18.955 | 23.810 |
| 1.97266 | 3.819 | 8.608 | 15.166 | 25.168 | 37.864 |
| 0.986328 | 4.444 | 8.536 | 15.883 | 30.116 | 51.641 |
| 0.493164 | 4.408 | 8.711 | 17.683 | 33.169 | 60.509 |
| 0 | 4.976 | 8.884 | 18.339 | 36.443 | 71.934 |

*Cpd.* **17***: Determination of* $IC_{50}$ *and* $k_1^*$

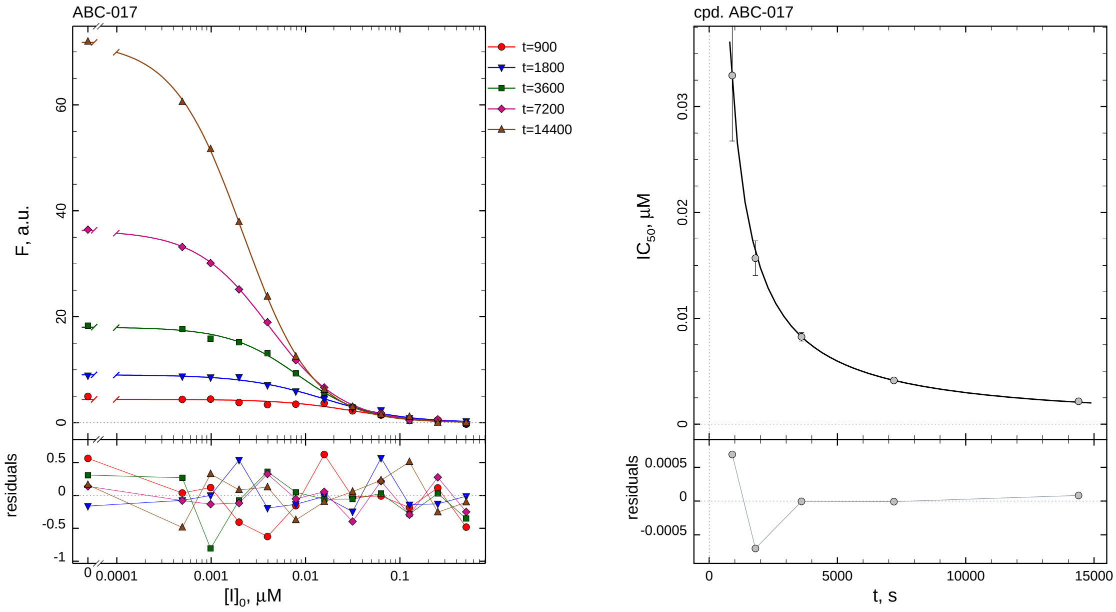

| $i$ | $t$, min | $I_{50}$, nM ± std.err. | CV, % | $k_1^*$, $mM^{-1}s^{-1}$ |
|---|---|---|---|---|
| 1 | 15 | 32.94 ± 6.187 | 18.8 | 53.75 |
| 2 | 30 | 15.68 ± 1.645 | 10.5 | 56.47 |
| 3 | 60 | 8.248 ± 0.3986 | 4.8 | 53.67 |
| 4 | 120 | 4.134 ± 0.1098 | 2.7 | 53.54 |
| 5 | 240 | 2.158 ± 0.02852 | 1.3 | 51.27 |

*Cpd.* **17***: Mechanistic analysis*

| parameter | unit | "true" (T) | calculated (C) | C/T ratio | note |
|---|---|---|---|---|---|
| mechanism | | **C2S** | **C1** | | |
| $k_{\text{eff}}$ | $mM^{-1}s^{-1}$ | 99.01 | 107.1 | 1.08 | $= k_1^*\ (1 + [S]_0/K_M)$ |

| | |
|---|---|
| Reaction times used for analysis: | 30 and 120 min |
| Maximum GSD for accepting one-step model **C1**: | 1.25 |
| Observed GSD: | 1.04 |
| Assumed $[S]_0/K_M$ ratio: | 1.0 |

*2.18. Compound No. 18*

*Cpd.* **18***: Simulated data*

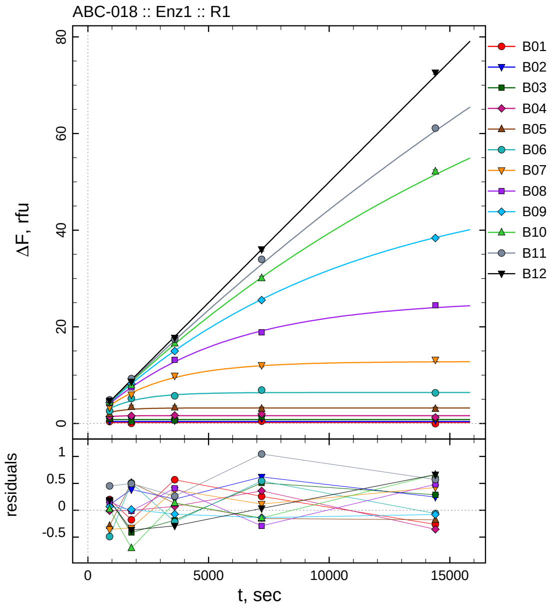

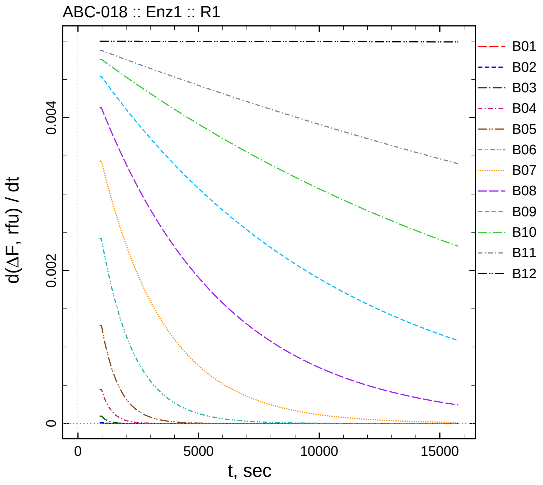

| $[I]_0$, nM | t = 15 | 30 | 60 | 120 | 240 min |
|---|---|---|---|---|---|
| 5050 | 0.401 | 0.023 | 0.768 | 0.458 | -0.063 |
| 2525 | 0.500 | 0.785 | 0.606 | 1.019 | 0.646 |
| 1262.5 | 0.962 | 0.386 | 0.613 | 1.307 | 1.089 |
| 631.25 | 1.405 | 1.576 | 1.673 | 1.960 | 1.246 |
| 315.625 | 1.965 | 3.431 | 3.304 | 3.046 | 3.022 |
| 157.812 | 2.577 | 5.171 | 5.714 | 6.909 | 6.340 |
| 78.9062 | 3.318 | 5.967 | 9.868 | 12.057 | 13.163 |
| 39.4531 | 4.211 | 7.453 | 13.164 | 18.859 | 24.449 |
| 19.7266 | 4.335 | 8.194 | 14.978 | 25.535 | 38.376 |
| 9.86328 | 4.402 | 7.873 | 16.566 | 30.074 | 52.158 |
| 4.93164 | 4.890 | 9.277 | 17.448 | 33.969 | 61.103 |
| 0 | 4.683 | 8.630 | 17.705 | 36.017 | 72.596 |

*Cpd.* **18***: Determination of* $IC_{50}$ *and* $k_1^*$

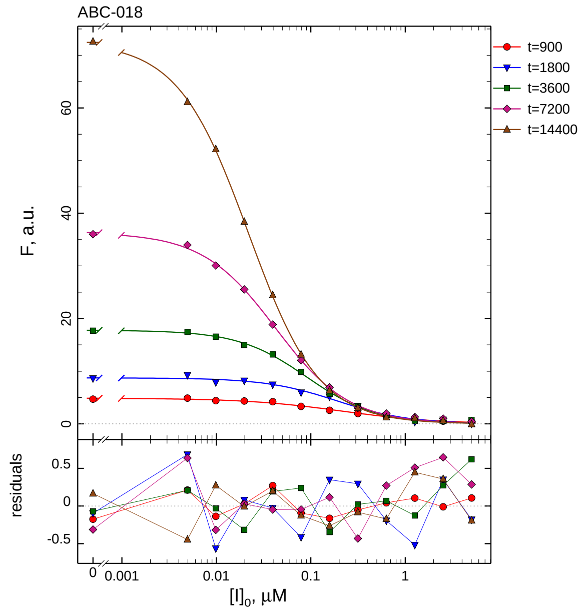

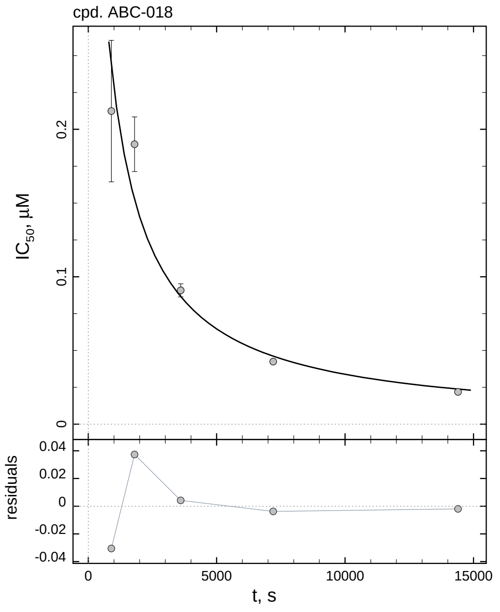

| $i$ | $t$, min | $I_{50}$, nM ± std.err. | CV, % | $k_1^*$, $mM^{-1}s^{-1}$ |
|---|---|---|---|---|
| 1 | 15 | 212.4 ± 47.99 | 22.6 | 8.337 |
| 2 | 30 | 189.9 ± 18.54 | 9.8 | 4.661 |
| 3 | 60 | 90.78 ± 4.453 | 4.9 | 4.876 |
| 4 | 120 | 42.44 ± 1.142 | 2.7 | 5.215 |
| 5 | 240 | 21.87 ± 0.2919 | 1.3 | 5.061 |

*Cpd.* **18***: Mechanistic analysis*

| parameter | unit | "true" (T) | calculated (C) | C/T ratio | note |
|---|---|---|---|---|---|
| mechanism | | **C2S** | **C1** | | |
| $k_{\mathrm{eff}}$ | $mM^{-1}s^{-1}$ | 9.901 | 10.43 | 1.05 | $= k_1^* \, (1 + [S]_0/K_M)$ |

| | |
|---|---|
| Reaction times used for analysis: | 30 and 120 min |
| Maximum GSD for accepting one-step model **C1**: | 1.25 |
| Observed GSD: | 1.08 |
| Assumed $[S]_0/K_M$ ratio: | 1.0 |

*2.19. Compound No. 19*

*Cpd.* **19***: Simulated data*

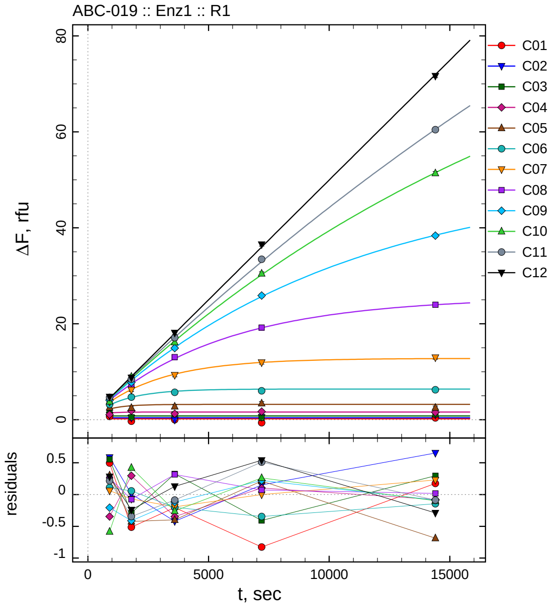

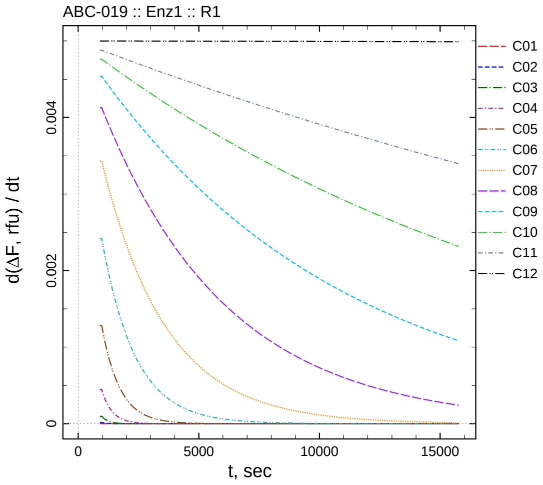

| $[I]_0$, nM | t = 15 | 30 | 60 | 120 | 240 min |
|---|---|---|---|---|---|
| 50500 | 0.697 | -0.314 | -0.004 | -0.625 | 0.374 |
| 25250 | 0.982 | 0.408 | -0.021 | 0.552 | 1.053 |
| 12625 | 1.329 | 0.509 | 1.125 | 0.391 | 1.096 |
| 6312.5 | 1.067 | 1.874 | 1.252 | 1.702 | 1.521 |
| 3156.25 | 2.556 | 2.496 | 2.777 | 3.411 | 2.513 |
| 1578.13 | 3.178 | 4.721 | 5.728 | 6.020 | 6.253 |
| 789.063 | 3.735 | 6.225 | 9.299 | 11.943 | 12.969 |
| 394.531 | 4.312 | 7.392 | 13.069 | 19.221 | 23.983 |
| 197.266 | 4.060 | 7.766 | 14.931 | 25.893 | 38.368 |
| 98.6328 | 3.796 | 8.997 | 16.170 | 30.478 | 51.397 |
| 49.3164 | 4.656 | 8.436 | 17.099 | 33.436 | 60.444 |
| 0 | 4.780 | 8.758 | 18.123 | 36.522 | 71.644 |

*Cpd.* **19***: Determination of $IC_{50}$ and $k_1^*$*

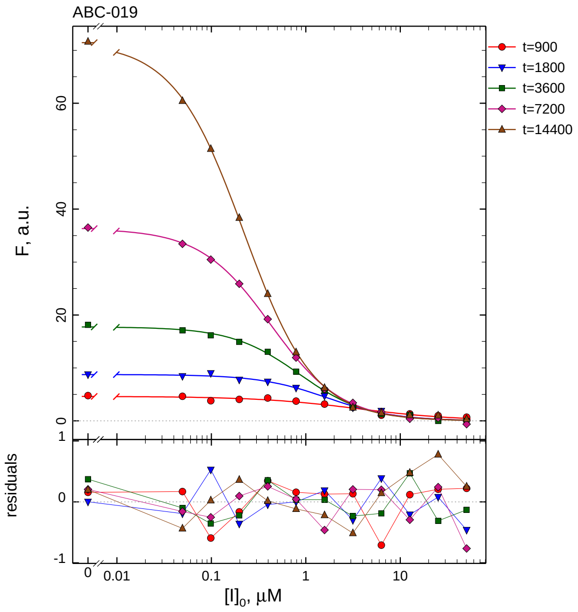

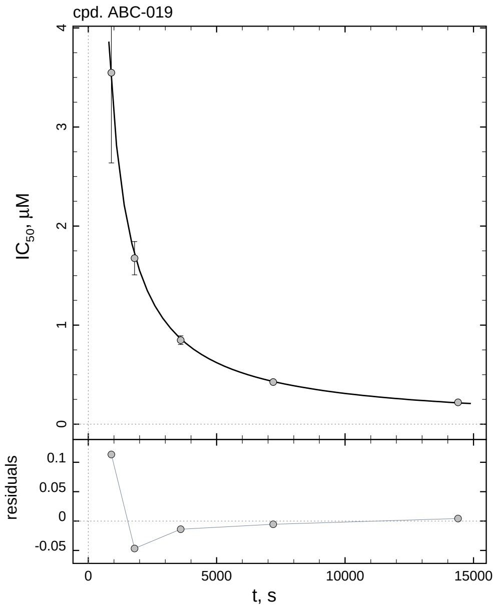

| $i$ | $t$, min | $I_{50}$, nM ± std.err. | CV, % | $k_1^*$, mM$^{-1}$s$^{-1}$ |
|---|---|---|---|---|
| 1 | 15 | 3548.6 ± 911.4 | 25.7 | 0.4990 |
| 2 | 30 | 1675.5 ± 168.3 | 10.0 | 0.5284 |
| 3 | 60 | 848.2 ± 44.11 | 5.2 | 0.5219 |
| 4 | 120 | 425.9 ± 11.81 | 2.8 | 0.5197 |
| 5 | 240 | 220.1 ± 3.161 | 1.4 | 0.5029 |

*Cpd.* **19***: Mechanistic analysis*

| parameter | unit | "true" (T) | calculated (C) | C/T ratio | note |
|---|---|---|---|---|---|
| mechanism | | **C2S** | **C1** | | |
| $k_{\mathrm{eff}}$ | mM$^{-1}$s$^{-1}$ | 0.9901 | 1.039 | 1.05 | $= k_1^* \, (1 + [\mathrm{S}]_0/K_\mathrm{M})$ |

| | |
|---|---|
| Reaction times used for analysis: | 30 and 120 min |
| Maximum GSD for accepting one-step model **C1**: | 1.25 |
| Observed GSD: | 1.01 |
| Assumed $[\mathrm{S}]_0/K_\mathrm{M}$ ratio: | 1.0 |

*2.20. Compound No. 20*

*Cpd.* **20***: Simulated data*

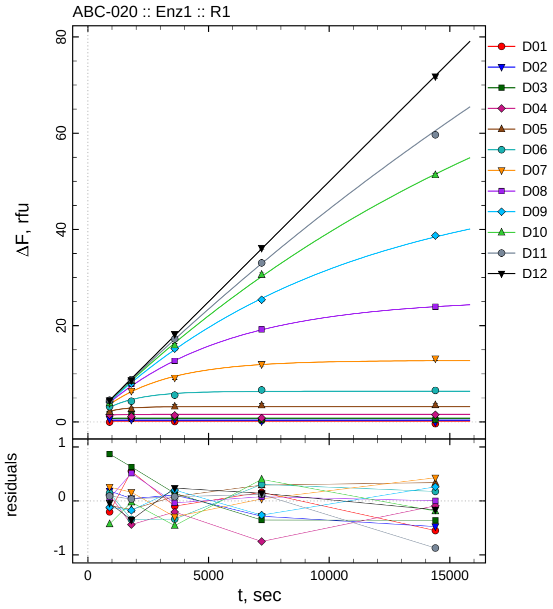

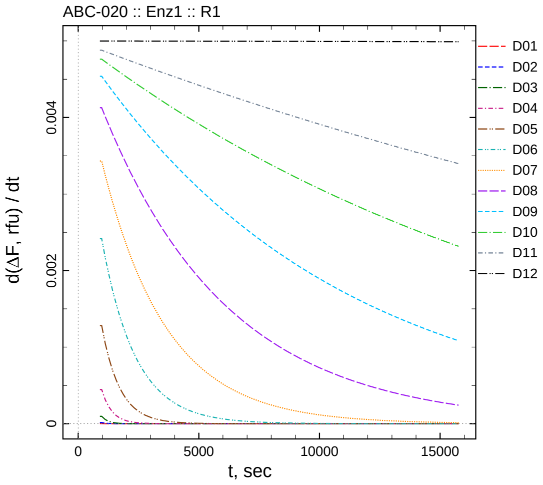

| $[I]_0$, nM | t = 15 | 30 | 60 | 120 | 240 min |
|---|---|---|---|---|---|
| 505000 | -0.003 | 0.752 | 0.105 | 0.361 | -0.350 |
| 252500 | 0.584 | 0.444 | 0.509 | 0.116 | -0.076 |
| 126250 | 1.645 | 1.432 | 0.943 | 0.446 | 0.442 |
| 63125 | 1.523 | 1.139 | 1.394 | 0.848 | 1.510 |
| 31562.5 | 2.164 | 2.761 | 3.258 | 3.493 | 3.541 |
| 15781.3 | 3.216 | 4.321 | 5.591 | 6.672 | 6.576 |
| 7890.63 | 3.932 | 6.455 | 9.199 | 11.980 | 13.169 |
| 3945.31 | 4.104 | 7.980 | 12.717 | 19.229 | 23.969 |
| 1972.66 | 4.150 | 8.004 | 15.253 | 25.410 | 38.718 |
| 986.328 | 3.952 | 8.551 | 15.971 | 30.616 | 51.307 |
| 493.164 | 4.530 | 8.818 | 17.265 | 33.047 | 59.658 |
| 0 | 4.460 | 8.646 | 18.234 | 36.125 | 71.770 |

*Cpd.* **20***: Determination of* $IC_{50}$ *and* $k_1^*$

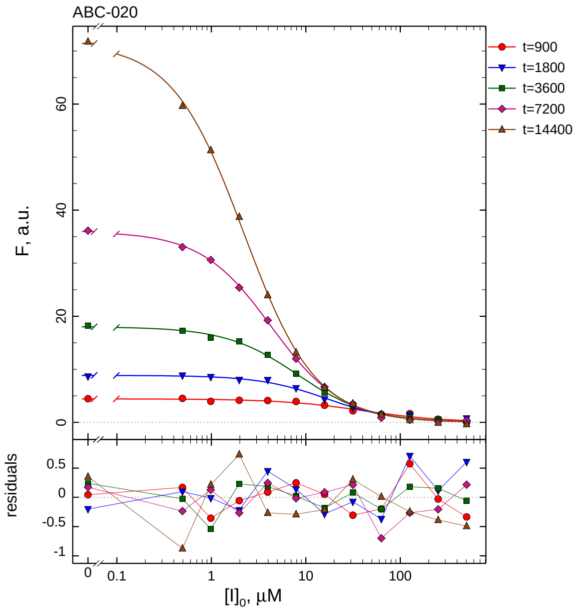

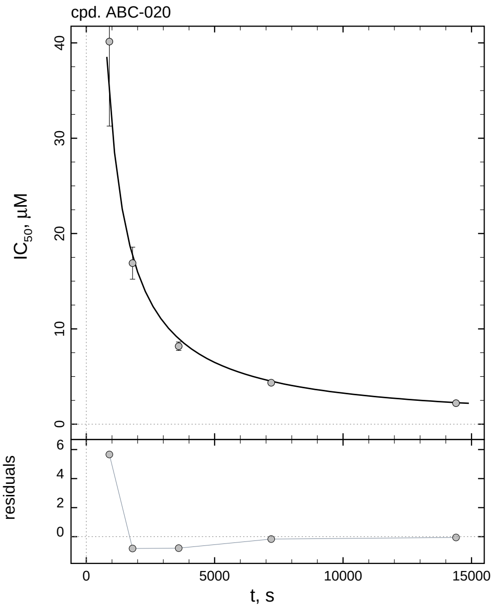

| $i$ | $t$, min | $I_{50}$, nM | ± | std.err. | CV, % | $k_1^*$, $\text{mM}^{-1}\text{s}^{-1}$ |
|---|---|---|---|---|---|---|
| 1 | 15 | 40139 | ± | 8866.7 | 22.1 | 0.04411 |
| 2 | 30 | 16891 | ± | 1676.4 | 9.9 | 0.05242 |
| 3 | 60 | 8173.6 | ± | 445.2 | 5.4 | 0.05416 |
| 4 | 120 | 4346.1 | ± | 122.5 | 2.8 | 0.05093 |
| 5 | 240 | 2205.3 | ± | 32.31 | 1.5 | 0.05018 |

*Cpd.* **20***: Mechanistic analysis*

| parameter | unit | "true" (T) | calculated (C) | C/T ratio | note |
|---|---|---|---|---|---|
| mechanism | | **C2S** | **C1** | | |
| $k_{\text{eff}}$ | $\text{mM}^{-1}\text{s}^{-1}$ | 0.09901 | 0.1019 | 1.03 | $= k_1^* \, (1 + [\text{S}]_0/K_\text{M})$ |

| | |
|---|---|
| Reaction times used for analysis: | 30 and 120 min |
| Maximum GSD for accepting one-step model **C1**: | 1.25 |
| Observed GSD: | 1.02 |
| Assumed $[\text{S}]_0/K_\text{M}$ ratio: | 1.0 |

*2.21. Compound No. 21*

*Cpd.* **21***: Simulated data*

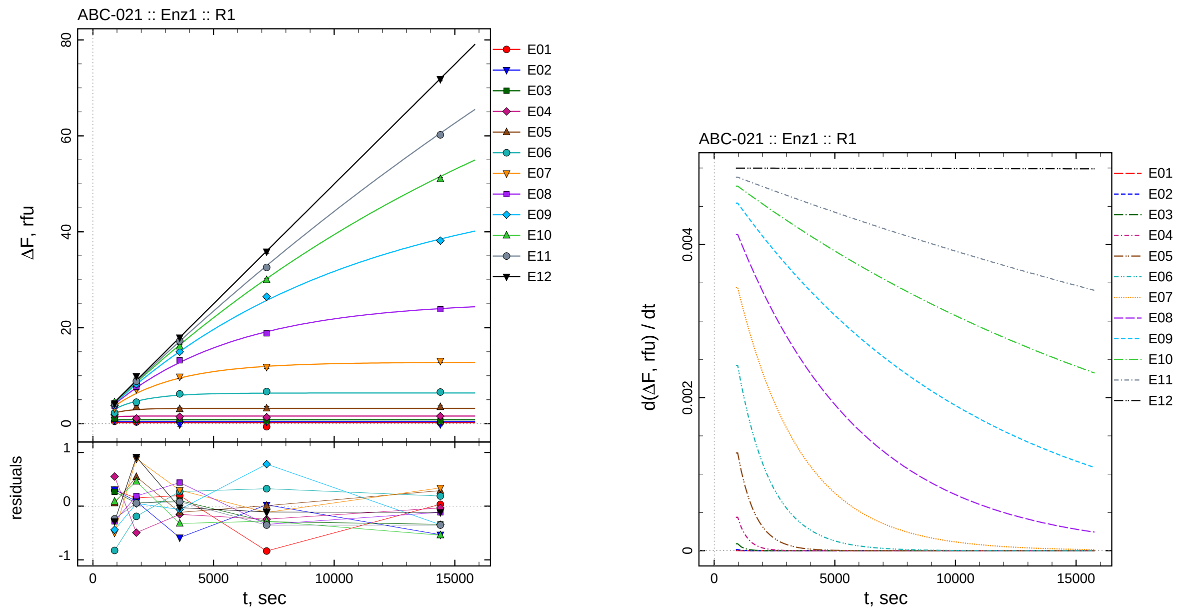

| $[I]_0$, nM | t = 15 | 30 | 60 | 120 | 240 min |
|---:|---:|---:|---:|---:|---:|
| 55 | 0.496 | 0.352 | 0.396 | -0.635 | 0.235 |
| 27.5 | 0.707 | 0.486 | -0.185 | 0.423 | -0.132 |
| 13.75 | 1.045 | 0.856 | 0.902 | 0.509 | 0.461 |
| 6.875 | 1.970 | 1.086 | 1.445 | 1.356 | 1.566 |
| 3.4375 | 2.320 | 3.472 | 3.060 | 3.207 | 3.488 |
| 1.71875 | 2.254 | 4.487 | 6.207 | 6.691 | 6.588 |
| 0.859375 | 3.189 | 7.191 | 9.804 | 11.838 | 13.086 |
| 0.429688 | 3.775 | 7.665 | 13.208 | 18.832 | 23.877 |
| 0.214844 | 3.830 | 8.235 | 14.999 | 26.474 | 38.152 |
| 0.107422 | 4.467 | 9.037 | 16.113 | 29.951 | 50.996 |
| 0.0537109 | 4.202 | 8.841 | 17.271 | 32.580 | 60.208 |
| 0 | 4.214 | 9.911 | 17.966 | 35.872 | 71.819 |

*Cpd.* **21***: Determination of* $IC_{50}$ *and* $k_1^*$

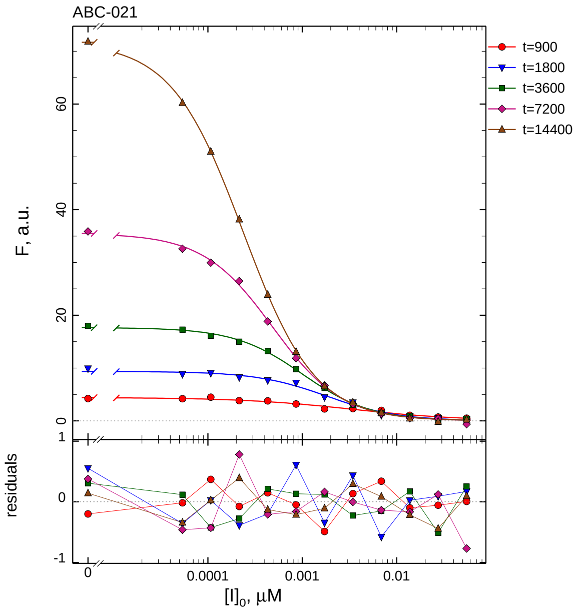

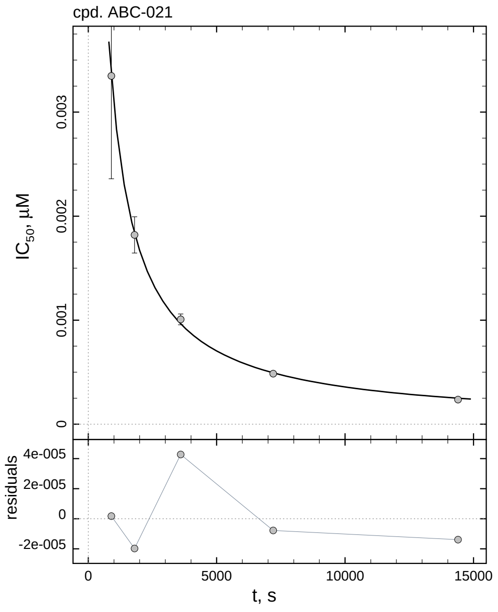

| $i$ | $t$, min | $I_{50}$, nM ± std.err. | CV, % | $k_1^*$, mM$^{-1}$s$^{-1}$ |
|---|---|---|---|---|
| 1 | 15 | 3.348 ± 0.9884 | 29.5 | 528.9 |
| 2 | 30 | 1.820 ± 0.1744 | 9.6 | 486.5 |
| 3 | 60 | 1.007 ± 0.05176 | 5.1 | 439.4 |
| 4 | 120 | 0.4862 ± 0.01342 | 2.8 | 455.3 |
| 5 | 240 | 0.2360 ± 0.003387 | 1.4 | 469.0 |

*Cpd.* **21***: Mechanistic analysis*

| parameter | unit | "true" (T) | calculated (C) | C/T ratio | note |
|---|---|---|---|---|---|
| mechanism | | **C2S** | **C1** | | |
| $k_{\text{eff}}$ | mM$^{-1}$s$^{-1}$ | 909.1 | 910.5 | 1.00 | $= k_1^*\ (1 + [\text{S}]_0/K_\text{M})$ |

| | |
|---|---|
| Reaction times used for analysis: | 30 and 120 min |
| Maximum GSD for accepting one-step model **C1**: | 1.25 |
| Observed GSD: | 1.05 |
| Assumed $[\text{S}]_0/K_\text{M}$ ratio: | 1.0 |

*2.22. Compound No. 22*

*Cpd.* **22***: Simulated data*

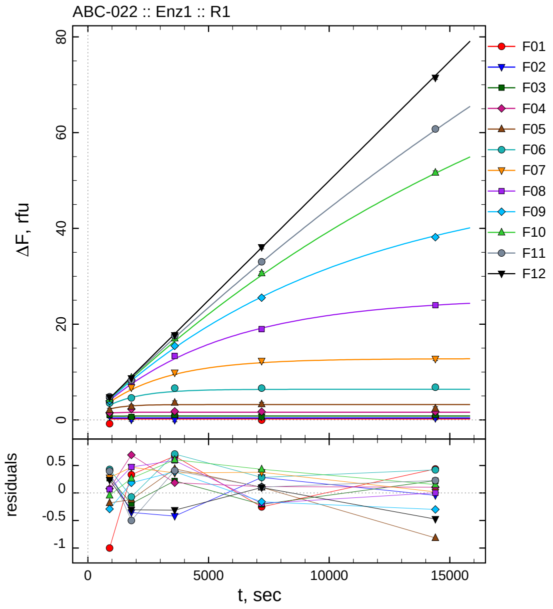

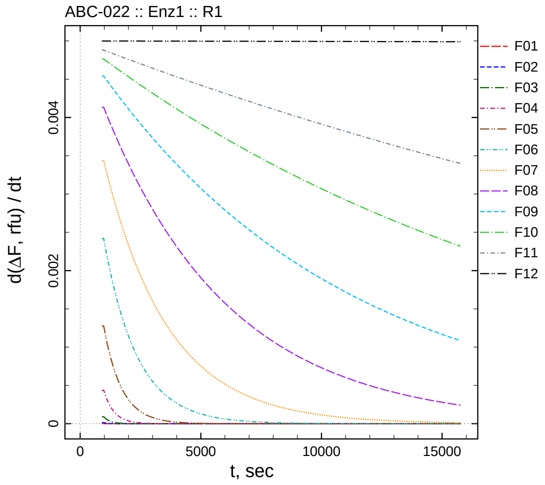

| $[I]_0$, nM | t = 15 | 30 | 60 | 120 | 240 min |
|---|---|---|---|---|---|
| 550 | -0.800 | 0.532 | 0.871 | -0.054 | 0.638 |
| 275 | 0.758 | 0.046 | -0.020 | 0.682 | 0.358 |
| 137.5 | 1.045 | 0.606 | 1.020 | 0.592 | 1.023 |
| 68.75 | 1.499 | 2.272 | 1.787 | 1.716 | 1.706 |
| 34.375 | 2.084 | 2.805 | 3.615 | 3.302 | 2.382 |
| 17.1875 | 3.509 | 4.605 | 6.643 | 6.648 | 6.820 |
| 8.59375 | 3.964 | 6.776 | 9.871 | 12.327 | 12.759 |
| 4.29688 | 4.127 | 7.948 | 13.360 | 18.974 | 23.977 |
| 2.14844 | 3.980 | 8.369 | 15.448 | 25.526 | 38.173 |
| 1.07422 | 4.340 | 8.843 | 17.039 | 30.657 | 51.664 |
| 0.537109 | 4.837 | 8.284 | 17.599 | 33.039 | 60.770 |
| 0 | 4.736 | 8.693 | 17.683 | 36.082 | 71.460 |

*Cpd.* **22***: Determination of* $IC_{50}$ *and* $k_1^*$

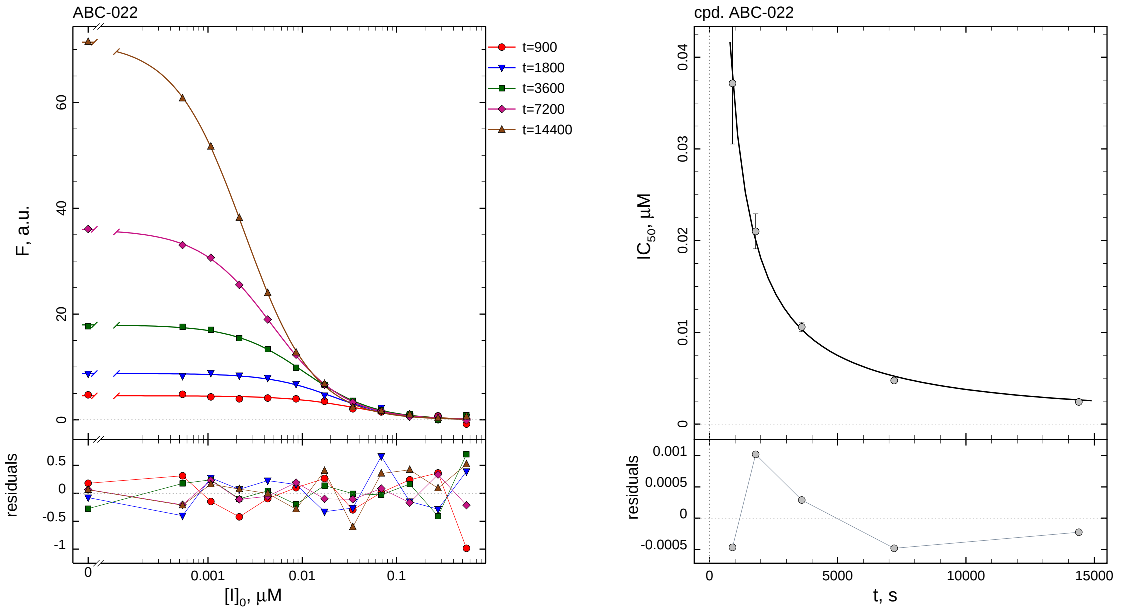

| $i$ | $t$, min | $I_{50}$, nM | ± | std.err. | CV, % | $k_1^*$, mM$^{-1}$s$^{-1}$ |
|---|---|---|---|---|---|---|
| 1 | 15 | 37.14 | ± | 6.605 | 17.8 | 47.67 |
| 2 | 30 | 21.01 | ± | 1.905 | 9.1 | 42.14 |
| 3 | 60 | 10.59 | ± | 0.5305 | 5.0 | 41.79 |
| 4 | 120 | 4.747 | ± | 0.1291 | 2.7 | 46.63 |
| 5 | 240 | 2.409 | ± | 0.03296 | 1.4 | 45.94 |

*Cpd.* **22***: Mechanistic analysis*

| parameter | unit | "true" (T) | calculated (C) | C/T ratio | note |
|---|---|---|---|---|---|
| mechanism | | **C2S** | **C1** | | |
| $k_{\mathrm{eff}}$ | mM$^{-1}$s$^{-1}$ | 90.91 | 93.26 | 1.03 | $= k_1^*\ (1 + [\mathrm{S}]_0/K_\mathrm{M})$ |

| | |
|---|---|
| Reaction times used for analysis: | 30 and 120 min |
| Maximum GSD for accepting one-step model **C1**: | 1.25 |
| Observed GSD: | 1.07 |
| Assumed $[\mathrm{S}]_0/K_\mathrm{M}$ ratio: | 1.0 |

*2.23. Compound No. 23*

*Cpd.* **23***: Simulated data*

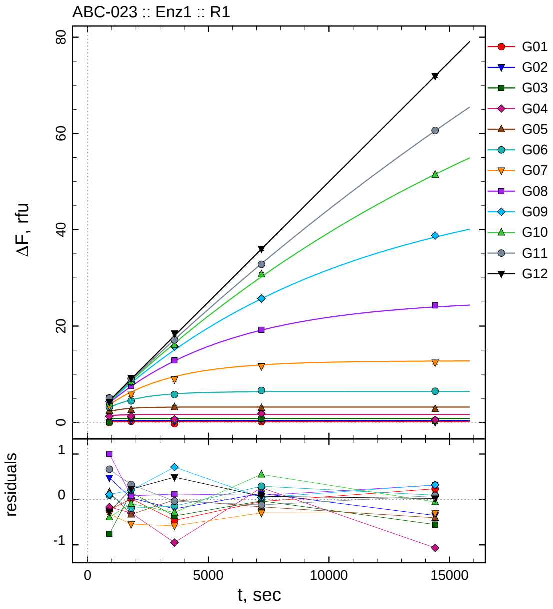

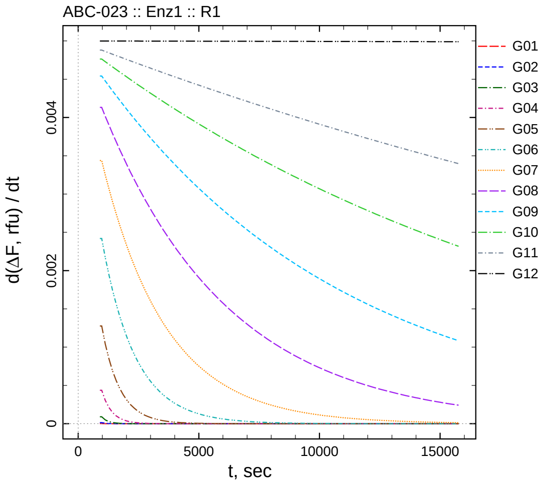

| $[I]_0$, nM | t = 15 | 30 | 60 | 120 | 240 min |
|---|---|---|---|---|---|
| 5500 | -0.033 | 0.218 | -0.264 | 0.155 | 0.435 |
| 2750 | 0.875 | 0.428 | 0.193 | 0.530 | 0.045 |
| 1375 | 0.015 | 0.949 | 0.436 | 0.766 | 0.247 |
| 687.5 | 1.253 | 1.280 | 0.650 | 1.856 | 0.534 |
| 343.75 | 2.425 | 2.594 | 3.168 | 3.034 | 2.795 |
| 171.875 | 3.171 | 4.483 | 5.787 | 6.655 | 6.491 |
| 85.9375 | 3.366 | 5.763 | 8.925 | 11.655 | 12.438 |
| 42.9688 | 5.061 | 7.554 | 12.886 | 19.253 | 24.284 |
| 21.4844 | 4.380 | 8.386 | 15.771 | 25.725 | 38.789 |
| 10.7422 | 3.990 | 8.483 | 16.145 | 30.777 | 51.452 |
| 5.37109 | 5.103 | 9.111 | 17.152 | 32.810 | 60.623 |
| 0 | 4.230 | 9.225 | 18.483 | 36.049 | 71.954 |

*Cpd.* **23***: Determination of* $IC_{50}$ *and* $k_1^*$

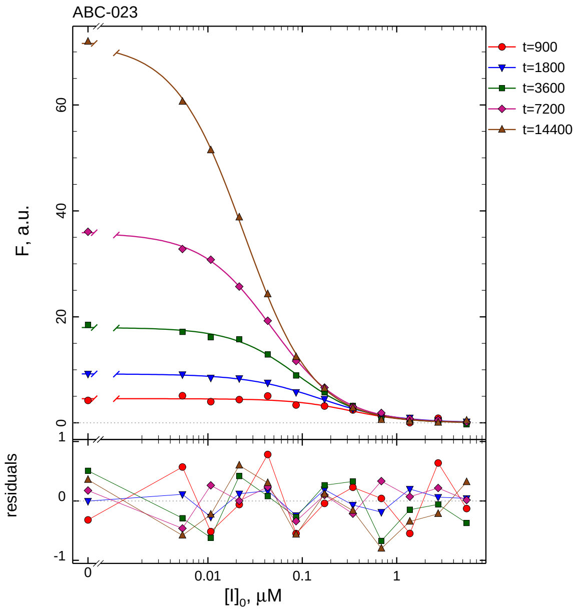

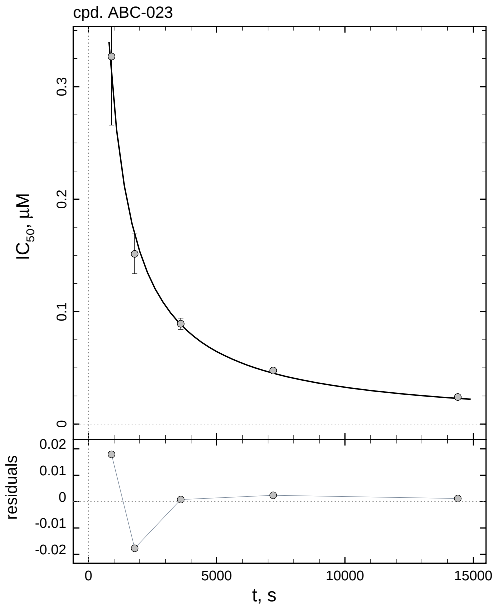

| $i$ | $t$, min | $I_{50}$, nM | ± | std.err. | CV, % | $k_1^*$, mM$^{-1}$s$^{-1}$ |
|---|---|---|---|---|---|---|
| 1 | 15 | 326.9 | ± | 60.94 | 18.6 | 5.417 |
| 2 | 30 | 151.4 | ± | 17.69 | 11.7 | 5.848 |
| 3 | 60 | 89.24 | ± | 5.026 | 5.6 | 4.960 |
| 4 | 120 | 47.65 | ± | 1.493 | 3.1 | 4.645 |
| 5 | 240 | 24.06 | ± | 0.3832 | 1.6 | 4.600 |

*Cpd.* **23***: Mechanistic analysis*

| parameter | unit | "true" (T) | calculated (C) | C/T ratio | note |
|---|---|---|---|---|---|
| mechanism | | **C2S** | **C1** | | |
| $k_{\mathrm{eff}}$ | mM$^{-1}$s$^{-1}$ | 9.091 | 9.290 | 1.02 | $= k_1^* \, (1 + [S]_0/K_M)$ |

| | |
|---|---|
| Reaction times used for analysis: | 30 and 120 min |
| Maximum GSD for accepting one-step model **C1**: | 1.25 |
| Observed GSD: | 1.18 |
| Assumed $[S]_0/K_M$ ratio: | 1.0 |

#### 2.24. Compound No. 24

*Cpd.* **24**: *Simulated data*

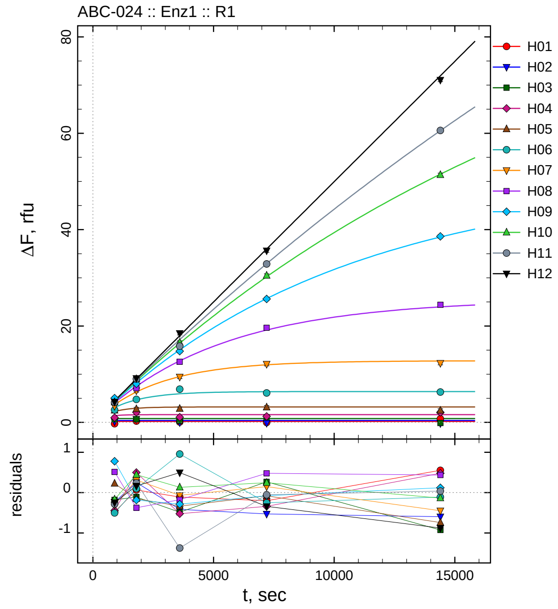

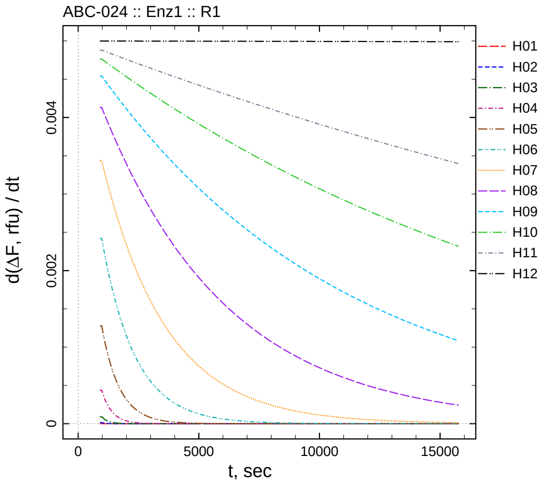

| $[I]_0$, nM | t = 15 | 30 | 60 | 120 | 240 min |
|---:|---:|---:|---:|---:|---:|
| 55000 | -0.279 | 0.266 | 0.091 | -0.002 | 0.752 |
| 27500 | 0.167 | 0.651 | -0.015 | -0.127 | -0.196 |
| 13750 | 0.597 | 0.689 | 0.316 | 1.067 | -0.127 |
| 6875 | 0.978 | 2.077 | 1.077 | 1.260 | 2.079 |
| 3437.5 | 2.490 | 2.777 | 2.842 | 3.121 | 2.457 |
| 1718.75 | 2.579 | 4.779 | 6.891 | 6.113 | 6.288 |
| 859.375 | 3.382 | 6.619 | 9.438 | 12.100 | 12.292 |
| 429.688 | 4.574 | 7.100 | 12.593 | 19.641 | 24.409 |
| 214.844 | 5.047 | 8.003 | 14.775 | 25.610 | 38.590 |
| 107.422 | 4.214 | 9.021 | 16.567 | 30.472 | 51.374 |
| 53.7109 | 4.155 | 9.022 | 15.819 | 32.878 | 60.578 |
| 0 | 4.251 | 9.165 | 18.490 | 35.643 | 71.057 |

*Cpd.* **24**: *Determination of* $IC_{50}$ *and* $k_1^*$

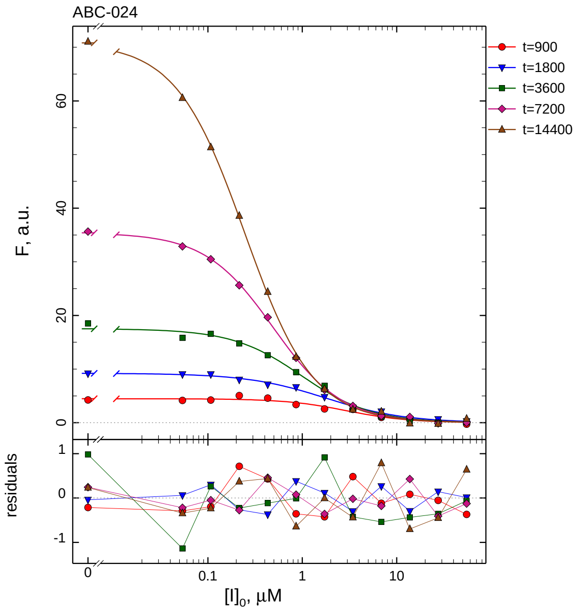

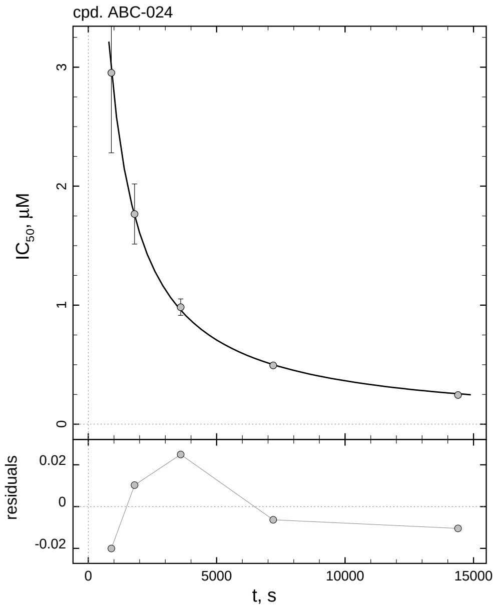

| $i$ | $t$, min | $I_{50}$, nM | ± | std.err. | CV, % | $k_1^*$, mM$^{-1}$s$^{-1}$ |
|---|---|---|---|---|---|---|
| 1 | 15 | 2951.8 | ± | 672.1 | 22.8 | 0.5999 |
| 2 | 30 | 1765.8 | ± | 252.6 | 14.3 | 0.5014 |
| 3 | 60 | 983.0 | ± | 69.05 | 7.0 | 0.4503 |
| 4 | 120 | 494.2 | ± | 17.84 | 3.6 | 0.4478 |
| 5 | 240 | 245.4 | ± | 4.572 | 1.9 | 0.4510 |

*Cpd.* **24**: *Mechanistic analysis*

| parameter | unit | “true” (T) | calculated (C) | C/T ratio | note |
|---|---|---|---|---|---|
| mechanism | | **C2S** | **C1** | | |
| $k_{\mathrm{eff}}$ | mM$^{-1}$s$^{-1}$ | 0.9091 | 0.8957 | 0.99 | $= k_1^* \, (1 + [S]_0/K_M)$ |

| | |
|---|---|
| Reaction times used for analysis: | 30 and 120 min |
| Maximum GSD for accepting one-step model **C1**: | 1.25 |
| Observed GSD: | 1.08 |
| Assumed $[S]_0/K_M$ ratio: | 1.0 |

### 2.25. *Compound No. 25*

*Cpd.* **25***: Simulated data*

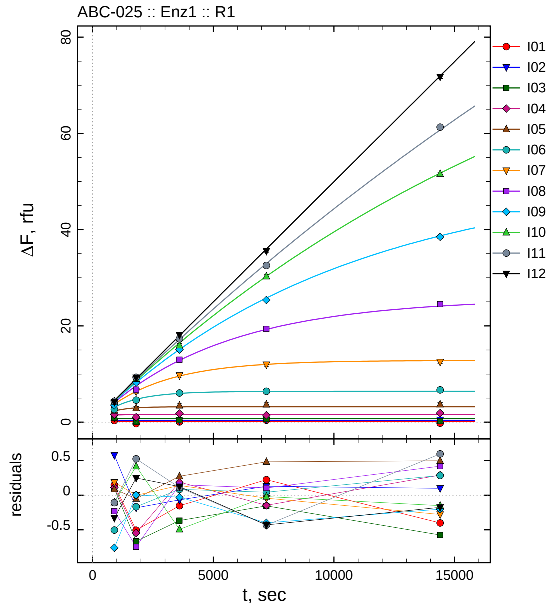

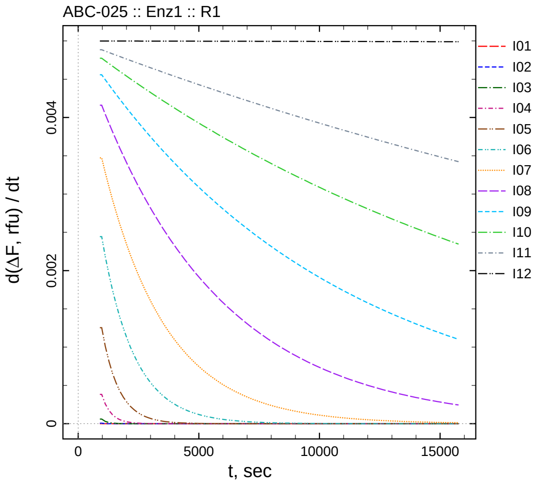

| $[I]_0$, nM | t = 15 | 30 | 60 | 120 | 240 min |
|---:|---:|---:|---:|---:|---:|
| 10 | 0.349 | -0.306 | 0.048 | 0.423 | -0.201 |
| 5 | 0.977 | 0.218 | 0.332 | 0.529 | 0.499 |
| 2.5 | 0.882 | 0.134 | 0.433 | 0.646 | 0.224 |
| 1.25 | 1.567 | 1.041 | 1.789 | 1.454 | 1.891 |
| 0.625 | 2.419 | 2.914 | 3.458 | 3.685 | 3.700 |
| 0.3125 | 2.644 | 4.581 | 6.061 | 6.423 | 6.694 |
| 0.15625 | 3.921 | 6.378 | 9.726 | 11.969 | 12.507 |
| 0.078125 | 3.861 | 6.782 | 12.998 | 19.379 | 24.535 |
| 0.0390625 | 3.528 | 8.220 | 15.090 | 25.396 | 38.503 |
| 0.0195313 | 4.287 | 9.013 | 15.977 | 30.286 | 51.584 |
| 0.00976563 | 4.337 | 9.319 | 17.320 | 32.545 | 61.295 |
| 0 | 4.169 | 9.247 | 18.119 | 35.554 | 71.759 |

*Cpd.* **25***: Determination of* $IC_{50}$ *and* $k_1^*$

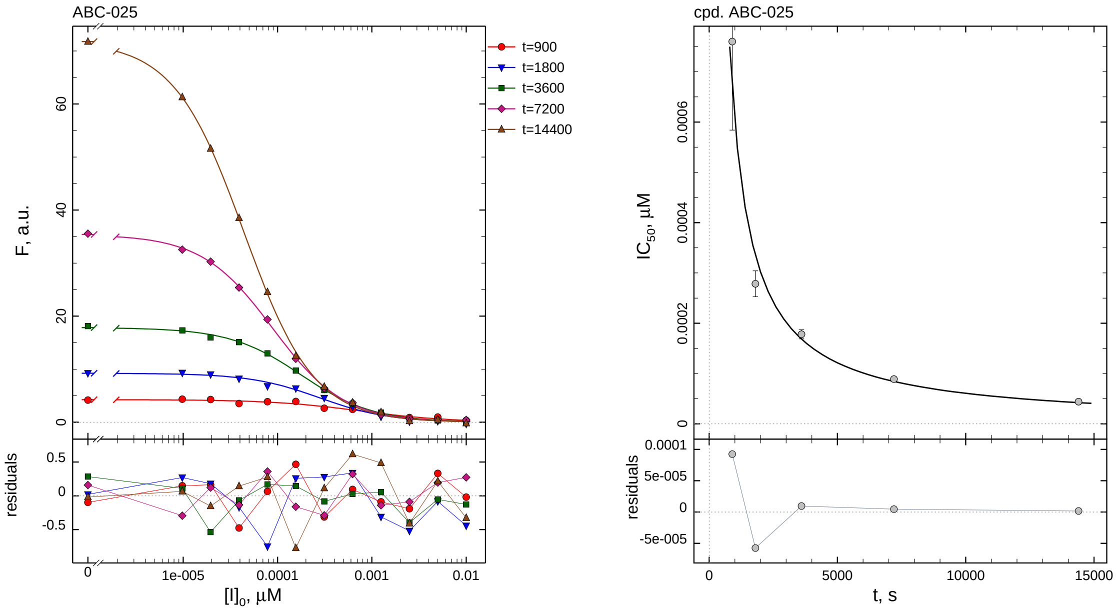

| $i$ | $t$, min | $I_{50}$, nM | ± | std.err. | CV, % | $k_1^*$, mM$^{-1}$s$^{-1}$ |
|---|---|---|---|---|---|---|
| 1 | 15 | 0.7600 | ± | 0.1761 | 23.2 | 2329.8 |
| 2 | 30 | 0.2784 | ± | 0.02570 | 9.2 | 3179.7 |
| 3 | 60 | 0.1780 | ± | 0.009055 | 5.1 | 2486.9 |
| 4 | 120 | 0.08893 | ± | 0.002385 | 2.7 | 2488.8 |
| 5 | 240 | 0.04396 | ± | 0.0005905 | 1.3 | 2517.7 |

*Cpd.* **25***: Mechanistic analysis*

| parameter | unit | "true" (T) | calculated (C) | C/T ratio | note |
|---|---|---|---|---|---|
| mechanism | | **C2S** | **C1** | | |
| $k_{\text{eff}}$ | mM$^{-1}$s$^{-1}$ | 5000.0 | 4977.5 | 1.00 | $= k_1^* \ (1 + [\text{S}]_0/K_\text{M})$ |

| | |
|---|---|
| Reaction times used for analysis: | 30 and 120 min |
| Maximum GSD for accepting one-step model **C1**: | 1.25 |
| Observed GSD: | 1.19 |
| Assumed $[\text{S}]_0/K_\text{M}$ ratio: | 1.0 |

*Cpd.* **26***: Simulated data*

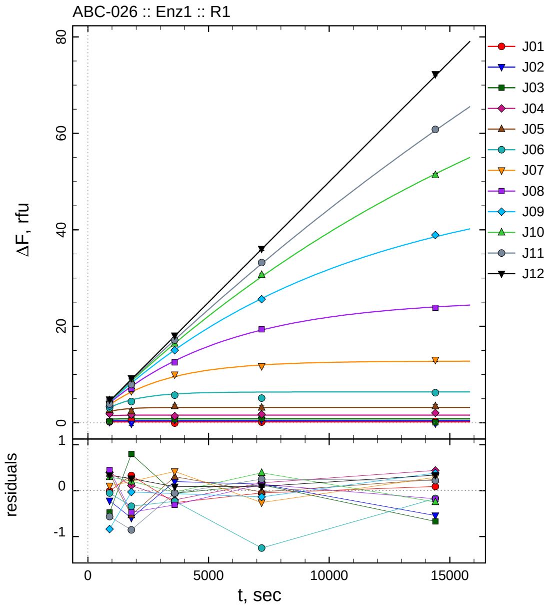

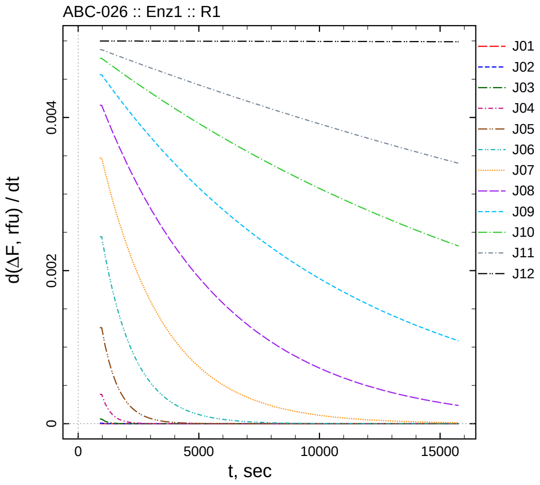

| $[I]_0$, nM | t = 15 | 30 | 60 | 120 | 240 min |
|---:|---:|---:|---:|---:|---:|
| 100 | 0.192 | 0.531 | -0.062 | 0.150 | 0.291 |
| 50 | 0.178 | -0.194 | 0.598 | 0.540 | -0.142 |
| 25 | 0.313 | 1.597 | 0.743 | 0.928 | 0.129 |
| 12.5 | 1.855 | 1.696 | 1.398 | 1.770 | 2.040 |
| 6.25 | 2.700 | 2.430 | 3.485 | 3.172 | 3.443 |
| 3.125 | 3.095 | 4.406 | 5.740 | 5.122 | 6.229 |
| 1.5625 | 3.839 | 6.574 | 9.993 | 11.728 | 13.040 |
| 0.78125 | 4.539 | 7.052 | 12.527 | 19.376 | 23.842 |
| 0.390625 | 3.450 | 8.186 | 15.047 | 25.623 | 38.939 |
| 0.195313 | 4.689 | 8.800 | 16.434 | 30.668 | 51.348 |
| 0.0976563 | 3.880 | 7.938 | 17.154 | 33.213 | 60.823 |
| 0 | 4.833 | 9.260 | 18.080 | 36.074 | 72.272 |

*Cpd.* **26***: Determination of* $IC_{50}$ *and* $k_1^*$

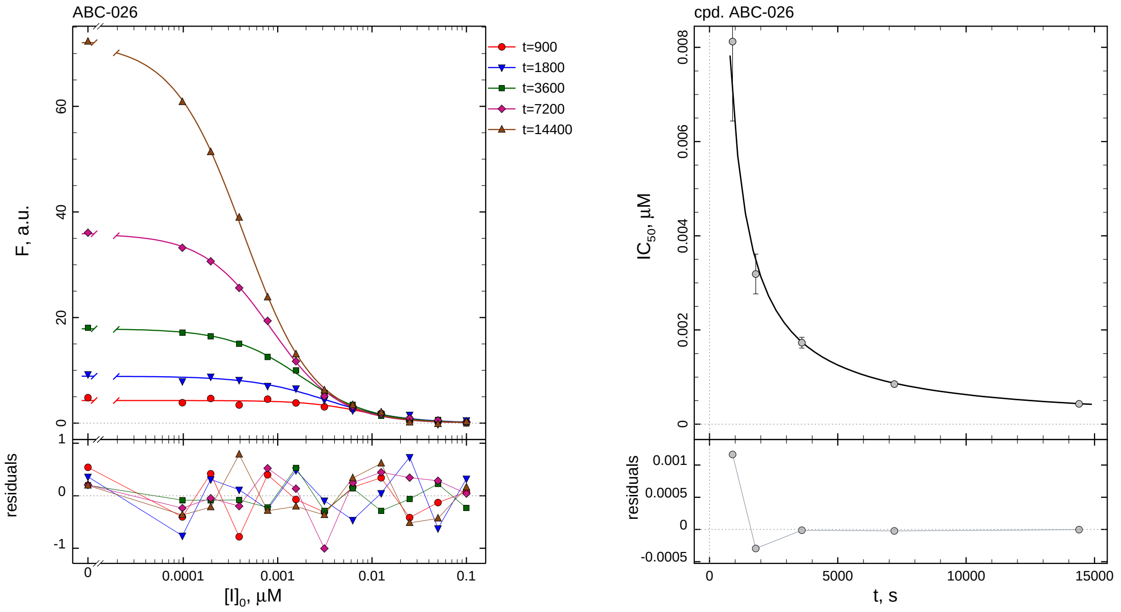

| $i$ | $t$, min | $I_{50}$, nM | ± | std.err. | CV, % | $k_1^*$, mM$^{-1}$s$^{-1}$ |
|---|---|---|---|---|---|---|
| 1 | 15 | 8.123 | ± | 1.687 | 20.8 | 218.0 |
| 2 | 30 | 3.189 | ± | 0.4231 | 13.3 | 277.6 |
| 3 | 60 | 1.731 | ± | 0.1148 | 6.6 | 255.7 |
| 4 | 120 | 0.8501 | ± | 0.02826 | 3.3 | 260.4 |
| 5 | 240 | 0.4323 | ± | 0.007594 | 1.8 | 256.0 |

*Cpd.* **26***: Mechanistic analysis*

| parameter | unit | "true" (T) | calculated (C) | C/T ratio | note |
|---|---|---|---|---|---|
| mechanism | | **C2S** | **C1** | | |
| $k_{\mathrm{eff}}$ | mM$^{-1}$s$^{-1}$ | 500.0 | 520.7 | 1.04 | $= k_1^* \, (1 + [\mathrm{S}]_0/K_\mathrm{M})$ |

| | |
|---|---|
| Reaction times used for analysis: | 30 and 120 min |
| Maximum GSD for accepting one-step model **C1**: | 1.25 |
| Observed GSD: | 1.05 |
| Assumed $[\mathrm{S}]_0/K_\mathrm{M}$ ratio: | 1.0 |

*2.27. Compound No. 27*

*Cpd.* ***27****: Simulated data*

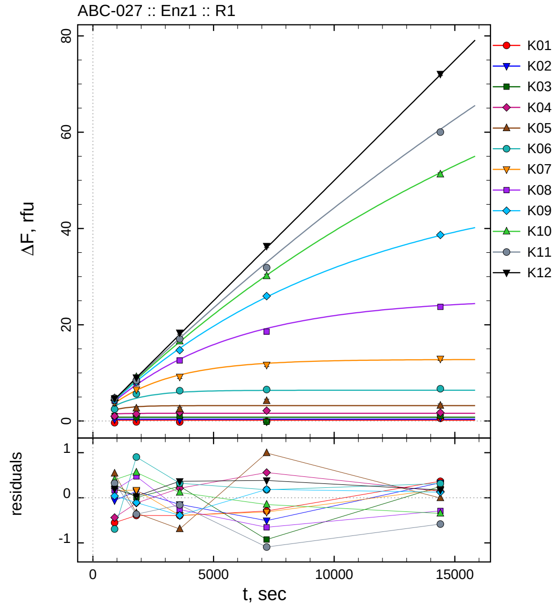

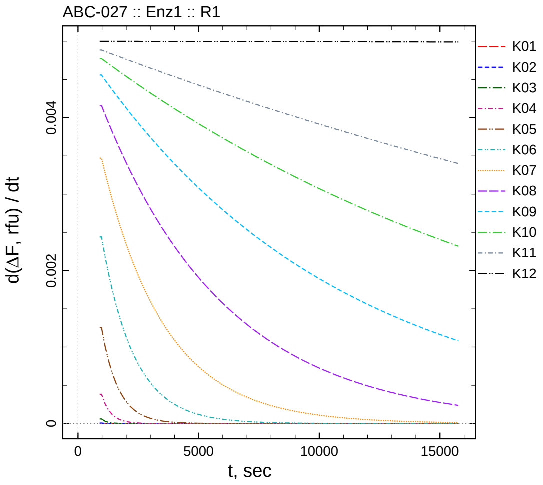

| $[I]_0$, nM | t = 15 | 30 | 60 | 120 | 240 min |
|---|---|---|---|---|---|
| 1000 | -0.352 | -0.191 | -0.195 | -0.094 | 0.567 |
| 500 | 0.341 | 0.542 | 0.253 | -0.106 | 0.734 |
| 250 | 1.092 | 0.807 | 1.054 | -0.126 | 1.051 |
| 125 | 1.017 | 1.471 | 1.811 | 2.160 | 1.743 |
| 62.5 | 2.868 | 2.622 | 2.491 | 4.191 | 3.194 |
| 31.25 | 2.455 | 5.647 | 6.294 | 6.548 | 6.715 |
| 15.625 | 3.891 | 6.548 | 9.186 | 11.669 | 12.934 |
| 7.8125 | 4.259 | 8.000 | 12.576 | 18.580 | 23.718 |
| 3.90625 | 4.327 | 8.109 | 14.721 | 25.942 | 38.666 |
| 1.95313 | 4.789 | 9.157 | 16.583 | 30.128 | 51.235 |
| 0.976563 | 4.767 | 8.429 | 17.049 | 31.871 | 60.009 |
| 0 | 4.696 | 9.034 | 18.362 | 36.368 | 72.108 |

*Cpd.* **27***: Determination of* $IC_{50}$ *and* $k_1^*$

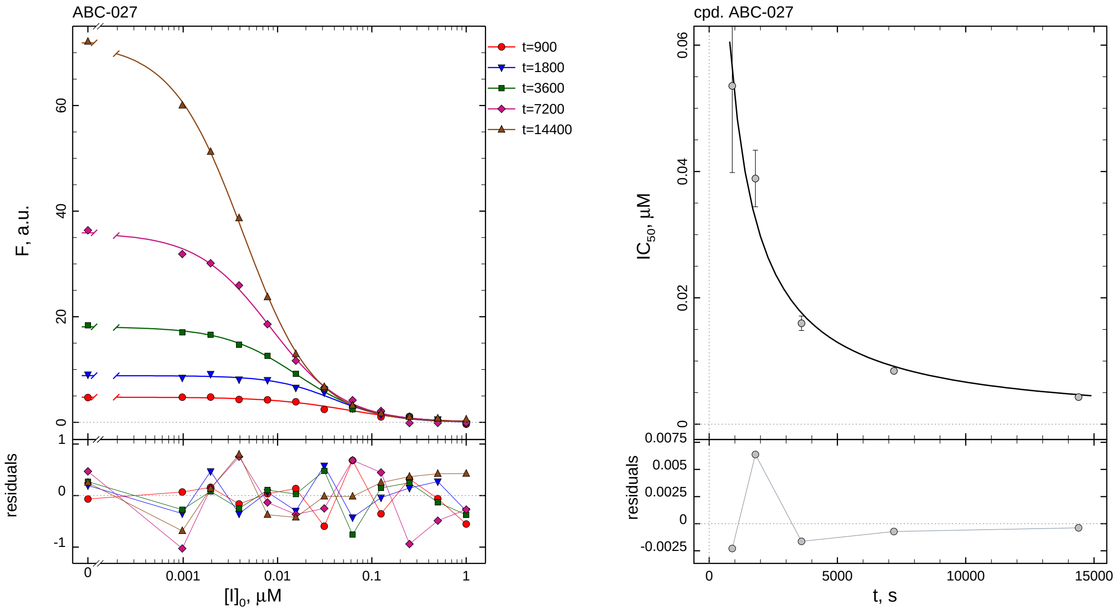

| $i$ | $t$, min | $I_{50}$, nM | ± | std.err. | CV, % | $k_1^*$, $\text{mM}^{-1}\text{s}^{-1}$ |
|---|---|---|---|---|---|---|
| 1 | 15 | 53.54 | ± | 13.72 | 25.6 | 33.07 |
| 2 | 30 | 38.89 | ± | 4.478 | 11.5 | 22.77 |
| 3 | 60 | 15.97 | ± | 1.148 | 7.2 | 27.72 |
| 4 | 120 | 8.435 | ± | 0.3210 | 3.8 | 26.24 |
| 5 | 240 | 4.294 | ± | 0.08128 | 1.9 | 25.77 |

*Cpd.* **27***: Mechanistic analysis*

| parameter | unit | "true" (T) | calculated (C) | C/T ratio | note |
|---|---|---|---|---|---|
| mechanism | | **C2S** | **C1** | | |
| $k_{\text{eff}}$ | $\text{mM}^{-1}\text{s}^{-1}$ | 50.00 | 52.48 | 1.05 | $= k_1^* \, (1 + [S]_0/K_M)$ |

| | |
|---|---|
| Reaction times used for analysis: | 30 and 120 min |
| Maximum GSD for accepting one-step model **C1**: | 1.25 |
| Observed GSD: | 1.11 |
| Assumed $[S]_0/K_M$ ratio: | 1.0 |

#### 2.28. *Compound No. 28*

*Cpd.* **28***: Simulated data*

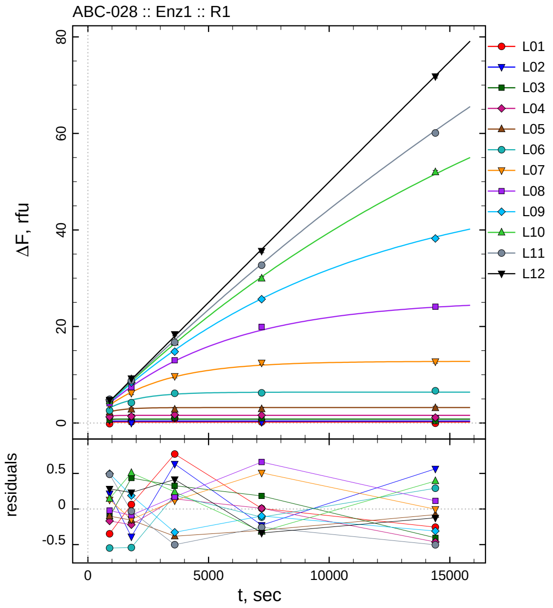

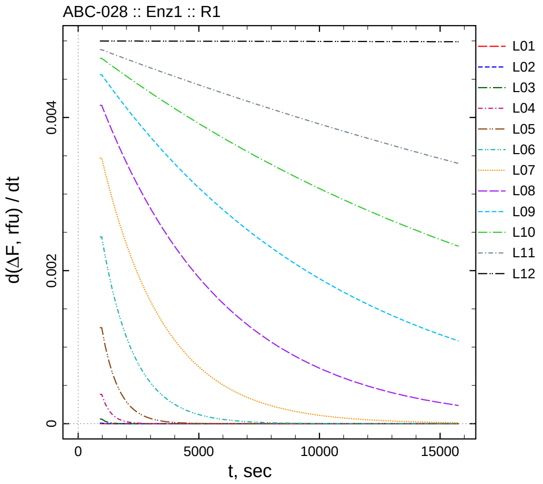

| $[I]_0$, nM | t = 15 | 30 | 60 | 120 | 240 min |
|---:|---:|---:|---:|---:|---:|
| 10000 | -0.149 | 0.263 | 0.973 | 0.208 | -0.055 |
| 5000 | 0.615 | 0.011 | 1.033 | 0.173 | 0.965 |
| 2500 | 0.682 | 1.236 | 1.128 | 0.985 | 0.396 |
| 1250 | 1.288 | 1.366 | 1.743 | 1.611 | 1.137 |
| 625 | 2.230 | 2.805 | 2.799 | 2.900 | 3.122 |
| 312.5 | 2.600 | 4.203 | 6.163 | 6.256 | 6.691 |
| 156.25 | 3.864 | 6.231 | 9.695 | 12.494 | 12.741 |
| 78.125 | 4.068 | 7.440 | 13.010 | 19.894 | 24.122 |
| 39.0625 | 4.784 | 8.404 | 14.784 | 25.658 | 38.230 |
| 19.5313 | 4.537 | 9.103 | 16.706 | 29.960 | 51.973 |
| 9.76563 | 4.932 | 8.763 | 16.705 | 32.705 | 60.087 |
| 0 | 4.780 | 9.228 | 18.410 | 35.646 | 71.810 |

*Cpd.* **28***: Determination of* $IC_{50}$ *and* $k_1^*$

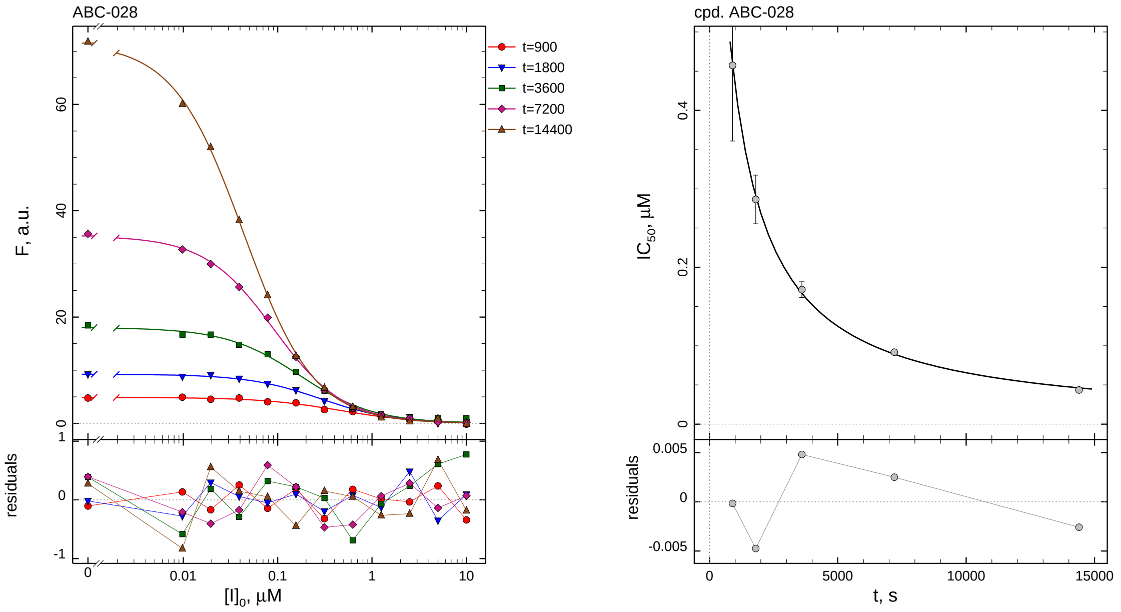

| $i$ | $t$, min | $I_{50}$, nM | ± | std.err. | CV, % | $k_1^*$, mM$^{-1}$s$^{-1}$ |
|---|---|---|---|---|---|---|
| 1 | 15 | 457.4 | ± | 96.48 | 21.1 | 3.871 |
| 2 | 30 | 286.5 | ± | 30.98 | 10.8 | 3.090 |
| 3 | 60 | 171.5 | ± | 10.07 | 5.9 | 2.581 |
| 4 | 120 | 91.83 | ± | 2.728 | 3.0 | 2.410 |
| 5 | 240 | 43.66 | ± | 0.6633 | 1.5 | 2.535 |

*Cpd.* **28***: Mechanistic analysis*

| parameter | unit | "true" (T) | calculated (C) | C/T ratio | note |
|---|---|---|---|---|---|
| mechanism | | **C2S** | **C1** | | |
| $k_{\mathrm{eff}}$ | mM$^{-1}$s$^{-1}$ | 5.000 | 4.820 | 0.96 | $= k_1^* \, (1 + [\mathrm{S}]_0/K_\mathrm{M})$ |

| | |
|---|---|
| Reaction times used for analysis: | 30 and 120 min |
| Maximum GSD for accepting one-step model **C1**: | 1.25 |
| Observed GSD: | 1.19 |
| Assumed $[\mathrm{S}]_0/K_\mathrm{M}$ ratio: | 1.0 |

*2.29. Compound No. 29*

*Cpd.* **29***: Simulated data*

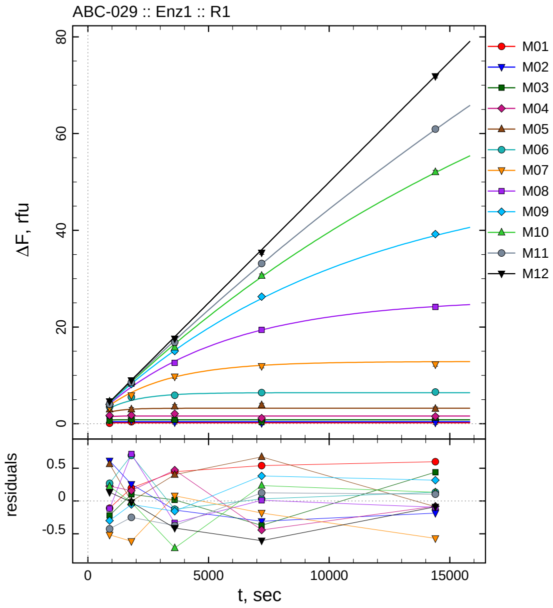

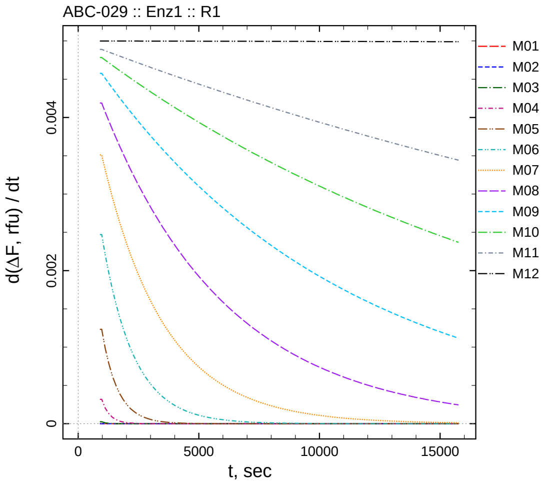

| $[I]_0$, nM | t = 15 | 30 | 60 | 120 | 240 min |
|---:|---:|---:|---:|---:|---:|
| 5.5 | 0.090 | 0.398 | 0.646 | 0.740 | 0.799 |
| 2.75 | 1.012 | 0.656 | 0.263 | 0.092 | 0.214 |
| 1.375 | 0.576 | 0.902 | 0.817 | 0.429 | 1.240 |
| 0.6875 | 1.721 | 1.752 | 2.072 | 1.160 | 1.521 |
| 0.34375 | 2.964 | 2.997 | 3.592 | 3.877 | 3.117 |
| 0.171875 | 3.487 | 5.514 | 5.893 | 6.425 | 6.551 |
| 0.0859375 | 3.268 | 5.837 | 9.742 | 11.887 | 12.251 |
| 0.0429688 | 4.002 | 8.294 | 12.593 | 19.389 | 24.143 |
| 0.0214844 | 4.005 | 8.194 | 15.019 | 26.279 | 39.225 |
| 0.0107422 | 4.621 | 8.591 | 15.787 | 30.612 | 52.064 |
| 0.00537109 | 4.023 | 8.551 | 16.853 | 33.145 | 60.939 |
| 0 | 4.632 | 8.979 | 17.582 | 35.378 | 71.846 |

*Cpd.* **29***: Determination of* $IC_{50}$ *and* $k_1^*$

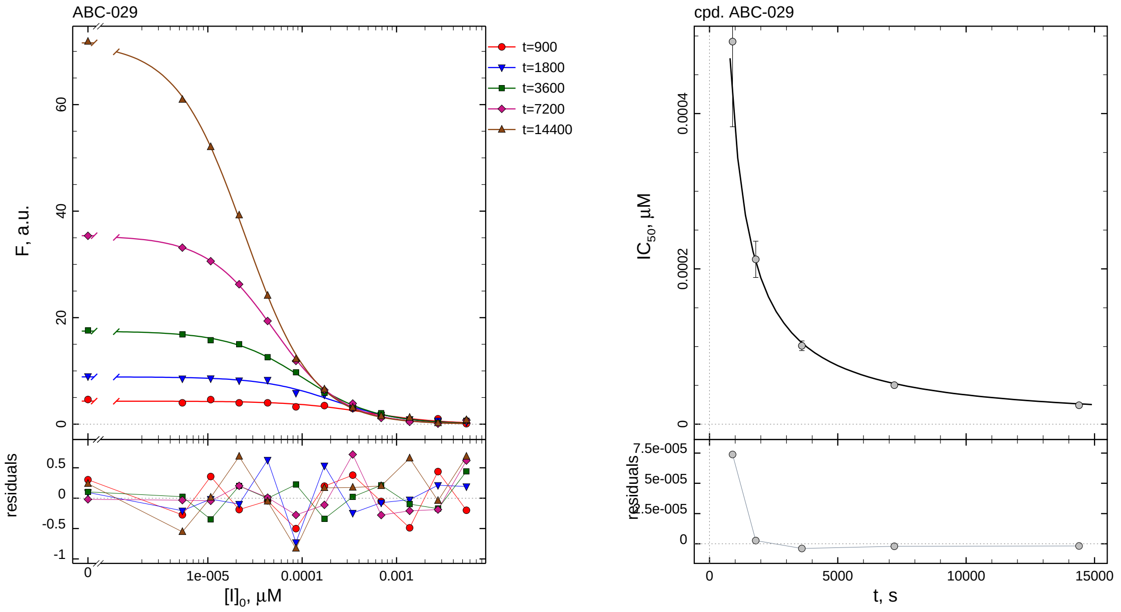

| $i$ | $t$, min | $I_{50}$, nM | ± | std.err. | CV, % | $k_1^*$, mM$^{-1}$s$^{-1}$ |
|---|---|---|---|---|---|---|
| 1 | 15 | 0.4928 | ± | 0.1098 | 22.3 | 3593.2 |
| 2 | 30 | 0.2123 | ± | 0.02336 | 11.0 | 4170.9 |
| 3 | 60 | 0.1009 | ± | 0.006223 | 6.2 | 4385.4 |
| 4 | 120 | 0.05031 | ± | 0.001501 | 3.0 | 4399.7 |
| 5 | 240 | 0.02443 | ± | 0.0003758 | 1.5 | 4529.6 |

*Cpd.* **29***: Mechanistic analysis*

| parameter | unit | "true" (T) | calculated (C) | C/T ratio | note |
|---|---|---|---|---|---|
| mechanism | | **C2S** | **C1** | | |
| $k_{\mathrm{eff}}$ | mM$^{-1}$s$^{-1}$ | 9090.9 | 8799.3 | 0.97 | $= k_1^* \, (1 + [\mathrm{S}]_0/K_\mathrm{M})$ |

| | |
|---|---|
| Reaction times used for analysis: | 30 and 120 min |
| Maximum GSD for accepting one-step model **C1**: | 1.25 |
| Observed GSD: | 1.04 |
| Assumed $[\mathrm{S}]_0/K_\mathrm{M}$ ratio: | 1.0 |

*2.30. Compound No. 30*

*Cpd.* **30***: Simulated data*

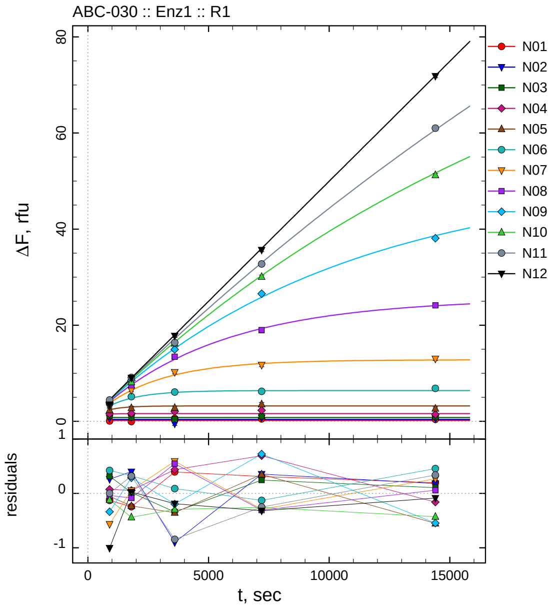

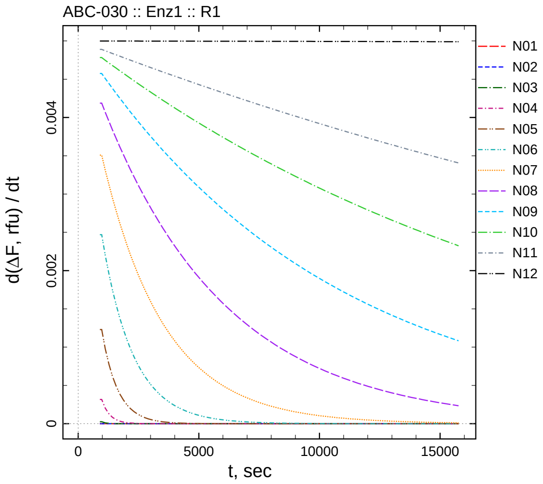

| $[I]_0$, nM | t = 15 | 30 | 60 | 120 | 240 min |
|---|---|---|---|---|---|
| 55 | 0.072 | -0.045 | 0.595 | 0.510 | 0.398 |
| 27.5 | 0.663 | 0.800 | -0.502 | 0.755 | 0.582 |
| 13.75 | 1.112 | 0.817 | 0.459 | 1.045 | 0.907 |
| 6.875 | 1.570 | 1.647 | 2.038 | 2.294 | 1.439 |
| 3.4375 | 2.346 | 2.770 | 2.832 | 3.551 | 2.649 |
| 1.71875 | 3.639 | 5.140 | 6.097 | 6.249 | 6.857 |
| 0.859375 | 3.216 | 6.504 | 10.239 | 11.735 | 13.026 |
| 0.429688 | 4.083 | 7.491 | 13.457 | 19.003 | 24.122 |
| 0.214844 | 3.966 | 8.532 | 14.965 | 26.558 | 38.093 |
| 0.107422 | 4.281 | 8.171 | 16.204 | 30.076 | 51.250 |
| 0.0537109 | 4.450 | 9.120 | 16.379 | 32.744 | 60.999 |
| 0 | 3.494 | 9.023 | 17.805 | 35.663 | 71.847 |

*Cpd.* **30***: Determination of* $IC_{50}$ *and* $k_1^*$

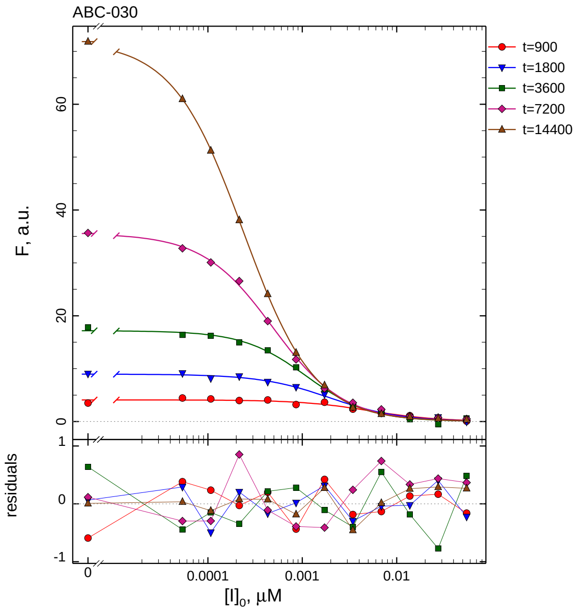

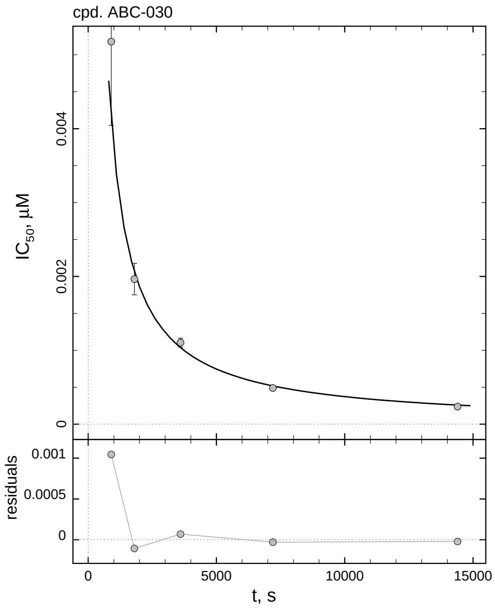

| $i$ | $t$, min | $I_{50}$, nM | ± | std.err. | CV, % | $k_1^*$, $mM^{-1}s^{-1}$ |
|---|---|---|---|---|---|---|
| 1 | 15 | 5.179 | ± | 1.135 | 21.9 | 341.9 |
| 2 | 30 | 1.964 | ± | 0.2126 | 10.8 | 450.8 |
| 3 | 60 | 1.105 | ± | 0.05994 | 5.4 | 400.7 |
| 4 | 120 | 0.4890 | ± | 0.01497 | 3.1 | 452.7 |
| 5 | 240 | 0.2376 | ± | 0.003710 | 1.6 | 465.9 |

*Cpd.* **30***: Mechanistic analysis*

| parameter | unit | "true" (T) | calculated (C) | C/T ratio | note |
|---|---|---|---|---|---|
| mechanism | | **C2S** | **C1** | | |
| $k_{\text{eff}}$ | $mM^{-1}s^{-1}$ | 909.1 | 905.3 | 1.00 | $= k_1^* \, (1 + [S]_0/K_M)$ |

| | |
|---|---|
| Reaction times used for analysis: | 30 and 120 min |
| Maximum GSD for accepting one-step model **C1**: | 1.25 |
| Observed GSD: | 1.00 |
| Assumed $[S]_0/K_M$ ratio: | 1.0 |

*Cpd.* **31***: Simulated data*

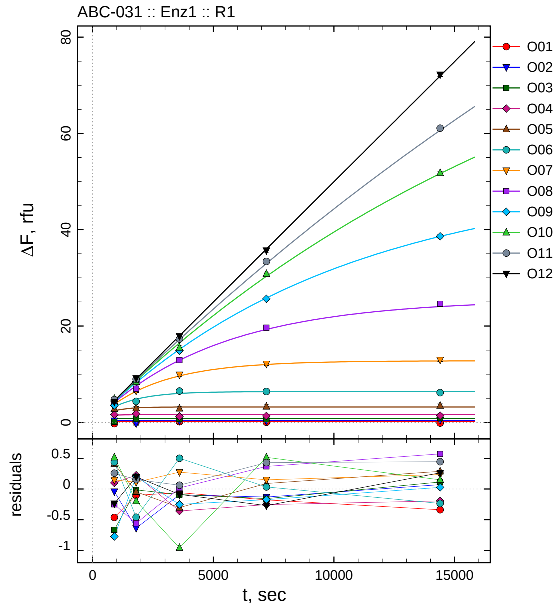

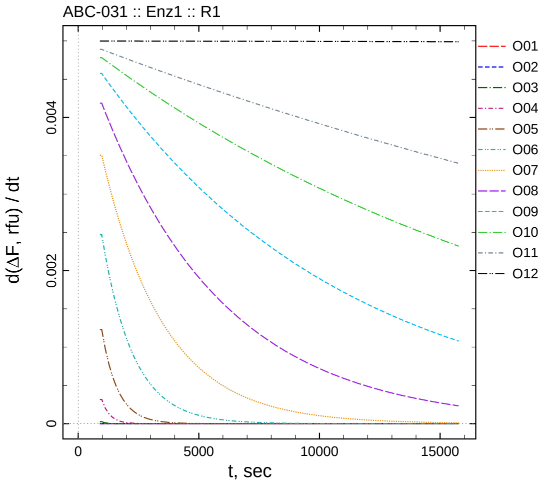

| $[I]_0$, nM | t = 15 | 30 | 60 | 120 | 240 min |
|---|---|---|---|---|---|
| 550 | -0.264 | 0.098 | 0.140 | 0.022 | -0.139 |
| 275 | 0.359 | -0.236 | 0.306 | 0.271 | 0.471 |
| 137.5 | 0.130 | 0.785 | 0.711 | 0.644 | 0.915 |
| 68.75 | 1.594 | 1.818 | 1.242 | 1.347 | 1.406 |
| 34.375 | 2.809 | 2.955 | 2.885 | 3.290 | 3.488 |
| 17.1875 | 3.653 | 4.356 | 6.509 | 6.410 | 6.165 |
| 8.59375 | 3.931 | 6.558 | 9.920 | 12.170 | 12.985 |
| 4.29688 | 3.870 | 7.014 | 12.925 | 19.672 | 24.616 |
| 2.14844 | 3.530 | 8.446 | 14.912 | 25.655 | 38.640 |
| 1.07422 | 4.917 | 8.415 | 15.538 | 30.844 | 51.804 |
| 0.537109 | 4.707 | 8.956 | 17.286 | 33.422 | 61.089 |
| 0 | 4.262 | 9.197 | 17.908 | 35.713 | 72.198 |

*Cpd.* **31***: Determination of* $IC_{50}$ *and* $k_1^*$

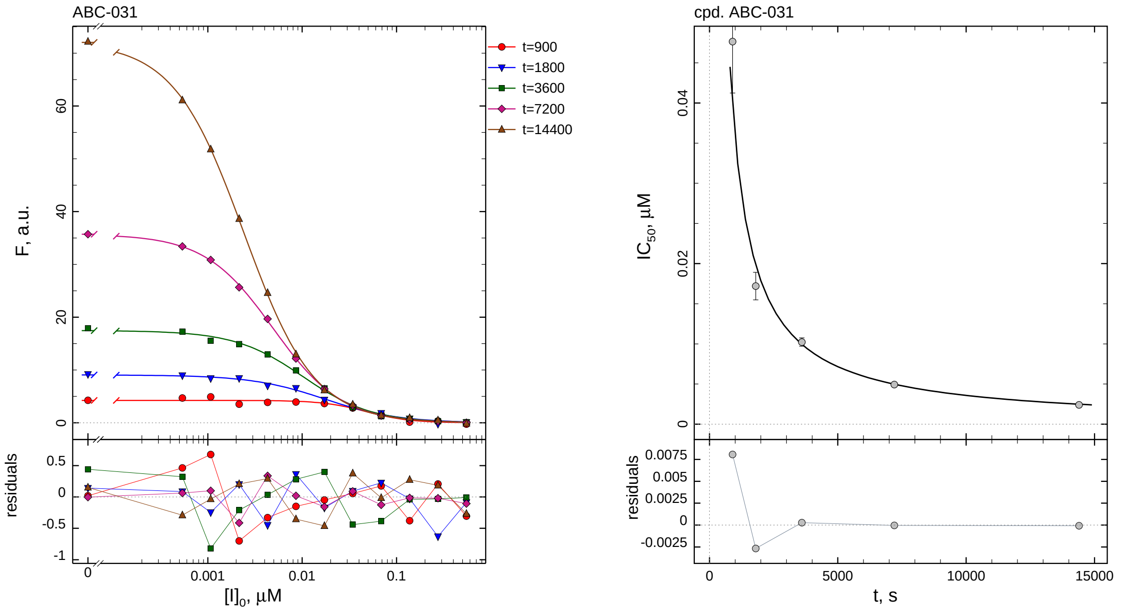

| $i$ | $t$, min | $I_{50}$, nM | ± | std.err. | CV, % | $k_1^*$, $mM^{-1}s^{-1}$ |
|---|---|---|---|---|---|---|
| 1 | 15 | 47.65 | ± | 6.411 | 13.5 | 37.16 |
| 2 | 30 | 17.20 | ± | 1.713 | 10.0 | 51.48 |
| 3 | 60 | 10.23 | ± | 0.5188 | 5.1 | 43.28 |
| 4 | 120 | 4.937 | ± | 0.1301 | 2.6 | 44.84 |
| 5 | 240 | 2.410 | ± | 0.03299 | 1.4 | 45.92 |

*Cpd.* **31***: Mechanistic analysis*

| parameter | unit | "true" (T) | calculated (C) | C/T ratio | note |
|---|---|---|---|---|---|
| mechanism | | **C2S** | **C1** | | |
| $k_{eff}$ | $mM^{-1}s^{-1}$ | 90.91 | 89.67 | 0.99 | $= k_1^* \, (1 + [S]_0/K_M)$ |

| | |
|---|---|
| Reaction times used for analysis: | 30 and 120 min |
| Maximum GSD for accepting one-step model **C1**: | 1.25 |
| Observed GSD: | 1.10 |
| Assumed $[S]_0/K_M$ ratio: | 1.0 |

*2.32. Compound No. 32*

*Cpd.* ***32****: Simulated data*

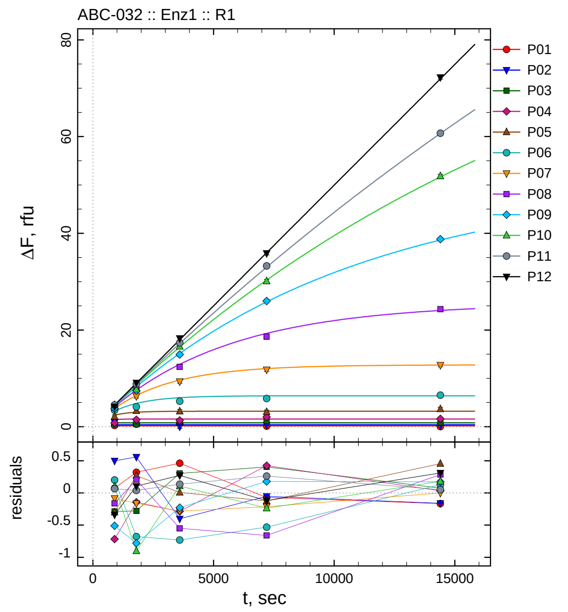

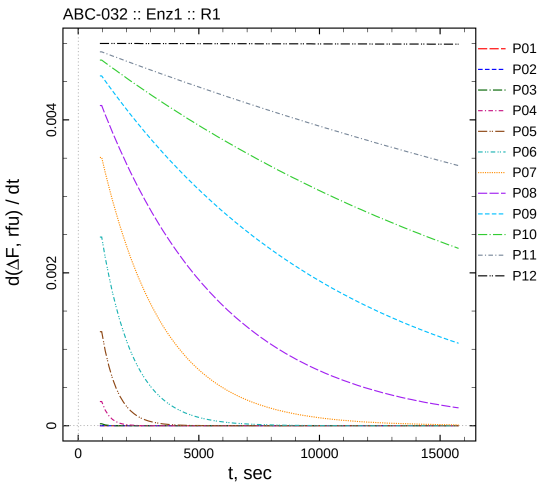

| $[I]_0$, nM | t = 15 | 30 | 60 | 120 | 240 min |
|---:|---:|---:|---:|---:|---:|
| 5500 | 0.273 | 0.522 | 0.662 | 0.130 | 0.034 |
| 2750 | 0.899 | 0.958 | -0.001 | 0.348 | 0.239 |
| 1375 | 0.506 | 0.522 | 1.104 | 1.205 | 0.875 |
| 687.5 | 0.778 | 1.449 | 1.313 | 2.024 | 1.634 |
| 343.75 | 2.099 | 3.268 | 3.194 | 3.076 | 3.654 |
| 171.875 | 3.416 | 4.138 | 5.274 | 5.843 | 6.527 |
| 85.9375 | 3.699 | 6.283 | 9.364 | 11.806 | 12.749 |
| 42.9688 | 3.958 | 7.782 | 12.359 | 18.642 | 24.328 |
| 21.4844 | 3.792 | 7.464 | 14.932 | 26.005 | 38.782 |
| 10.7422 | 4.499 | 7.708 | 16.605 | 30.090 | 51.830 |
| 5.37109 | 4.515 | 8.840 | 17.359 | 33.253 | 60.694 |
| 0 | 4.160 | 9.102 | 18.268 | 35.864 | 72.245 |

#### *Cpd.* **32***: Determination of* $IC_{50}$ *and* $k_1^*$

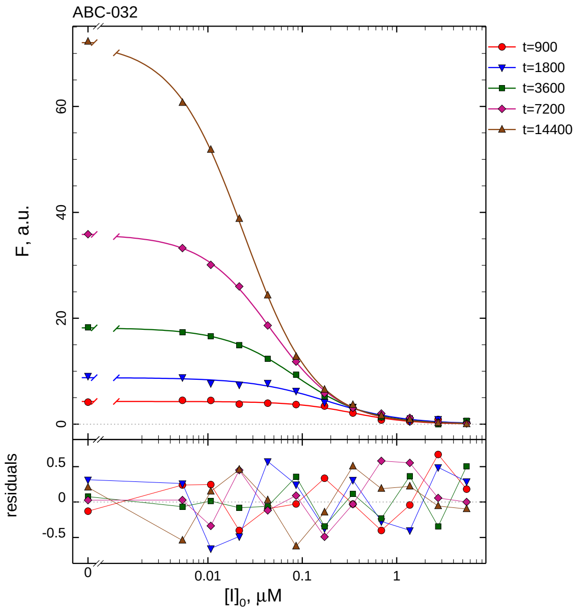

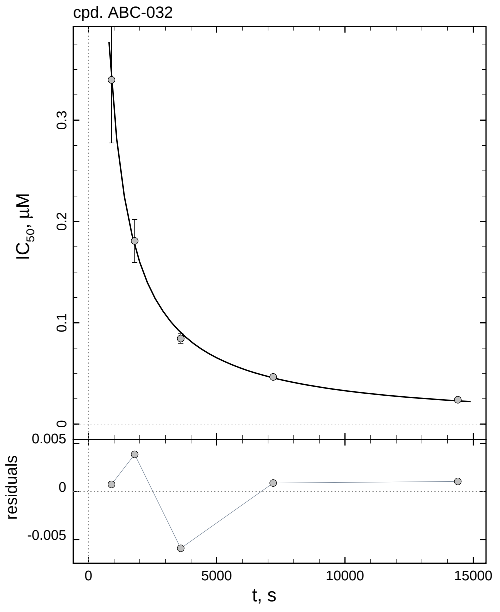

| $i$ | $t$, min | $I_{50}$, nM | ± | std.err. | CV, % | $k_1^*$, $mM^{-1}s^{-1}$ |
|---|---|---|---|---|---|---|
| 1 | 15 | 339.8 | ± | 62.31 | 18.3 | 5.212 |
| 2 | 30 | 180.8 | ± | 21.23 | 11.7 | 4.898 |
| 3 | 60 | 84.46 | ± | 4.796 | 5.7 | 5.241 |
| 4 | 120 | 46.55 | ± | 1.358 | 2.9 | 4.755 |
| 5 | 240 | 24.00 | ± | 0.3608 | 1.5 | 4.612 |

#### *Cpd.* **32***: Mechanistic analysis*

| parameter | unit | "true" (T) | calculated (C) | C/T ratio | note |
|---|---|---|---|---|---|
| mechanism | | **C2S** | **C1** | | |
| $k_{eff}$ | $mM^{-1}s^{-1}$ | 9.091 | 9.510 | 1.05 | $= k_1^* \, (1 + [S]_0/K_M)$ |

| | |
|---|---|
| Reaction times used for analysis: | 30 and 120 min |
| Maximum GSD for accepting one-step model **C1**: | 1.25 |
| Observed GSD: | 1.02 |
| Assumed $[S]_0/K_M$ ratio: | 1.0 |

*2.33. Compound No. 33*

*Cpd.* **33***: Simulated data*

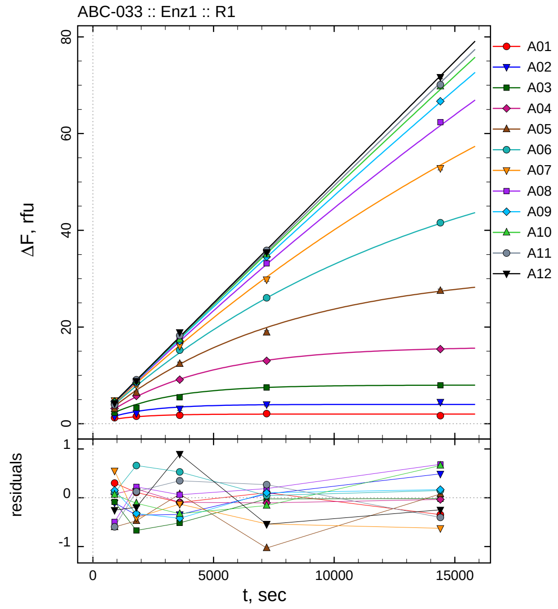

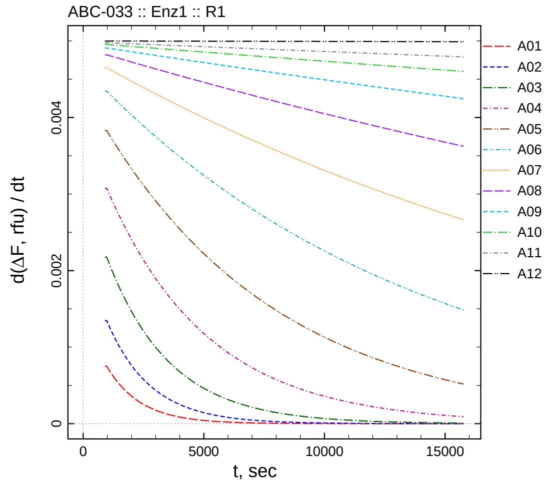

| $[I]_0$, nM | t = 15 | 30 | 60 | 120 | 240 min |
|---|---|---|---|---|---|
| 500.5 | 1.249 | 1.555 | 1.752 | 2.093 | 1.654 |
| 250.25 | 1.471 | 2.175 | 3.117 | 4.003 | 4.488 |
| 125.125 | 2.256 | 3.327 | 5.481 | 7.473 | 7.949 |
| 62.5625 | 3.163 | 5.759 | 9.117 | 13.011 | 15.441 |
| 31.2812 | 3.064 | 6.437 | 12.402 | 18.871 | 27.492 |
| 15.6406 | 4.142 | 8.486 | 15.223 | 26.052 | 41.577 |
| 7.82031 | 4.811 | 7.964 | 16.070 | 29.805 | 52.842 |
| 3.91016 | 3.880 | 8.899 | 17.118 | 33.157 | 62.349 |
| 1.95508 | 4.587 | 8.513 | 17.093 | 34.524 | 66.669 |
| 0.977539 | 4.531 | 8.802 | 17.425 | 35.030 | 69.799 |
| 0.48877 | 3.887 | 9.084 | 18.222 | 35.851 | 70.115 |
| 0 | 4.246 | 8.804 | 18.890 | 35.440 | 71.688 |

*Cpd.* **33***: Determination of* $IC_{50}$ *and* $k_1^*$

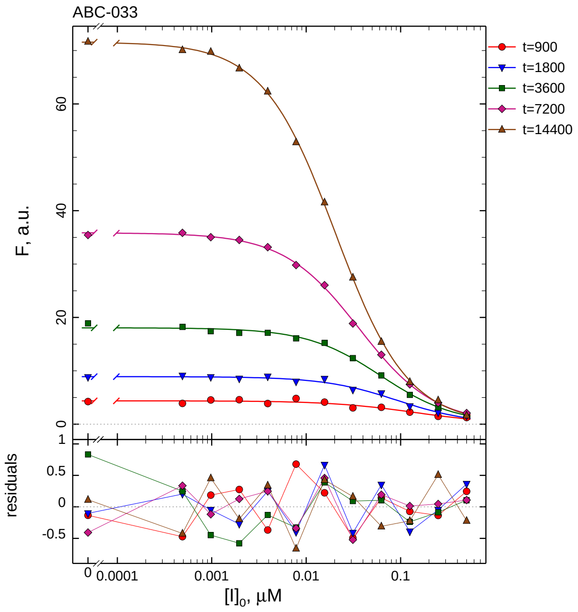

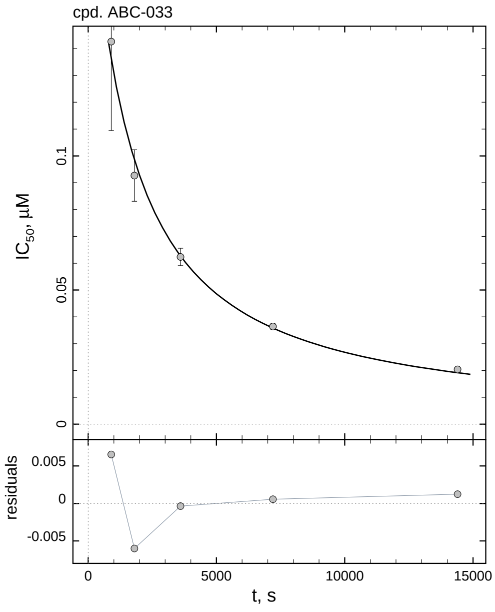

| $i$ | $t$, min | $I_{50}$, nM ± std.err. | CV, % | $k_1^*$, $mM^{-1}s^{-1}$ |
|---|---|---|---|---|
| 1 | 15 | 142.6 ± 33.16 | 23.3 | 12.42 |
| 2 | 30 | 92.68 ± 9.604 | 10.4 | 9.552 |
| 3 | 60 | 62.34 ± 3.256 | 5.2 | 7.101 |
| 4 | 120 | 36.41 ± 1.006 | 2.8 | 6.080 |
| 5 | 240 | 20.43 ± 0.2839 | 1.4 | 5.418 |

*Cpd.* **33***: Mechanistic analysis*

| parameter | unit | "true" (T) | calculated (C) | C/T ratio | note |
|---|---|---|---|---|---|
| mechanism | | **C2S** | **C2F** | | |
| $k_{eff}$ | $mM^{-1}s^{-1}$ | 9.990 | 10.45 | 1.05 | $= k_{inact}/K_i$ |
| $K_i$ | nM | 100.1 | 95.60 | 0.96 | |
| $k_{inact}$ | $s^{-1}$ | 0.001 | 0.0009993 | 1.00 | |

| | |
|---|---|
| Reaction times used for analysis: | 30 and 120 min |
| Maximum GSD for accepting one-step model **C1**: | 1.25 |
| Observed GSD: | 1.38 |
| Assumed $[S]_0/K_M$ ratio: | 1.0 |

*2.34. Compound No. 34*

*Cpd.* **34***: Simulated data*

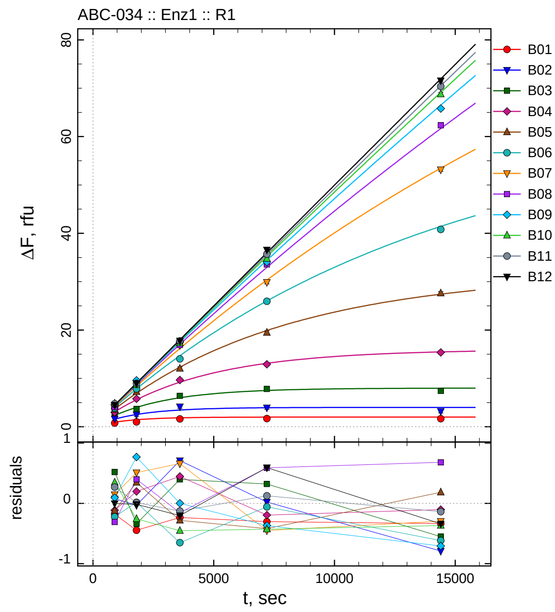

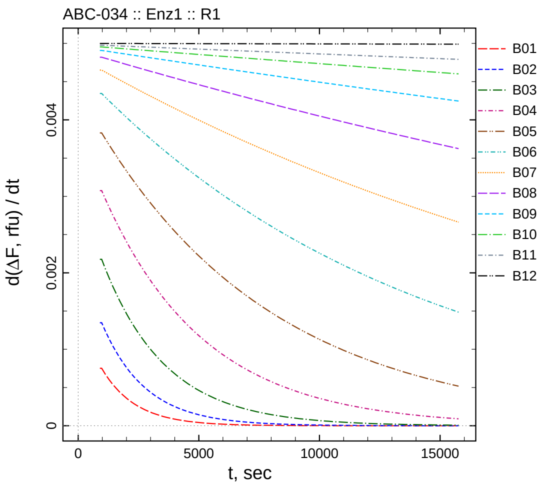

| $[I]_0$, nM | t = 15 | 30 | 60 | 120 | 240 min |
|---|---|---|---|---|---|
| 5005 | 0.770 | 1.001 | 1.613 | 1.686 | 1.665 |
| 2502.5 | 1.652 | 2.498 | 4.168 | 3.948 | 3.209 |
| 1251.25 | 2.861 | 3.660 | 6.394 | 7.818 | 7.416 |
| 625.625 | 2.972 | 5.774 | 9.655 | 12.921 | 15.371 |
| 312.812 | 3.546 | 7.255 | 12.043 | 19.479 | 27.597 |
| 156.406 | 3.819 | 7.842 | 14.042 | 25.950 | 40.811 |
| 78.2031 | 4.404 | 8.882 | 16.858 | 29.892 | 53.170 |
| 39.1016 | 4.067 | 9.075 | 16.901 | 33.558 | 62.345 |
| 19.5508 | 4.530 | 9.601 | 17.515 | 34.055 | 65.796 |
| 9.77539 | 4.815 | 8.657 | 17.299 | 34.761 | 68.771 |
| 4.8877 | 4.752 | 8.969 | 17.745 | 35.707 | 70.379 |
| 0 | 4.495 | 8.982 | 17.793 | 36.575 | 71.595 |

*Cpd.* **34**: *Determination of $IC_{50}$ and $k_1^*$*

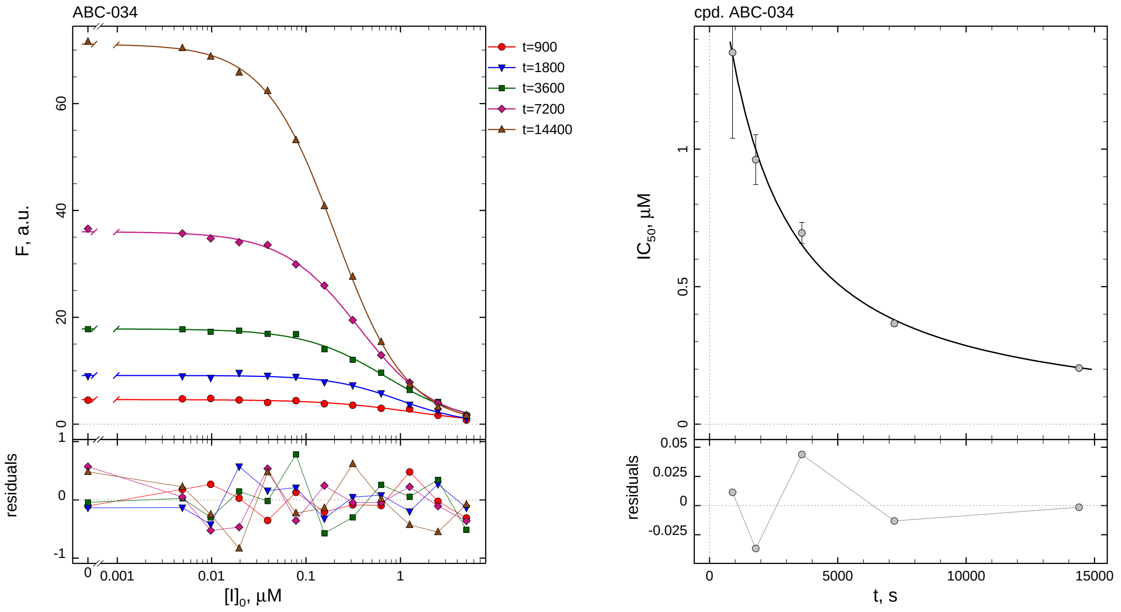

| $i$ | $t$, min | $I_{50}$, nM ± std.err. | CV, % | $k_1^*$, mM$^{-1}$s$^{-1}$ |
|---|---|---|---|---|
| 1 | 15 | 1351.6 ± 311.5 | 23.0 | 1.310 |
| 2 | 30 | 962.2 ± 90.75 | 9.4 | 0.9201 |
| 3 | 60 | 695.5 ± 38.50 | 5.5 | 0.6364 |
| 4 | 120 | 366.4 ± 9.944 | 2.7 | 0.6041 |
| 5 | 240 | 204.0 ± 2.753 | 1.3 | 0.5425 |

*Cpd.* **34**: *Mechanistic analysis*

| parameter | unit | "true" (T) | calculated (C) | C/T ratio | note |
|---|---|---|---|---|---|
| mechanism | | **C2S** | **C2F** | | |
| $k_{\text{eff}}$ | mM$^{-1}$s$^{-1}$ | 0.9990 | 1.057 | 1.06 | $= k_{\text{inact}}/K_{\text{i}}$ |
| $K_{\text{i}}$ | nM | 1001 | 1050.6 | 1.05 | |
| $k_{\text{inact}}$ | s$^{-1}$ | 0.001 | 0.001110 | 1.11 | |

| | |
|---|---|
| Reaction times used for analysis: | 30 and 120 min |
| Maximum GSD for accepting one-step model **C1**: | 1.25 |
| Observed GSD: | 1.35 |
| Assumed $[S]_0/K_M$ ratio: | 1.0 |

*2.35. Compound No. 35*

*Cpd.* ***35****: Simulated data*

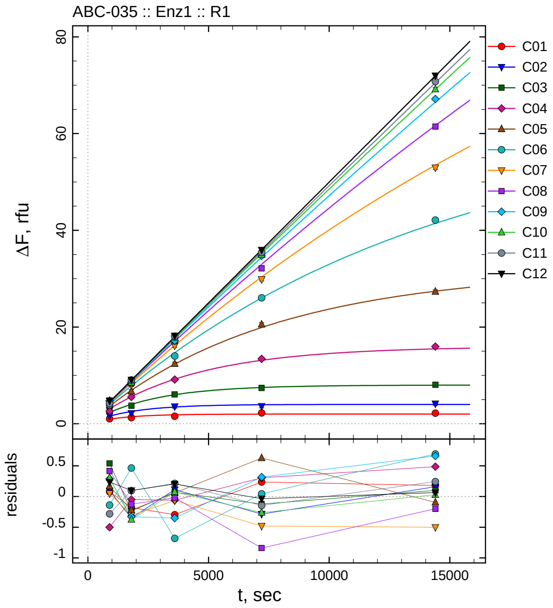

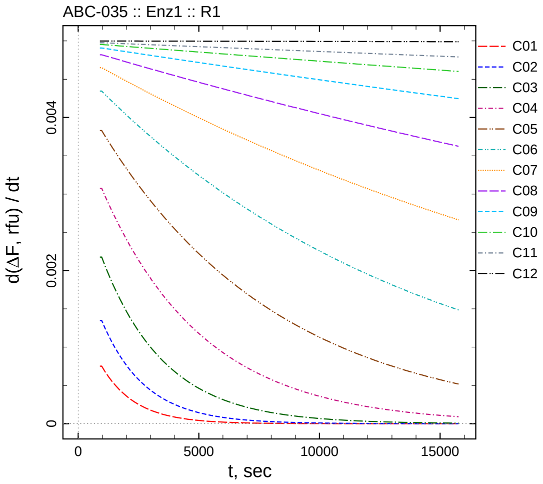

| $[I]_0$, nM | t = 15 | 30 | 60 | 120 | 240 min |
|---|---|---|---|---|---|
| 50050 | 1.027 | 1.269 | 1.551 | 2.225 | 2.180 |
| 25025 | 1.669 | 2.178 | 3.571 | 3.641 | 4.158 |
| 12512.5 | 2.881 | 3.761 | 6.066 | 7.374 | 8.067 |
| 6256.25 | 2.587 | 5.528 | 9.148 | 13.421 | 15.964 |
| 3128.12 | 3.833 | 6.687 | 12.384 | 20.526 | 27.320 |
| 1564.06 | 3.900 | 8.289 | 14.011 | 26.051 | 42.121 |
| 782.031 | 4.308 | 8.068 | 16.134 | 29.862 | 52.966 |
| 391.016 | 4.790 | 8.550 | 17.036 | 32.130 | 61.465 |
| 195.508 | 4.724 | 8.503 | 17.163 | 34.739 | 67.164 |
| 97.7539 | 4.774 | 8.539 | 17.828 | 34.928 | 69.168 |
| 48.877 | 4.202 | 9.052 | 18.087 | 35.437 | 70.759 |
| 0 | 4.735 | 9.097 | 18.200 | 35.948 | 71.999 |

*Cpd.* **35***: Determination of* $IC_{50}$ *and* $k_1^*$

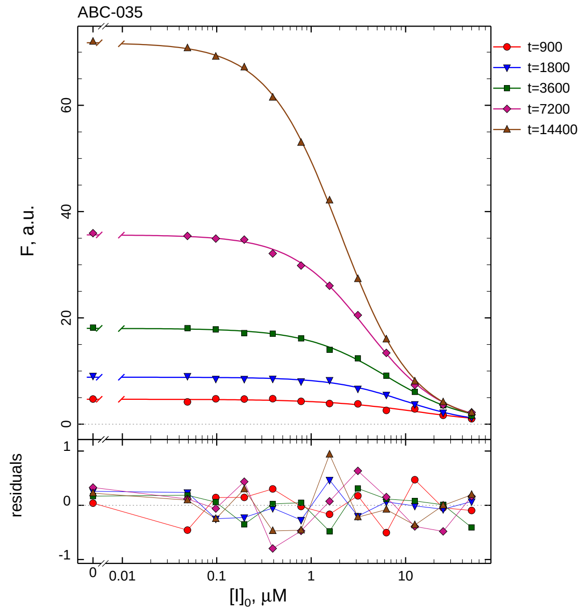

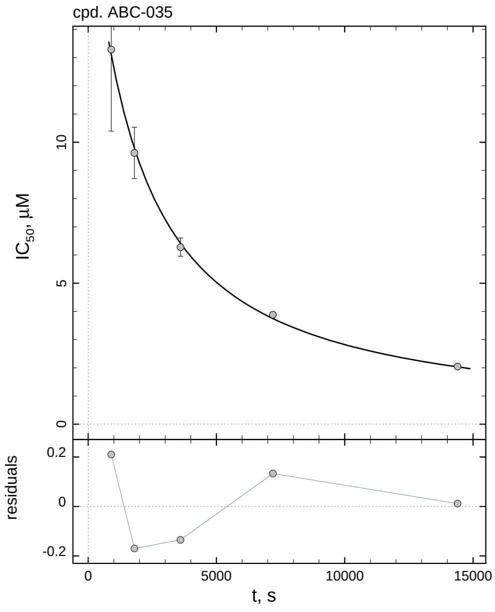

| $i$ | $t$, min | $I_{50}$, nM | ± | std.err. | CV, % | $k_1^*$, $mM^{-1}s^{-1}$ |
|---|---|---|---|---|---|---|
| 1 | 15 | 13288 | ± | 2896.6 | 21.8 | 0.1333 |
| 2 | 30 | 9620.0 | ± | 909.8 | 9.5 | 0.09203 |
| 3 | 60 | 6278.3 | ± | 323.2 | 5.1 | 0.07051 |
| 4 | 120 | 3879.4 | ± | 97.79 | 2.5 | 0.05705 |
| 5 | 240 | 2043.2 | ± | 26.32 | 1.3 | 0.05416 |

*Cpd.* **35***: Mechanistic analysis*

| parameter | unit | "true" (T) | calculated (C) | C/T ratio | note |
|---|---|---|---|---|---|
| mechanism | | **C2S** | **C2F** | | |
| $k_{eff}$ | $mM^{-1}s^{-1}$ | 0.09990 | 0.09659 | 0.97 | $= k_{inact}/K_i$ |
| $K_i$ | nM | 10010 | 9491.8 | 0.95 | |
| $k_{inact}$ | $s^{-1}$ | 0.001 | 0.0009168 | 0.92 | |

| | |
|---|---|
| Reaction times used for analysis: | 30 and 120 min |
| Maximum GSD for accepting one-step model **C1**: | 1.25 |
| Observed GSD: | 1.40 |
| Assumed $[S]_0/K_M$ ratio: | 1.0 |

#### *2.36. Compound No. 36*

*Cpd.* **36***: Simulated data*

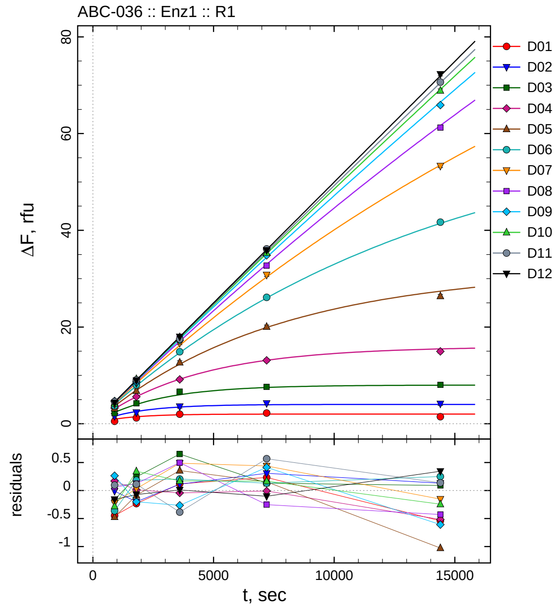

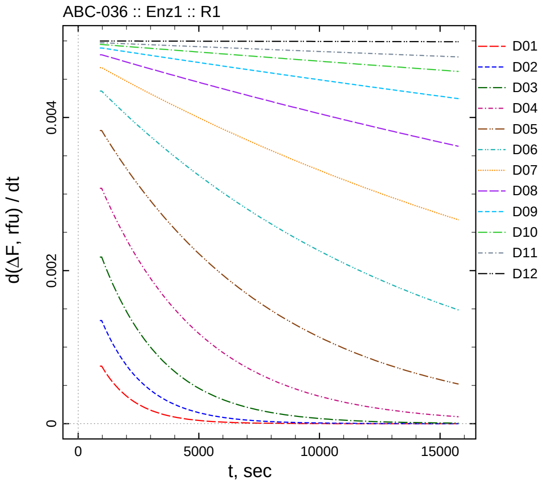

| $[I]_0$, nM | t = 15 | 30 | 60 | 120 | 240 min |
|---|---|---|---|---|---|
| 500500 | 0.501 | 1.214 | 1.966 | 2.222 | 1.460 |
| 250250 | 1.565 | 2.328 | 3.571 | 4.234 | 4.129 |
| 125125 | 2.182 | 4.240 | 6.648 | 7.629 | 8.057 |
| 62562.5 | 3.256 | 5.589 | 9.166 | 13.108 | 14.959 |
| 31281.3 | 3.192 | 6.826 | 12.683 | 20.054 | 26.389 |
| 15640.6 | 3.678 | 8.009 | 14.898 | 26.138 | 41.679 |
| 7820.31 | 4.035 | 8.406 | 16.687 | 30.778 | 53.312 |
| 3910.16 | 4.437 | 8.798 | 17.555 | 32.716 | 61.236 |
| 1955.08 | 4.700 | 8.634 | 17.249 | 34.836 | 65.894 |
| 977.539 | 4.189 | 9.259 | 17.925 | 35.357 | 68.895 |
| 488.77 | 4.579 | 9.069 | 17.488 | 36.151 | 70.652 |
| 0 | 4.339 | 8.926 | 18.006 | 35.881 | 72.279 |

*Cpd.* **36***: Determination of* $IC_{50}$ *and* $k_1^*$

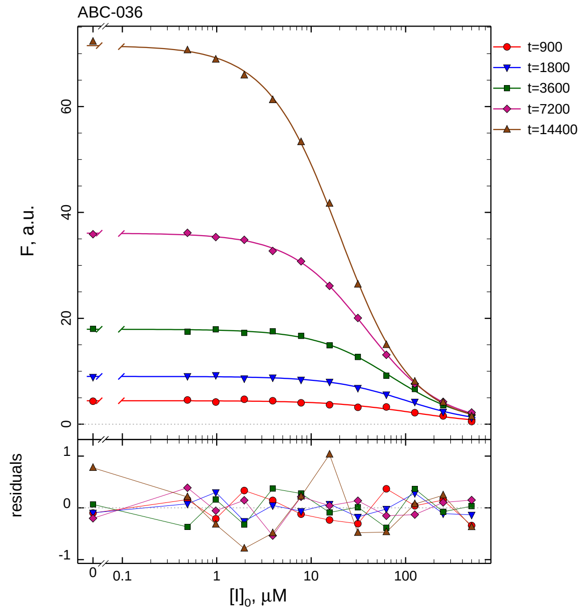

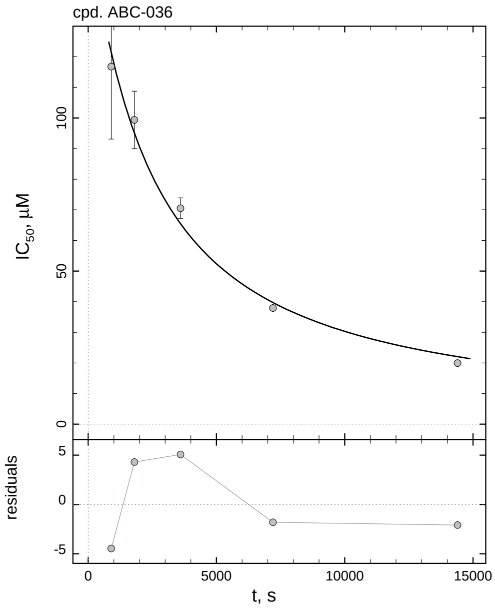

| $i$ | $t$, min | $I_{50}$, nM | ± | std.err. | CV, % | $k_1^*$, mM$^{-1}$s$^{-1}$ |
|---|---|---|---|---|---|---|
| 1 | 15 | 116765 | ± | 23663 | 20.3 | 0.01516 |
| 2 | 30 | 99372 | ± | 9370.1 | 9.4 | 0.008909 |
| 3 | 60 | 70516 | ± | 3386.9 | 4.8 | 0.006278 |
| 4 | 120 | 37924 | ± | 935.5 | 2.5 | 0.005836 |
| 5 | 240 | 19962 | ± | 251.1 | 1.3 | 0.005544 |

*Cpd.* **36***: Mechanistic analysis*

| parameter | unit | "true" (T) | calculated (C) | C/T ratio | note |
|---|---|---|---|---|---|
| mechanism | | **C2S** | **C2F** | | |
| $k_{\text{eff}}$ | mM$^{-1}$s$^{-1}$ | 0.009990 | 0.01020 | 1.02 | $= k_{\text{inact}}/K_{\text{i}}$ |
| $K_{\text{i}}$ | nM | 100100 | 108037 | 1.08 | |
| $k_{\text{inact}}$ | s$^{-1}$ | 0.001 | 0.001102 | 1.10 | |

| | |
|---|---|
| Reaction times used for analysis: | 30 and 120 min |
| Maximum GSD for accepting one-step model **C1**: | 1.25 |
| Observed GSD: | 1.35 |
| Assumed $[S]_0/K_M$ ratio: | 1.0 |

*2.37. Compound No. 37*

*Cpd.* ***37****: Simulated data*

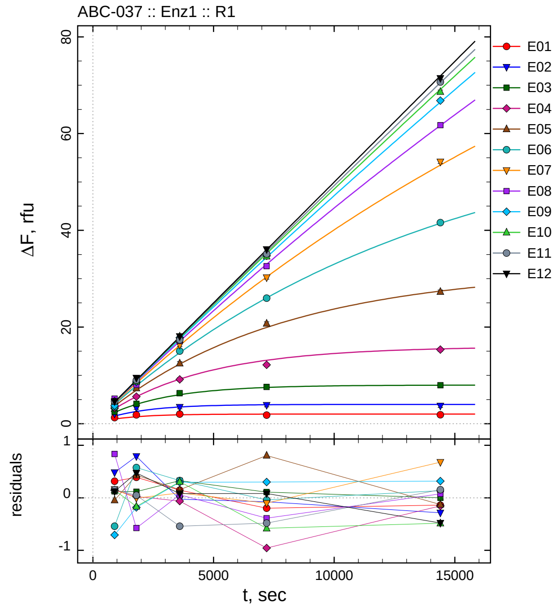

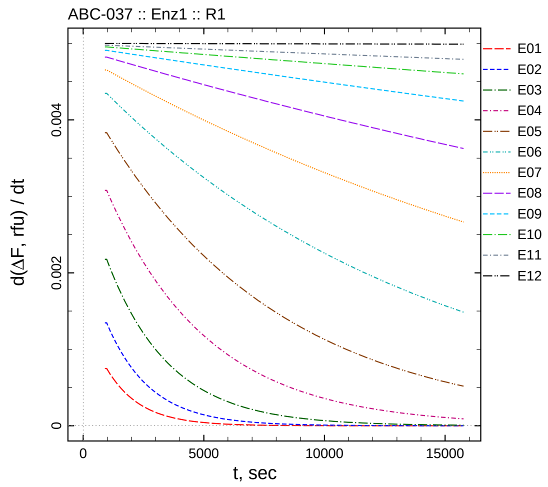

| $[I]_0$, nM | t = 15 | 30 | 60 | 120 | 240 min |
|---|---|---|---|---|---|
| 50.5 | 1.271 | 1.842 | 1.995 | 1.791 | 1.862 |
| 25.25 | 2.067 | 3.328 | 3.436 | 3.853 | 3.713 |
| 12.625 | 2.474 | 4.125 | 6.333 | 7.610 | 7.970 |
| 6.3125 | 3.241 | 5.614 | 9.160 | 12.170 | 15.327 |
| 3.15625 | 3.629 | 7.351 | 12.494 | 20.720 | 27.300 |
| 1.57813 | 3.504 | 8.408 | 15.032 | 25.980 | 41.585 |
| 0.789063 | 4.421 | 8.373 | 16.298 | 30.250 | 54.164 |
| 0.394531 | 5.210 | 8.102 | 17.118 | 32.586 | 61.752 |
| 0.197266 | 3.729 | 8.652 | 17.814 | 34.726 | 66.828 |
| 0.0986328 | 4.614 | 8.751 | 18.060 | 34.611 | 68.659 |
| 0.0493164 | 4.638 | 9.005 | 17.333 | 35.101 | 70.673 |
| 0 | 4.621 | 9.471 | 18.075 | 36.066 | 71.455 |

*Cpd.* **37***: Determination of* $IC_{50}$ *and* $k_1^*$

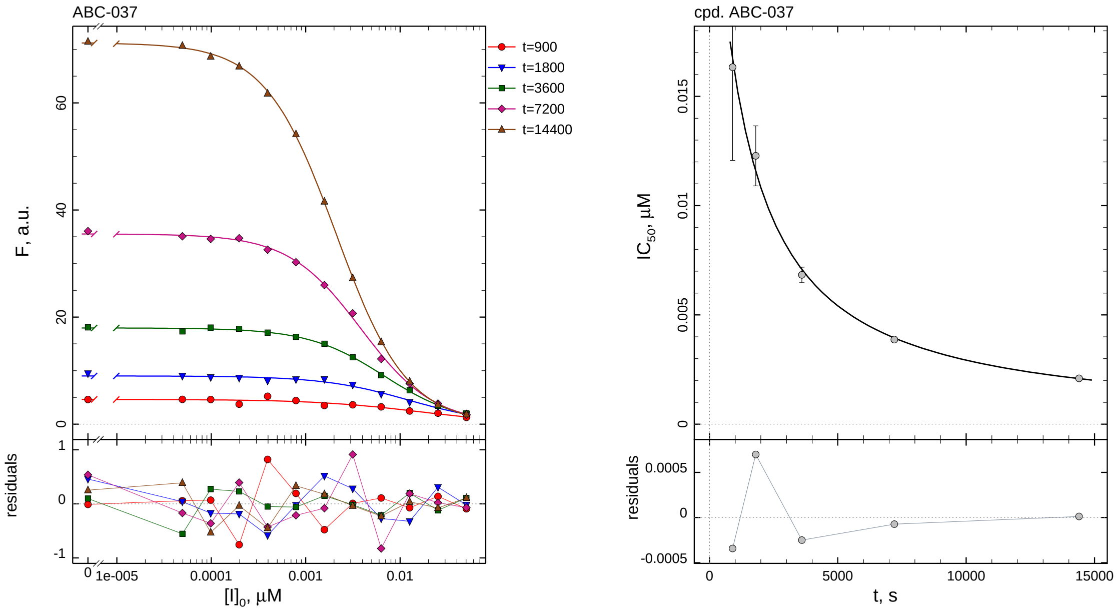

| $i$ | $t$, min | $I_{50}$, nM ± std.err. | CV, % | $k_1^*$, mM$^{-1}$s$^{-1}$ |
|---|---|---|---|---|
| 1 | 15 | 16.34 ± 4.262 | 26.1 | 108.4 |
| 2 | 30 | 12.28 ± 1.373 | 11.2 | 72.10 |
| 3 | 60 | 6.833 ± 0.3555 | 5.2 | 64.79 |
| 4 | 120 | 3.874 ± 0.1010 | 2.6 | 57.13 |
| 5 | 240 | 2.097 ± 0.02786 | 1.3 | 52.77 |

*Cpd.* **37***: Mechanistic analysis*

| parameter | unit | "true" (T) | calculated (C) | C/T ratio | note |
|---|---|---|---|---|---|
| mechanism | | **C2S** | **C1** | | |
| $k_{eff}$ | mM$^{-1}$s$^{-1}$ | 99.01 | 114.3 | 1.15 | $= k_1^* \, (1 + [S]_0/K_M)$ |

| | |
|---|---|
| Reaction times used for analysis: | 30 and 120 min |
| Maximum GSD for accepting one-step model **C1**: | 1.25 |
| Observed GSD: | 1.18 |
| Assumed $[S]_0/K_M$ ratio: | 1.0 |

*2.38. Compound No. 38*

*Cpd.* **38***: Simulated data*

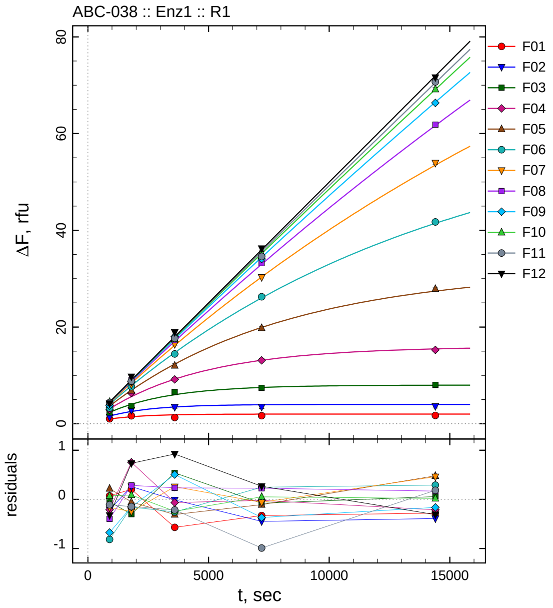

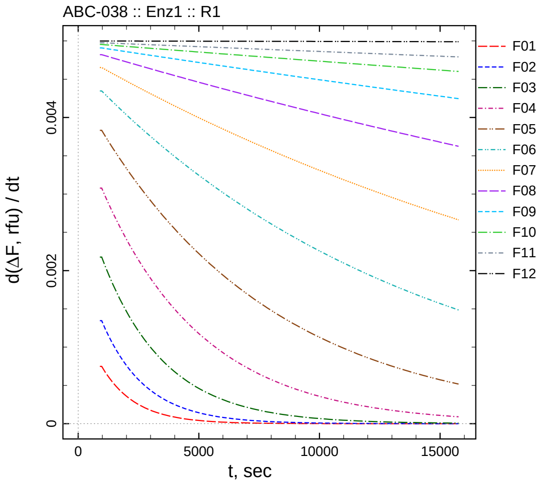

| $[I]_0$, nM | t = 15 | 30 | 60 | 120 | 240 min |
|---|---|---|---|---|---|
| 505 | 1.027 | 1.653 | 1.279 | 1.659 | 1.720 |
| 252.5 | 1.373 | 2.796 | 3.451 | 3.479 | 3.608 |
| 126.25 | 2.304 | 3.706 | 6.543 | 7.421 | 8.024 |
| 63.125 | 2.886 | 6.343 | 9.156 | 13.105 | 15.267 |
| 31.5625 | 3.893 | 6.867 | 12.030 | 19.805 | 27.903 |
| 15.7813 | 3.230 | 7.676 | 14.446 | 26.267 | 41.733 |
| 7.89063 | 4.094 | 8.129 | 16.461 | 30.290 | 53.941 |
| 3.94531 | 3.979 | 8.962 | 17.292 | 33.202 | 61.843 |
| 1.97266 | 3.761 | 8.680 | 18.020 | 34.058 | 66.346 |
| 0.986328 | 4.537 | 9.005 | 17.509 | 35.243 | 69.159 |
| 0.493164 | 4.373 | 8.809 | 17.658 | 34.594 | 70.701 |
| 0 | 4.174 | 9.731 | 18.916 | 36.248 | 71.623 |

*Cpd.* **38***: Determination of* $IC_{50}$ *and* $k_1^*$

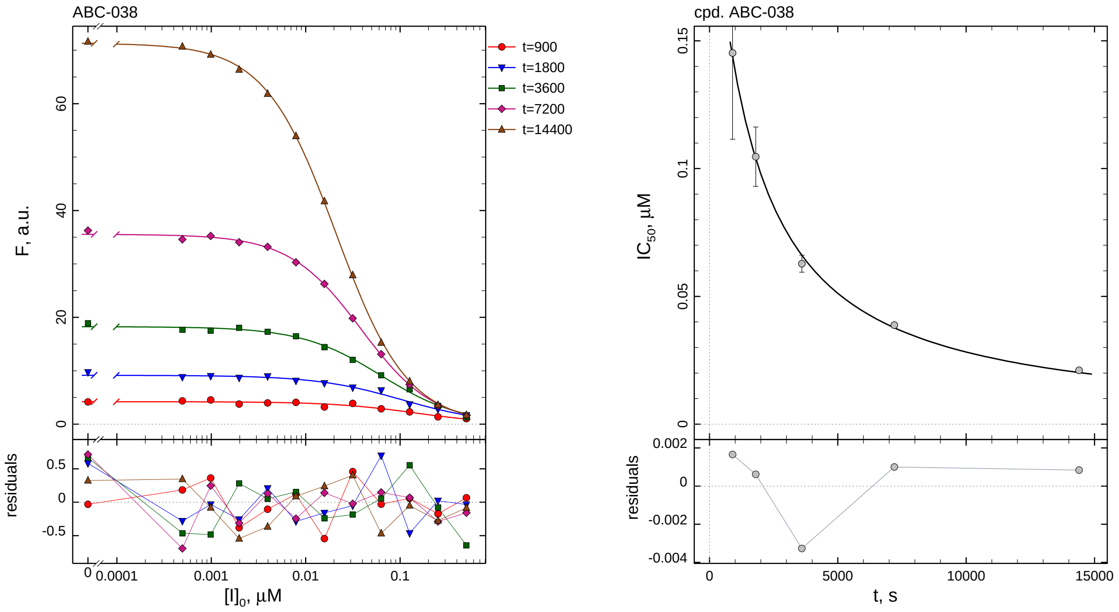

| $i$ | $t$, min | $I_{50}$, nM | ± | std.err. | CV, % | $k_1^*$, $mM^{-1}s^{-1}$ |
|---|---|---|---|---|---|---|
| 1 | 15 | 145.2 | ± | 33.76 | 23.2 | 12.19 |
| 2 | 30 | 104.6 | ± | 11.62 | 11.1 | 8.460 |
| 3 | 60 | 62.78 | ± | 3.342 | 5.3 | 7.051 |
| 4 | 120 | 38.74 | ± | 0.9986 | 2.6 | 5.713 |
| 5 | 240 | 21.05 | ± | 0.2797 | 1.3 | 5.258 |

*Cpd.* **38***: Mechanistic analysis*

| parameter | unit | "true" (T) | calculated (C) | C/T ratio | note |
|---|---|---|---|---|---|
| mechanism | | **C2S** | **C2F** | | |
| $k_{\text{eff}}$ | $mM^{-1}s^{-1}$ | 9.901 | 10.14 | 1.02 | $= k_{\text{inact}}/K_{\text{i}}$ |
| $K_{\text{i}}$ | nM | 101 | 120.9 | 1.20 | |
| $k_{\text{inact}}$ | $s^{-1}$ | 0.001 | 0.001226 | 1.23 | |

| | |
|---|---|
| Reaction times used for analysis: | 30 and 120 min |
| Maximum GSD for accepting one-step model **C1**: | 1.25 |
| Observed GSD: | 1.32 |
| Assumed $[S]_0/K_M$ ratio: | 1.0 |

#### 2.39. *Compound No. 39*

*Cpd.* **39***: Simulated data*

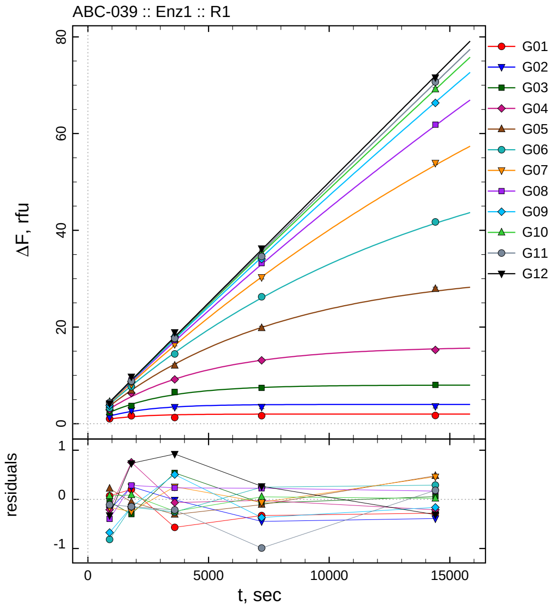

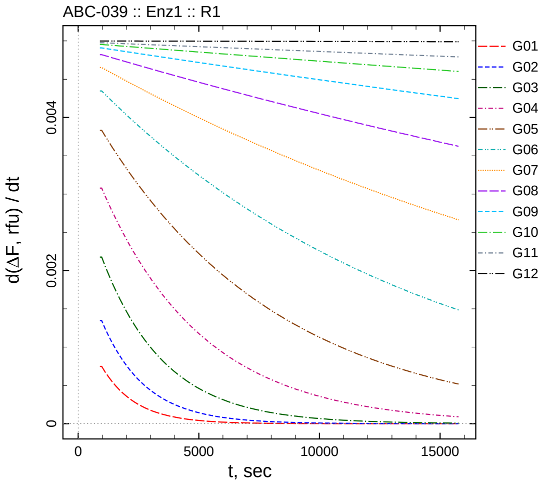

| $[I]_0$, nM | t = 15 | 30 | 60 | 120 | 240 min |
|---:|---:|---:|---:|---:|---:|
| 5050 | 1.027 | 1.653 | 1.279 | 1.659 | 1.720 |
| 2525 | 1.373 | 2.796 | 3.451 | 3.479 | 3.608 |
| 1262.5 | 2.304 | 3.706 | 6.543 | 7.421 | 8.024 |
| 631.25 | 2.886 | 6.343 | 9.156 | 13.105 | 15.267 |
| 315.625 | 3.893 | 6.867 | 12.030 | 19.805 | 27.903 |
| 157.812 | 3.230 | 7.676 | 14.446 | 26.267 | 41.733 |
| 78.9062 | 4.094 | 8.129 | 16.461 | 30.290 | 53.941 |
| 39.4531 | 3.979 | 8.962 | 17.292 | 33.202 | 61.843 |
| 19.7266 | 3.761 | 8.680 | 18.020 | 34.058 | 66.346 |
| 9.86328 | 4.537 | 9.005 | 17.509 | 35.243 | 69.159 |
| 4.93164 | 4.373 | 8.809 | 17.658 | 34.594 | 70.701 |
| 0 | 4.174 | 9.731 | 18.916 | 36.248 | 71.623 |

*Cpd.* **39***: Determination of* $IC_{50}$ *and* $k_1^*$

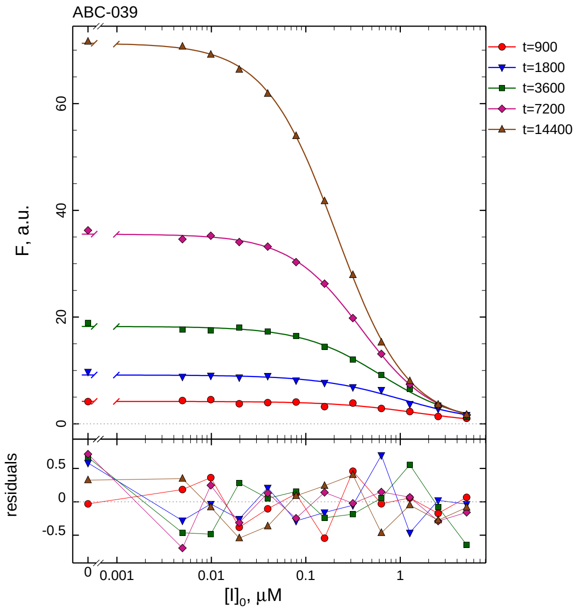

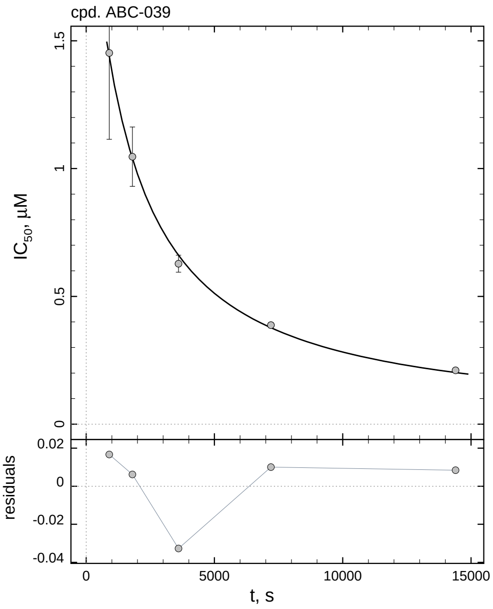

| $i$ | $t$, min | $I_{50}$, nM | ± | std.err. | CV, % | $k_1^*$, mM$^{-1}$s$^{-1}$ |
|---|---|---|---|---|---|---|
| 1 | 15 | 1452.2 | ± | 337.5 | 23.2 | 1.219 |
| 2 | 30 | 1046.5 | ± | 116.2 | 11.1 | 0.8460 |
| 3 | 60 | 627.8 | ± | 33.42 | 5.3 | 0.7051 |
| 4 | 120 | 387.4 | ± | 9.986 | 2.6 | 0.5713 |
| 5 | 240 | 210.5 | ± | 2.797 | 1.3 | 0.5258 |

*Cpd.* **39***: Mechanistic analysis*

| parameter | unit | "true" (T) | calculated (C) | C/T ratio | note |
|---|---|---|---|---|---|
| mechanism | | **C2S** | **C2F** | | |
| $k_{\text{eff}}$ | mM$^{-1}$s$^{-1}$ | 0.9901 | 1.014 | 1.02 | $= k_{\text{inact}}/K_{\text{i}}$ |
| $K_{\text{i}}$ | nM | 1010 | 1208.6 | 1.20 | |
| $k_{\text{inact}}$ | s$^{-1}$ | 0.001 | 0.001226 | 1.23 | |

| | |
|---|---|
| Reaction times used for analysis: | 30 and 120 min |
| Maximum GSD for accepting one-step model **C1**: | 1.25 |
| Observed GSD: | 1.32 |
| Assumed $[S]_0/K_M$ ratio: | 1.0 |

*2.40. Compound No. 40*

*Cpd.* **40***: Simulated data*

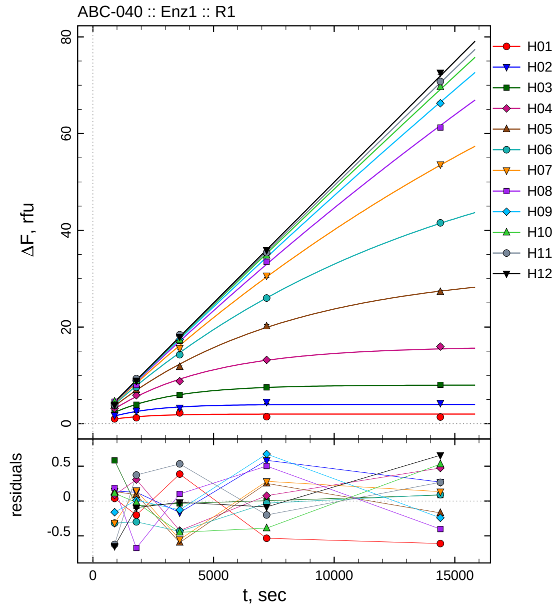

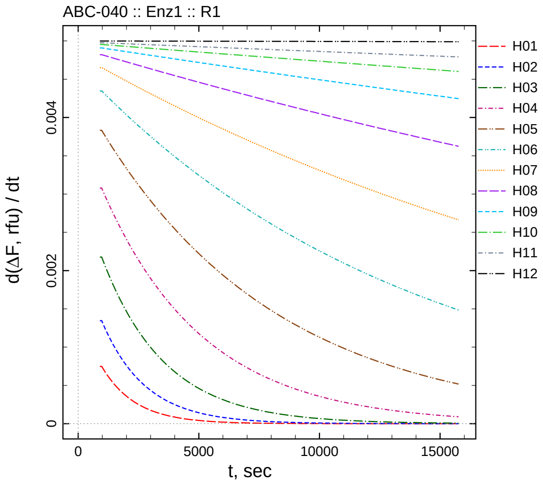

| $[I]_0$, nM | t = 15 | 30 | 60 | 120 | 240 min |
|---|---|---|---|---|---|
| 50500 | 0.992 | 1.249 | 2.236 | 1.454 | 1.389 |
| 25250 | 1.725 | 2.662 | 3.297 | 4.512 | 4.269 |
| 12625 | 2.936 | 3.940 | 5.969 | 7.505 | 8.052 |
| 6312.5 | 3.184 | 5.897 | 8.797 | 13.202 | 15.953 |
| 3156.25 | 3.809 | 7.009 | 11.743 | 20.167 | 27.254 |
| 1578.13 | 3.728 | 7.533 | 14.267 | 25.991 | 41.535 |
| 789.063 | 3.946 | 8.529 | 15.652 | 30.633 | 53.614 |
| 394.531 | 4.563 | 8.003 | 17.162 | 33.477 | 61.271 |
| 197.266 | 4.276 | 8.847 | 17.388 | 35.101 | 66.267 |
| 98.6328 | 4.579 | 8.894 | 17.304 | 34.800 | 69.672 |
| 49.3164 | 3.860 | 9.330 | 18.407 | 35.384 | 70.785 |
| 0 | 3.850 | 8.904 | 17.976 | 35.898 | 72.592 |

*Cpd.* **40**: *Determination of $IC_{50}$ and $k_1^*$*

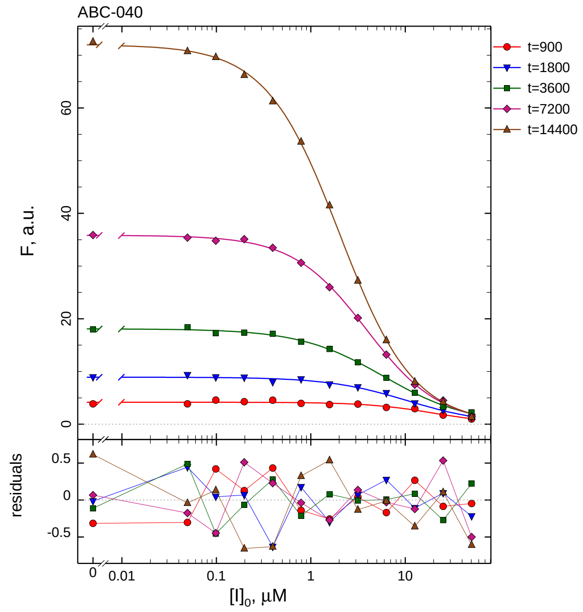

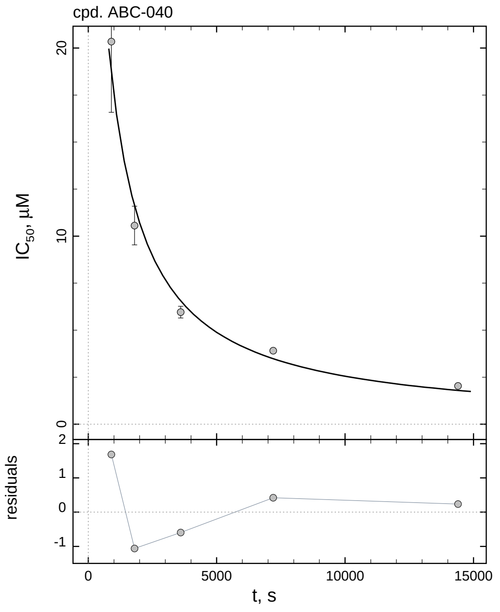

| $i$ | $t$, min | $I_{50}$, nM | ± | std.err. | CV, % | $k_1^*$, $\mathrm{mM^{-1}s^{-1}}$ |
|---|---|---|---|---|---|---|
| 1 | 15 | 20342 | ± | 3761.9 | 18.5 | 0.08704 |
| 2 | 30 | 10562 | ± | 1028.7 | 9.7 | 0.08383 |
| 3 | 60 | 5958.2 | ± | 312.0 | 5.2 | 0.07430 |
| 4 | 120 | 3904.6 | ± | 94.05 | 2.4 | 0.05669 |
| 5 | 240 | 2032.3 | ± | 25.70 | 1.3 | 0.05445 |

*Cpd.* **40**: *Mechanistic analysis*

| parameter | unit | "true" (T) | calculated (C) | C/T ratio | note |
|---|---|---|---|---|---|
| mechanism | | **C2S** | **C2F** | | |
| $k_{\mathrm{eff}}$ | $\mathrm{mM^{-1}s^{-1}}$ | 0.09901 | 0.1007 | 1.02 | $= k_{\mathrm{inact}}/K_{\mathrm{i}}$ |
| $K_{\mathrm{i}}$ | nM | 10100 | 12233 | 1.21 | |
| $k_{\mathrm{inact}}$ | $\mathrm{s^{-1}}$ | 0.001 | 0.001232 | 1.23 | |

| | |
|---|---|
| Reaction times used for analysis: | 30 and 120 min |
| Maximum GSD for accepting one-step model **C1**: | 1.25 |
| Observed GSD: | 1.32 |
| Assumed $[S]_0/K_M$ ratio: | 1.0 |

*2.41. Compound No. 41*

*Cpd.* **41***: Simulated data*

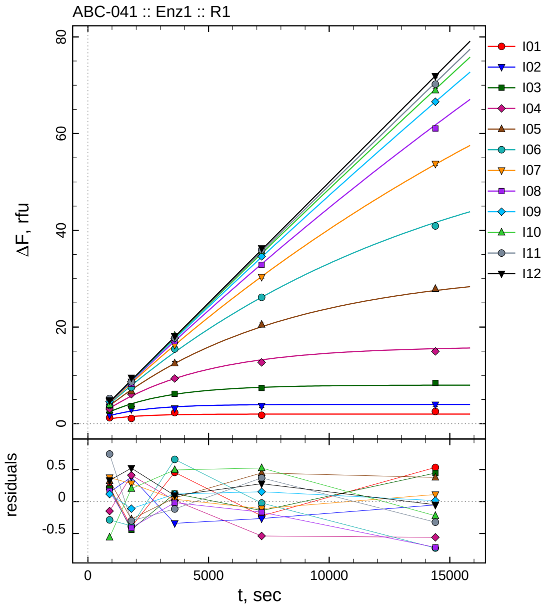

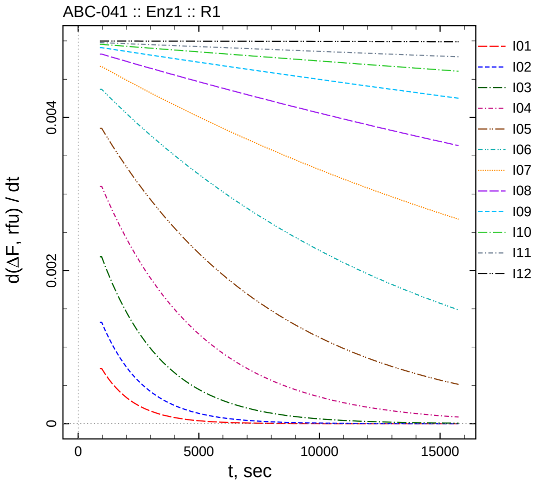

| $[I]_0$, nM | t = 15 | 30 | 60 | 120 | 240 min |
|---|---|---|---|---|---|
| 5.5 | 1.233 | 1.090 | 2.316 | 1.764 | 2.533 |
| 2.75 | 1.817 | 2.986 | 3.157 | 3.670 | 3.945 |
| 1.375 | 2.668 | 3.668 | 6.209 | 7.393 | 8.419 |
| 0.6875 | 3.048 | 6.116 | 9.360 | 12.680 | 14.958 |
| 0.34375 | 4.070 | 6.738 | 12.518 | 20.501 | 27.918 |
| 0.171875 | 3.802 | 7.514 | 15.453 | 26.129 | 40.890 |
| 0.0859375 | 4.660 | 8.694 | 16.297 | 30.333 | 53.747 |
| 0.0429688 | 4.552 | 8.287 | 17.073 | 32.864 | 61.066 |
| 0.0214844 | 4.560 | 8.733 | 17.650 | 34.610 | 66.586 |
| 0.0107422 | 3.911 | 9.121 | 18.256 | 35.731 | 68.952 |
| 0.00537109 | 5.227 | 8.656 | 17.760 | 35.960 | 70.212 |
| 0 | 4.828 | 9.519 | 18.089 | 36.261 | 71.883 |

*Cpd.* **41***: Determination of* $IC_{50}$ *and* $k_1^*$

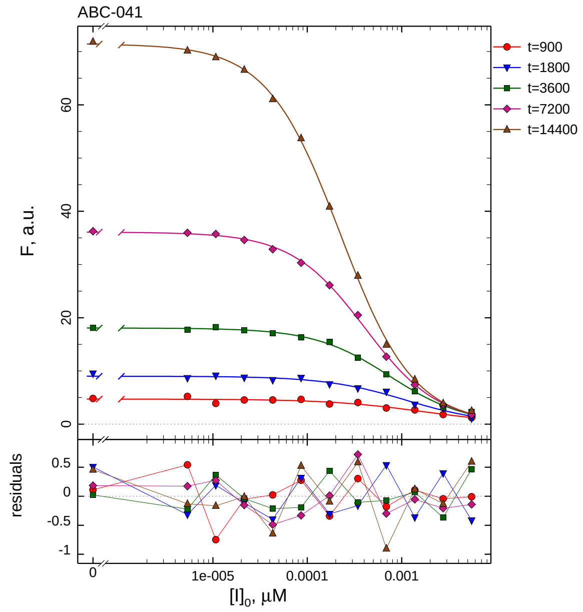

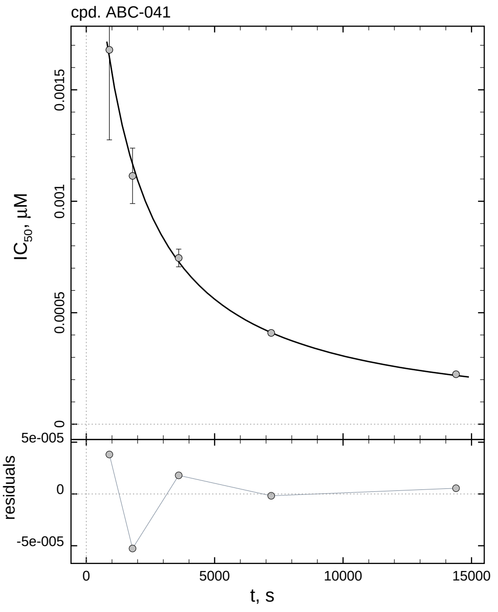

| $i$ | $t$, min | $I_{50}$, nM | ± | std.err. | CV, % | $k_1^*$, $mM^{-1}s^{-1}$ |
|---|---|---|---|---|---|---|
| 1 | 15 | 1.679 | ± | 0.4037 | 24.0 | 1054.4 |
| 2 | 30 | 1.114 | ± | 0.1245 | 11.2 | 794.7 |
| 3 | 60 | 0.7458 | ± | 0.03974 | 5.3 | 593.5 |
| 4 | 120 | 0.4093 | ± | 0.01110 | 2.7 | 540.8 |
| 5 | 240 | 0.2242 | ± | 0.003189 | 1.4 | 493.7 |

*Cpd.* **41***: Mechanistic analysis*

| parameter | unit | "true" (T) | calculated (C) | C/T ratio | note |
|---|---|---|---|---|---|
| mechanism | | **C2S** | **C2F** | | |
| $k_{eff}$ | $mM^{-1}s^{-1}$ | 909.1 | 963.7 | 1.06 | $= k_{inact}/K_i$ |
| $K_i$ | nM | 1.1 | 1.307 | 1.19 | |
| $k_{inact}$ | $s^{-1}$ | 0.001 | 0.001260 | 1.26 | |

| | |
|---|---|
| Reaction times used for analysis: | 30 and 120 min |
| Maximum GSD for accepting one-step model **C1**: | 1.25 |
| Observed GSD: | 1.31 |
| Assumed $[S]_0/K_M$ ratio: | 1.0 |

*2.42. Compound No. 42*

*Cpd.* **42***: Simulated data*

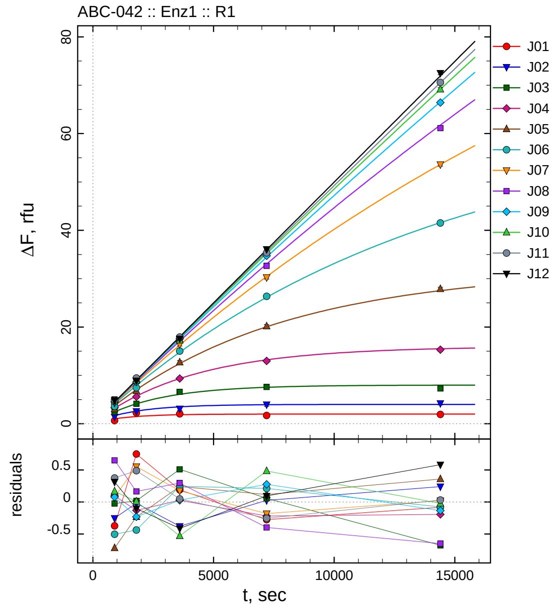

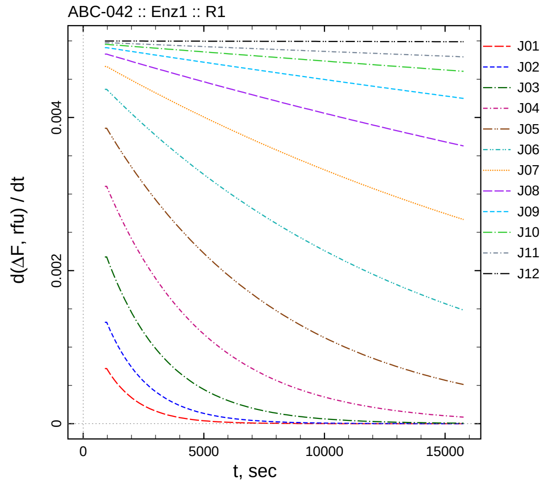

| $[I]_0$, nM | t = 15 | 30 | 60 | 120 | 240 min |
|---|---|---|---|---|---|
| 55 | 0.637 | 2.236 | 2.052 | 1.714 | 1.917 |
| 27.5 | 1.419 | 2.585 | 3.119 | 3.961 | 4.240 |
| 13.75 | 2.435 | 4.130 | 6.594 | 7.592 | 7.296 |
| 6.875 | 3.292 | 5.572 | 9.368 | 12.994 | 15.314 |
| 3.4375 | 3.027 | 6.781 | 12.700 | 20.161 | 27.877 |
| 1.71875 | 3.586 | 7.457 | 15.040 | 26.357 | 41.508 |
| 0.859375 | 4.364 | 8.968 | 16.438 | 30.257 | 53.632 |
| 0.429688 | 5.041 | 8.863 | 17.387 | 32.630 | 61.114 |
| 0.214844 | 4.521 | 8.613 | 17.562 | 34.731 | 66.426 |
| 0.107422 | 4.638 | 8.930 | 17.231 | 35.690 | 69.149 |
| 0.0537109 | 4.860 | 9.450 | 17.926 | 35.336 | 70.563 |
| 0 | 4.814 | 8.896 | 17.583 | 36.078 | 72.518 |

*Cpd.* **42***: Determination of* $IC_{50}$ *and* $k_1^*$

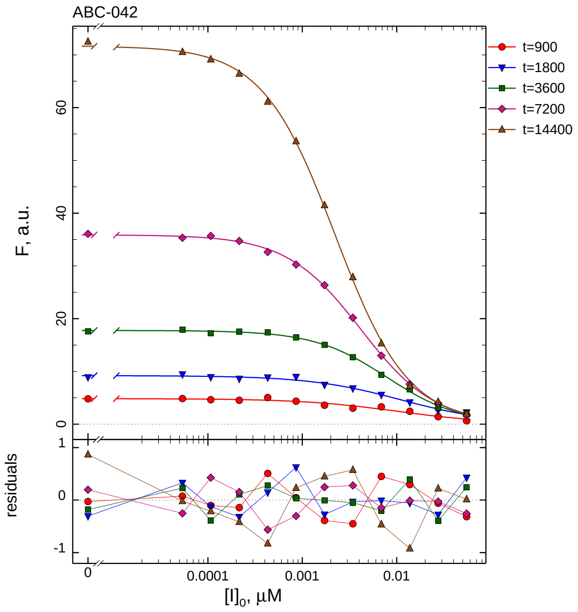

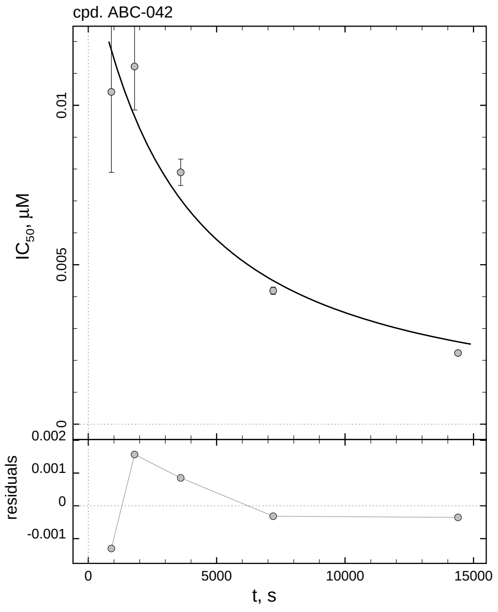

| $i$ | $t$, min | $I_{50}$, nM | ± | std.err. | CV, % | $k_1^*$, $mM^{-1}s^{-1}$ |
|---|---|---|---|---|---|---|
| 1 | 15 | 10.42 | ± | 2.524 | 24.2 | 169.9 |
| 2 | 30 | 11.21 | ± | 1.362 | 12.1 | 78.95 |
| 3 | 60 | 7.896 | ± | 0.4130 | 5.2 | 56.06 |
| 4 | 120 | 4.185 | ± | 0.1139 | 2.7 | 52.89 |
| 5 | 240 | 2.233 | ± | 0.03135 | 1.4 | 49.56 |

*Cpd.* **42***: Mechanistic analysis*

| parameter | unit | "true" (T) | calculated (C) | C/T ratio | note |
|---|---|---|---|---|---|
| mechanism | | **C2S** | **C2F** | | |
| $k_{eff}$ | $mM^{-1}s^{-1}$ | 90.91 | 93.50 | 1.03 | $= k_{inact}/K_i$ |
| $K_i$ | nM | 11 | 12.74 | 1.16 | |
| $k_{inact}$ | $s^{-1}$ | 0.001 | 0.001191 | 1.19 | |

| | |
|---|---|
| Reaction times used for analysis: | 30 and 120 min |
| Maximum GSD for accepting one-step model **C1**: | 1.25 |
| Observed GSD: | 1.33 |
| Assumed $[S]_0/K_M$ ratio: | 1.0 |

*2.43. Compound No. 43*

*Cpd.* **43***: Simulated data*

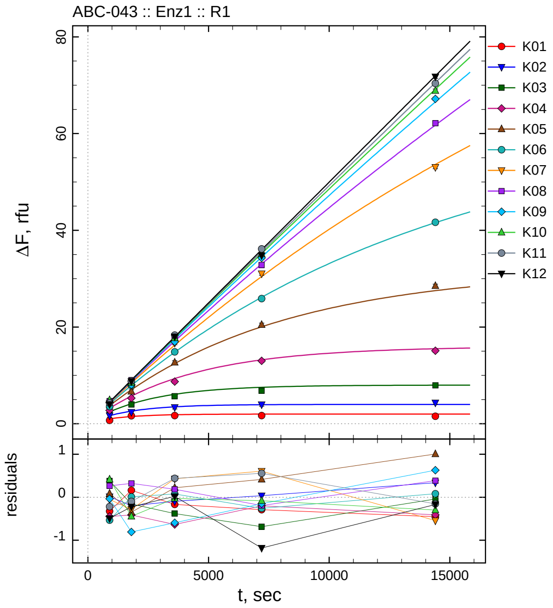

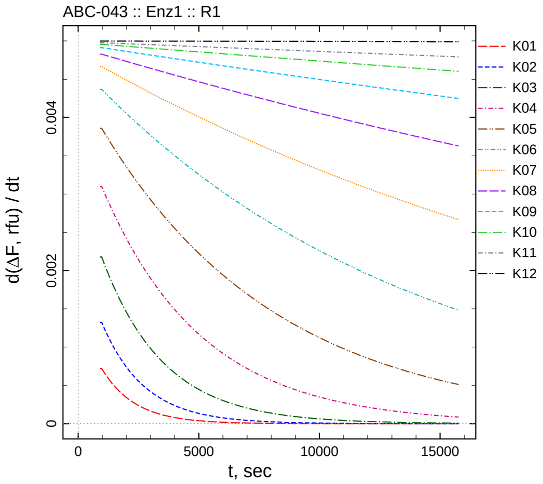

| $[I]_0$, nM | t = 15 | 30 | 60 | 120 | 240 min |
|---|---|---|---|---|---|
| 550 | 0.688 | 1.649 | 1.694 | 1.700 | 1.549 |
| 275 | 1.686 | 2.406 | 3.404 | 3.973 | 4.336 |
| 137.5 | 2.841 | 3.956 | 5.703 | 6.851 | 7.932 |
| 68.75 | 2.751 | 5.292 | 8.707 | 13.013 | 15.098 |
| 34.375 | 3.834 | 6.666 | 12.679 | 20.460 | 28.521 |
| 17.1875 | 3.557 | 7.904 | 14.870 | 25.888 | 41.666 |
| 8.59375 | 4.234 | 8.133 | 16.690 | 31.046 | 53.066 |
| 4.29688 | 4.656 | 9.020 | 17.278 | 32.834 | 62.147 |
| 2.14844 | 4.409 | 8.039 | 16.941 | 34.304 | 67.188 |
| 1.07422 | 4.886 | 8.477 | 17.735 | 35.124 | 68.872 |
| 0.537109 | 4.267 | 8.864 | 18.318 | 36.148 | 70.379 |
| 0 | 4.010 | 8.776 | 18.015 | 34.803 | 71.766 |

*Cpd.* **43***: Determination of* $IC_{50}$ *and* $k_1^*$

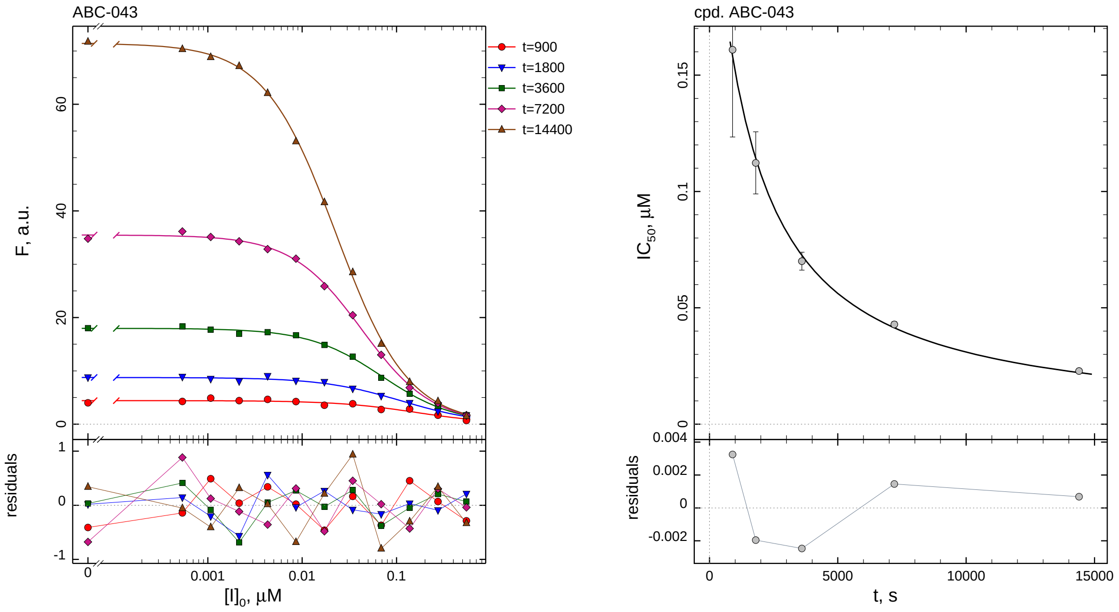

| $i$ | $t$, min | $I_{50}$, nM | ± | std.err. | CV, % | $k_1^*$, $mM^{-1}s^{-1}$ |
|---|---|---|---|---|---|---|
| 1 | 15 | 160.9 | ± | 37.50 | 23.3 | 11.00 |
| 2 | 30 | 112.3 | ± | 13.35 | 11.9 | 7.885 |
| 3 | 60 | 70.05 | ± | 3.856 | 5.5 | 6.319 |
| 4 | 120 | 42.88 | ± | 1.193 | 2.8 | 5.162 |
| 5 | 240 | 22.86 | ± | 0.3366 | 1.5 | 4.841 |

*Cpd.* **43***: Mechanistic analysis*

| parameter | unit | "true" (T) | calculated (C) | C/T ratio | note |
|---|---|---|---|---|---|
| mechanism | | **C2S** | **C2F** | | |
| $k_{eff}$ | $mM^{-1}s^{-1}$ | 9.091 | 9.016 | 0.99 | $= k_{inact}/K_i$ |
| $K_i$ | nM | 110 | 121.9 | 1.11 | |
| $k_{inact}$ | $s^{-1}$ | 0.001 | 0.001099 | 1.10 | |

| | |
|---|---|
| Reaction times used for analysis: | 30 and 120 min |
| Maximum GSD for accepting one-step model **C1**: | 1.25 |
| Observed GSD: | 1.35 |
| Assumed $[S]_0/K_M$ ratio: | 1.0 |

#### 2.44. *Compound No. 44*

*Cpd.* ***44****: Simulated data*

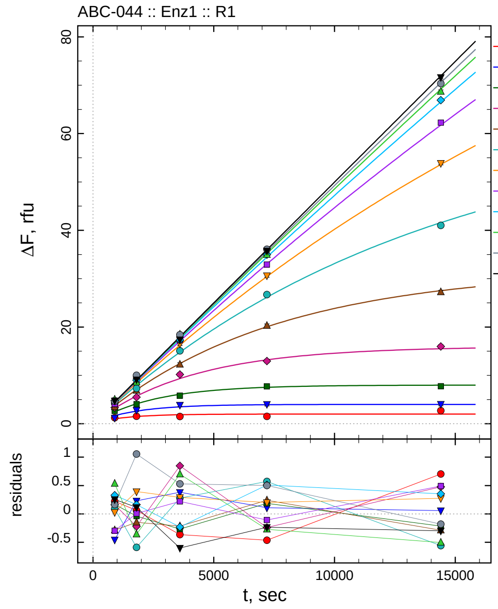

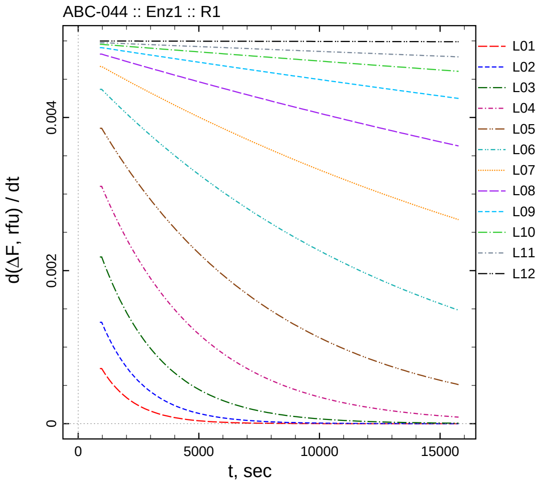

| $[I]_0$, nM | t = 15 | 30 | 60 | 120 | 240 min |
|---:|---:|---:|---:|---:|---:|
| 5500 | 1.232 | 1.548 | 1.498 | 1.527 | 2.704 |
| 2750 | 1.215 | 2.835 | 3.881 | 4.044 | 4.057 |
| 1375 | 2.627 | 4.072 | 5.817 | 7.719 | 7.757 |
| 687.5 | 3.358 | 5.484 | 10.188 | 12.973 | 15.985 |
| 343.75 | 3.460 | 6.866 | 12.245 | 20.285 | 27.228 |
| 171.875 | 4.181 | 7.305 | 15.072 | 26.716 | 41.025 |
| 85.9375 | 4.317 | 8.809 | 16.552 | 30.640 | 53.882 |
| 42.9688 | 4.094 | 8.705 | 17.312 | 32.920 | 62.253 |
| 21.4844 | 4.773 | 9.009 | 17.303 | 34.965 | 66.912 |
| 10.7422 | 5.005 | 8.563 | 18.461 | 34.935 | 68.665 |
| 5.37109 | 4.650 | 10.014 | 18.407 | 36.092 | 70.351 |
| 0 | 4.753 | 9.115 | 17.395 | 35.748 | 71.636 |

*Cpd.* **44**: *Determination of* $IC_{50}$ *and* $k_1^*$

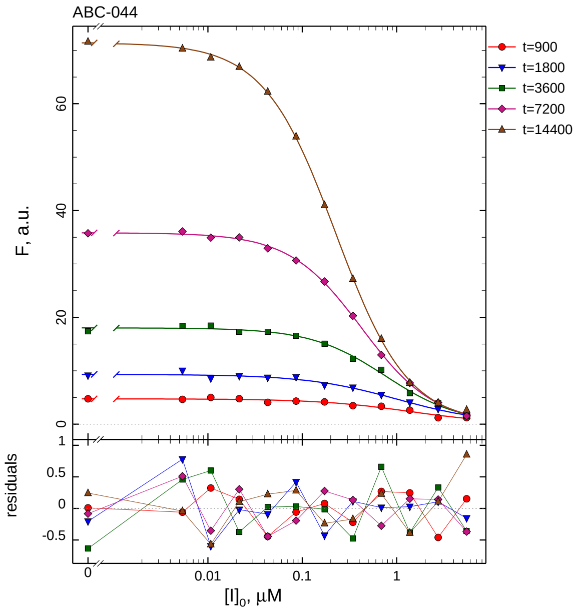

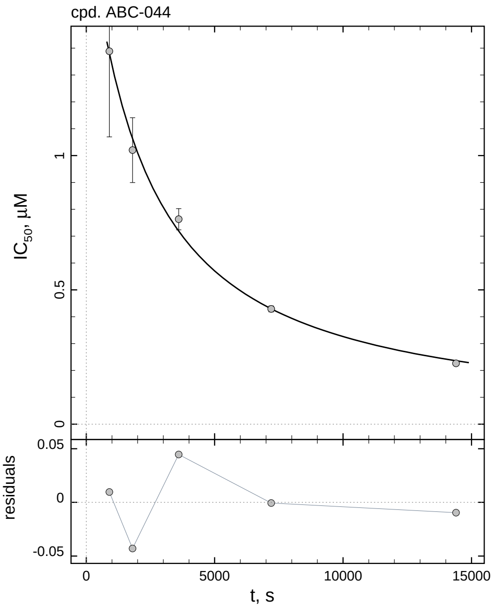

| $i$ | $t$, min | $I_{50}$, nM | ± | std.err. | CV, % | $k_1^*$, mM$^{-1}$s$^{-1}$ |
|---|---|---|---|---|---|---|
| 1 | 15 | 1388.3 | ± | 318.9 | 23.0 | 1.275 |
| 2 | 30 | 1020.3 | ± | 120.7 | 11.8 | 0.8677 |
| 3 | 60 | 763.0 | ± | 39.54 | 5.2 | 0.5802 |
| 4 | 120 | 429.0 | ± | 11.26 | 2.6 | 0.5159 |
| 5 | 240 | 226.6 | ± | 3.109 | 1.4 | 0.4884 |

*Cpd.* **44**: *Mechanistic analysis*

| parameter | unit | "true" (T) | calculated (C) | C/T ratio | note |
|---|---|---|---|---|---|
| mechanism | | **C2S** | **C2F** | | |
| $k_{\mathrm{eff}}$ | mM$^{-1}$s$^{-1}$ | 0.9091 | 0.8507 | 0.94 | $= k_{\mathrm{inact}}/K_{\mathrm{i}}$ |
| $K_{\mathrm{i}}$ | nM | 1100 | 943.6 | 0.86 | |
| $k_{\mathrm{inact}}$ | s$^{-1}$ | 0.001 | 0.0008028 | 0.80 | |

| | |
|---|---|
| Reaction times used for analysis: | 30 and 120 min |
| Maximum GSD for accepting one-step model **C1**: | 1.25 |
| Observed GSD: | 1.44 |
| Assumed $[S]_0/K_M$ ratio: | 1.0 |

*2.45. Compound No. 45*

*Cpd.* **45***: Simulated data*

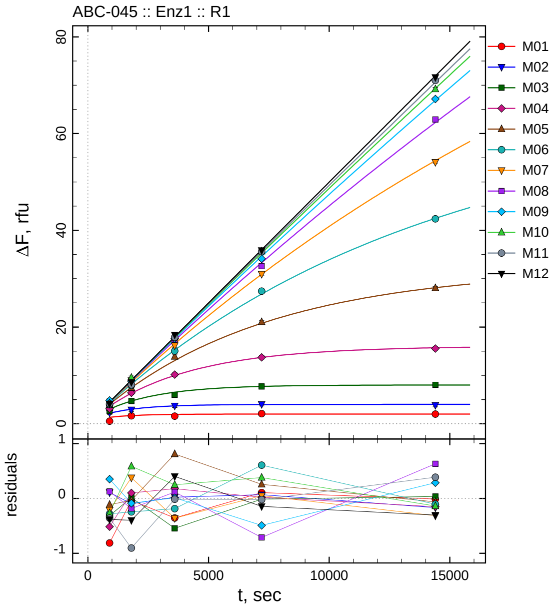

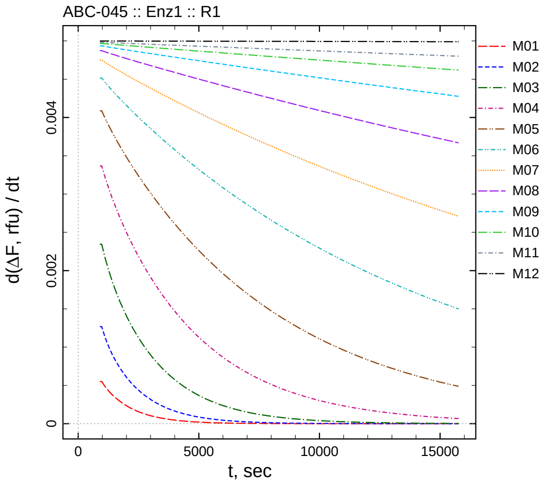

| $[I]_0$, nM | t = 15 | 30 | 60 | 120 | 240 min |
|---:|---:|---:|---:|---:|---:|
| 1 | 0.524 | 1.645 | 1.571 | 2.105 | 1.991 |
| 0.5 | 2.275 | 2.896 | 3.713 | 4.044 | 3.839 |
| 0.25 | 2.689 | 4.701 | 5.980 | 7.704 | 8.041 |
| 0.125 | 3.122 | 6.390 | 10.162 | 13.729 | 15.560 |
| 0.0625 | 3.921 | 7.422 | 13.886 | 21.028 | 28.067 |
| 0.03125 | 3.977 | 7.928 | 15.051 | 27.421 | 42.370 |
| 0.015625 | 4.165 | 8.950 | 16.170 | 30.951 | 54.090 |
| 0.0078125 | 4.565 | 8.596 | 17.356 | 32.590 | 62.933 |
| 0.00390625 | 4.820 | 8.802 | 17.623 | 34.113 | 67.160 |
| 0.00195313 | 4.218 | 9.528 | 18.050 | 35.664 | 69.202 |
| 0.000976563 | 4.134 | 8.068 | 17.884 | 35.610 | 71.008 |
| 0 | 4.116 | 8.600 | 18.398 | 35.839 | 71.632 |

*Cpd.* **45***: Determination of* $IC_{50}$ *and* $k_1^*$

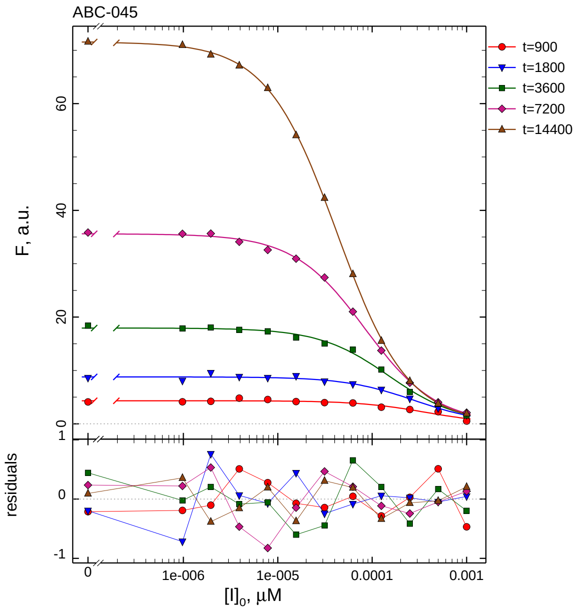

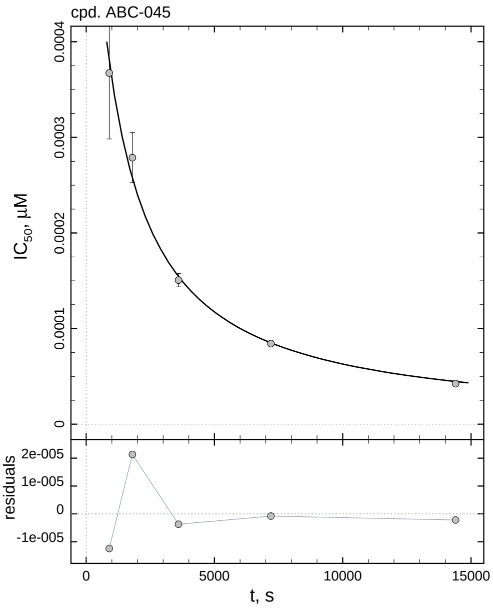

| $i$ | $t$, min | $I_{50}$, nM | ± | std.err. | CV, % | $k_1^*$, $mM^{-1}s^{-1}$ |
|---|---|---|---|---|---|---|
| 1 | 15 | 0.3673 | ± | 0.06891 | 18.8 | 4820.7 |
| 2 | 30 | 0.2788 | ± | 0.02624 | 9.4 | 3175.0 |
| 3 | 60 | 0.1507 | ± | 0.007109 | 4.7 | 2936.9 |
| 4 | 120 | 0.08425 | ± | 0.002099 | 2.5 | 2626.9 |
| 5 | 240 | 0.04245 | ± | 0.0005457 | 1.3 | 2606.9 |

*Cpd.* **45***: Mechanistic analysis*

| parameter | unit | "true" (T) | calculated (C) | C/T ratio | note |
|---|---|---|---|---|---|
| mechanism | | **C2S** | **C1** | | |
| $k_{eff}$ | $mM^{-1}s^{-1}$ | 5000.0 | 5253.9 | 1.05 | $= k_1^* \ (1 + [S]_0/K_M)$ |

| | |
|---|---|
| Reaction times used for analysis: | 30 and 120 min |
| Maximum GSD for accepting one-step model **C1**: | 1.25 |
| Observed GSD: | 1.14 |
| Assumed $[S]_0/K_M$ ratio: | 1.0 |

*Cpd.* **46***: Simulated data*

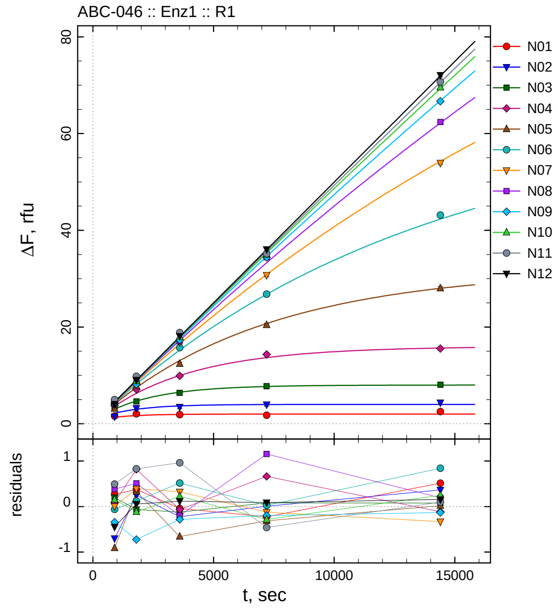

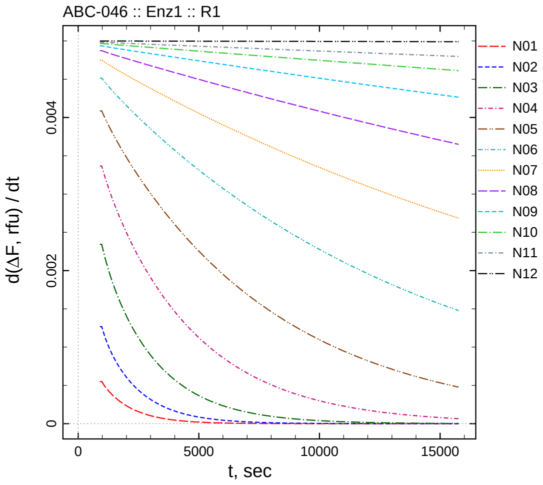

| $[I]_0$, nM | t = 15 | 30 | 60 | 120 | 240 min |
|---:|---:|---:|---:|---:|---:|
| 10 | 1.602 | 2.055 | 1.883 | 1.771 | 2.515 |
| 5 | 1.464 | 3.237 | 3.463 | 3.977 | 4.366 |
| 2.5 | 3.201 | 4.618 | 6.391 | 7.781 | 8.066 |
| 1.25 | 3.725 | 7.104 | 9.904 | 14.340 | 15.540 |
| 0.625 | 3.122 | 7.783 | 12.405 | 20.415 | 28.024 |
| 0.3125 | 4.194 | 8.290 | 15.744 | 26.797 | 43.153 |
| 0.15625 | 4.378 | 8.962 | 16.848 | 30.749 | 53.929 |
| 0.078125 | 4.820 | 9.291 | 17.036 | 34.439 | 62.397 |
| 0.0390625 | 4.124 | 8.162 | 17.329 | 34.398 | 66.686 |
| 0.0195313 | 4.635 | 8.824 | 18.026 | 34.996 | 69.556 |
| 0.00976563 | 4.983 | 9.798 | 18.859 | 35.167 | 70.703 |
| 0 | 4.051 | 9.053 | 18.114 | 36.069 | 72.090 |

*Cpd.* **46***: Determination of* $IC_{50}$ *and* $k_1^*$

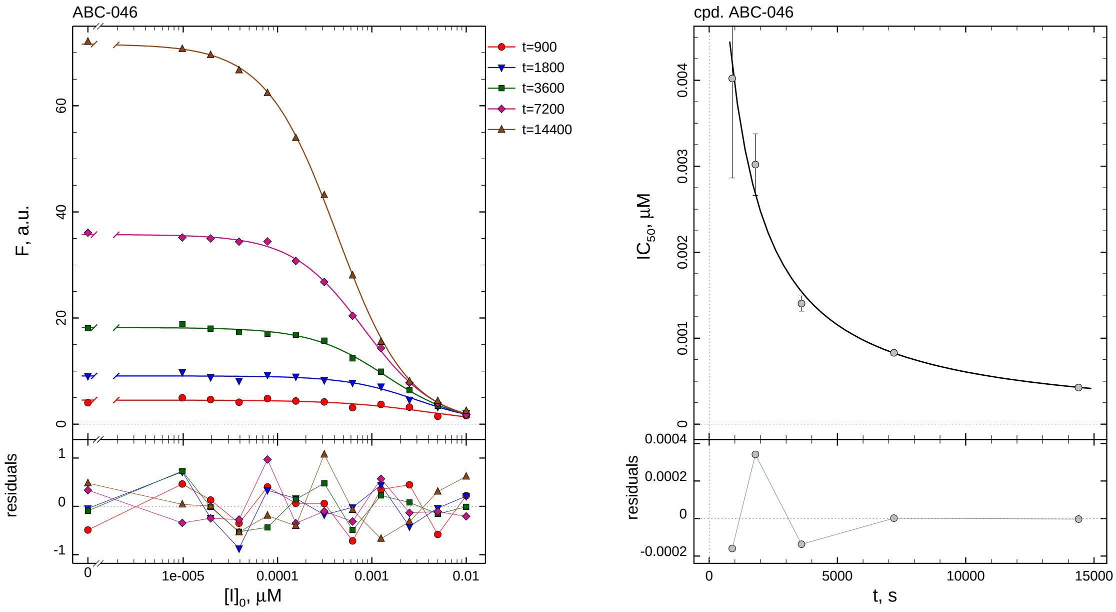

| $i$ | $t$, min | $I_{50}$, nM | ± | std.err. | CV, % | $k_1^*$, $mM^{-1}s^{-1}$ |
|---|---|---|---|---|---|---|
| 1 | 15 | 4.020 | ± | 1.157 | 28.8 | 440.4 |
| 2 | 30 | 3.019 | ± | 0.3570 | 11.8 | 293.2 |
| 3 | 60 | 1.403 | ± | 0.08762 | 6.2 | 315.5 |
| 4 | 120 | 0.8298 | ± | 0.02633 | 3.2 | 266.7 |
| 5 | 240 | 0.4261 | ± | 0.007055 | 1.7 | 259.7 |

*Cpd.* **46***: Mechanistic analysis*

| parameter | unit | "true" (T) | calculated (C) | C/T ratio | note |
|---|---|---|---|---|---|
| mechanism | | **C2S** | **C1** | | |
| $k_{eff}$ | $mM^{-1}s^{-1}$ | 500.0 | 533.4 | 1.07 | $= k_1^* \ (1 + [S]_0/K_M)$ |

| | |
|---|---|
| Reaction times used for analysis: | 30 and 120 min |
| Maximum GSD for accepting one-step model **C1**: | 1.25 |
| Observed GSD: | 1.07 |
| Assumed $[S]_0/K_M$ ratio: | 1.0 |

*2.47. Compound No. 47*

*Cpd.* ***47****: Simulated data*

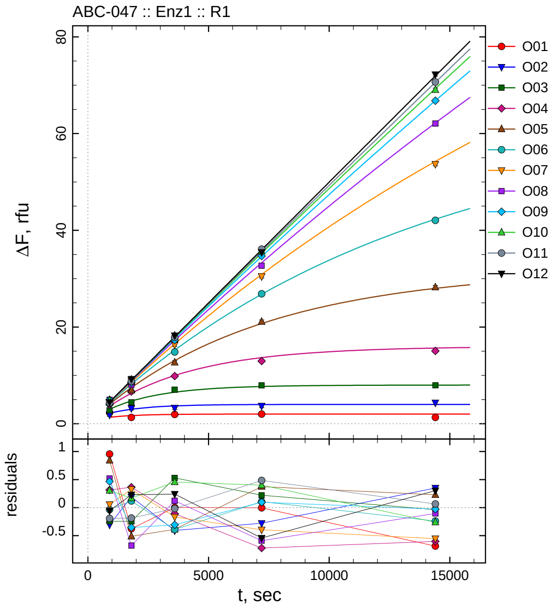

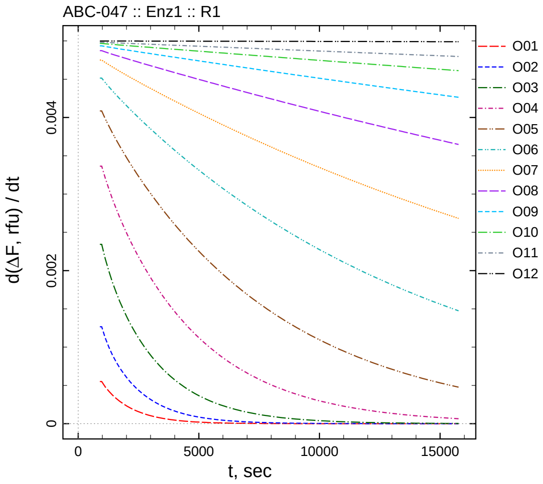

| $[I]_0$, nM | t = 15 | 30 | 60 | 120 | 240 min |
|---:|---:|---:|---:|---:|---:|
| 100 | 2.289 | 1.303 | 1.927 | 1.990 | 1.314 |
| 50 | 1.859 | 3.179 | 3.278 | 3.690 | 4.349 |
| 25 | 2.757 | 4.435 | 7.043 | 7.920 | 7.949 |
| 12.5 | 3.949 | 6.649 | 9.861 | 12.953 | 15.047 |
| 6.25 | 4.873 | 6.942 | 12.676 | 21.098 | 28.221 |
| 3.125 | 4.220 | 8.289 | 14.836 | 26.877 | 42.054 |
| 1.5625 | 4.432 | 8.896 | 16.357 | 30.477 | 53.694 |
| 0.78125 | 4.953 | 8.102 | 17.358 | 32.693 | 62.091 |
| 0.390625 | 4.933 | 8.534 | 17.301 | 34.690 | 66.786 |
| 0.195313 | 4.787 | 9.111 | 18.256 | 35.680 | 69.045 |
| 0.0976563 | 4.298 | 8.784 | 17.880 | 36.112 | 70.668 |
| 0 | 4.442 | 9.227 | 18.235 | 35.440 | 72.237 |

*Cpd.* **47***: Determination of* $IC_{50}$ *and* $k_1^*$

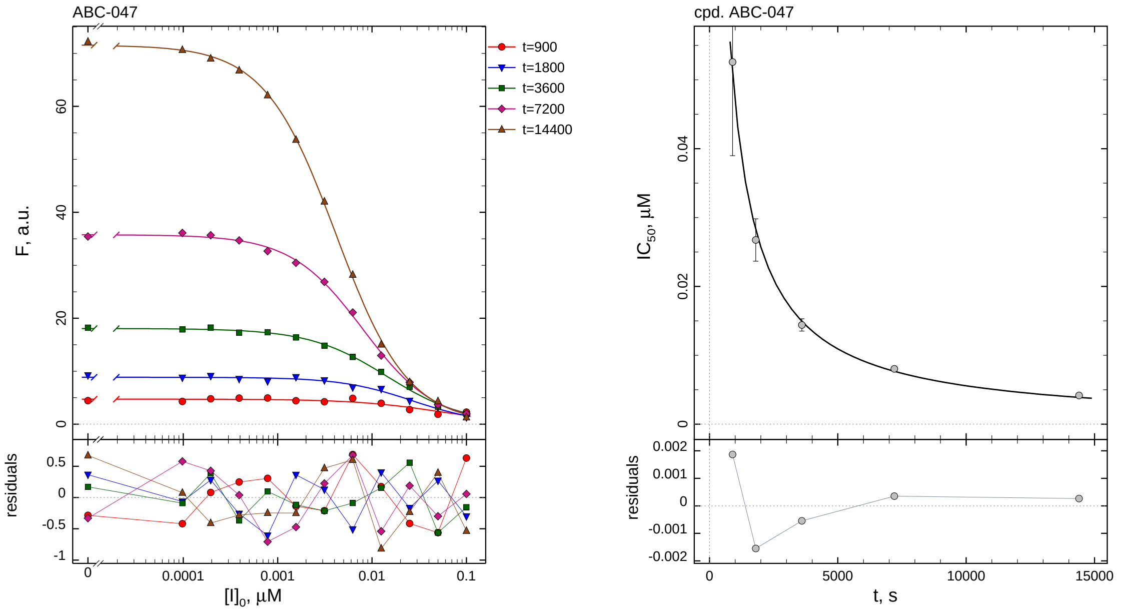

| $i$ | $t$, min | $I_{50}$, nM | ± | std.err. | CV, % | $k_1^*$, mM$^{-1}$s$^{-1}$ |
|---|---|---|---|---|---|---|
| 1 | 15 | 52.57 | ± | 13.60 | 25.9 | 33.68 |
| 2 | 30 | 26.74 | ± | 3.068 | 11.5 | 33.11 |
| 3 | 60 | 14.41 | ± | 0.8862 | 6.2 | 30.72 |
| 4 | 120 | 8.037 | ± | 0.2422 | 3.0 | 27.54 |
| 5 | 240 | 4.162 | ± | 0.06530 | 1.6 | 26.59 |

*Cpd.* **47***: Mechanistic analysis*

| parameter | unit | "true" (T) | calculated (C) | C/T ratio | note |
|---|---|---|---|---|---|
| mechanism | | **C2S** | **C1** | | |
| $k_{\text{eff}}$ | mM$^{-1}$s$^{-1}$ | 50.00 | 55.08 | 1.10 | $= k_1^* \, (1 + [S]_0/K_M)$ |

| | |
|---|---|
| Reaction times used for analysis: | 30 and 120 min |
| Maximum GSD for accepting one-step model **C1**: | 1.25 |
| Observed GSD: | 1.14 |
| Assumed $[S]_0/K_M$ ratio: | 1.0 |

*2.48. Compound No. 48*

*Cpd.* **48***: Simulated data*

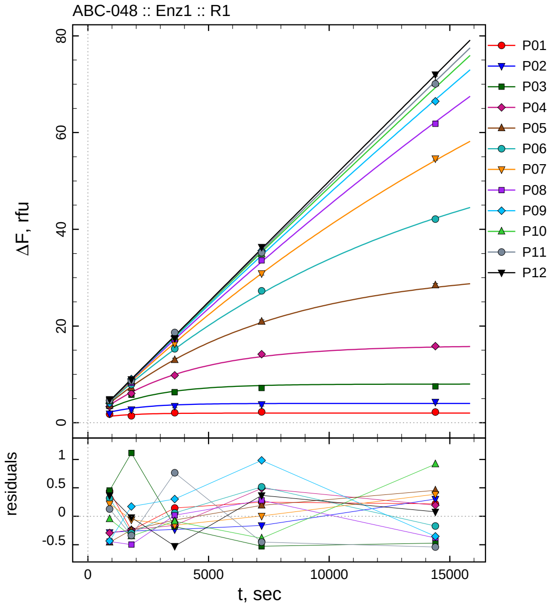

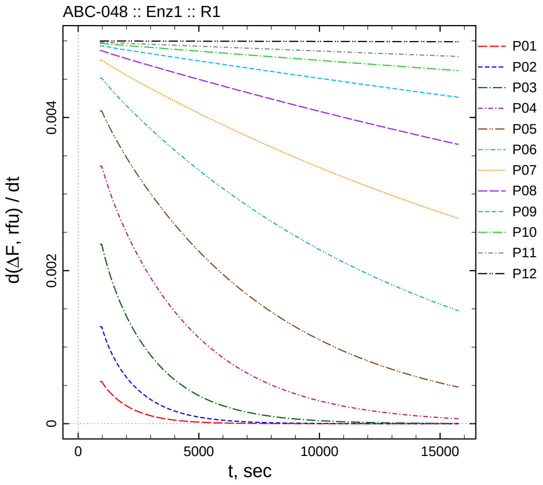

| $[I]_0$, nM | t = 15 | 30 | 60 | 120 | 240 min |
|---|---|---|---|---|---|
| 1000 | 1.777 | 1.430 | 2.071 | 2.244 | 2.219 |
| 500 | 1.881 | 2.729 | 3.454 | 3.807 | 4.303 |
| 250 | 3.453 | 5.793 | 6.332 | 7.174 | 7.513 |
| 125 | 3.341 | 6.047 | 9.811 | 14.163 | 15.842 |
| 62.5 | 3.574 | 7.213 | 12.997 | 20.914 | 28.448 |
| 31.25 | 4.573 | 7.875 | 15.286 | 27.286 | 42.121 |
| 15.625 | 4.597 | 8.508 | 16.365 | 30.877 | 54.634 |
| 7.8125 | 3.991 | 8.281 | 17.253 | 33.565 | 61.813 |
| 3.90625 | 4.034 | 9.059 | 17.912 | 35.578 | 66.458 |
| 1.95313 | 4.435 | 8.593 | 17.714 | 34.891 | 70.223 |
| 0.976563 | 4.617 | 8.633 | 18.663 | 35.173 | 70.062 |
| 0 | 4.866 | 8.978 | 17.470 | 36.350 | 72.013 |

*Cpd.* **48***: Determination of $IC_{50}$ and $k_1^*$*

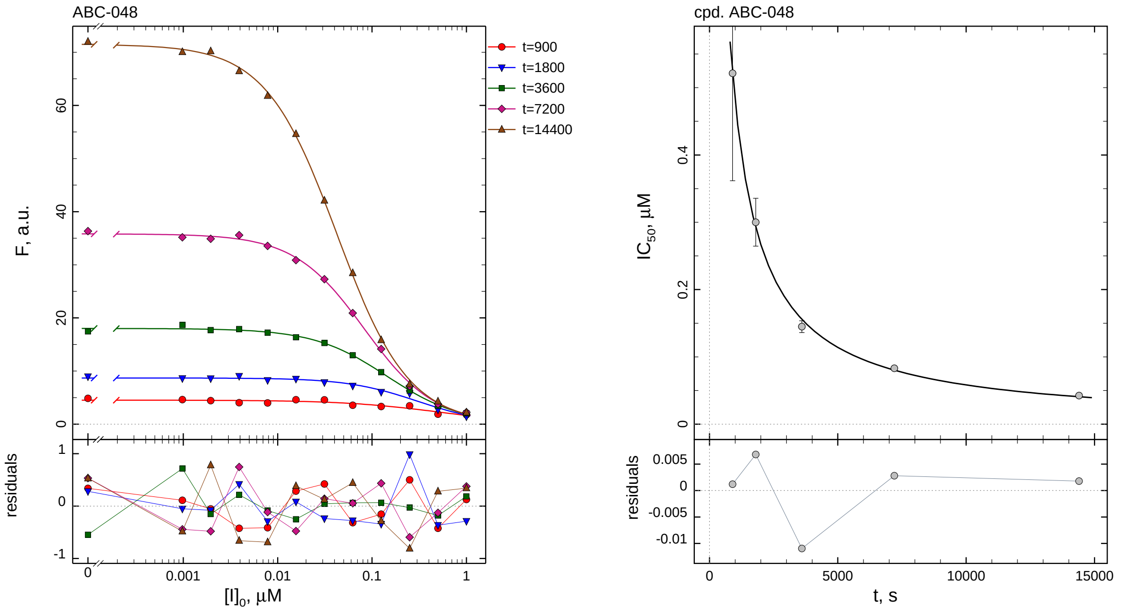

| $i$ | $t$, min | $I_{50}$, nM | ± | std.err. | CV, % | $k_1^*$, $mM^{-1}s^{-1}$ |
|---|---|---|---|---|---|---|
| 1 | 15 | 521.4 | ± | 159.7 | 30.6 | 3.396 |
| 2 | 30 | 300.1 | ± | 35.51 | 11.8 | 2.951 |
| 3 | 60 | 144.9 | ± | 8.770 | 6.1 | 3.054 |
| 4 | 120 | 83.14 | ± | 2.506 | 3.0 | 2.662 |
| 5 | 240 | 42.56 | ± | 0.6835 | 1.6 | 2.601 |

*Cpd.* **48***: Mechanistic analysis*

| parameter | unit | "true" (T) | calculated (C) | C/T ratio | note |
|---|---|---|---|---|---|
| mechanism | | **C2S** | **C1** | | |
| $k_{\text{eff}}$ | $mM^{-1}s^{-1}$ | 5.000 | 5.324 | 1.06 | $= k_1^* \, (1 + [S]_0/K_M)$ |

| | |
|---|---|
| Reaction times used for analysis: | 30 and 120 min |
| Maximum GSD for accepting one-step model **C1**: | 1.25 |
| Observed GSD: | 1.08 |
| Assumed $[S]_0/K_M$ ratio: | 1.0 |

#### *2.49. Compound No. 49*

*Cpd.* **49***: Simulated data*

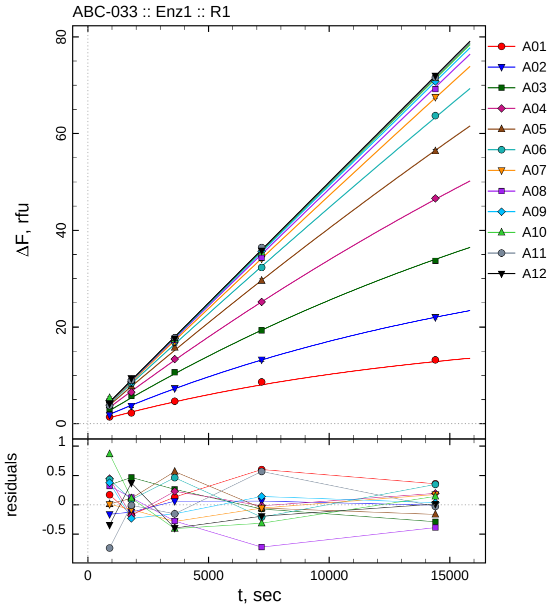

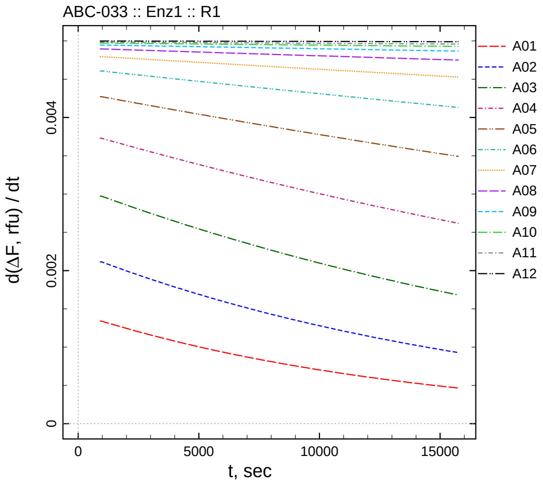

| $[I]_0$, nM | t = 15 | 30 | 60 | 120 | 240 min |
|---|---|---|---|---|---|
| 500.05 | 1.416 | 2.246 | 4.666 | 8.642 | 13.205 |
| 250.025 | 1.789 | 3.683 | 7.313 | 13.247 | 22.009 |
| 125.013 | 3.060 | 5.817 | 10.606 | 19.274 | 33.713 |
| 62.5062 | 3.839 | 6.529 | 13.370 | 25.179 | 46.600 |
| 31.2531 | 3.883 | 7.799 | 15.756 | 29.600 | 56.376 |
| 15.6266 | 4.591 | 8.379 | 16.937 | 32.311 | 63.708 |
| 7.81328 | 4.331 | 8.551 | 16.917 | 34.109 | 67.565 |
| 3.90664 | 4.727 | 8.933 | 17.314 | 34.331 | 69.204 |
| 1.95332 | 4.829 | 8.671 | 17.637 | 35.652 | 70.781 |
| 0.97666 | 5.340 | 9.062 | 17.485 | 35.431 | 71.479 |
| 0.48833 | 3.752 | 8.965 | 17.790 | 36.431 | 71.604 |
| 0 | 4.156 | 9.371 | 17.599 | 35.788 | 71.946 |

*Cpd.* **49***: Determination of* $IC_{50}$ *and* $k_1^*$

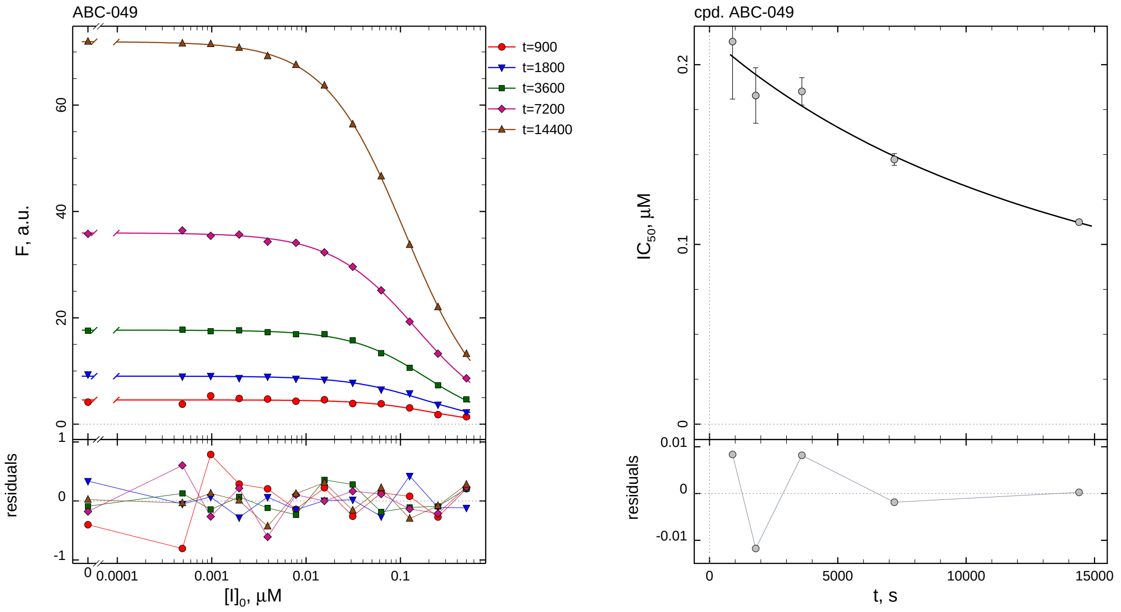

| $i$ | $t$, min | $I_{50}$, nM ± std.err. | CV, % | $k_1^*$, mM$^{-1}$s$^{-1}$ |
|---|---|---|---|---|
| 1 | 15 | 212.8 ± 31.98 | 15.0 | 8.319 |
| 2 | 30 | 182.8 ± 15.43 | 8.4 | 4.842 |
| 3 | 60 | 185.1 ± 7.617 | 4.1 | 2.391 |
| 4 | 120 | 147.2 ± 3.269 | 2.2 | 1.503 |
| 5 | 240 | 112.4 ± 1.207 | 1.1 | 0.9849 |

*Cpd.* **49***: Mechanistic analysis*

| parameter | unit | "true" (T) | calculated (C) | C/T ratio | note |
|---|---|---|---|---|---|
| mechanism | | **C2S** | **C2F** | | |
| $k_{eff}$ | mM$^{-1}$s$^{-1}$ | 0.9999 | 0.8757 | 0.88 | $= k_{inact}/K_i$ |
| $K_i$ | nM | 100.01 | 99.43 | 0.99 | |
| $k_{inact}$ | s$^{-1}$ | 0.0001 | 8.707e-005 | 0.87 | |

| | |
|---|---|
| Reaction times used for analysis: | 30 and 120 min |
| Maximum GSD for accepting one-step model **C1**: | 1.25 |
| Observed GSD: | 2.29 |
| Assumed $[S]_0/K_M$ ratio: | 1.0 |

*2.50. Compound No. 50*

*Cpd.* **50***: Simulated data*

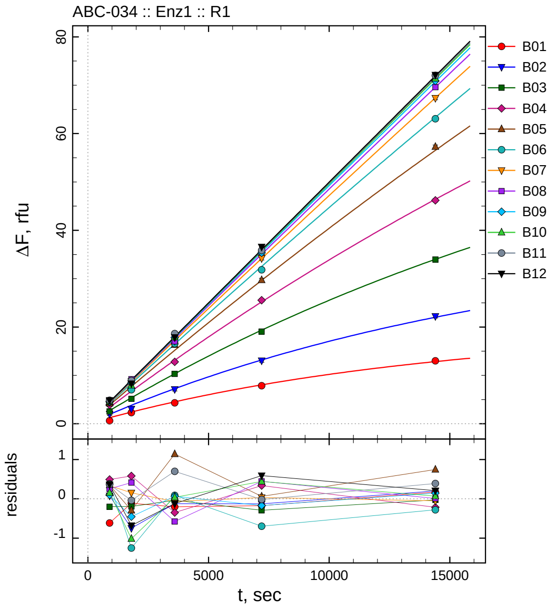

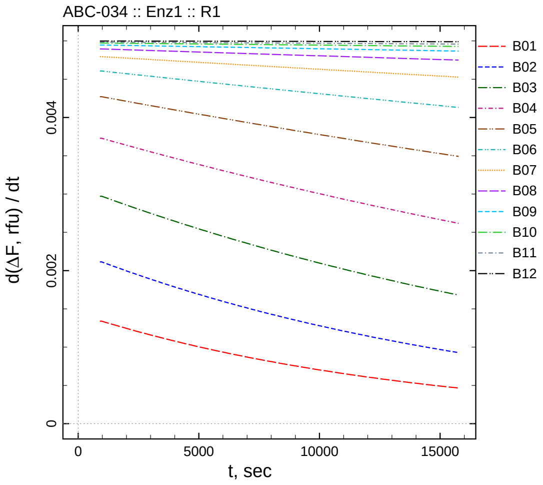

| $[I]_0$, nM | t = 15 | 30 | 60 | 120 | 240 min |
|---|---|---|---|---|---|
| 5000.5 | 0.629 | 2.301 | 4.331 | 7.866 | 13.016 |
| 2500.25 | 2.031 | 3.061 | 7.126 | 13.055 | 22.216 |
| 1250.12 | 2.522 | 5.157 | 10.306 | 19.053 | 33.971 |
| 625.062 | 3.885 | 7.294 | 12.792 | 25.534 | 46.190 |
| 312.531 | 4.217 | 7.395 | 16.330 | 29.717 | 57.283 |
| 156.266 | 4.540 | 7.041 | 16.564 | 31.824 | 63.081 |
| 78.1328 | 4.660 | 8.785 | 17.133 | 34.199 | 67.339 |
| 39.0664 | 4.663 | 9.222 | 17.011 | 35.497 | 69.602 |
| 19.5332 | 4.551 | 8.447 | 17.856 | 35.341 | 70.889 |
| 9.7666 | 4.631 | 7.935 | 17.931 | 36.188 | 71.402 |
| 4.8833 | 4.853 | 8.933 | 18.641 | 35.848 | 72.023 |
| 0 | 4.862 | 8.317 | 17.872 | 36.574 | 72.141 |

#### *Cpd.* **50**: *Determination of $IC_{50}$ and $k_1^*$*

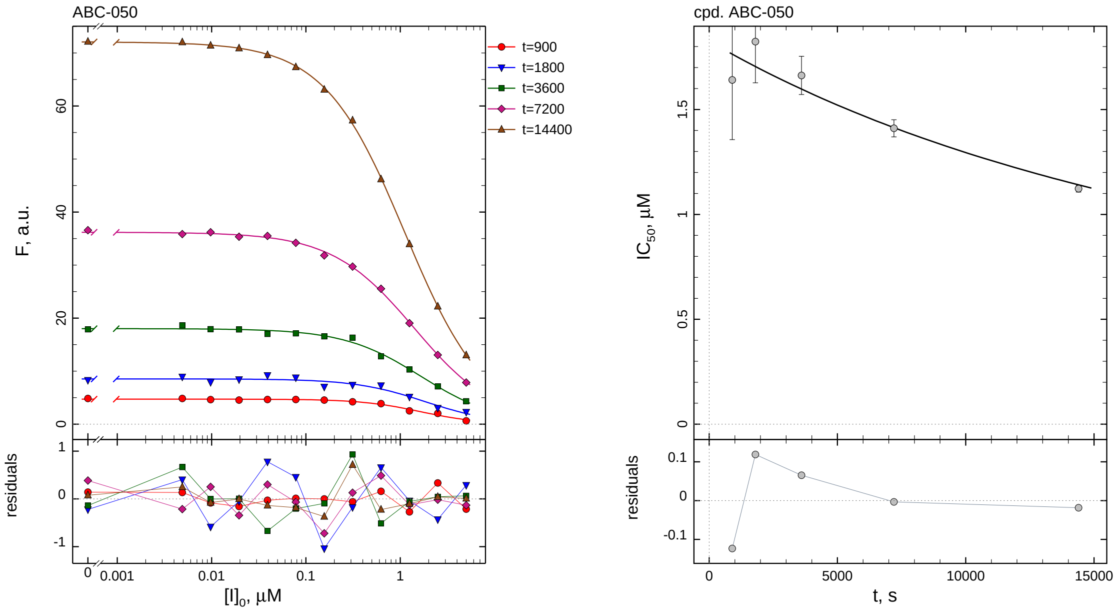

| $i$ | $t$, min | $I_{50}$, nM ± std.err. | CV, % | $k_1^*$, $mM^{-1}s^{-1}$ |
|---|---|---|---|---|
| 1 | 15 | 1640.7 ± 284.2 | 17.3 | 1.079 |
| 2 | 30 | 1823.8 ± 196.5 | 10.8 | 0.4854 |
| 3 | 60 | 1662.6 ± 91.12 | 5.5 | 0.2663 |
| 4 | 120 | 1410.4 ± 40.53 | 2.9 | 0.1569 |
| 5 | 240 | 1122.7 ± 16.05 | 1.4 | 0.09857 |

#### *Cpd.* **50**: *Mechanistic analysis*

| parameter | unit | "true" (T) | calculated (C) | C/T ratio | note |
|---|---|---|---|---|---|
| mechanism | | **C2S** | **C2F** | | |
| $k_{eff}$ | $mM^{-1}s^{-1}$ | 0.09999 | 0.1059 | 1.06 | $= k_{inact}/K_i$ |
| $K_i$ | nM | 1000.1 | 1010.7 | 1.01 | |
| $k_{inact}$ | $s^{-1}$ | 0.0001 | 0.0001071 | 1.07 | |

| | |
|---|---|
| Reaction times used for analysis: | 30 and 120 min |
| Maximum GSD for accepting one-step model **C1**: | 1.25 |
| Observed GSD: | 2.22 |
| Assumed $[S]_0/K_M$ ratio: | 1.0 |

*2.51. Compound No. 51*

*Cpd.* **51***: Simulated data*

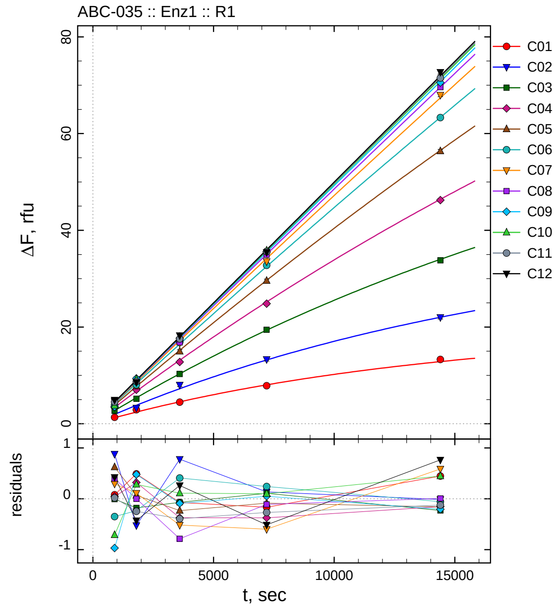

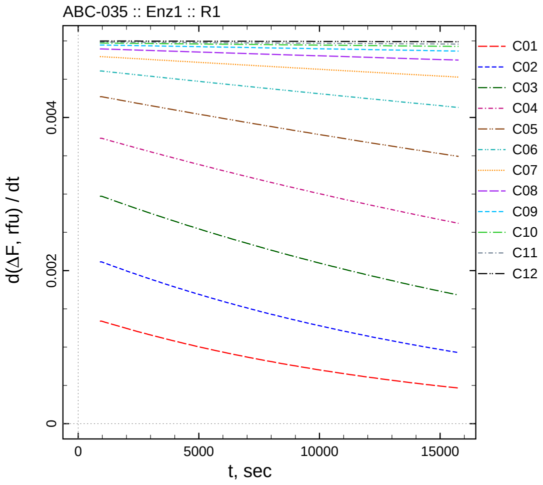

| $[I]_0$, nM | t = 15 | 30 | 60 | 120 | 240 min |
|---|---|---|---|---|---|
| 50005 | 1.325 | 2.901 | 4.463 | 7.872 | 13.293 |
| 25002.5 | 2.830 | 3.284 | 8.028 | 13.319 | 22.003 |
| 12501.3 | 2.715 | 5.172 | 10.273 | 19.453 | 33.769 |
| 6250.63 | 3.435 | 7.032 | 12.767 | 24.822 | 46.253 |
| 3125.31 | 4.490 | 7.755 | 14.959 | 29.568 | 56.380 |
| 1562.66 | 3.810 | 8.057 | 16.884 | 32.762 | 63.303 |
| 781.328 | 4.618 | 8.740 | 16.685 | 33.567 | 67.977 |
| 390.664 | 4.806 | 8.810 | 16.804 | 34.941 | 69.599 |
| 195.332 | 3.486 | 9.382 | 17.699 | 35.561 | 70.540 |
| 97.666 | 3.766 | 9.231 | 18.003 | 35.845 | 71.780 |
| 48.833 | 4.505 | 8.728 | 17.549 | 35.596 | 71.509 |
| 0 | 4.922 | 8.569 | 18.257 | 35.471 | 72.698 |

*Cpd.* **51***: Determination of* $IC_{50}$ *and* $k_1^*$

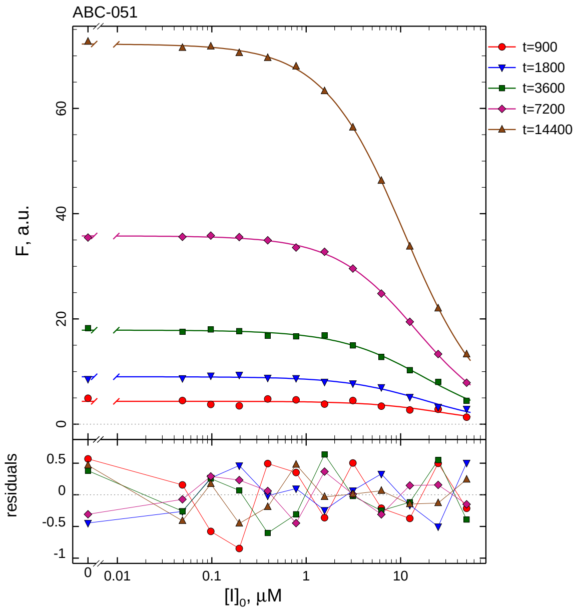

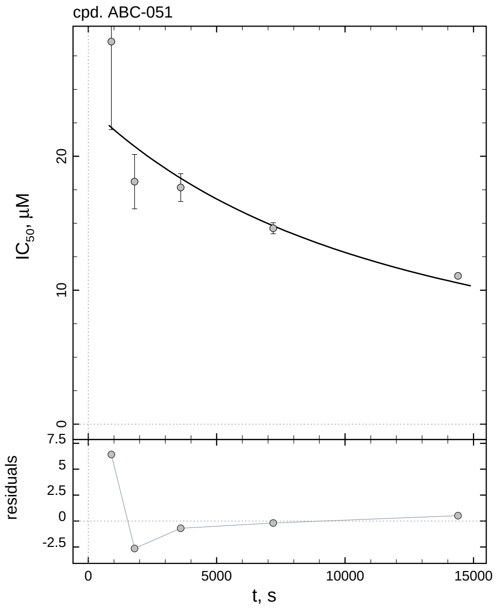

| $i$ | $t$, min | $I_{50}$, nM | ± | std.err. | CV, % | $k_1^*$, mM$^{-1}$s$^{-1}$ |
|---|---|---|---|---|---|---|
| 1 | 15 | 28564 | ± | 6573.0 | 23.0 | 0.06199 |
| 2 | 30 | 18108 | ± | 2029.7 | 11.2 | 0.04889 |
| 3 | 60 | 17668 | ± | 1043.0 | 5.9 | 0.02505 |
| 4 | 120 | 14628 | ± | 408.4 | 2.8 | 0.01513 |
| 5 | 240 | 11069 | ± | 154.9 | 1.4 | 0.009998 |

*Cpd.* **51***: Mechanistic analysis*

| parameter | unit | "true" (T) | calculated (C) | C/T ratio | note |
|---|---|---|---|---|---|
| mechanism | | **C2S** | **C2F** | | |
| $k_{\mathrm{eff}}$ | mM$^{-1}$s$^{-1}$ | 0.009999 | 0.008704 | 0.87 | $= k_{\mathrm{inact}}/K_{\mathrm{i}}$ |
| $K_{\mathrm{i}}$ | nM | 10001 | 9833.9 | 0.98 | |
| $k_{\mathrm{inact}}$ | s$^{-1}$ | 0.0001 | 8.56e-005 | 0.86 | |

| | |
|---|---|
| Reaction times used for analysis: | 30 and 120 min |
| Maximum GSD for accepting one-step model **C1**: | 1.25 |
| Observed GSD: | 2.29 |
| Assumed $[S]_0/K_M$ ratio: | 1.0 |

*2.52. Compound No. 52*

*Cpd.* ***52****: Simulated data*

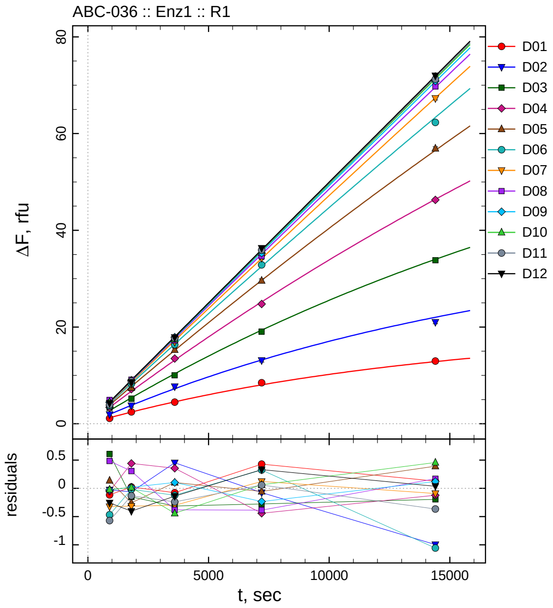

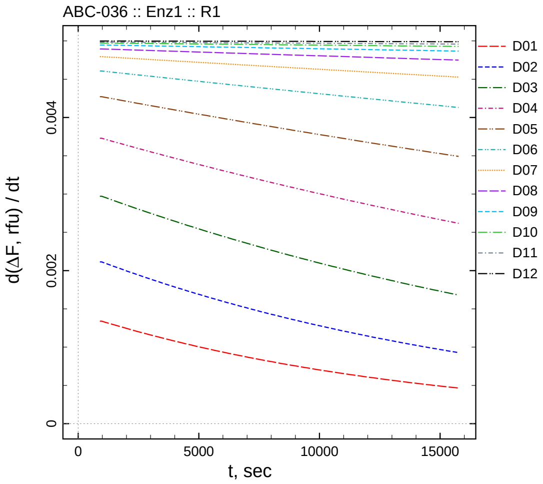

| $[I]_0$, nM | t = 15 | 30 | 60 | 120 | 240 min |
|---|---|---|---|---|---|
| 500050 | 1.130 | 2.437 | 4.457 | 8.469 | 12.974 |
| 250025 | 1.924 | 3.744 | 7.702 | 13.108 | 21.021 |
| 125013 | 3.328 | 5.198 | 10.028 | 19.068 | 33.804 |
| 62506.3 | 3.342 | 7.151 | 13.496 | 24.754 | 46.283 |
| 31253.1 | 4.004 | 7.433 | 15.294 | 29.601 | 56.929 |
| 15626.6 | 3.692 | 8.274 | 16.354 | 32.845 | 62.304 |
| 7813.28 | 3.999 | 8.314 | 16.900 | 34.286 | 67.299 |
| 3906.64 | 4.892 | 9.114 | 17.209 | 34.664 | 69.761 |
| 1953.32 | 4.427 | 8.924 | 17.892 | 35.277 | 70.863 |
| 976.66 | 4.455 | 8.964 | 17.450 | 35.807 | 71.791 |
| 488.33 | 3.915 | 8.842 | 17.703 | 35.922 | 71.266 |
| 0 | 4.241 | 8.592 | 17.855 | 36.312 | 71.969 |

*Cpd.* **52***: Determination of* $IC_{50}$ *and* $k_1^*$

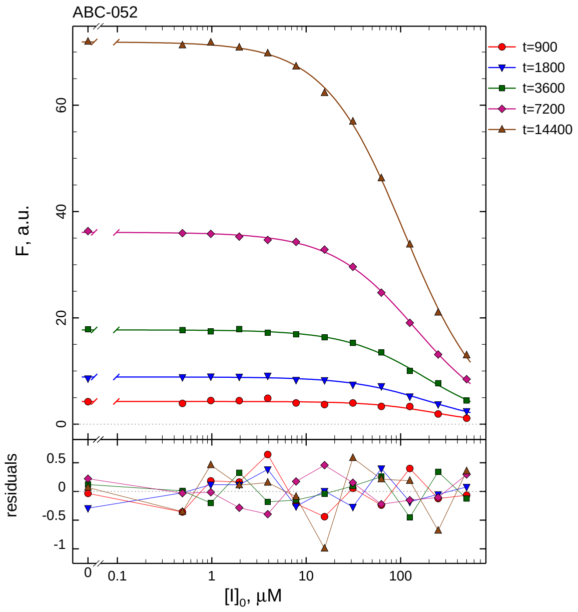

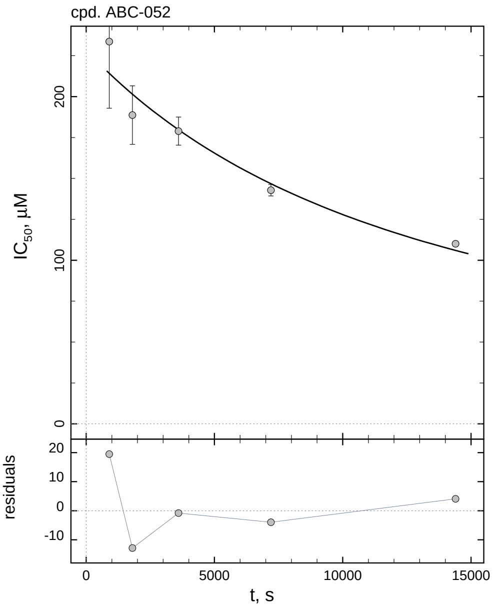

| $i$ | $t$, min | $I_{50}$, nM | ± | std.err. | CV, % | $k_1^*$, $mM^{-1}s^{-1}$ |
|---|---|---|---|---|---|---|
| 1 | 15 | 233656 | ± | 40729 | 17.4 | 0.007578 |
| 2 | 30 | 188690 | ± | 17887 | 9.5 | 0.004692 |
| 3 | 60 | 178925 | ± | 8588.8 | 4.8 | 0.002474 |
| 4 | 120 | 142807 | ± | 3489.9 | 2.4 | 0.001550 |
| 5 | 240 | 110090 | ± | 1311.6 | 1.2 | 0.001005 |

*Cpd.* **52***: Mechanistic analysis*

| parameter | unit | "true" (T) | calculated (C) | C/T ratio | note |
|---|---|---|---|---|---|
| mechanism | | **C2S** | **C2F** | | |
| $k_{eff}$ | $mM^{-1}s^{-1}$ | 0.0009999 | 0.001120 | 1.12 | $= k_{inact}/K_i$ |
| $K_i$ | nM | 100010 | 105661 | 1.06 | |
| $k_{inact}$ | $s^{-1}$ | 0.0001 | 0.0001183 | 1.18 | |

| | |
|---|---|
| Reaction times used for analysis: | 30 and 120 min |
| Maximum GSD for accepting one-step model **C1**: | 1.25 |
| Observed GSD: | 2.19 |
| Assumed $[S]_0/K_M$ ratio: | 1.0 |

*2.53. Compound No. 53*

*Cpd.* **53***: Simulated data*

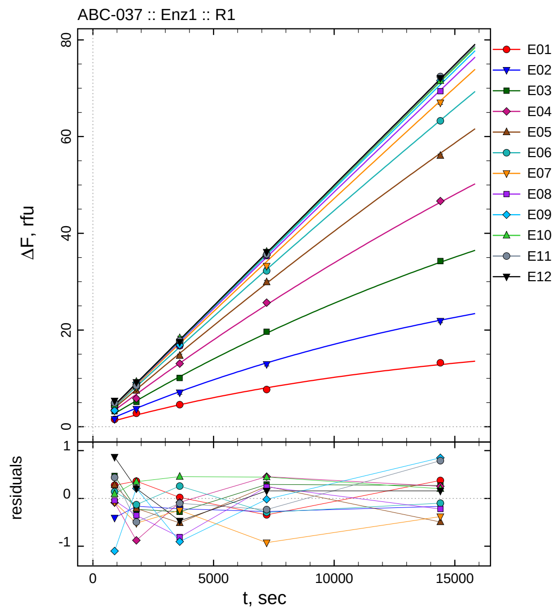

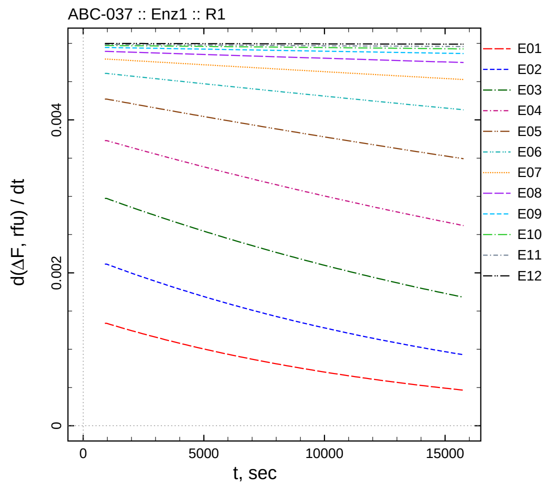

| $[I]_0$, nM | t = 15 | 30 | 60 | 120 | 240 min |
|---|---|---|---|---|---|
| 50.05 | 1.530 | 2.783 | 4.563 | 7.702 | 13.228 |
| 25.025 | 1.556 | 3.653 | 7.044 | 12.920 | 21.859 |
| 12.5125 | 3.204 | 5.136 | 10.071 | 19.649 | 34.282 |
| 6.25625 | 3.313 | 5.842 | 13.067 | 25.664 | 46.674 |
| 3.12812 | 4.154 | 7.484 | 14.686 | 29.916 | 56.057 |
| 1.56406 | 4.304 | 8.164 | 16.738 | 32.234 | 63.269 |
| 0.782031 | 4.241 | 8.116 | 16.956 | 33.242 | 67.015 |
| 0.391016 | 4.368 | 8.456 | 16.782 | 35.294 | 69.373 |
| 0.195508 | 3.354 | 9.114 | 16.887 | 35.494 | 71.596 |
| 0.0977539 | 4.572 | 9.300 | 18.348 | 36.191 | 71.539 |
| 0.048877 | 4.925 | 8.481 | 17.840 | 35.630 | 72.424 |
| 0 | 5.364 | 9.205 | 17.526 | 36.140 | 72.091 |

*Cpd.* **53***: Determination of* $IC_{50}$ *and* $k_1^*$

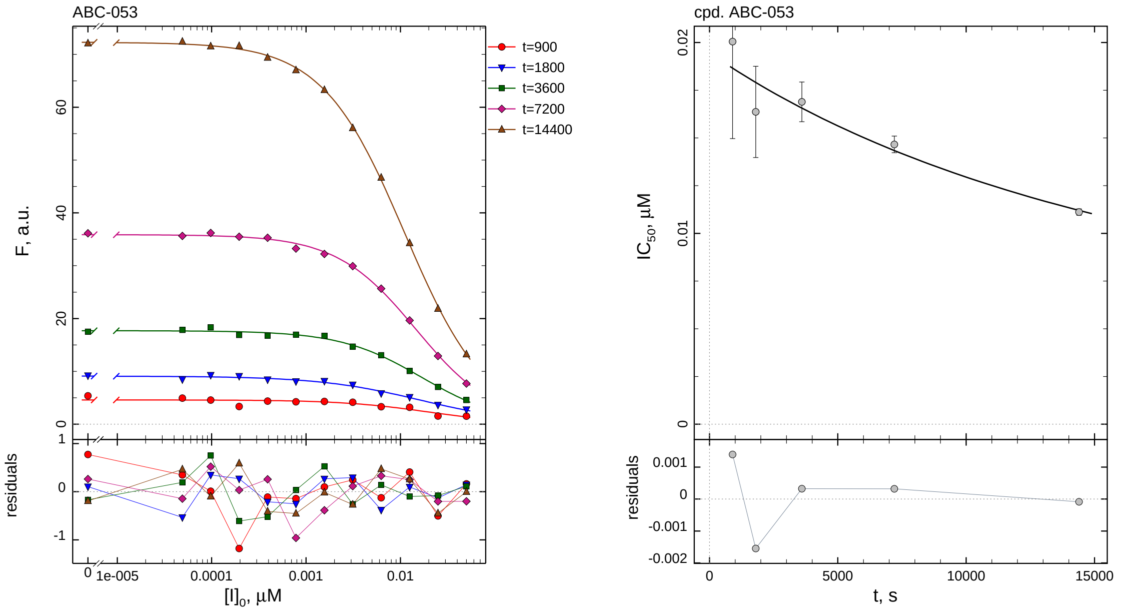

| $i$ | $t$, min | $I_{50}$, nM ± std.err. | CV, % | $k_1^*$, $mM^{-1}s^{-1}$ |
|---|---|---|---|---|
| 1 | 15 | 20.05 ± 5.081 | 25.3 | 88.31 |
| 2 | 30 | 16.37 ± 2.390 | 14.6 | 54.10 |
| 3 | 60 | 16.89 ± 1.041 | 6.2 | 26.21 |
| 4 | 120 | 14.66 ± 0.4360 | 3.0 | 15.10 |
| 5 | 240 | 11.12 ± 0.1698 | 1.5 | 9.954 |

*Cpd.* **53***: Mechanistic analysis*

| parameter | unit | "true" (T) | calculated (C) | C/T ratio | note |
|---|---|---|---|---|---|
| mechanism | | **C2S** | **C2F** | | |
| $k_{eff}$ | $mM^{-1}s^{-1}$ | 9.990 | 4.789 | 0.48 | $= k_{inact}/K_i$ |
| $K_i$ | nM | 10.01 | 8.513 | 0.85 | |
| $k_{inact}$ | $s^{-1}$ | 0.0001 | 4.077e-005 | 0.41 | |

| | |
|---|---|
| Reaction times used for analysis: | 30 and 120 min |
| Maximum GSD for accepting one-step model **C1**: | 1.25 |
| Observed GSD: | 2.47 |
| Assumed $[S]_0/K_M$ ratio: | 1.0 |

*2.54. Compound No. 54*

*Cpd.* **54***: Simulated data*

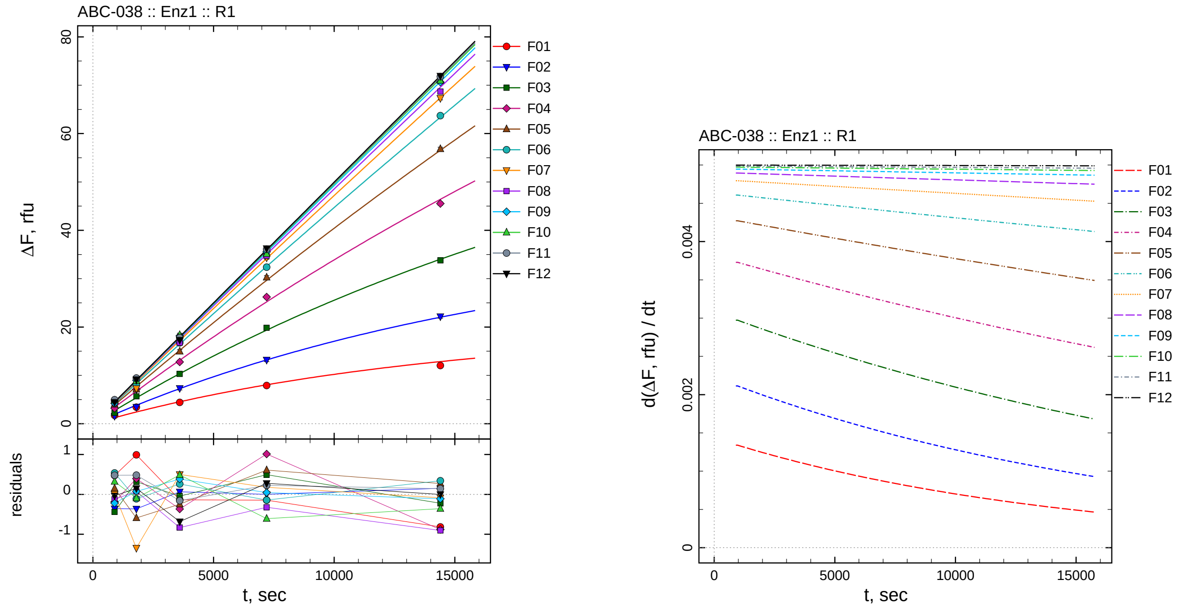

| $[I]_0$, nM | t = 15 | 30 | 60 | 120 | 240 min |
|---:|---:|---:|---:|---:|---:|
| 500.5 | 1.729 | 3.412 | 4.407 | 7.901 | 12.035 |
| 250.25 | 1.605 | 3.454 | 7.327 | 13.195 | 22.177 |
| 125.125 | 2.294 | 5.680 | 10.310 | 19.849 | 33.789 |
| 62.5625 | 3.212 | 7.115 | 12.783 | 26.217 | 45.536 |
| 31.2812 | 4.023 | 7.102 | 14.951 | 30.271 | 56.820 |
| 15.6406 | 4.700 | 8.185 | 16.739 | 32.382 | 63.707 |
| 7.82031 | 4.331 | 7.289 | 17.703 | 34.345 | 67.310 |
| 3.91016 | 4.286 | 8.883 | 16.758 | 34.729 | 68.689 |
| 1.95508 | 4.233 | 8.976 | 18.173 | 35.556 | 70.635 |
| 0.977539 | 4.797 | 8.858 | 18.380 | 35.137 | 70.977 |
| 0.48877 | 4.967 | 9.456 | 17.787 | 36.090 | 71.777 |
| 0 | 4.463 | 9.148 | 17.311 | 36.260 | 71.933 |

*Cpd.* **54***: Determination of* $IC_{50}$ *and* $k_1^*$

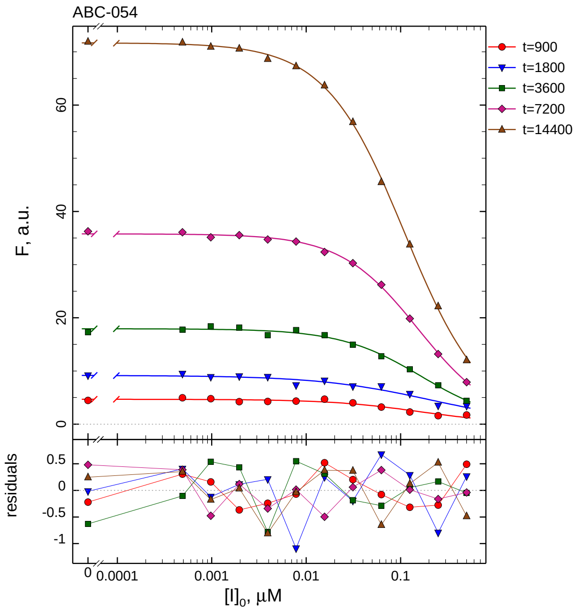

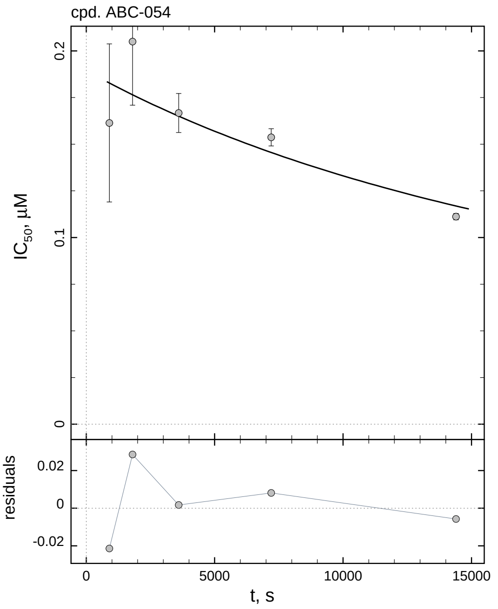

| $i$ | $t$, min | $I_{50}$, nM | ± | std.err. | CV, % | $k_1^*$, $mM^{-1}s^{-1}$ |
|---|---|---|---|---|---|---|
| 1 | 15 | 161.4 | ± | 42.31 | 26.2 | 10.97 |
| 2 | 30 | 205.0 | ± | 34.11 | 16.6 | 4.319 |
| 3 | 60 | 166.7 | ± | 10.44 | 6.3 | 2.655 |
| 4 | 120 | 153.7 | ± | 4.616 | 3.0 | 1.440 |
| 5 | 240 | 111.2 | ± | 1.715 | 1.5 | 0.9952 |

*Cpd.* **54***: Mechanistic analysis*

| parameter | unit | “true” (T) | calculated (C) | C/T ratio | note |
|---|---|---|---|---|---|
| mechanism | | **C2S** | **C2F** | | |
| $k_{eff}$ | $mM^{-1}s^{-1}$ | 0.9990 | 1.070 | 1.07 | $= k_{inact}/K_i$ |
| $K_i$ | nM | 100.1 | 115.3 | 1.15 | |
| $k_{inact}$ | $s^{-1}$ | 0.0001 | 0.0001234 | 1.23 | |

| | |
|---|---|
| Reaction times used for analysis: | 30 and 120 min |
| Maximum GSD for accepting one-step model **C1**: | 1.25 |
| Observed GSD: | 2.17 |
| Assumed $[S]_0/K_M$ ratio: | 1.0 |

*2.55. Compound No. 55*

*Cpd.* **55**: *Simulated data*

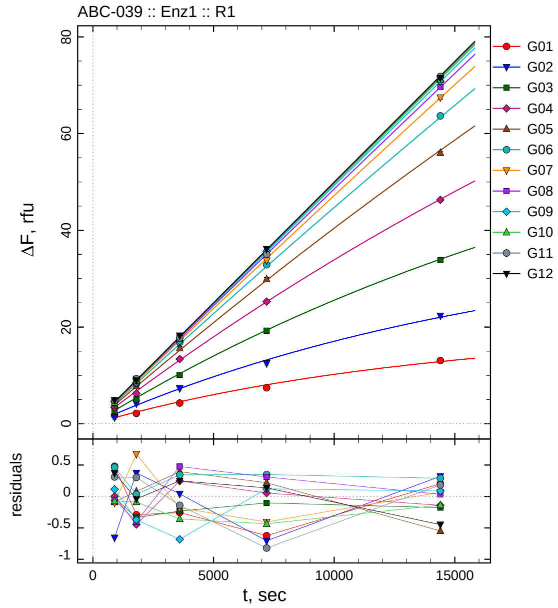

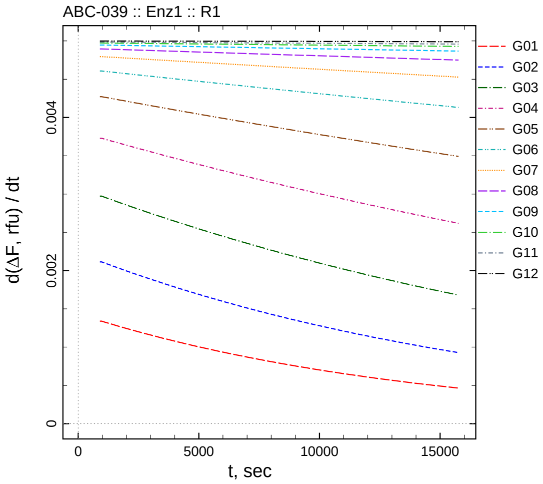

| $[I]_0$, nM | t = 15 | 30 | 60 | 120 | 240 min |
|---|---|---|---|---|---|
| 5005 | 1.735 | 2.131 | 4.287 | 7.423 | 13.053 |
| 2502.5 | 1.306 | 4.192 | 7.303 | 12.489 | 22.346 |
| 1251.25 | 2.712 | 5.026 | 10.123 | 19.254 | 33.835 |
| 625.625 | 3.402 | 6.274 | 13.397 | 25.262 | 46.278 |
| 312.812 | 3.786 | 7.774 | 15.592 | 29.881 | 56.000 |
| 156.406 | 4.628 | 8.330 | 16.823 | 32.873 | 63.658 |
| 78.2031 | 4.224 | 9.307 | 17.022 | 33.762 | 67.462 |
| 39.1016 | 4.359 | 8.409 | 18.067 | 35.362 | 69.629 |
| 19.5508 | 4.569 | 8.532 | 17.109 | 35.635 | 70.834 |
| 9.77539 | 4.403 | 8.863 | 17.534 | 35.311 | 71.198 |
| 4.8877 | 4.797 | 9.277 | 17.802 | 35.044 | 71.816 |
| 0 | 4.871 | 8.959 | 18.242 | 36.119 | 71.486 |

*Cpd.* **55***: Determination of* $IC_{50}$ *and* $k_1^*$

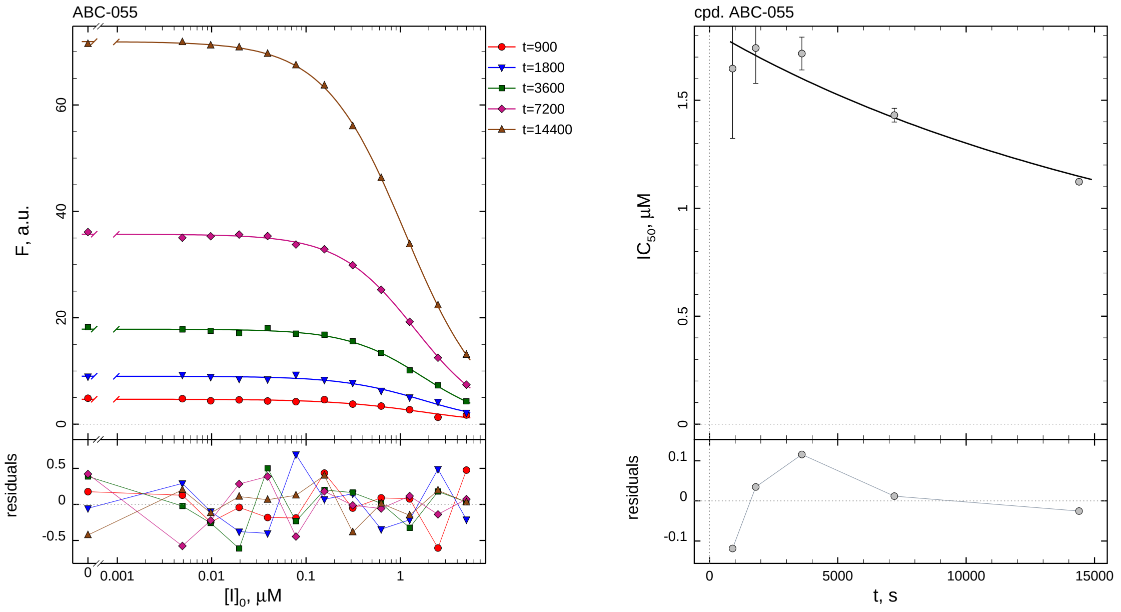

| $i$ | $t$, min | $I_{50}$, nM | ± | std.err. | CV, % | $k_1^*$, $mM^{-1}s^{-1}$ |
|---|---|---|---|---|---|---|
| 1 | 15 | 1646.4 | ± | 323.0 | 19.6 | 1.075 |
| 2 | 30 | 1742.0 | ± | 164.2 | 9.4 | 0.5082 |
| 3 | 60 | 1716.3 | ± | 75.88 | 4.4 | 0.2579 |
| 4 | 120 | 1430.9 | ± | 31.78 | 2.2 | 0.1547 |
| 5 | 240 | 1122.4 | ± | 13.12 | 1.2 | 0.09860 |

*Cpd.* **55***: Mechanistic analysis*

| parameter | unit | "true" (T) | calculated (C) | C/T ratio | note |
|---|---|---|---|---|---|
| mechanism | | **C2S** | **C2F** | | |
| $k_{\mathrm{eff}}$ | $mM^{-1}s^{-1}$ | 0.09990 | 0.08289 | 0.83 | $= k_{\mathrm{inact}}/K_{\mathrm{i}}$ |
| $K_{\mathrm{i}}$ | nM | 1001 | 939.1 | 0.94 | |
| $k_{\mathrm{inact}}$ | $s^{-1}$ | 0.0001 | 7.784e-005 | 0.78 | |

| | |
|---|---|
| Reaction times used for analysis: | 30 and 120 min |
| Maximum GSD for accepting one-step model **C1**: | 1.25 |
| Observed GSD: | 2.32 |
| Assumed $[S]_0/K_M$ ratio: | 1.0 |

*Cpd.* **56***: Simulated data*

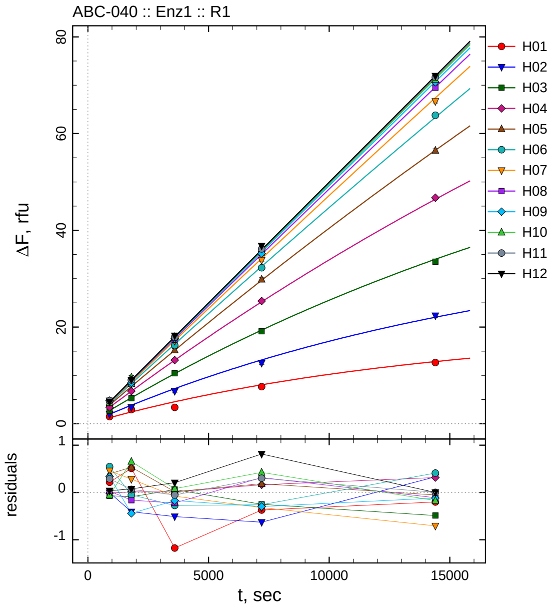

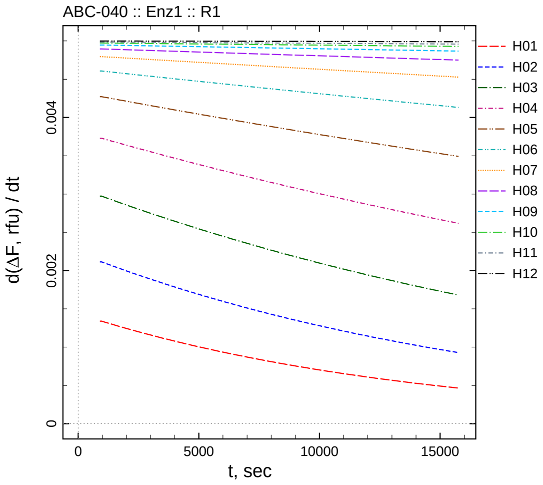

| $[I]_0$, nM | t = 15 | 30 | 60 | 120 | 240 min |
|---|---|---|---|---|---|
| 50050 | 1.471 | 2.931 | 3.367 | 7.674 | 12.650 |
| 25025 | 1.956 | 3.409 | 6.749 | 12.565 | 22.355 |
| 12512.5 | 2.665 | 5.269 | 10.414 | 19.106 | 33.523 |
| 6256.25 | 3.386 | 6.733 | 13.162 | 25.371 | 46.734 |
| 3128.12 | 4.268 | 8.229 | 15.215 | 29.848 | 56.494 |
| 1564.06 | 4.709 | 8.244 | 16.204 | 32.264 | 63.776 |
| 782.031 | 4.784 | 8.915 | 17.127 | 33.849 | 66.687 |
| 391.016 | 4.437 | 8.647 | 17.368 | 35.368 | 69.482 |
| 195.508 | 4.790 | 8.461 | 17.618 | 35.212 | 70.627 |
| 97.7539 | 4.417 | 9.604 | 17.971 | 36.172 | 71.162 |
| 48.877 | 4.776 | 9.041 | 17.891 | 36.164 | 71.635 |
| 0 | 4.534 | 9.073 | 18.202 | 36.795 | 71.933 |

*Cpd.* **56**: *Determination of* $IC_{50}$ *and* $k_1^*$

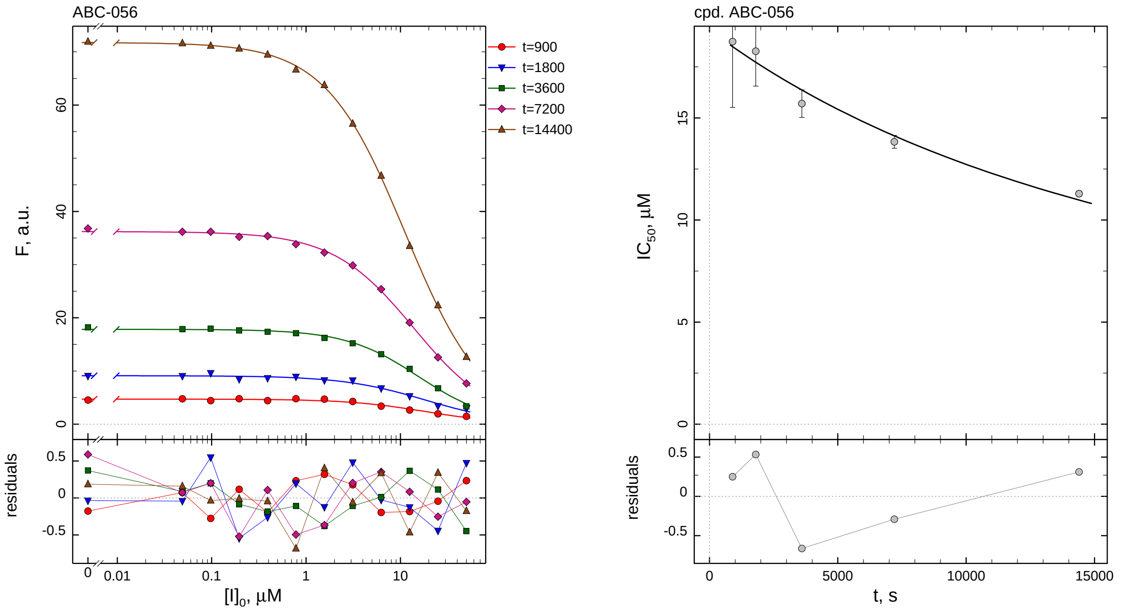

| $i$ | $t$, min | $I_{50}$, nM | ± | std.err. | CV, % | $k_1^*$, $mM^{-1}s^{-1}$ |
|---|---|---|---|---|---|---|
| 1 | 15 | 18740 | ± | 3219.5 | 17.2 | 0.09449 |
| 2 | 30 | 18265 | ± | 1709.0 | 9.4 | 0.04847 |
| 3 | 60 | 15702 | ± | 674.4 | 4.3 | 0.02819 |
| 4 | 120 | 13831 | ± | 317.1 | 2.3 | 0.01600 |
| 5 | 240 | 11290 | ± | 130.2 | 1.2 | 0.009802 |

*Cpd.* **56**: *Mechanistic analysis*

| parameter | unit | "true" (T) | calculated (C) | C/T ratio | note |
|---|---|---|---|---|---|
| mechanism | | **C2S** | **C2F** | | |
| $k_{eff}$ | $mM^{-1}s^{-1}$ | 0.009990 | 0.01155 | 1.16 | $= k_{inact}/K_i$ |
| $K_i$ | nM | 10010 | 10225 | 1.02 | |
| $k_{inact}$ | $s^{-1}$ | 0.0001 | 0.0001181 | 1.18 | |

| | |
|---|---|
| Reaction times used for analysis: | 30 and 120 min |
| Maximum GSD for accepting one-step model **C1**: | 1.25 |
| Observed GSD: | 2.19 |
| Assumed $[S]_0/K_M$ ratio: | 1.0 |

*2.57. Compound No. 57*

*Cpd.* ***57****: Simulated data*

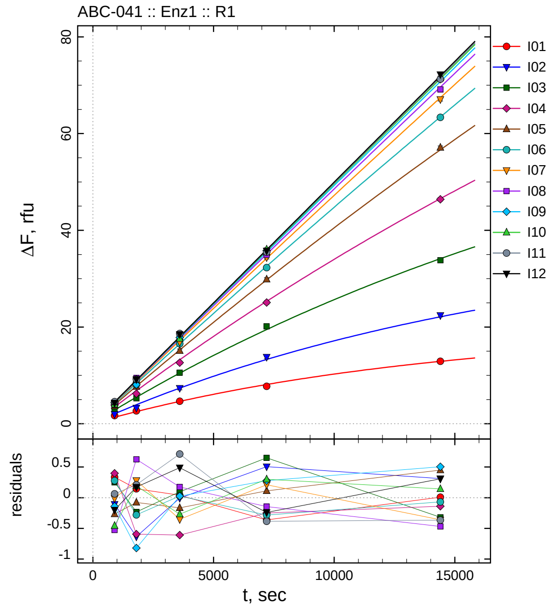

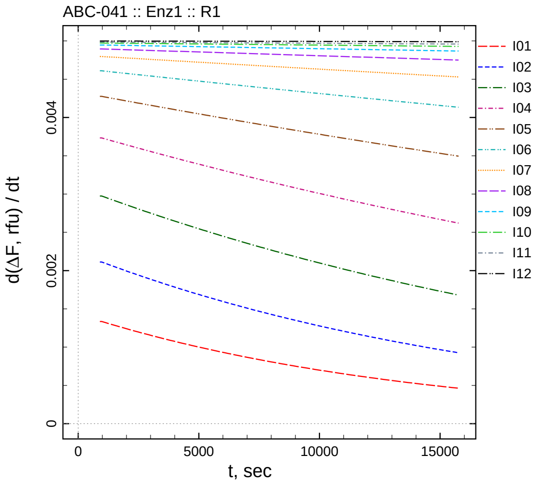

| $[I]_0$, nM | t = 15 | 30 | 60 | 120 | 240 min |
|---:|---:|---:|---:|---:|---:|
| 5.05 | 1.678 | 2.651 | 4.662 | 7.753 | 12.907 |
| 2.525 | 1.966 | 3.291 | 7.360 | 13.799 | 22.423 |
| 1.2625 | 3.094 | 5.239 | 10.565 | 20.125 | 33.815 |
| 0.63125 | 3.883 | 6.214 | 12.636 | 25.066 | 46.408 |
| 0.315625 | 3.663 | 7.675 | 15.099 | 29.853 | 57.101 |
| 0.157813 | 4.473 | 8.050 | 16.545 | 32.290 | 63.366 |
| 0.0789063 | 4.331 | 8.933 | 16.872 | 34.407 | 67.074 |
| 0.0394531 | 3.892 | 9.446 | 17.777 | 34.922 | 69.146 |
| 0.0197266 | 4.318 | 8.089 | 17.823 | 35.798 | 71.261 |
| 0.00986328 | 4.021 | 9.144 | 17.626 | 36.050 | 71.484 |
| 0.00493164 | 4.552 | 9.186 | 18.656 | 35.480 | 71.270 |
| 0 | 4.299 | 9.172 | 18.484 | 35.743 | 72.242 |

*Cpd.* **57***: Determination of* $IC_{50}$ *and* $k_1^*$

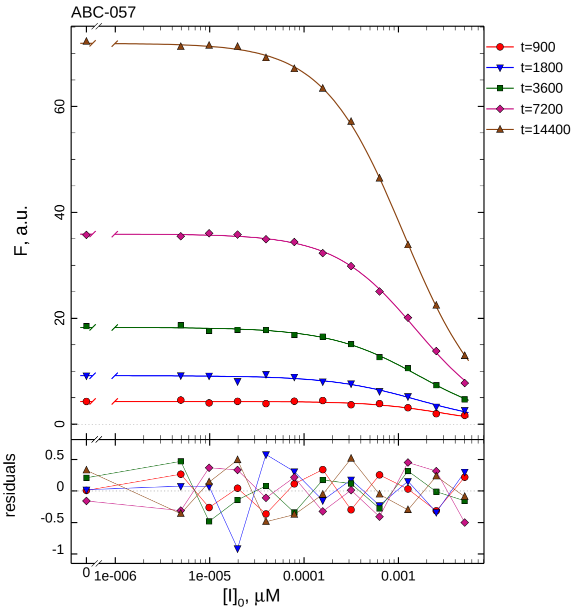

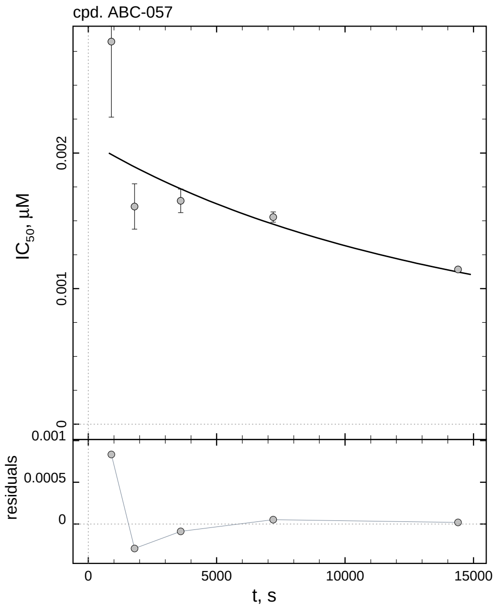

| $i$ | $t$, min | $I_{50}$, nM | ± | std.err. | CV, % | $k_1^*$, $mM^{-1}s^{-1}$ |
|---|---|---|---|---|---|---|
| 1 | 15 | 2.822 | ± | 0.5573 | 19.7 | 627.5 |
| 2 | 30 | 1.606 | ± | 0.1670 | 10.4 | 551.4 |
| 3 | 60 | 1.647 | ± | 0.08675 | 5.3 | 268.7 |
| 4 | 120 | 1.527 | ± | 0.03768 | 2.5 | 144.9 |
| 5 | 240 | 1.141 | ± | 0.01406 | 1.2 | 96.97 |

*Cpd.* **57***: Mechanistic analysis*

| parameter | unit | "true" (T) | calculated (C) | C/T ratio | note |
|---|---|---|---|---|---|
| mechanism | | **C2S** | **C2F** | | |
| $k_{eff}$ | $mM^{-1}s^{-1}$ | 99.01 | 21.86 | 0.22 | $= k_{inact}/K_i$ |
| $K_i$ | nM | 1.01 | 0.8167 | 0.81 | |
| $k_{inact}$ | $s^{-1}$ | 0.0001 | 1.785e-005 | 0.18 | |

| | |
|---|---|
| Reaction times used for analysis: | 30 and 120 min |
| Maximum GSD for accepting one-step model **C1**: | 1.25 |
| Observed GSD: | 2.57 |
| Assumed $[S]_0/K_M$ ratio: | 1.0 |

*2.58. Compound No. 58*

*Cpd.* **58***: Simulated data*

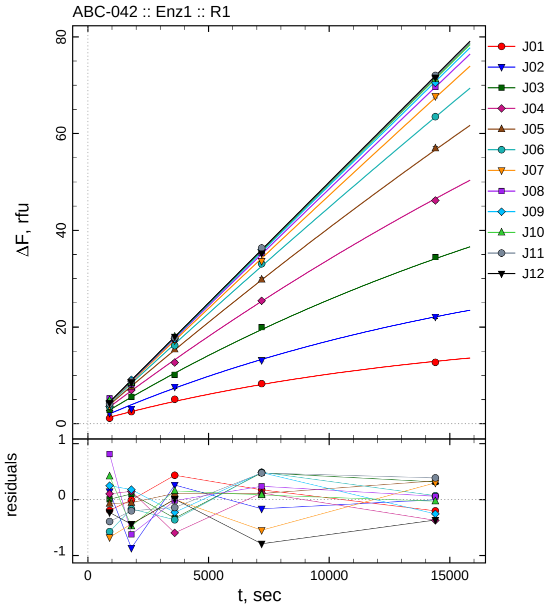

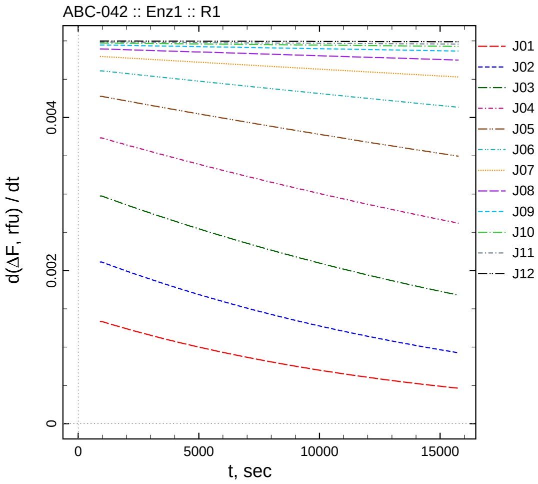

| $[I]_0$, nM | t = 15 | 30 | 60 | 120 | 240 min |
|---|---|---|---|---|---|
| 50.5 | 1.149 | 2.498 | 5.056 | 8.287 | 12.697 |
| 25.25 | 2.216 | 3.060 | 7.627 | 13.129 | 22.106 |
| 12.625 | 2.847 | 5.571 | 10.133 | 19.951 | 34.442 |
| 6.3125 | 3.587 | 6.956 | 12.649 | 25.430 | 46.168 |
| 3.15625 | 3.857 | 7.699 | 15.375 | 29.849 | 56.980 |
| 1.57813 | 3.621 | 8.165 | 16.158 | 33.061 | 63.501 |
| 0.789063 | 3.667 | 8.216 | 17.225 | 33.647 | 67.731 |
| 0.394531 | 5.233 | 8.200 | 17.583 | 35.305 | 69.674 |
| 0.197266 | 4.707 | 9.088 | 17.569 | 36.001 | 70.501 |
| 0.0986328 | 4.895 | 8.479 | 18.051 | 35.832 | 71.312 |
| 0.0493164 | 4.096 | 8.775 | 17.803 | 36.342 | 72.024 |
| 0 | 4.268 | 8.556 | 18.003 | 35.193 | 71.568 |

*Cpd.* **58***: Determination of* $IC_{50}$ *and* $k_1^*$

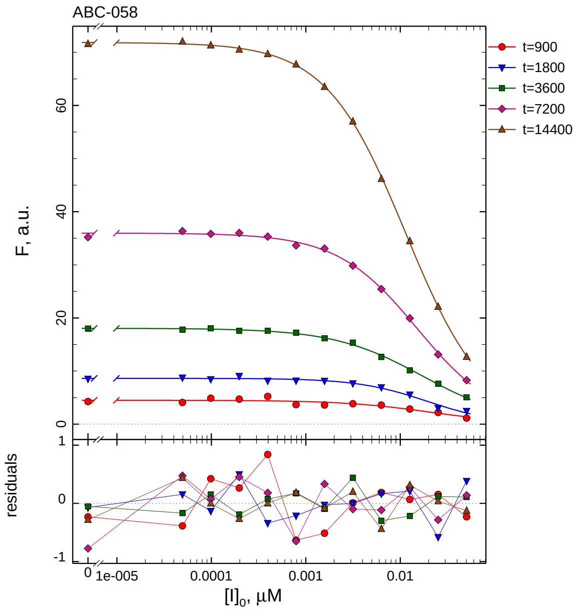

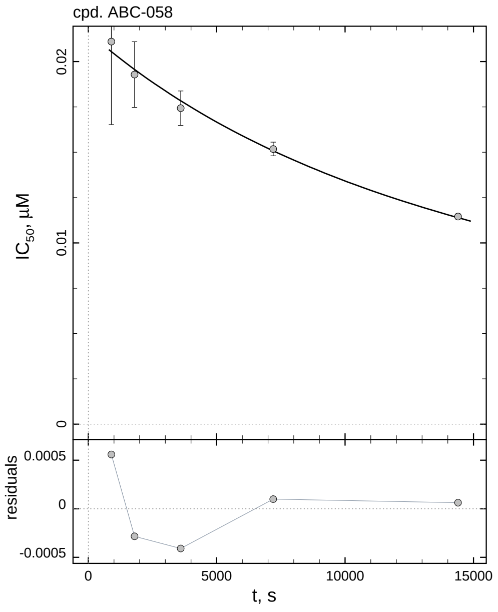

| $i$ | $t$, min | $I_{50}$, nM | ± | std.err. | CV, % | $k_1^*$, $mM^{-1}s^{-1}$ |
|---|---|---|---|---|---|---|
| 1 | 15 | 21.10 | ± | 4.581 | 21.7 | 83.90 |
| 2 | 30 | 19.29 | ± | 1.808 | 9.4 | 45.91 |
| 3 | 60 | 17.43 | ± | 0.9494 | 5.4 | 25.40 |
| 4 | 120 | 15.18 | ± | 0.3778 | 2.5 | 14.58 |
| 5 | 240 | 11.46 | ± | 0.1425 | 1.2 | 9.661 |

*Cpd.* **58***: Mechanistic analysis*

| parameter | unit | "true" (T) | calculated (C) | C/T ratio | note |
|---|---|---|---|---|---|
| mechanism | | **C2S** | **C2F** | | |
| $k_{eff}$ | $mM^{-1}s^{-1}$ | 9.901 | 9.266 | 0.94 | $= k_{inact}/K_i$ |
| $K_i$ | nM | 10.1 | 10.60 | 1.05 | |
| $k_{inact}$ | $s^{-1}$ | 0.0001 | 9.821e-005 | 0.98 | |

| | |
|---|---|
| Reaction times used for analysis: | 30 and 120 min |
| Maximum GSD for accepting one-step model **C1**: | 1.25 |
| Observed GSD: | 2.25 |
| Assumed $[S]_0/K_M$ ratio: | 1.0 |

*2.59. Compound No. 59*

*Cpd.* **59***: Simulated data*

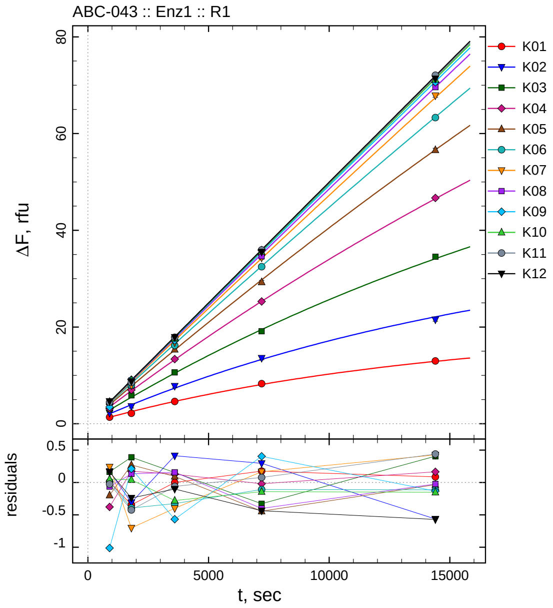

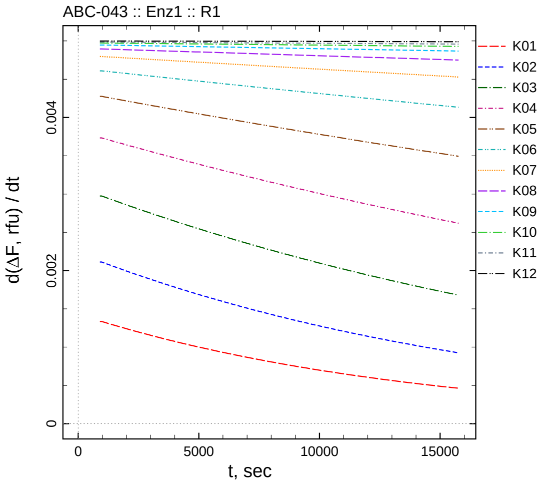

| $[I]_0$, nM | t = 15 | 30 | 60 | 120 | 240 min |
|---|---|---|---|---|---|
| 505 | 1.353 | 2.160 | 4.621 | 8.288 | 12.985 |
| 252.5 | 2.245 | 3.615 | 7.780 | 13.589 | 21.542 |
| 126.25 | 3.005 | 5.862 | 10.620 | 19.144 | 34.533 |
| 63.125 | 3.107 | 6.982 | 13.373 | 25.294 | 46.704 |
| 31.5625 | 3.731 | 8.026 | 15.357 | 29.296 | 56.611 |
| 15.7813 | 4.206 | 7.936 | 16.192 | 32.465 | 63.316 |
| 7.89063 | 4.583 | 7.949 | 16.827 | 34.362 | 67.857 |
| 3.94531 | 4.356 | 8.955 | 17.761 | 34.666 | 69.590 |
| 1.97266 | 3.447 | 9.120 | 17.228 | 35.924 | 70.624 |
| 0.986328 | 4.532 | 8.995 | 17.616 | 35.607 | 71.189 |
| 0.493164 | 4.464 | 8.552 | 17.884 | 35.944 | 72.076 |
| 0 | 4.661 | 8.758 | 17.895 | 35.547 | 71.363 |

*Cpd.* **59***: Determination of* $IC_{50}$ *and* $k_1^*$

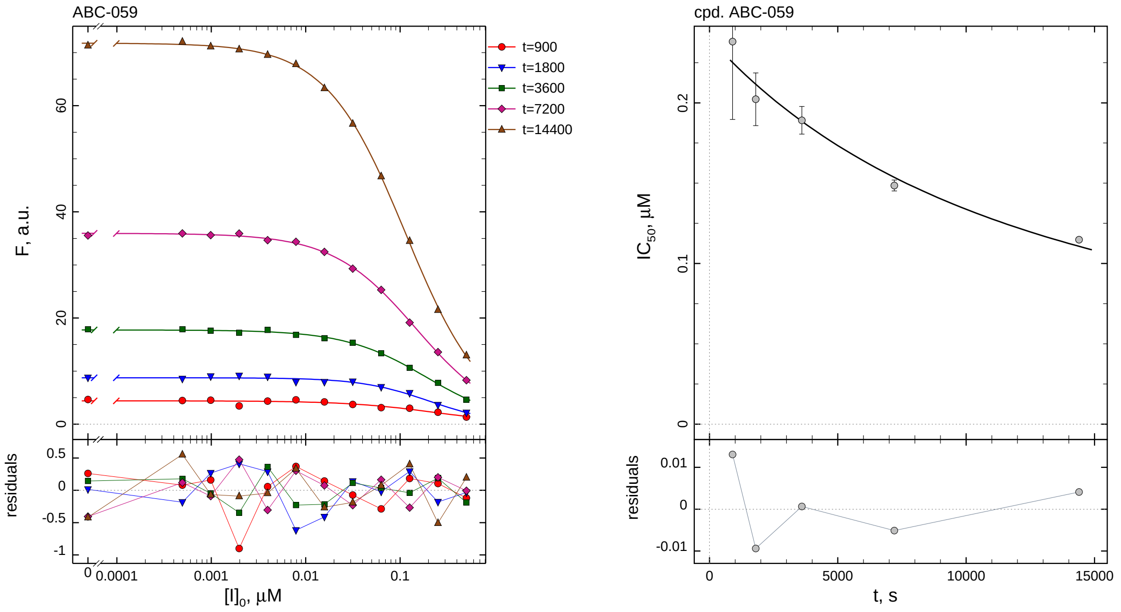

| $i$ | $t$, min | $I_{50}$, nM ± std.err. | CV, % | $k_1^*$, $mM^{-1}s^{-1}$ |
|---|---|---|---|---|
| 1 | 15 | 238.1 ± 48.41 | 20.3 | 7.436 |
| 2 | 30 | 202.2 ± 16.38 | 8.1 | 4.378 |
| 3 | 60 | 189.1 ± 8.612 | 4.6 | 2.340 |
| 4 | 120 | 148.5 ± 3.389 | 2.3 | 1.490 |
| 5 | 240 | 114.8 ± 1.268 | 1.1 | 0.9640 |

*Cpd.* **59***: Mechanistic analysis*

| parameter | unit | "true" (T) | calculated (C) | C/T ratio | note |
|---|---|---|---|---|---|
| mechanism | | **C2S** | **C2F** | | |
| $k_{eff}$ | $mM^{-1}s^{-1}$ | 0.9901 | 1.172 | 1.18 | $= k_{inact}/K_i$ |
| $K_i$ | nM | 101 | 115.0 | 1.14 | |
| $k_{inact}$ | $s^{-1}$ | 0.0001 | 0.0001348 | 1.35 | |

| | |
|---|---|
| Reaction times used for analysis: | 30 and 120 min |
| Maximum GSD for accepting one-step model **C1**: | 1.25 |
| Observed GSD: | 2.14 |
| Assumed $[S]_0/K_M$ ratio: | 1.0 |

*2.60. Compound No. 60*

*Cpd.* **60***: Simulated data*

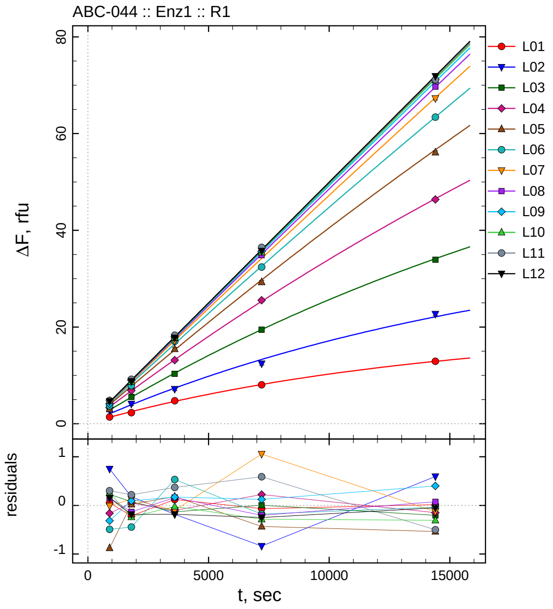

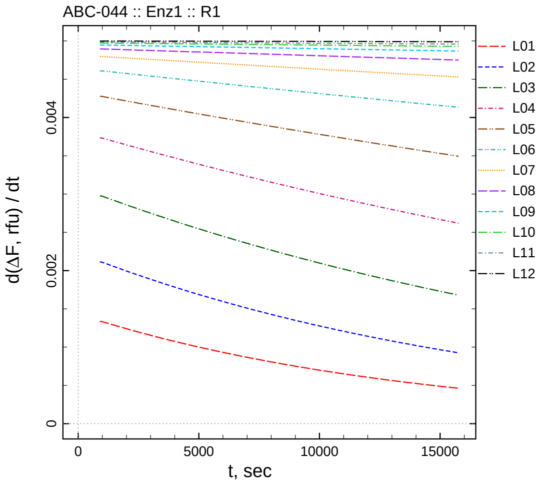

| $[I]_0$, nM | t = 15 | 30 | 60 | 120 | 240 min |
|---|---|---|---|---|---|
| 5050 | 1.396 | 2.291 | 4.742 | 8.046 | 12.913 |
| 2525 | 2.823 | 4.104 | 7.182 | 12.453 | 22.702 |
| 1262.5 | 3.070 | 5.545 | 10.333 | 19.475 | 33.922 |
| 631.25 | 3.324 | 6.858 | 13.142 | 25.535 | 46.388 |
| 315.625 | 3.052 | 7.776 | 15.445 | 29.305 | 56.106 |
| 157.812 | 3.702 | 7.884 | 17.051 | 32.393 | 63.392 |
| 78.9062 | 4.334 | 8.777 | 17.122 | 35.252 | 67.304 |
| 39.4531 | 4.576 | 8.688 | 17.765 | 34.862 | 69.689 |
| 19.7266 | 4.143 | 9.002 | 17.968 | 35.642 | 71.156 |
| 9.86328 | 4.738 | 8.714 | 17.872 | 35.465 | 71.035 |
| 4.93164 | 4.790 | 9.195 | 18.314 | 36.455 | 71.135 |
| 0 | 4.654 | 8.811 | 17.812 | 35.734 | 71.884 |

*Cpd.* **60***: Determination of* $IC_{50}$ *and* $k_1^*$

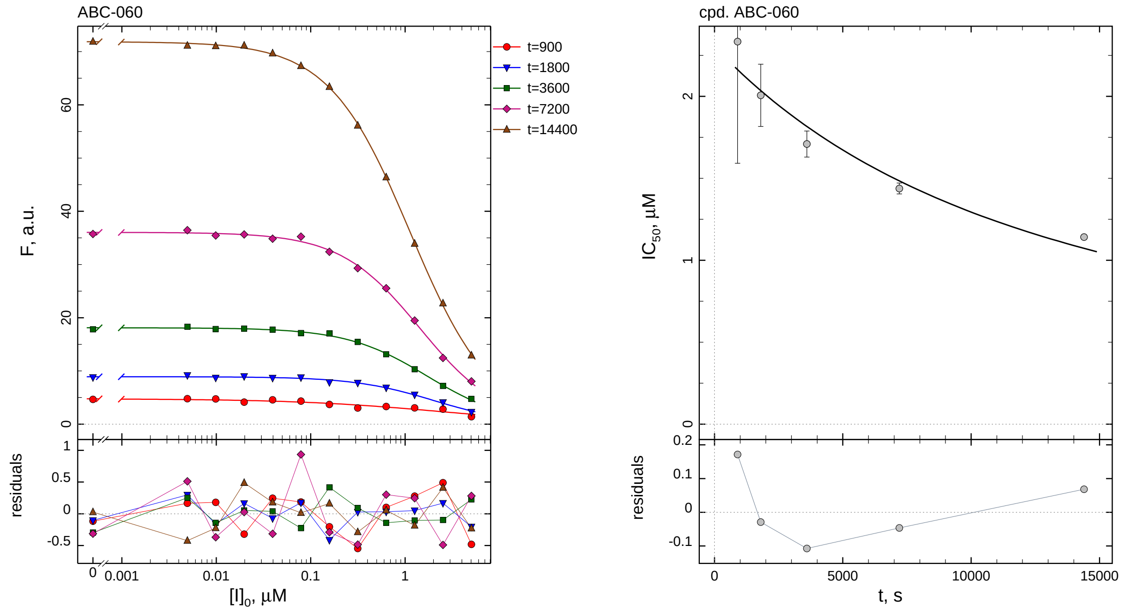

| $i$ | $t$, min | $I_{50}$, nM | ± | std.err. | CV, % | $k_1^*$, $mM^{-1}s^{-1}$ |
|---|---|---|---|---|---|---|
| 1 | 15 | 2334.4 | ± | 742.5 | 31.8 | 0.7585 |
| 2 | 30 | 2006.5 | ± | 190.0 | 9.5 | 0.4412 |
| 3 | 60 | 1709.3 | ± | 79.53 | 4.7 | 0.2590 |
| 4 | 120 | 1438.0 | ± | 33.25 | 2.3 | 0.1539 |
| 5 | 240 | 1141.4 | ± | 13.55 | 1.2 | 0.09696 |

*Cpd.* **60***: Mechanistic analysis*

| parameter | unit | "true" (T) | calculated (C) | C/T ratio | note |
|---|---|---|---|---|---|
| mechanism | | **C2S** | **C2F** | | |
| $k_{eff}$ | $mM^{-1}s^{-1}$ | 0.09901 | 0.1289 | 1.30 | $= k_{inact}/K_i$ |
| $K_i$ | nM | 1010 | 1155.6 | 1.14 | |
| $k_{inact}$ | $s^{-1}$ | 0.0001 | 0.000149 | 1.49 | |

| | |
|---|---|
| Reaction times used for analysis: | 30 and 120 min |
| Maximum GSD for accepting one-step model **C1**: | 1.25 |
| Observed GSD: | 2.11 |
| Assumed $[S]_0/K_M$ ratio: | 1.0 |

#### 2.61. *Compound No. 61*

*Cpd.* **61***: Simulated data*

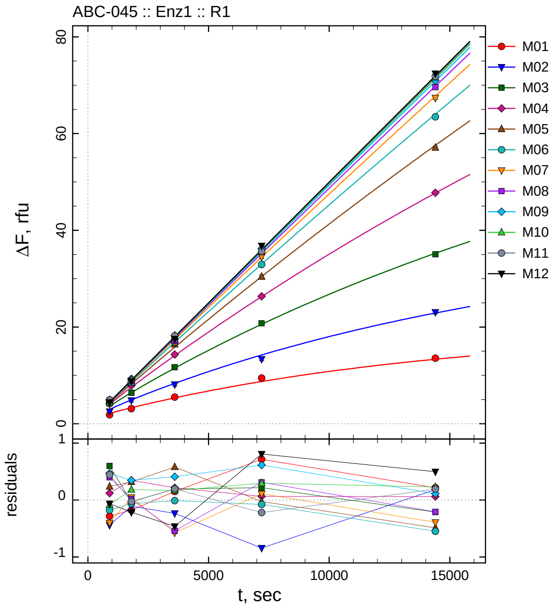

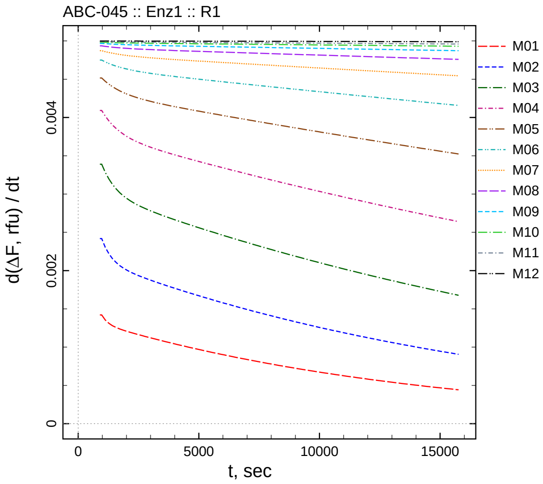

| $[I]_0$, nM | t = 15 | 30 | 60 | 120 | 240 min |
|---|---|---|---|---|---|
| 0.55 | 1.850 | 3.117 | 5.502 | 9.446 | 13.551 |
| 0.275 | 2.521 | 4.811 | 8.117 | 13.375 | 23.078 |
| 0.1375 | 4.202 | 6.401 | 11.701 | 20.779 | 35.034 |
| 0.06875 | 4.133 | 7.880 | 14.312 | 26.355 | 47.769 |
| 0.034375 | 4.482 | 8.529 | 16.413 | 30.411 | 57.081 |
| 0.0171875 | 4.192 | 8.527 | 16.841 | 32.920 | 63.480 |
| 0.00859375 | 4.031 | 8.837 | 16.833 | 34.542 | 67.383 |
| 0.00429688 | 4.867 | 8.908 | 17.156 | 35.504 | 69.592 |
| 0.00214844 | 4.951 | 9.292 | 18.252 | 36.197 | 70.964 |
| 0.00107422 | 4.407 | 9.154 | 18.086 | 36.077 | 71.618 |
| 0.000537109 | 4.940 | 8.957 | 18.152 | 35.662 | 71.854 |
| 0 | 4.432 | 8.783 | 17.532 | 36.789 | 72.431 |

*Cpd.* **61***: Determination of* $IC_{50}$ *and* $k_1^*$

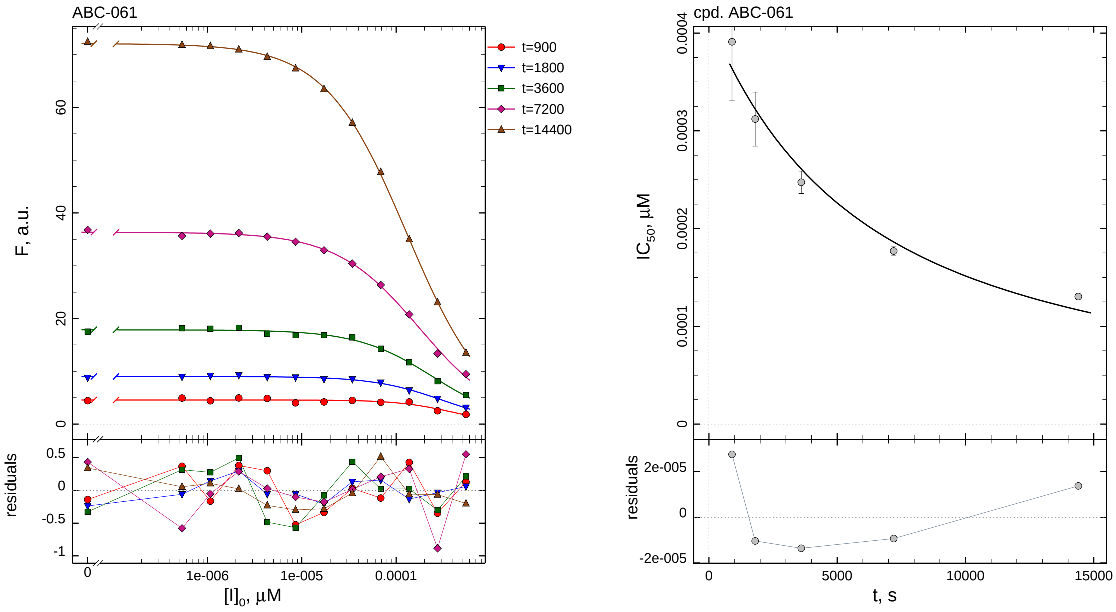

| $i$ | $t$, min | $I_{50}$, nM | ± | std.err. | CV, % | $k_1^*$, mM$^{-1}$s$^{-1}$ |
|---|---|---|---|---|---|---|
| 1 | 15 | 0.3911 | ± | 0.06042 | 15.4 | 4527.9 |
| 2 | 30 | 0.3120 | ± | 0.02767 | 8.9 | 2837.4 |
| 3 | 60 | 0.2473 | ± | 0.01144 | 4.6 | 1789.8 |
| 4 | 120 | 0.1770 | ± | 0.004241 | 2.4 | 1250.3 |
| 5 | 240 | 0.1304 | ± | 0.001568 | 1.2 | 848.4 |

*Cpd.* **61***: Mechanistic analysis*

| parameter | unit | "true" (T) | calculated (C) | C/T ratio | note |
|---|---|---|---|---|---|
| mechanism | | **C2S** | **C2F** | | |
| $k_{\mathrm{eff}}$ | mM$^{-1}$s$^{-1}$ | 909.1 | 1570.8 | 1.73 | $= k_{\mathrm{inact}}/K_{\mathrm{i}}$ |
| $K_{\mathrm{i}}$ | nM | 0.11 | 0.2092 | 1.90 | |
| $k_{\mathrm{inact}}$ | s$^{-1}$ | 0.0001 | 0.0003286 | 3.29 | |

| | |
|---|---|
| Reaction times used for analysis: | 30 and 120 min |
| Maximum GSD for accepting one-step model **C1**: | 1.25 |
| Observed GSD: | 1.79 |
| Assumed $[S]_0/K_M$ ratio: | 1.0 |

*2.62. Compound No. 62*

*Cpd.* **62***: Simulated data*

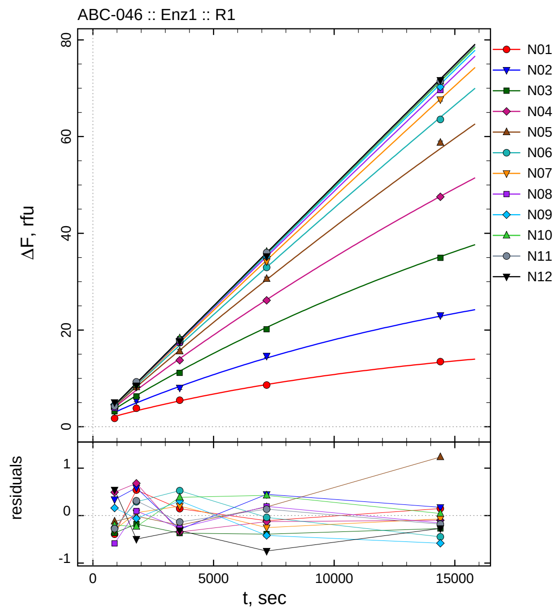

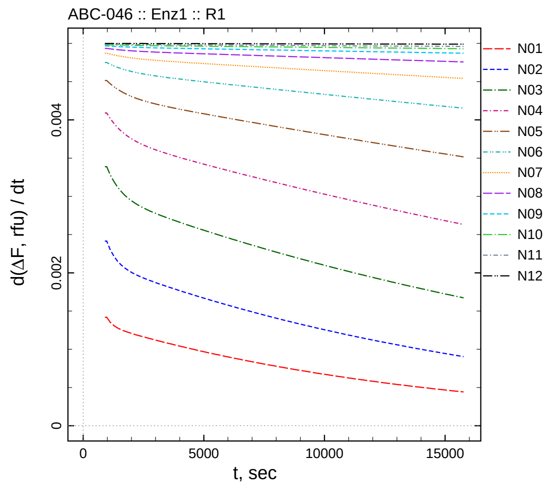

| $[I]_0$, nM | t = 15 | 30 | 60 | 120 | 240 min |
|---:|---:|---:|---:|---:|---:|
| 5.5 | 1.740 | 3.832 | 5.494 | 8.617 | 13.470 |
| 2.75 | 3.299 | 5.503 | 8.052 | 14.650 | 23.038 |
| 1.375 | 3.240 | 6.251 | 11.130 | 20.148 | 34.918 |
| 0.6875 | 4.505 | 8.213 | 13.745 | 26.140 | 47.544 |
| 0.34375 | 4.123 | 8.199 | 15.626 | 30.602 | 58.746 |
| 0.171875 | 4.104 | 8.880 | 17.375 | 32.950 | 63.547 |
| 0.0859375 | 4.133 | 8.851 | 17.604 | 34.173 | 67.683 |
| 0.0429688 | 3.883 | 8.988 | 17.435 | 35.383 | 69.631 |
| 0.0214844 | 4.643 | 8.887 | 18.155 | 35.165 | 70.264 |
| 0.0107422 | 4.291 | 8.740 | 18.305 | 36.206 | 71.430 |
| 0.00537109 | 4.219 | 9.296 | 17.823 | 36.018 | 71.479 |
| 0 | 5.041 | 8.503 | 17.680 | 35.238 | 71.664 |

*Cpd.* **62***: Determination of* $IC_{50}$ *and* $k_1^*$

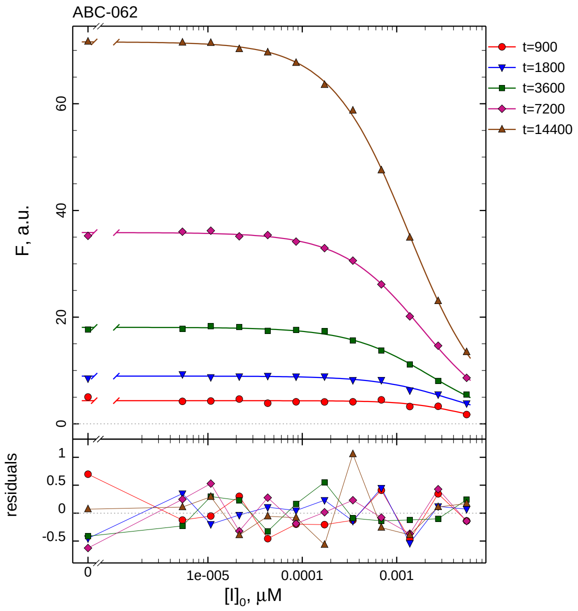

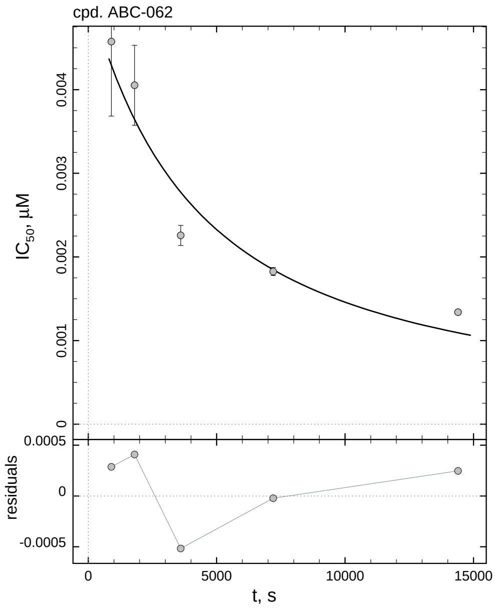

| $i$ | $t$, min | $I_{50}$, nM | ± | std.err. | CV, % | $k_1^*$, mM$^{-1}$s$^{-1}$ |
|---|---|---|---|---|---|---|
| 1 | 15 | 4.575 | ± | 0.8903 | 19.5 | 387.1 |
| 2 | 30 | 4.053 | ± | 0.4785 | 11.8 | 218.5 |
| 3 | 60 | 2.257 | ± | 0.1204 | 5.3 | 196.1 |
| 4 | 120 | 1.826 | ± | 0.04785 | 2.6 | 121.2 |
| 5 | 240 | 1.339 | ± | 0.01732 | 1.3 | 82.62 |

*Cpd.* **62***: Mechanistic analysis*

| parameter | unit | "true" (T) | calculated (C) | C/T ratio | note |
|---|---|---|---|---|---|
| mechanism | | **C2S** | **C2F** | | |
| $k_{\mathrm{eff}}$ | mM$^{-1}$s$^{-1}$ | 90.91 | 190.5 | 2.10 | $= k_{\mathrm{inact}}/K_{\mathrm{i}}$ |
| $K_{\mathrm{i}}$ | nM | 1.1 | 3.415 | 3.10 | |
| $k_{\mathrm{inact}}$ | s$^{-1}$ | 0.0001 | 0.0006505 | 6.50 | |

| | |
|---|---|
| Reaction times used for analysis: | 30 and 120 min |
| Maximum GSD for accepting one-step model **C1**: | 1.25 |
| Observed GSD: | 1.52 |
| Assumed $[S]_0/K_M$ ratio: | 1.0 |

*2.63. Compound No. 63*

*Cpd.* **63***: Simulated data*

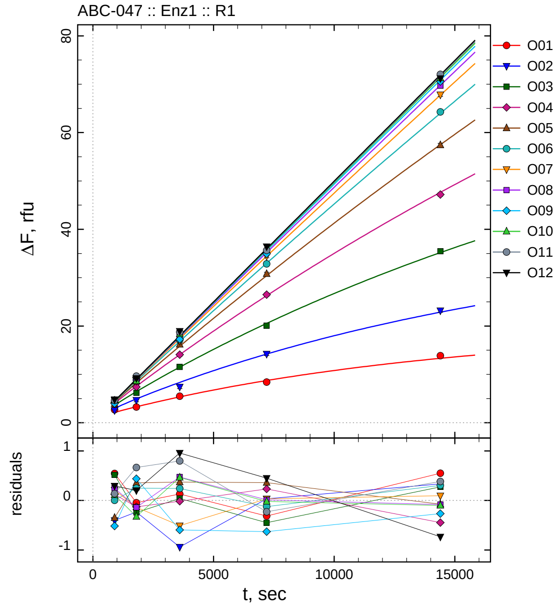

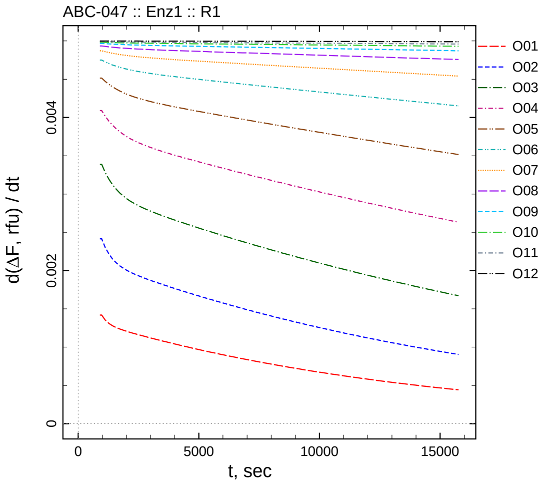

| $[I]_0$, nM | t = 15 | 30 | 60 | 120 | 240 min |
|---|---|---|---|---|---|
| 55 | 2.678 | 3.249 | 5.478 | 8.409 | 13.872 |
| 27.5 | 2.557 | 4.684 | 7.404 | 14.233 | 23.194 |
| 13.75 | 4.119 | 6.188 | 11.538 | 20.087 | 35.458 |
| 6.875 | 4.135 | 7.402 | 14.073 | 26.498 | 47.189 |
| 3.4375 | 3.896 | 8.566 | 16.199 | 30.778 | 57.432 |
| 1.71875 | 4.371 | 8.837 | 17.091 | 32.861 | 64.293 |
| 0.859375 | 4.642 | 8.654 | 16.896 | 34.459 | 67.846 |
| 0.429688 | 4.715 | 8.753 | 18.169 | 35.199 | 69.702 |
| 0.214844 | 3.966 | 9.383 | 17.250 | 34.952 | 70.580 |
| 0.107422 | 4.601 | 8.645 | 18.384 | 35.762 | 71.284 |
| 0.0537109 | 4.632 | 9.650 | 18.753 | 35.652 | 72.041 |
| 0 | 4.792 | 9.199 | 18.951 | 36.432 | 71.201 |

*Cpd.* **63***: Determination of* $IC_{50}$ *and* $k_1^*$

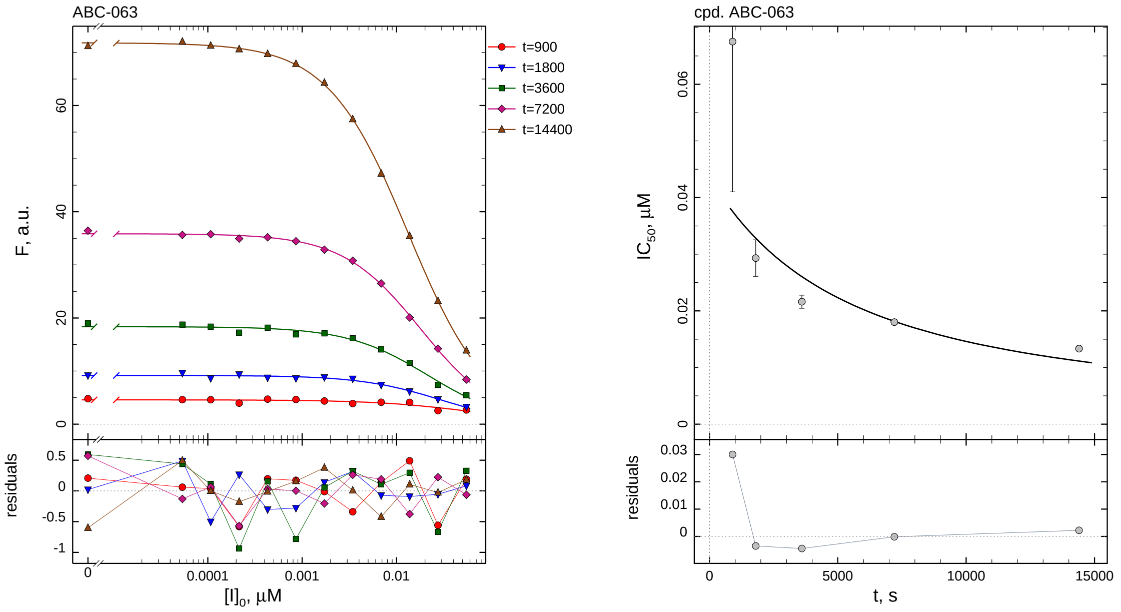

| $i$ | $t$, min | $I_{50}$, nM ± std.err. | CV, % | $k_1^*$, $mM^{-1}s^{-1}$ |
|---|---|---|---|---|
| 1 | 15 | 67.54 ± 26.53 | 39.3 | 26.22 |
| 2 | 30 | 29.33 ± 3.221 | 11.0 | 30.19 |
| 3 | 60 | 21.63 ± 1.155 | 5.3 | 20.47 |
| 4 | 120 | 18.02 ± 0.4733 | 2.6 | 12.28 |
| 5 | 240 | 13.34 ± 0.1796 | 1.3 | 8.293 |

*Cpd.* **63***: Mechanistic analysis*

| parameter | unit | "true" (T) | calculated (C) | C/T ratio | note |
|---|---|---|---|---|---|
| mechanism | | **C2S** | **C2F** | | |
| $k_{eff}$ | $mM^{-1}s^{-1}$ | 9.091 | 13.83 | 1.52 | $= k_{inact}/K_i$ |
| $K_i$ | nM | 11 | 18.54 | 1.69 | |
| $k_{inact}$ | $s^{-1}$ | 0.0001 | 0.0002563 | 2.56 | |

| | |
|---|---|
| Reaction times used for analysis: | 30 and 120 min |
| Maximum GSD for accepting one-step model **C1**: | 1.25 |
| Observed GSD: | 1.89 |
| Assumed $[S]_0/K_M$ ratio: | 1.0 |

*2.64. Compound No. 64*

*Cpd.* **64***: Simulated data*

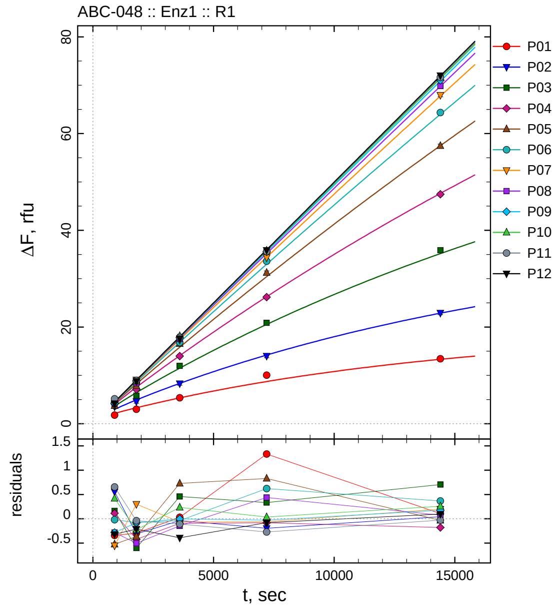

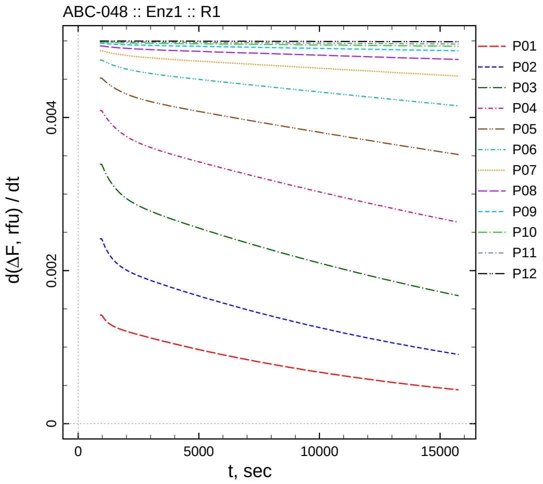

| $[I]_0$, nM | t = 15 | 30 | 60 | 120 | 240 min |
|---|---|---|---|---|---|
| 550 | 1.789 | 2.996 | 5.379 | 10.056 | 13.428 |
| 275 | 3.518 | 4.606 | 8.305 | 14.006 | 22.903 |
| 137.5 | 3.768 | 5.829 | 11.957 | 20.871 | 35.888 |
| 68.75 | 4.126 | 7.110 | 13.993 | 26.183 | 47.456 |
| 34.375 | 3.721 | 7.848 | 16.555 | 31.249 | 57.475 |
| 17.1875 | 4.350 | 8.513 | 16.813 | 33.607 | 64.355 |
| 8.59375 | 3.882 | 9.092 | 17.340 | 34.363 | 67.951 |
| 4.29688 | 4.183 | 8.391 | 17.551 | 35.626 | 69.863 |
| 2.14844 | 4.200 | 8.862 | 17.842 | 35.553 | 71.021 |
| 1.07422 | 4.910 | 8.743 | 18.155 | 35.818 | 71.644 |
| 0.537109 | 5.150 | 8.945 | 17.849 | 35.611 | 71.632 |
| 0 | 4.180 | 8.786 | 17.605 | 35.904 | 72.031 |

*Cpd.* **64***: Determination of* $IC_{50}$ *and* $k_1^*$

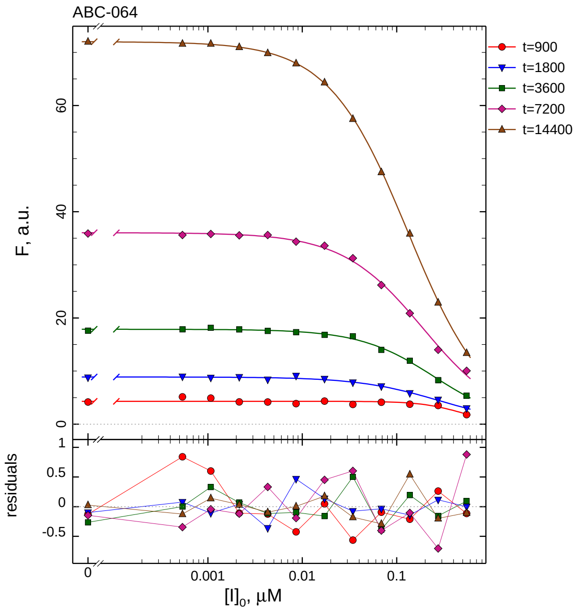

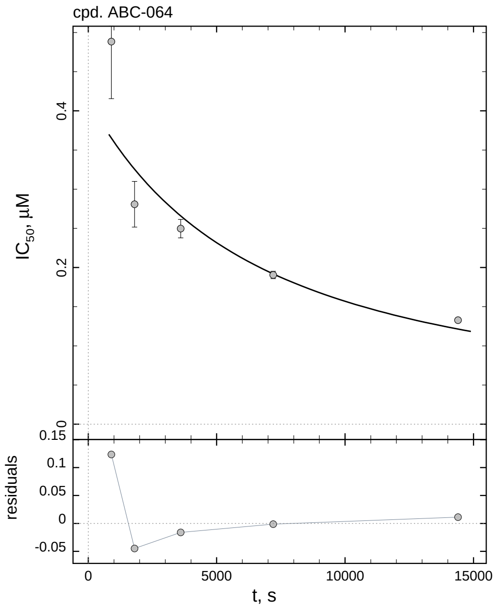

| $i$ | $t$, min | $I_{50}$, nM | ± | std.err. | CV, % | $k_1^*$, $mM^{-1}s^{-1}$ |
|---|---|---|---|---|---|---|
| 1 | 15 | 488.4 | ± | 72.85 | 14.9 | 3.625 |
| 2 | 30 | 280.8 | ± | 29.18 | 10.4 | 3.153 |
| 3 | 60 | 249.7 | ± | 11.78 | 4.7 | 1.773 |
| 4 | 120 | 190.5 | ± | 4.678 | 2.5 | 1.162 |
| 5 | 240 | 132.7 | ± | 1.613 | 1.2 | 0.8339 |

*Cpd.* **64***: Mechanistic analysis*

| parameter | unit | "true" (T) | calculated (C) | C/T ratio | note |
|---|---|---|---|---|---|
| mechanism | | **C2S** | **C2F** | | |
| $k_{eff}$ | $mM^{-1}s^{-1}$ | 0.9091 | 1.099 | 1.21 | $= k_{inact}/K_i$ |
| $K_i$ | nM | 110 | 166.7 | 1.52 | |
| $k_{inact}$ | $s^{-1}$ | 0.0001 | 0.0001832 | 1.83 | |

| | |
|---|---|
| Reaction times used for analysis: | 30 and 120 min |
| Maximum GSD for accepting one-step model **C1**: | 1.25 |
| Observed GSD: | 2.03 |
| Assumed $[S]_0/K_M$ ratio: | 1.0 |
