## Supplementary material for "A two-point IC_50_ method for evaluating the biochemical potency of irreversible enzyme inhibitors": Simulated data: notes.txt

In file names located in directory 'data', the notation 
"I01", "I02", ..., "I64" represents compound numbers.

A file named "simtr-I(NN)-R1.csv", where (NN) is 01, 02, ... 64,
contains simulated fluorescence intensities vs. inhibitor
concentrations at times t = 15, 30, ... 120 min.

A file named "ic50-I(NN)-R1.csv" contains the corresponding 
IC50 values determined by a fit of simulated dose-response 
curves to the three-parameter logistic equation.
